# Global Phylogeny of the Brassicaceae Provides Important Insights into Gene Discordance

**DOI:** 10.1101/2022.09.01.506188

**Authors:** Kasper P. Hendriks, Christiane Kiefer, Ihsan A. Al-Shehbaz, C. Donovan Bailey, Alex Hooft van Huysduynen, Lachezar A. Nikolov, Lars Nauheimer, Alexandre R. Zuntini, Dmitry A. German, Andreas Franzke, Marcus A. Koch, Martin A. Lysak, Óscar Toro-Núñez, Barış Özüdoğru, Vanessa R. Invernón, Nora Walden, Olivier Maurin, Nikolai M. Hay, Philip Shushkov, Terezie Mandáková, Mats Thulin, Michael D. Windham, Ivana Rešetnik, Stanislav Španiel, Elfy Ly, J. Chris Pires, Alex Harkess, Barbara Neuffer, Robert Vogt, Christian Bräuchler, Heimo Rainer, Steven B. Janssens, Michaela Schmull, Alan Forrest, Alessia Guggisberg, Sue Zmarzty, Brendan J. Lepschi, Neville Scarlett, Fred W. Stauffer, Ines Schönberger, Peter Heenan, William J. Baker, Félix Forest, Klaus Mummenhoff, Frederic Lens

**Affiliations:** Department of Biology, Botany, University of Osnabrück, Barbarastraße 11, D-49076 Osnabrück, Germany; Naturalis Biodiversity Center, Leiden University, PO Box 9517, Leiden, 2300 RA, The Netherlands; Centre for Organismal Studies (COS) Heidelberg, Department of Biodiversity and Plant Systematics, Heidelberg University, Im Neuenheimer Feld 345, 69120 Heidelberg, Germany; Missouri Botanical Garden, St. Louis, MO 63166-0299, USA; Department of Biology, New Mexico State University, Las Cruces, NM, USA; Department of Biology, University of Antwerp, Universiteitsplein 1, 2610, Antwerpen, Belgium; Department of Molecular, Cell and Developmental Biology and Molecular Biology Institute, University of California, Los Angeles, Los Angeles CA 90095, USA; Australian Tropical Herbarium, James Cook University, McGregor Road, Smithfield, Queensland 4878, Australia; Royal Botanic Gardens, Kew, Richmond, Surrey, TW9 3AE, United Kingdom; South-Siberian Botanical Garden, Altai State University, Lenin Ave. 61, 656049 Barnaul, Russia; CEITEC - Central European Institute of Technology and NCBR, Faculty of Science, Masaryk University, Kamenice 5, Brno, 625 00, Czech Republic; Departamento de Botánica, Facultad de Ciencias Naturales y Oceanográficas, Universidad de Concepción, Concepción, Chile; Applied Biology Section, Department of Biology, Faculty of Science, Hacettepe University, Ankara, Turkey; Institut de Systématique, Évolution, Biodiversité (ISYEB), Muséum National d’Histoire Naturelle, Sorbonne Université, École Pratique des Hautes Études, CNRS, Université des Antilles, 57 Rue Cuvier, CP 39, 75005, Paris, France; Department of Biology, Box 90338, Duke University, Durham, NC 27708, USA; Department of Chemistry, Indiana University, 800 E Kirkwood Ave, Bloomington, IN 47405, USA; Systematic Biology, Department of Organismal Biology, EBC, Uppsala University, Norbyvägen 18D, SE-752 36 Uppsala, Sweden; Department of Biology, Faculty of Science, University of Zagreb, Marulićev trg 20/II, HR-1000 Zagreb, Croatia; Institute of Botany, Plant Science and Biodiversity Centre, Slovak Academy of Sciences, Dúbravská cesta 9, SK-845 23 Bratislava, Slovakia; Wetsus, European Centre of Excellence for Sustainable Water Technology, Leeuwarden, The Netherlands; Department of Biotechnology, Delft University of Technology, Delft, The Netherlands; Biological Sciences, Bond Life Sciences Center, University of Missouri, Columbia, MO 65211-7310, USA; HudsonAlpha Institute for Biotechnology, Huntsville, AL, 35806, USA; Botanischer Garten und Botanisches Museum, Freie Universität Berlin, Königin-Luise-Str. 6-8, 14195 Berlin, Germany; Department of Botany, Natural History Museum Vienna, Burgring 7, 1010, Vienna, Austria; Department of Biology, KU Leuven, Kasteelpark Arenberg 31, BE-3001 Leuven, Belgium; Meise Botanic Garden, Nieuwelaan 38, BE-1860 Meise, Belgium; Harvard University Herbaria, 22 Divinity Ave, Cambridge, MA 02138, USA; Centre for Middle Eastern Plants, Royal Botanic Garden Edinburgh, 20a Inverleith Row, Edinburgh EH3 5LR, Scotland, United Kingdom; Institut für Integrative Biologie, ETH Zürich, Universitätstrasse 16, 8092 Zürich, Switzerland; Australian National Herbarium, Centre for Australian National Biodiversity Research, GPO Box 1700, Canberra, ACT, 2601, Australia; La Trobe University, Melbourne Victoria 3086, Australia; Department of Botany and Plant Biology, University of Geneva, Geneva, Switzerland; Allan Herbarium (CHR) Manaaki Whenua Landcare Research, Lincoln, New Zealand; Institute of Biology Leiden, Plant Sciences, Leiden University, Sylviusweg 72, 2333 BE Leiden, The Netherlands

**Author notes:** Corresponding authors* Kasper P. Hendriks, Klaus Mummenhoff, Frederic Lens.

**Keywords:** mustard family, Tree of Life, Angiosperms353, coalescence, phylogenomics, taxonomy

## Abstract

The mustard family (Brassicaceae) is a scientifically and economically important family, containing the model plant *Arabidopsis thaliana* and numerous crop species that feed billions worldwide. Despite its relevance, most published family phylogenies are incompletely sampled, generally contain massive polytomies, and/or show incongruent topologies between datasets. Here, we present the most complete Brassicaceae genus-level family phylogenies to date (Brassicaceae Tree of Life, or BrassiToL) based on nuclear (>1,000 genes, almost all 349 genera and 53 tribes) and plastome (60 genes, 79% of the genera, all tribes) data. We found cytonuclear discordance between nuclear and plastome-derived phylogenies, which is likely a result of rampant hybridisation among closely and more distantly related species, and highlight rogue taxa. To evaluate the impact of this rampant hybridisation on the nuclear phylogeny reconstruction, we performed four different sampling routines that increasingly removed variable data and likely paralogs. Our resulting cleaned subset of 297 nuclear genes revealed high support for the tribes, while support for the main lineages remained relatively low. Calibration based on the 20 most clock-like nuclear genes suggests a late Eocene to late Oligocene ‘icehouse origin’ of the family. Finally, we propose five new or re-established tribes, including the recognition of Arabidopsideae, a monotypic tribe to accommodate *Arabidopsis*. With a worldwide community of thousands of researchers working on this family, our new, densely sampled family phylogeny will be an indispensable tool to further highlight Brassicaceae as an excellent model family for studies on biodiversity and plant biology.

## Introduction

The mustard family (Brassicaceae) is a medium-sized plant family with huge economic and scientific impact. The family contains numerous crop species grown for food and biofuel, such as cabbage, cauliflower, mustard, cress, rapeseed and pennycress, as well as many model species, which have provided fundamental insights into plant biology, including *Arabidopsis thaliana and A. lyrata*, *Arabis alpina, Brassica* spp., and *Capsella* spp. (Franzke et al. 2011; Warwick 2011; Jabeen 2020). Moreover, the overwhelming availability of genetic resources in these species make Brassicaceae an ideal model family in flowering plants, facilitating, among others, plant developmental studies that disentangle genotype-phenotype interactions (Nikolov and Tsiantis 2017; Rahimi et al. 2022), investigate impacts of differently aged whole genome duplications (Hohmann et al. 2015; Mandáková and Lysak 2018), and unravel the variation in evolutionary pathways leading to a huge diversity in secondary metabolites that fuel the plant-pollinator or plant-herbivore arms-race (Franzke et al. 2011; Kiefer et al. 2014; Edger et al. 2015). In many of these genetic studies, comparative studies to extend what is known from model species to close or more distant allies require a robust evolutionary framework encompassing the ~4,000 currently accepted species that are divided among 349 genera and 50-60 tribes (Koch et al. 2018). This is also true for studies on the wild relatives of crops that aim to deliberately introgress desirable traits (e.g., increased drought tolerance, disease resistance) from closely related species in the wild into crop species using plant breeding (Castañeda-Álvarez et al. 2016; Castillo-Lorenzo et al. 2019; Quezada-Martinez et al. 2021). It is unsurprising then that the taxonomy and evolution of Brassicaceae have been the subject of study for a long time, with the ultimate goal of producing a complete and robust family phylogeny (Al-Shehbaz et al. 2006; Bailey et al. 2006; Franzke et al. 2009; Beilstein et al. 2010; Couvreur et al. 2010; Warwick et al. 2010; Huang et al. 2016; Nikolov et al. 2019; Huang et al. 2020; Walden et al. 2020).

Despite various seminal contributions pushing forward the Brassicaceae Tree of Life (hereafter named BrassiTol), a robust, densely sampled family phylogeny is still lacking. First attempts using molecular markers in the 1980s showed that earlier phylogenetic inferences, which mainly relied on flower and fruit traits (von Hayek 1911; Schulz 1919; Janchen 1942), were often misled by rampant convergent evolution (Koch et al. 2003; Mitchell-Olds et al. 2005; Koch and Mummenhoff 2006). Similarly, the use of one to few molecular markers generated debatable, poorly-supported phylogenies that were unlikely to resemble species phylogenies and intrinsically suffered from unresolved deeper nodes leading to uninformative polytomies (e.g., Koch et al. 2001; Al-Shehbaz et al. 2006; Bailey et al. 2006). This has traditionally been attributed to an early rapid radiation in the family (Couvreur et al. 2010), combined with multiple whole genome duplication and hybridisation events (Franzke et al. 2011). Although it has been suggested that the use of more genes is the best way forward to increase resolution and nodal support in a phylogenetic tree (Rokas and Carroll 2005), early multi-gene (phylogenomic) work also showed that discordance among different genes may reduce nodal support (Jeffroy et al. 2006). More recent family-wide phylogenomic studies in Brassicaceae offered support for three (Koch and Al-Shehbaz 2009), four (Franzke et al. 2011), or five (Huang et al. 2016; Nikolov et al. 2019; Liu et al. 2020; Mabry et al. 2020; Beric et al. 2021) main lineages in addition to tribe Aethionemeae, sister to the rest of Brassicaceae, which is a likely result of the generally low support for the deeper nodes based on measures like gene discordance. In addition to the traditional explanation why the deeper Brassicaceae nodes are poorly supported, more recent studies on causes for this gene discordance are focusing on among others incomplete lineage sorting (ILS), ancient hybridization and introgression (Forsythe et al. 2020), (ancient) inter-tribal and even inter-lineage hybridization (Mandáková, Pouch, et al. 2017; Guo et al. 2020), whole-genome duplication (WGD) and post-polyploid diploidization (PPD), and complex combinations of the above (Maddison 1997; Yue et al. 2009; German et al. 2011; Mandáková, Li, et al. 2017; Mandáková and Lysak 2018; Dogan et al. 2021). To make things more complicated, there is a general mismatch in the position and resolution of the deeper (main lineages) and shallower (tribal) nodes between phylogenies constructed from nuclear and plastome data, which most likely reflects the different evolutionary trajectories of the nuclear genome and the maternally-inherited plastome (Forsythe et al. 2020; Mabry et al. 2020; Walden et al. 2020).

Although debates about the best way to reconstruct species trees are ongoing, there is increasing evidence that large genomic datasets representing hundreds or even thousands of genes (e.g., transcriptomes) are the way forward (Rannala and Yang 2008). They allow the reconstruction of the evolutionary histories of different parts of the genome using multiple gene trees, and permit an understanding of the landscape of conflict and concordance among different genes and gene histories (Smith et al. 2018). Importantly, more complex genomic datasets also allow for more informative species trees and nodal support, for instance by using gene and site concordance factors (Degnan and Rosenberg 2009; Sayyari and Mirarab 2016; Minh, Hahn, et al. 2020), compared to traditional bootstrap values that are often artificially high (100%) in large datasets. Furthermore, genomic datasets also enable tracing the histories of gene duplications, which can facilitate orthology assessment (Zhang et al. 2020). Over the past 10 years, novel methodologies in reduced-representation sequencing have generated these genomic datasets, with target capture sequencing (or hyb-seq) turning out to be the most influential in the field of phylogenomics (Lemmon and Lemmon 2013; Dodsworth et al. 2019). This method offers several important advantages over full genome sequencing and other modern sequencing methods, notably a combination of broad taxonomic application with high efficiency on non-model species, broad phylogenetic informativeness at both deep and shallow nodes, and cost-efficiency (Lemmon and Lemmon 2013). Because the method relies on short DNA sequences (200-400 bp), may work with very low DNA starting amount (down to 5 ng), and is rather insensitive to DNA degradation, it is also ideally suited to work with (up to centuries-old) material from natural history museums, allowing access to rare and previously ‘locked’ specimens (Särkinen et al. 2012; Hart et al. 2016; Brewer et al. 2019; Forrest et al. 2019; Alsos et al. 2020; Bakker et al. 2020; Folk et al. 2021; Kates et al. 2021). In addition, target capture sequencing of leaves generates a considerable sequence bycatch in the form of off-target reads (‘genome skimming’) that represent a wealth of chloroplast genes, allowing simultaneous reconstruction of the plastome (Weitemier et al. 2014).

Here, we present results from the largest global Brassicaceae phylogenetic study to date, with the following five objectives: (1) reconstruct the most complete and robust family-wide BrassiToL based on nuclear genomic data (using a mixed baits approach containing over 1,000 genes) and plastome data; (2) investigate the influence of paralogous genes and polyploid species on phylogeny reconstruction; (3) explore cytonuclear discordance between the nuclear and plastome-derived phylogenies; (4) reevaluate the time of origin of the Brassicaceae and its main lineages and tribes; and (5) assess delimitation of tribes and major lineages. This improved phylogenetic framework will set the stage to further develop Brassicaceae as a model family in flowering plants.

## Results

### Target capture sequencing datasets

Our nuclear family ingroup sampling included 380 samples, covering 375 species, 319 of the 349 currently accepted genera (91%) and 51 of the 52 accepted tribes (or 57 of 58 after inclusion of newly proposed tribes, see below; excluding only the monotypic Hillielleae). In addition, we sampled 23 outgroup species representing all Brassicales families (table 1, supplementary tables S1-2) and 14 Brassicaceae genera formerly considered valid taxa, but now treated as synonymous to one of the 349 accepted genera. Most of the 30 genera missing from our dataset could either not be sampled or failed during the laboratory process; of these, three genera (*Hilliella*, *Onuris*, and *Thelypodium*) were included in the plastome phylogeny, though (supplementary table S3).

**Table 1.** Definition of sampling routines and threshold settings in HybPhaser used to select nuclear genes and subsequently gene trees in the coalescent approach to reconstruct the nuclear species tree, with ASTRAL-III output results.

| Sampling routine | inclusive | strict | superstrict | superstrict by tribe |
| --- | --- | --- | --- | --- |
| **Gene and sample recovery thresholds** |  |  |  |  |
| Gene minimum proportion of samples recovered | 0.1 | 0.2 | 0.2 | 0.2 |
| Gene minimum proportion of target length recovered | 0.1 | 0.2 | 0.2 | 0.2 |
| Sample minimum proportion of total target length recovered | 0.0 | 0.4 | 0.4 | 0.4 |
| Sample minimum proportion of genes recovered | 0.0 | 0.2 | 0.2 | 0.2 |
| Gene SNPs proportion threshold for all samples | none | outliers | 0.02 | 0.02 |
| Remove outlier genes per sample? | yes | yes | yes | yes |
| **Results** |  |  |  |  |
| Number of genes retained | 1,018 | 1,013 | 297 | 1,031* |
| Number of samples | 380 | 361 | 361 | 361 |
| Mean sample occupancy across all genes retained | 818.5 | 861.5 | 243.9 | 394.3 |
| Number of species | 375 | 356 | 356 | 356 |
| Number of genera | 332 | 317 | 317 | 317 |
| Number of tribes ** | 57 | 56 | 56 | 56 |
| LPP (mean of all nodes) | 0.94 | 0.95 | 0.92 | 0.88 |
| LPP (median of all nodes) | 1 | 1 | 1 | 1 |
| First quartile proportion (mean of all nodes) | 0.55 | 0.56 | 0.57 | 0.56 |
| First quartile proportion (median of all nodes) | 0.48 | 0.50 | 0.50 | 0.49 |

\* Mean number across samples, with genes retained different in different tribes (i.e., tribe-specific). As a result, more genes could be retained in total as compared to the 'inclusive' routine, but at a lower mean sample occupancy across all genes, thus leading to a relatively sparse matrix.
\*\* Tribal definition following taxonomic insights presented in this study, i.e. 58 tribes in total.

We recovered a total of 1,081 unique nuclear genes (supplementary fig. S1) across 397 samples (supplementary table S4). For the Brassicaceae specific bait set containing 764 nuclear genes (Nikolov et al. 2019; hereafter B764), we recovered on average (across all 397 samples) 91% of the total target length of 919,712 bp, and 84% of the 764 genes targeted. Using the PAFTOL bait set containing 353 nuclear genes (Johnson et al. 2019; William J. Baker et al. 2021; hereafter A353) average numbers were 83% of the total target length of 263,894 bp and 80% of 353 genes targeted. This indicates that recovery success was comparable between the two bait sets, as previously also found by Hendriks et al. (2021), but slightly higher for the B764 bait set. As expected, gene recovery was on average better for samples belonging to the Brassicaceae family than for the non-Brassicaceae outgroups, with on average 85% of the total target length of 1183,606 bp (from the two baits sets combined) recovered for Brassicaceae-samples vs. 41% for outgroup-samples, and 79% of the 1,081 targeted genes for the Brassicaceae-samples vs. 23% for outgroup-samples. Surprisingly, gene recovery hardly decreased with sample age, and we successfully obtained hundreds of nuclear genes, even from pre-1900 samples (supplementary fig. S2).

### Nuclear Brassicaceae phylogeny

We used paralog detection in HybPhaser v2.0 (Nauheimer et al. 2021) to define four ‘sampling routines’ (‘inclusive’, ‘strict’, ‘superstrict’, and ‘superstrict by tribe’) to select different nuclear gene subsets based on gene SNP proportions to reconstruct the mustard family phylogeny (table 1; details in supplementary table S4 and fig. S3-S4). In addition, we used both supermatrix maximum likelihood (ML) and coalescent approaches in ASTRAL-III (Zhang et al. 2018) and ASTRAL-Pro (Zhang et al. 2020) on these datasets to infer the species phylogeny (see Materials and Methods for details).

Node support was high for most tribes, regardless of the sampling routine or phylogenetic approach (fig. 1; details in supplementary table S4 and fig. S5-S11). This provides strong support for the monophyly of many (but not all) of the tribes, although it needs to be noted that some tribes were previously defined based on molecular studies only. From the supermatrix approach (concatenation of 297 target genes; total length 700,445 bp; 82.7% complete), the mean of bootstrap (BS) values across tribal nodes was 98%, the mean of gene concordance factors (gCF) was 47%, and the mean of site concordance factors (sCF) was 62%. From the coalescent approach (‘strict’ routine), the mean local posterior probability (LPP) across tribal nodes was 98% and support for the first quartet (Q1) 73%. Tribes Biscutelleae, Brassiceae, Microlepidieae, Subularieae, and Thelypodieae had an gCF<10%. Tribes Camelineae and Iberideae were polyphyletic in all nuclear phylogenies (supplementary fig. S5-S11).

**Fig. 1.**
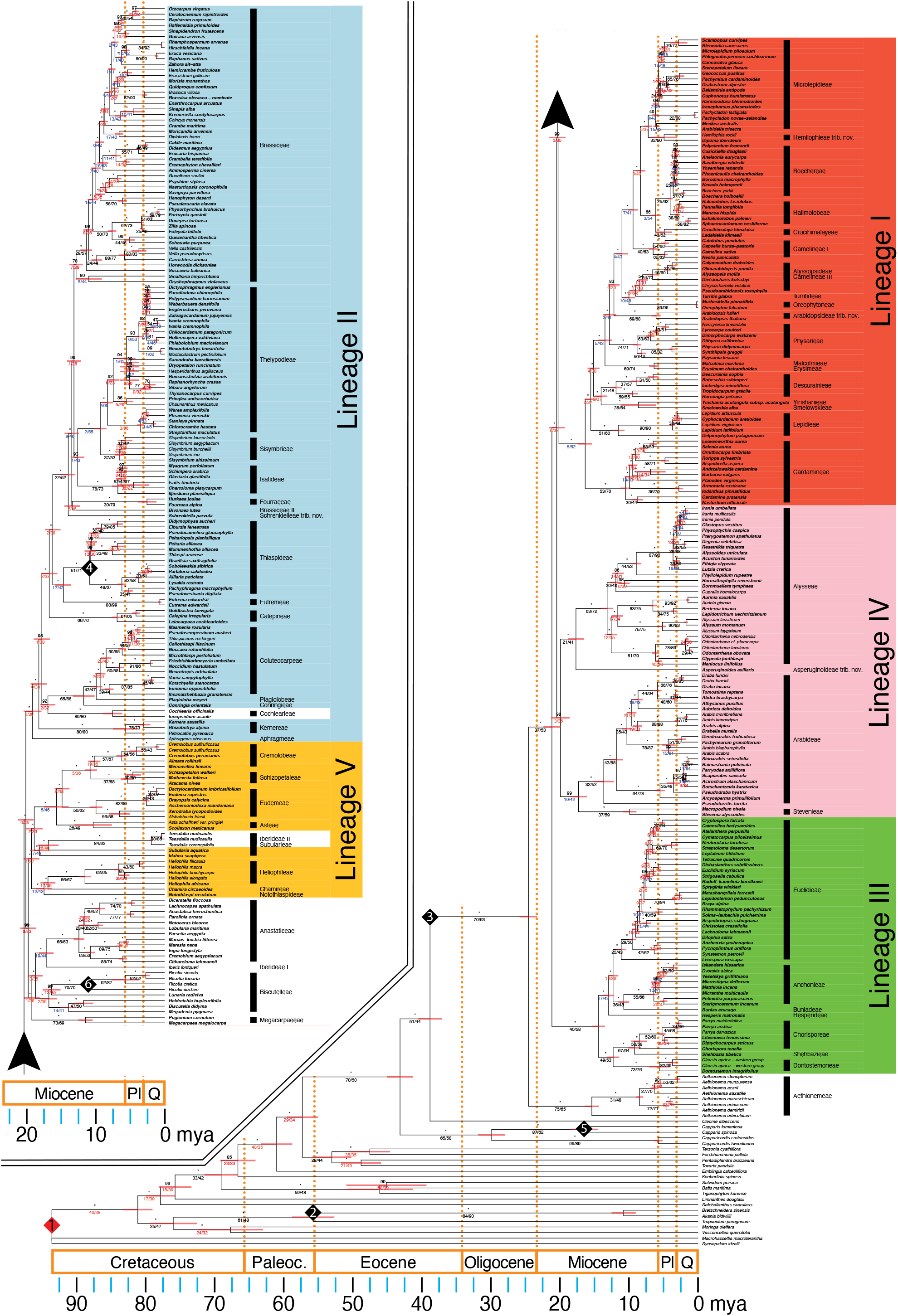
Mustard family genus-level phylogeny from a maximum likelihood analysis on a 297 nuclear genes supermatrix (‘superstrict routine’) with node calibration of 90 mya at the stem of order Brassicales. Genus type species are highlighted in bold. Numbers in diamonds refer to calibration points or checks listed in table 2 (note that not all numbers could be fitted into the plot).

Based on the nuclear dataset, we consistently found tribe Aethionemeae sister to the rest of the Brassicaceae (hereafter defined as core Brassicaceae), main lineage III sister to a clade of lineages I+II+IV+V, and lineages II+V sharing a common ancestor (fig. 2a, b), regardless of sampling routine and/or phylogenetic inference approach, as recovered previously (Nikolov et al. 2019). Whereas support for the split between tribe Aethionemeae and the core Brassicaceae was high (BS: 100%; gCF: 70%; sCF: 63%; LPP: 100%; Q1: 70%), support for the five remaining main lineages varied. Lineage III received medium support (100%; 40%; 58%; 100%; 91%, respectively), but support for the remaining main lineages was poor (supplementary table S5) in terms of gCF (ranging from 1 to 5%) and sCF (ranging from 36 to 58%) values. Remarkably, BS and LPP values for these main lineages were (nearly) always 100%, indicating that both values are poor estimators of node support in phylogenies based on hundreds of markers. The generally low support for the phylogenetic backbone was reflected in the general topological differences among the family phylogenies based on the different sampling routines (fig. 2a-c), and was visualised as a large, complex reticulate core in our split network analysis based on a supermatrix of the nuclear genes included in the ‘superstrict’ routine (fig. 2d). A more inclusive approach resulted in placement of main lineage I as sister to a clade of main lineages II+IV+V (fig. 2a). The stricter routines of the nuclear dataset showed a consistent placement of lineage IV as sister to a clade of lineages I+II+V (fig. 2b) in both the ML and coalescent approaches. This showed that removing a first set of most variable genes (‘strict’ routine) considerably impacts the backbone of the topology, whereas an additional removal of less variable genes (‘superstrict’, ‘superstrict by tribe’) has little effect.

**Table 2.**
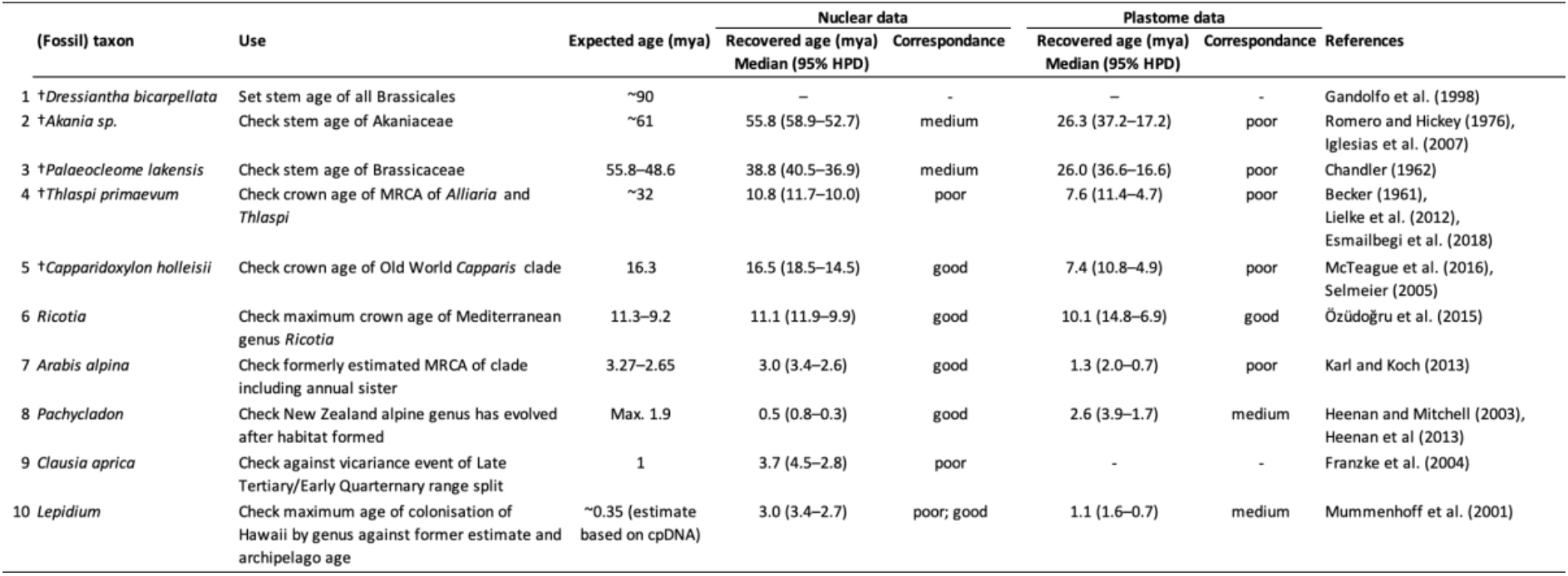
Data used for fossil calibration of mustard family phylogeny (1), fossils to check calibration (2-5), and biogeographical validation (6-10) of our new results. Case 9 could not be checked in the plastome phylogeny, because not all the populations of *Clausia aprica* were included in our sampling.

**Fig. 2.**
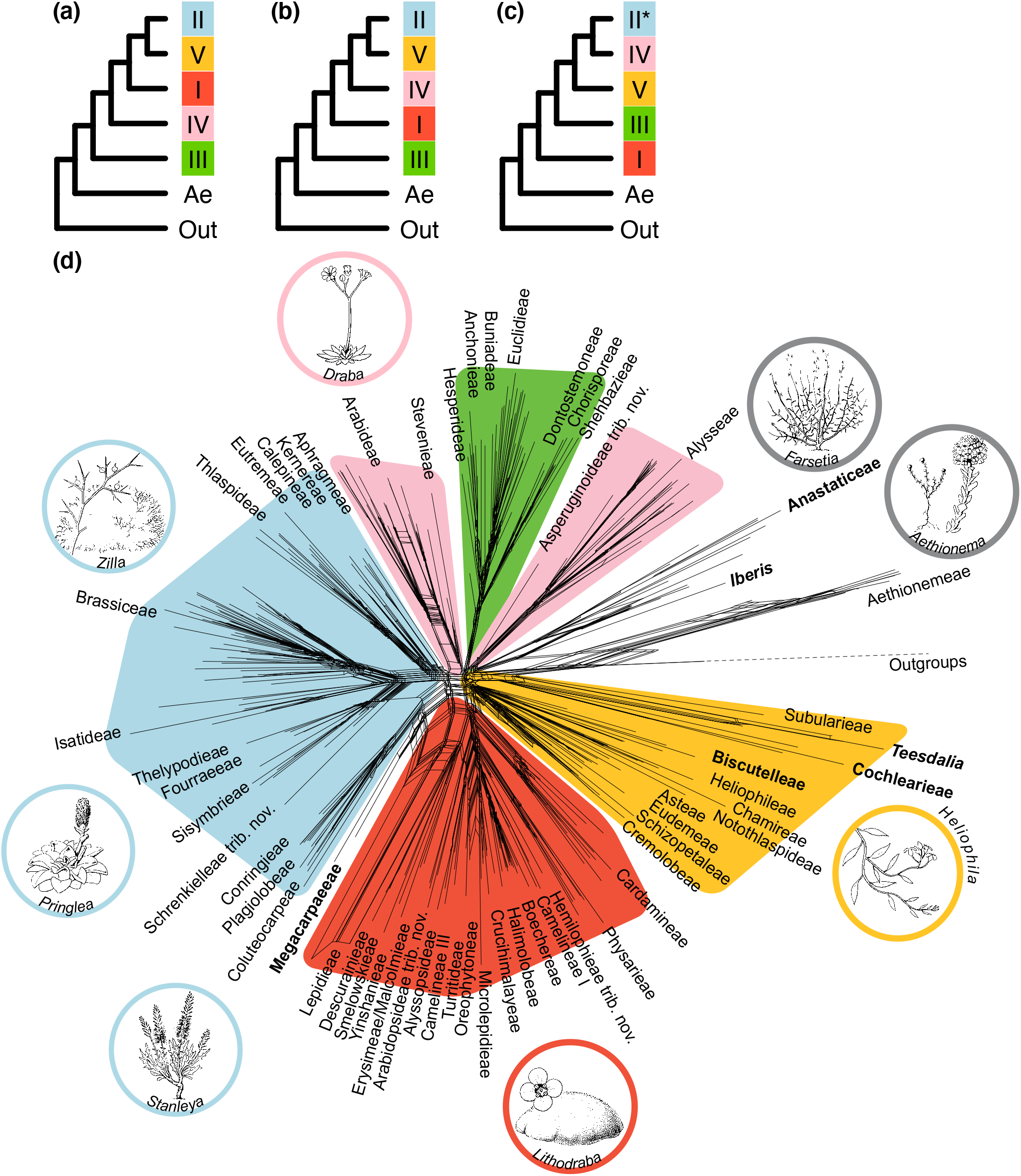
Overview of the main mustard family relationships from different phylogenetic reconstruction routines and approaches. Colours follow the main core Brassicaceae lineages (I–V) described by Nikolov, et al. (2019). (a-c) Cladograms showing topology of main lineages as recovered from (a) nuclear ‘inclusive’ routine and coalescent approach (using either ASTRAL-III and ASTRAL-Pro), (b) nuclear stricter routines (‘strict’, ‘superstrict’, and ‘superstrict by tribe’) using either supermatrix ML and coalescent approaches (ASTRAL-III), and (c) plastome supermatrix ML approach. The * indicates polyphyly in main lineage II in the plastome supermatrix ML phylogeny; the clade drawn represents most genera. (d) Split network of the mustard family tribes computed from uncorrected p-distances on a supermatrix of the nuclear genes covered by the ‘superstrict’ routine, covering 317 genera of 56 tribes. The network highlights the complex reticulate evolution both in the ancestors of extant main lineages, as well as within some of the main lineages themselves. Names in bold highlight rogue tribes as described in the main text. Insets show the high morphological diversity within the family, which contains growth forms such as tiny herbaceous, frutescent, and woody species, as well as lianas, shrubs and cushion plants. Drawings by Esmée Winkel, Naturalis Biodiversity Center.

We used Townsend’s phylogenetic informativeness (Townsend 2007) to quantify the impact of each of the 1,081 nuclear genes on the final species tree (supplementary fig. S12). Variation in informativeness from genes varied greatly, with mean values suggesting that genes from the B764 bait set were more informative than those of the A353 bait set, for genes with both low and high mean SNP proportions. This is expected to be the result of the B764 bait set being designed specifically for the mustard family, thereby capturing more of the family-specific genetic variation.

### Plastome Brassicaceae phylogeny

Our plastome family ingroup sampling included 502 samples, covering 438 species, 266 genera (76%, plus 10 Brassicaceae genera formerly considered valid taxa, but now synonymous to one of the 349 accepted genera) and all 58 tribes (following the new tribal delimitation suggested in this paper; see below).

Plastome tribal support was mostly high, similar to the nuclear phylogeny, with the mean of site concordance factors 78% and the mean of bootstrap (BS) values across tribal nodes 100% (supplementary tables S4-S5). As in the nuclear phylogeny, tribe Aethionemeae was sister to all remaining Brassicaceae (fig. 3). All tribes that had been assigned to lineage II in the nuclear phylogeny also formed one large clade in the plastome phylogeny. Contrary to the nuclear phylogeny, lineage I (not III) was sister to all remaining lineages (II–V), with lineage III sister to a clade formed by lineages II+IV+V. Whereas each of the main lineages formed a monophyletic group in the nuclear phylogeny, lineage II was polyphyletic in the plastome phylogeny, with several tribes (Aphragmeae, Coluteocarpeae, Conringieae, Kernereae, and Plagiolobeae) recovered within lineage V. Similarly, tribe Stevenieae, recovered within lineage IV in the nuclear phylogeny, was recovered within lineage I in the plastome phylogeny. The newly suggested tribe Asperuginoideae, assigned to lineage IV in the nuclear phylogeny, formed a clade with Biscutelleae, and these tribes together were sister to all other remaining lineages (II+IV+V).

**Fig. 3.**
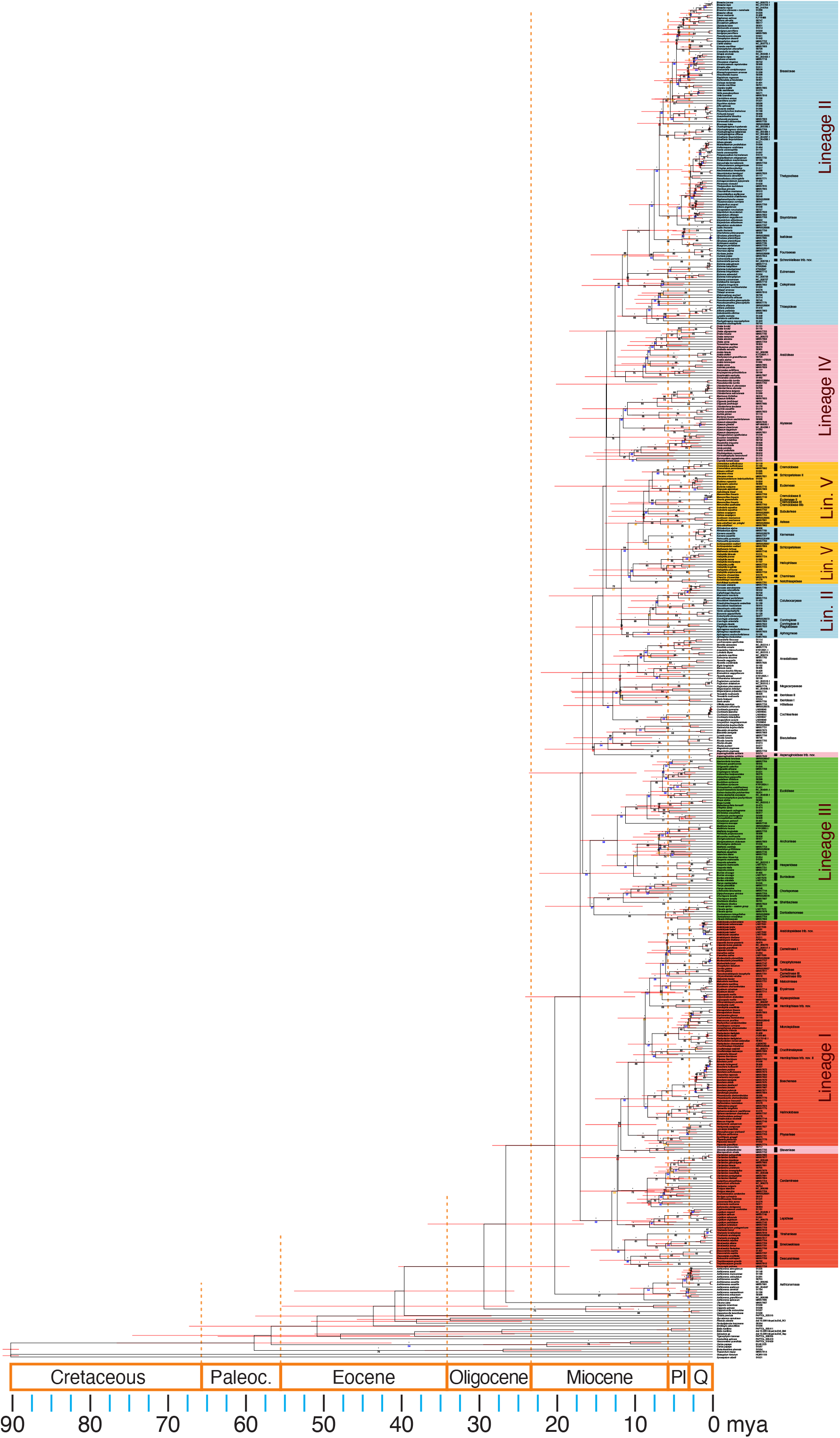
Mustard family genus-level phylogeny from a maximum likelihood analysis on a 60 plastome genes supermatrix with node calibration of 90 mya at the stem of order Brassicales. Genus type species are high-lighted in bold.

### Cytonuclear discordance

Brassicaceae family phylogenies derived from nuclear and plastome data show marked differences in topology (fig. 1 vs. fig. 3). We used a Procrustean Approach to Cophylogeny (PACo; Balbuena et al. 2013) to detect phylogenetic incongruences between nuclear and plastome supermatrix ML species trees, each first pruned to tribal level by selecting a random sample from each tribe (fig. 4a). The global PACo test of the hypothesis, stating that the similarity between the trees is not higher than expected by chance (Balbuena et al. 2013), was strongly rejected (*p*<0.001, m^2^=0.203, 1,000 permutations). However, there were clear incongruences in tribal positions between the two phylogenies (fig. 4b), where tribes in lineage II showed least incongruence (3 out of 13 tribes), and tribes in lineage IV most incongruence (3 out of 4 tribes).

**Fig. 4.**
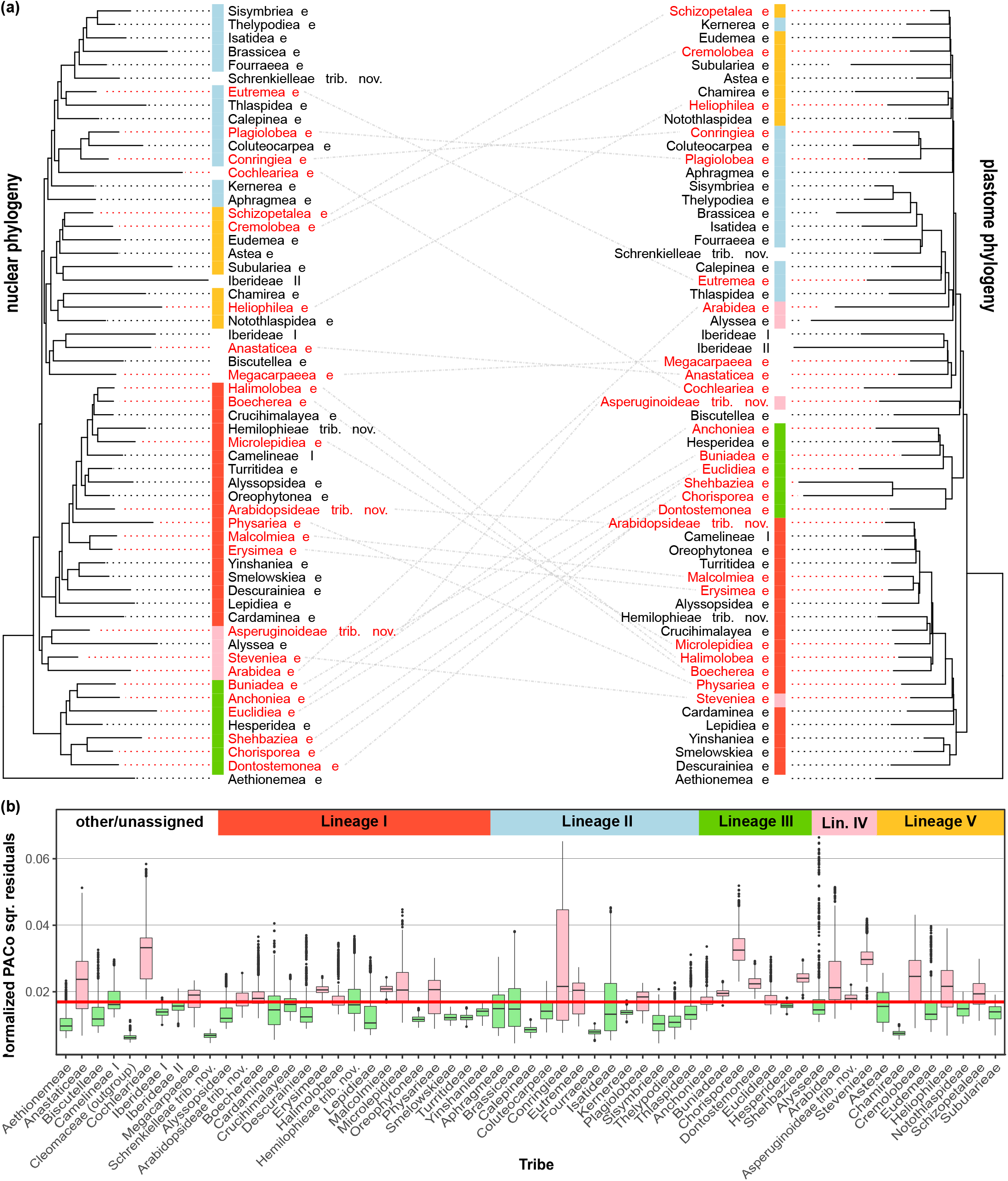
Results from Procrustean Approach to Cophylogeny (PACo) comparing the topologies of the nuclear and plastome-derived Brassicaceae family phylogenies at the tribe level. (a) Nuclear phylogeny (left) and plastome phylogeny (right), with incongruent tribes highlighted in red (corresponding to pink bars in panel b), and dashed lines connecting the incongruent tribes. Coloured bars represent the main lineages cf. (Nikolov et al. 2019). (b) Boxplot of normalised squared residual values 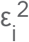 from individual nuclear-plastome associations using phylograms from 1,000 bootstrap replicates. The horizontal red line equals 1/n=0.0169, where *n*=59 is the number of nuclear-plastome tribe-associations, which equals the number of tribes under consideration. Median values above this threshold are expected to be linked to tribes that show incongruence between nuclear and plastome-derived trees, and associated boxes are highlighted in pink.

### Indications of polyploidy and rogue taxa

We assessed allelic variation using HybPhaser v2.0 (Nauheimer et al. 2021) to calculate the proportions of loci with divergent alleles (% locus heterozygosity, LH) and average divergence between alleles (% allele divergence, AD), using the 1,013 nuclear genes included in the ‘strict’ routine (ignoring 38 samples with a coverage too low to calculate LH and AD; table 1 and supplementary table S4). While most samples fell into the expected range of ‘normal’ species (roughly LH <90% and AD <1), a number of samples showed signs of polyploidization (fig. 5; supplementary fig. S13). We found mean values for both LH and AD to be very high at 75.1% and 2.25%, respectively, with no clear difference between genes from the B764 and A353 bait sets (supplementary table S6).

**Fig. 5.**
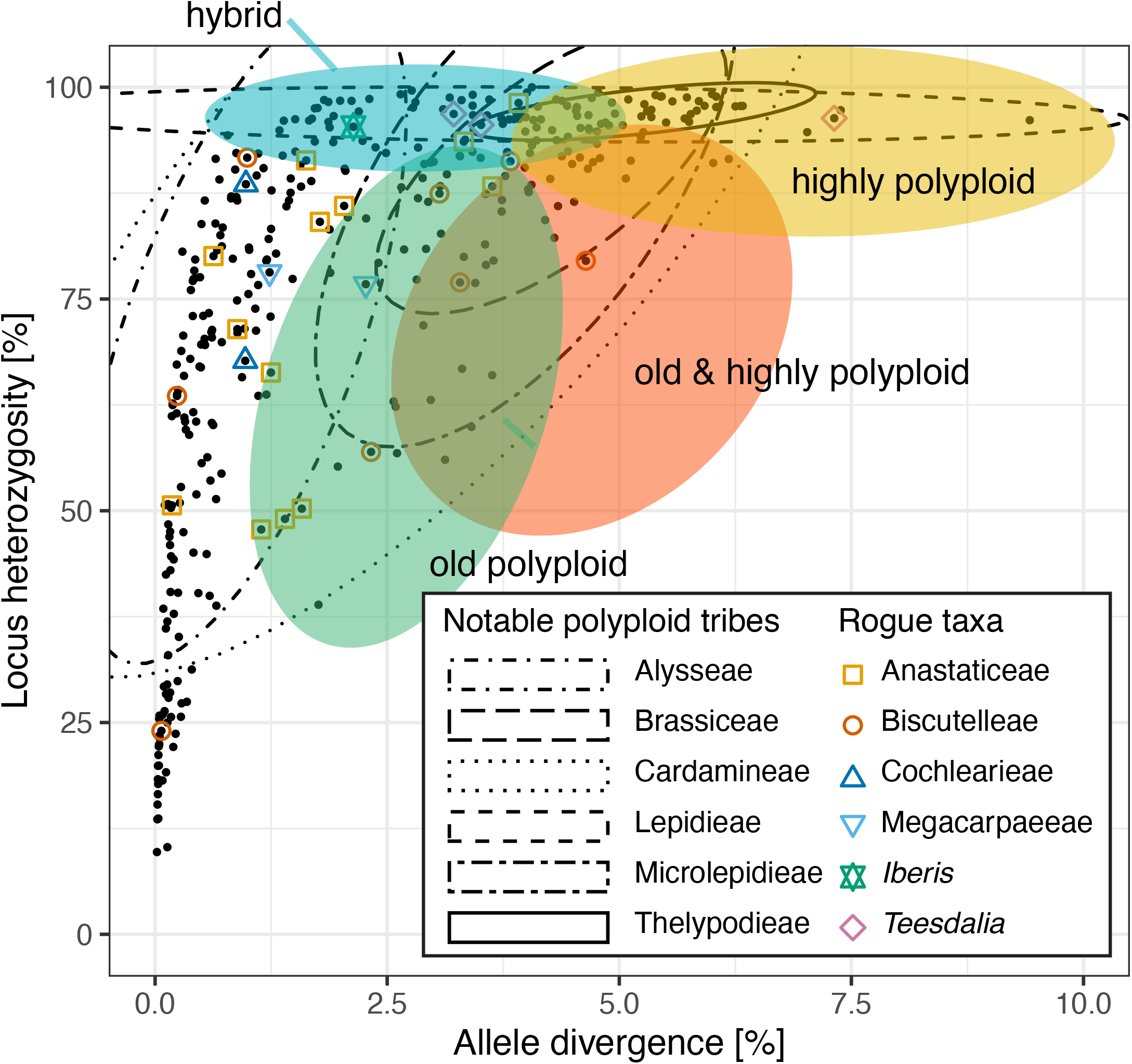
Scatterplot displaying the locus heterozygosity and allele divergence of all samples included in the ‘strict’ routine. Coloured ovals indicate four classes in the expected ‘realm of polyploids’, rough estimates of what could be considered likely hybrids, polyploids, and their ages (see main text for the rationale behind assigning these different classes). The overlap between these classes highlights the uncertainty. The six largest tribes (represented by >10 samples) for which (most) of their samples fall within this realm of polyploids are annotated using ellipses showing the 95% confidence level for a multivariate *t*-distribution of their data. Samples from rogue taxa are highlighted using different shapes and colours.

Based on the list of LH and AD values for all species, we tentatively distinguished four family-specific classes: ‘hybrid’, ‘old polyploid’, ‘highly polyploid’, and ‘old & highly polyploid’ (fig. 5; see Materials and Methods for details and class definition). Nearly all representatives of tribe Thelypodieae fell within the ‘highly polyploid’ class (with high LH and very high AD values), meaning that it is likely that the group experienced one or multiple recent polyploidization events leading to many neopolyploid species (see also supplementary tables S3 and S6). Similarly, all representatives of tribe Lepidieae fell within the ‘hybrid’ or ‘highly polyploid’ classes. Also, tribes Alysseae, Brassiceae, Cardamineae, and Microlepidieae had the bulk of their representatives in one of the polyploid classes (fig. 5).

Our phylogenetic results highlighted several jumpy or ‘rogue taxa’, mostly at the tribal level, that were recovered in different positions in the various species trees that resulted from the different routines and/or approaches. Most importantly, in the nuclear supermatrix ML approach and the stricter coalescent approaches (supplementary fig. S14a and d-f), Anastaticeae, Biscutelleae, and Megacarpaeeae formed a single clade sister to lineage V, whereas in the ‘inclusive’ coalescent approaches (ASTRAL-III and ASTRAL-Pro; supplementary fig. S14b-c), Megacarpaeeae was sister to Anastaticeae, with these tribes together sister to lineage II. However, the position of tribe Biscutelleae changed again depending on the coalescent approach used (ASTRAL-III vs. ASTRAL-Pro). Species that belong to one of the rogue tribes generally also show relatively high LH and AD values (17 out of 29 cases fall well within the ‘realm of polyploids’, with another 6 cases bordering it; fig. 5), suggesting their polyploid origin.

### Fossil calibration and tribal age estimates

We used the Turonian *Dressiantha bicarpellata* fossil, dated at 89.3–93.6 mya (Gandolfo et al. 1998), to calibrate the stem node of order Brassicales in both nuclear and plastome supermatrix ML phylogenies (fig. 1 and 3). To validate our results, we confronted our dating estimates with nine expected dates from fossil and biogeographical events (table 2), as suggested by Franzke et al. (2016). In general, results from our nuclear phylogeny supported age estimates, or were younger. In the plastome phylogeny, our age estimates were significantly younger, except for the following cases. We found good corroboration with age estimates for the Miocene *Capparidoxylon holleisii* fossil (16.3 mya; Selmeier 2005) in the nuclear phylogeny, the maximum crown age of the Mediterranean genus *Ricotia* (11.3–9.2 mya; Özüdoğru et al. 2015) in both phylogenies, the formerly estimated age of the most recent common ancestor (mrca) of the *Arabis alpina* clade (3.27–2.65 mya; Karl and Koch 2013) in the nuclear phylogeny, and the maximum age of the New Zealand alpine genus *Pachycladon* (max 1.9 mya; Heenan and Mitchell 2003; Heenan and McGlone 2013) in both phylogenies. The support for the Paleocene fossil *Akania* sp. (~61 mya; Romero and Hickey 1976) was medium in the nuclear phylogeny, for the early Tertiary *Palaeocleome lakensis* fossil (55.8–48.6 mya; Chandler 1962) medium (nuclear) to poor (plastome phylogeny), and poor for the vicariance event in *Clausia aprica* as well (Franzke et al. 2004) in the nuclear phylogeny. Corroboration with the Brassicaceae *Thlaspi primaevum* fossil (~32 mya; Becker 1961; Lielke et al. 2012) was poor (node estimated at 10.8 mya). However, the identification of this fossil was heavily debated (Franzke et al. 2016).

Based on our nuclear dataset, we found that the core Brassicales (Brassicaceae + Cleomaceae) and Capparaceae split around 43.2 mya (95% HPD: 45.0–41.1), in the middle Eocene (fig. 1). Subsequently, the Brassicaceae split from sister family Cleomaceae around 38.8 mya (95% HPD: 40.5–36.9), in the middle to late Eocene (table 3). The core Brassicaceae split from tribe Aethionemeae around 24.5 mya (95% HPD: 25.7–23.1), in the late Oligocene. All five main lineages originated early to middle Miocene (median stem ages between 21.2 and 19.8 mya; median crown ages between 19.9 and 14.4 mya; supplementary table S5). Mean median stem age across tribes (supplementary table S5) was 12.1 mya (max. 18.9 mya, Subularieae; min. 4.0 mya, Boechereae; *n*=52), while mean median crown age was 8.0 mya (max. 18.1 mya, Subularieae; min. 0.2 mya, Oreophytoneae; *n*=37). Results from our plastome dataset were much younger than those from our nuclear dataset, with a family crown age of 20.2 mya (95% HPD: 29.0–13.0) and a core Brassicaceae crown age of 16.9 mya (95% HPD: 24.3–10.2; supplementary table S5).

**Table 3.**
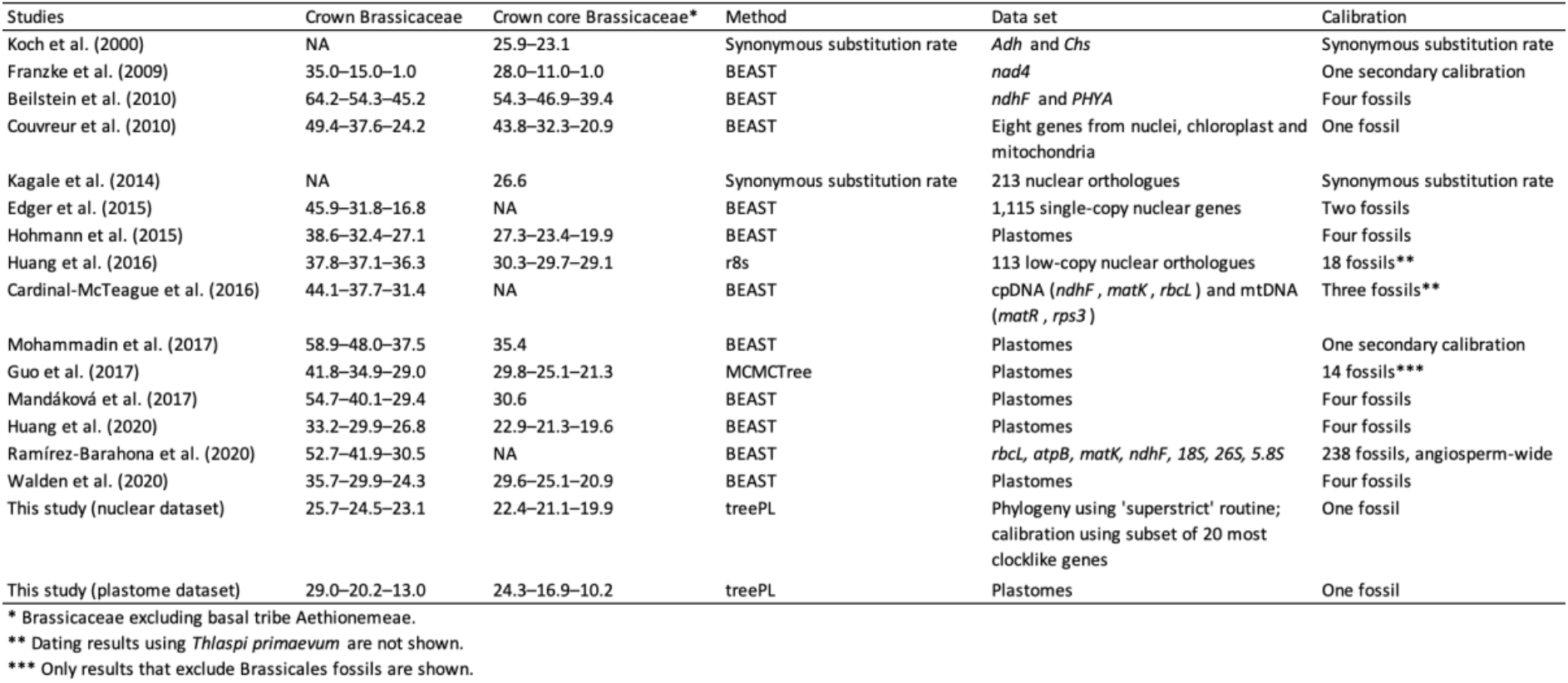
Comparison of divergence time estimates (mya) for Brassicaceae from past studies and new results from our study. Table copied from (Huang et al. 2020) and updated with more recent results and results from this study.

### Tribal taxonomic revisions

The Brassicaceae, one of the top 15 families of angiosperms in terms of number of species, are a family with a relatively large number of recognised tribes per species as compared to some of the other large angiosperm families (e.g., Asteraceae, Orchidaceae, Poaceae, and Fabaceae) (Koch et al. 2018). Nevertheless, based on our new phylogenetic framework, together with an analysis of former literature and comparative morphology, we find good support for the formal recognition of five more tribes (see also Discussion). With respect to *Arabidopsis*, our results agree with previous findings Huang et al. (2016) and Nikolov et al. (2019) that support the movement of this genus into a new monotypic, primarily Eurasian tribe, Arabidopsideae (lineage I), since it is not closely related to other genera of Camelineae, the tribe to which it was previously assigned. Furthermore, we propose the following taxonomic changes: erecting the monospecific and distinct genus *Asperuginoides* into its own new tribe, Asperuginoideae; combining the genera *Dipoma* and *Hemilophia* in the new tribe Hemilophieae; combining the genera *Idahoa* and *Subularia* in the re-established tribe Subularieae; and raising the monospecific and distinct genus *Schrenkiella* into its own tribe, Schrenkielleae.

Finally, genus *Iberis* requires further taxonomic study due to unexpected recovery in our nuclear phylogenies. Contrary to our plastome phylogeny and previous findings (Warwick et al. 2010), we recovered it as sister to tribe Anastaticeae, and not as sister to the other genus in tribe Iberideae, *Teesdalia*, which instead consistently grouped with main lineage II (but in various positions; supporting fig. S14).

## Discussion

### Nuclear and plastome genus-level phylogenies of the Brassicaceae

We present here the most complete nuclear and plastome-derived Brassicaceae Trees of Life (BrassiToLs) to date, together covering nearly all genera from which plant tissue is currently accessible, i.e., 92% of the 349 genera, representing all 58 tribes (following our suggested taxonomic revision). Such global genus-level BrassiToLs were long overdue, and are expected to become an indispensable tool for among Brassicaceae experts who want to understand the evolutionary patterns and processes that shaped biodiversity, and to extend the mechanistic insights derived from the study of model species to all major lineages of the mustard family, including the many food and oilseed crops.

The overall backbone of our nuclear BrassiToL largely agrees with the nuclear phylogeny published by (Nikolov et al. 2019), which included 63 species representing 50 out of the 52 tribes then recognised. More specifically, our nuclear BrassiToL confirms (1) the recognition of tribe Aethionemeae as sister to the rest of the Brassicaceae (i.e., core Brassicaceae), which includes the same five main lineages (I–V) as proposed by Nikolov et al. (2019), and (2) main lineage III (containing tribes Anchonieae, Buniadeae, Chorisporeae, Dontostemoneae, Euclidieae, Hesperideae, and Shehbazieae) as sister to the rest of the core Brassicaceae. This confirmation is not surprising, given that 71% of target genes in our total dataset were included in the Brassicaceae-specific bait set (Nikolov et al. 2019). The basal split of tribe Aethionemeae followed by lineage III was also supported by the recently published transcriptomic phylogenies of Mabry et al. (2020) and Beric et al. (2021) based on a much more reduced Brassicaceae sampling (43 genera). When comparing our results with the transcriptomic study of Huang et al. (2016), relationships among tribes were similar, but their main lineages A–E only roughly corresponded to our main lineages I–V. This highlights the well-known problem of recovering the deeper nodes in the BrassiToL backbone with great certainty, which we attempt to further disentangle given the uncertainties that still persist (see below). Whereas the gene selection in Huang et al. (2016) was based on low-copy occurrence among widely different angiosperms, comparable to the Angiosperms353 dataset (Johnson et al. 2019), the majority of genes we used were instead based on putatively single-copy loci among the Brassicaceae (Nikolov et al. 2019).

When comparing our plastome phylogeny (265 currently accepted genera, all 58 tribes) with the nuclear phylogeny (319 currently accepted genera, 57 tribes), we found strong cytonuclear discordance, as previously demonstrated by (Nikolov et al. 2019; Mabry et al. 2020). At a higher taxonomic level, tribe Aethionemeae remained sister to all remaining Brassicaceae in the plastome phylogeny, but lineage I (not III) was sister to all remaining lineages (II–V), with lineage III being sister to the rest. Moreover, lineage III is the only major lineage in the core Brassicaceae that turned out to be monophyletic, while the four others are either paraphyletic (lineage I and V) or polyphyletic (lineage IV and especially lineage II; fig. 3). At a lower taxonomic level, such discordance was further emphasised, as has also been recently studied in more detail for *Arabidopsis* and several close relatives (Forsythe et al. 2020; Guo et al. 2020), who showed that (ancient) hybridization and introgression are causing these topological incongruencies. Furthermore, Guo et al. (2021) showed that tribe Biscutelleae harbours four genus-specific WGDs based on hybridization among the same and/or closely related parental genomes. Then, despite the incidence of multiple independent WGDs, diploid and polyploid genomes may be retrieved as a monophyletic clade characterised by cytonuclear discordance. Our PACo test on species tree incongruence showed that, even though major lineages are placed in different positions across the BrassiToLs, the overall topologies are not significantly different.

When comparing our plastome phylogeny with the previously published plastome phylogeny of Walden et al. (2020; 152 genera, all 58 tribes), we find that tribal relationships within lineages I and III were very similar. However, the topology differed with respect to deeper nodes in lineages II, IV and V. Notably, tribe Alysseae (lineage IV) was sister to the clade containing tribes Arabideae and the largest clade of lineage II in our plastome phylogeny, while Walden et al. (2020) recovered tribe Alysseae as sister to most of the expanded lineage II (also including lineage IV-V).

### Toward solving the deeper nodes in the Brassicaceae backbone

Relationships at the shallower nodes are rather well-resolved in our phylogenies, leading to strong relationships among genera within tribes and among most of the Brassicaceae tribes (Q1 node support from the coalescent approach was >50% in 34 of 42 tribes; supplementary table S5). The nodes connecting the Brassicales families are generally well-resolved as well. However, the deeper nodes within the Brassicaceae family –reflecting the position of the five main lineages of the core Brassicaceae– have been notoriously hard to recover in the past. Some authors have claimed to have solved the true Brassicaceae backbone derived from an NGS dataset (Liu et al. 2020), but relied only on bootstrap and posterior probability values. However, we show that alternative sampling routines can recover high bootstrap and posterior probability values on conflicting topologies (supplementary table S5), questioning the value of these support metrics. Instead, metrics on node support should take underlying topological variation among gene trees and sites in the alignments into account to be more realistic, using for example gene and site concordance factors, something that has been shown in many recent large-scale phylogenomic studies (Minh, Hahn, et al. 2020). We show that the topology of the species tree backbone is highly sensitive to the selected (nuclear) gene set (supplementary fig. S14), and to a lesser extent also the phylogenetic reconstruction method applied.

Low backbone support in a phylogenetic tree can have several causes: (1) biological processes within the family such as gene duplication and loss, hybridisation, incomplete lineage sorting, and gene saturation, (2) one or more artefacts (e.g., deficient or erroneous data such as paralogs interpreted as orthologs), or (3) a combination thereof (Boussau and Scornavacca 2020). As gene saturation is a process that increases with time, we believe it is not influencing our dataset to a large extent, because node support among Brassicales families is generally higher than support for main lineages within Brassicaceae. In contrast, we found allele divergence and locus heterozygosity to be very high in many species of Brassicaceae, with mean values of 75.1% and 2.25%, respectively (supplementary tables S4 and S6). In comparison, these values were 52.3% and 0.21%, respectively, across all non-hybrid natural accessions of pitcher plants (*Nepenthes* spp.; Nauheimer et al. 2021), and 60.7% and 0.85% across a family-wide sampling of Australian Thelypterid ferns (Bloesch et al. 2022). This indicates the abundant presence of paralogs in our Brassicaceae dataset, likely caused by ample hybridisation and gene duplication events (fig. 5).

Indeed, within the Brassicaceae, many cases of hybridisation that are known to be rampant at different taxonomic levels have been described (see below), and it will be challenging to include such a wide variety of evolutionary oddballs into a single model that can recover the ‘true’ evolutionary history of the family, if possible at all; perhaps the phylogeny of the family cannot be described by a bifurcating tree, and a network approach represents these relationships more faithfully (Huson and Bryant 2006; fig. 2). Aware of the issues with data deficiency and paralogs, we specifically designed our analyses to try to disentangle some of the deeper nodes, by taking the following approaches into account: we first pruned samples with deficient data and likely paralogs (‘strict’: 1,013 genes, ‘superstrict’: 297 genes, and ‘superstrict by tribe routine’: 1,031 genes, but mean sample occupancy of 394 genes; table 1), and repeated the analysis explicitly including all gene copies (including possible paralogs) using ASTRAL-Pro.

In this regard, we advocate critically scrutinising all genes showing a SNP variation of >0.02, a value indicating the presence of paralogs, and in addition all samples with less than 20% of genes and/or 40% of recovered target length. This refined sampling approach corresponds to the ‘superstrict’ routine (fig. 1 and 2). Interestingly, we found that going from the ‘strict’ routine towards the ‘superstrict’ routine (see Materials and Methods for definitions), the phylogeny hardly changed (supplementary fig. S14). We therefore believe that the ‘superstrict’ routine provides the best estimate of the phylogeny, as it includes most samples and at the same time does not contain a whole set of extra genes that do not impact the topology nor node support. This is especially relevant in Brassicaceae due to the huge amount of SNP variation among the genes studied as a result of many polyploidisation events (see below), despite the fact that Brassicaceae specific B764 bait set was designed to only include single copy nuclear genes.

### Rogue taxa share a mesopolyploid past

Previous studies highlighted several ‘unplaced taxa’, including the tribes Anastaticeae, Biscutelleae, Cochlearieae, and Megacarpaeeae, and the genera *Iberis*, *Idahoa*, and *Subularia*, which have been difficult to resolve (Nikolov et al. 2019). Our results highlight the same ‘rogue taxa’. Contrary to the bulk of the tribes, which were consistently placed within a main lineage in our different nuclear phylogenies (i.e., from different routines and/or approaches), these rogue tribes ‘jumped’ positions across the different phylogenies (supplementary fig. S5-11, summarised in supplementary fig. S14). This has previously been interpreted as an indication of genus- or tribe-specific meso-polyploidisations, including distant (inter-tribal) hybridization, i.e., hybridisations between members of distantly related lineages.

The genus *Brassica* and the tribe Brassiceae were the first reported Brassicaceae taxa with a WGD that postdates the family-specific paleotetraploidization *At*-a event (e.g., Lysak et al. 2005; Parkin et al. 2005; Lysak et al. 2007). Since this pioneering work, more than a dozen genus- and tribe-specific mesopolyploid WGDs have been discovered throughout the Brassicaceae family tree (Hohmann et al. 2015; Mandáková, Li, et al. 2017; Guo et al. 2020; Dogan et al. 2022). Because most mesopolyploid taxa have an allopolyploid origin, often understood to result from (inter-tribal) hybridization between distantly related parental species (German and Friesen 2014; Mandáková, Pouch, et al. 2017; Guo et al. 2020), inferring their phylogenetic position within the family tree is a challenging endeavour. For all of the rogue taxa we identified, a mesopolyploid origin has been demonstrated or claimed. For tribe Cochlearieae, whole genome triplication has been demonstrated, whereas Anastaticeae and *Iberis* have a mesotetraploid origin (Mandáková, Li, et al. 2017). Biscutelleae harbour four different WGDs specific to *Biscutella*, *Heldreichia*, *Lunaria*, and *Ricotia* (Mandáková et al. 2018; Guo et al. 2020). Both genera of Megacarpaeeae (*Megacarpeae* and *Pugionium*) have been shown to be formed by independent WGDs (Yang et al. 2020; Hu et al. 2021). *Teesdalia* (2n=36 in *T. coronopifolia* and *T. nudicaulis*, 2n = 20 in *T. conferta*) most likely has a polyploid origin based on available chromosome numbers, but (cyto)genomic data are lacking.

### The Brassicaceae originated during an icehouse period

Based on our analysis of the 20 most clock-like nuclear genes, we recovered a middle to late Eocene stem age for the mustard family of 38.8 mya (95% HPD: 40.5–36.9), with a late Oligocene crown age of 24.5 mya (95% HPD: 25.7–23.1; fig. 1; table 3). Whereas results from previous family-wide studies ranged from 15.0 mya (Franzke et al. 2009) to 54.3 mya (Beilstein et al. 2010), more recent studies resulted in a converging family crown age of 37.1–32.4 mya, which seemed quite insensitive to data type (nuclear/plastome), methods, and fossils used (Huang et al. 2020). Ramírez-Barahona et al. (2020), in an angiosperm-wide study using 238 fossils, recovered a family crown age of 41.9 mya. Therefore, our new results suggest a somewhat younger age of the Brassicaceae than generally found. Consensus about the best approach to retrieve reliable dating estimates based on big genomic datasets is lacking, but our experience is that time calibrating analyses using a more restricted dataset that includes only the most clock-like genes outcompete those that use a less stringent approach or even the entire dataset.

The results from our most clock-like nuclear dataset suggest that the family’s origin and the onset of diversification coincides with the cooling of the Earth during the Eocene-Oligocene transition (so-called greenhouse to icehouse transition; Zanazzi et al. 2007; Sun et al. 2014). This period was characterised by a replacement of tropical forests with temperate forests, open vegetation and deserts, which all are associated with typical habitats of extant Brassicaceae. Tribe Aethionemeae and the other five main lineages originated quickly after (median stem ages ranging 21.2–19.8 mya; median crown ages ranging 19.9–14.4 mya; supplementary table S5) in the early Miocene.

Results from our plastome phylogeny generally show a ~5 mya forward shift in time relative to our nuclear phylogeny, with median crown ages of the family (20.2 vs 24.5 mya, respectively) and basal split of Aethionemeae (16.9 vs 21.1 mya, respectively; fig. 3 and table 3). Importantly, results from our plastome study are also much younger (nearly 10 mya) than found by Walden et al. (2020), and differences may derive from a different set of fossils used for calibration. While our phylogenies were calibrated using the *Dressiantha bicarpellata* fossil (Gandolfo et al. 1998) that can be used for setting the age range of the Brassicales, Walden et al. (2020) applied four fossils from outside the Brassicales for setting calibration points in their divergence time analysis that included more outgroup taxa across the angiosperms. This, along with the use of different algorithms for age estimation, may have led to the difference in age estimates.

### Taxonomic considerations

Based on the critical evaluation of morphology in light of molecular phylogenetic studies, the number of Brassicaceae tribes has been on the increase from initially 25 tribes (Al-Shehbaz et al. 2006) to about 50 in the following decade based on only a handful of molecular markers. With the tremendous advances in whole-genome sequencing, both nuclear and plastome, gene-level phylogenies such as presented in this study are providing solid bases to define tribal relationships and boundaries. As a result, the number of tribes is approaching 60, and for a family of ~4,000 species, this exceeds the number of tribes in the largest angiosperm families, such as the Asteraceae, Orchidaceae, Poaceae, and Fabaceae (Koch et al. 2018).

At the same time, phylogenetic data has been accumulating, enabling a proposition for a new phylogenetically-based infrafamilial classification of the Brassicaceae. We propose an updated classification that groups numerous tribes into few higher-level taxa according to the general topology of the family phylogeny, namely two subfamilies and five supertribes in one of them. Presence of rogue taxa and remaining discordance between nuclear and plastome phylogenies might change the composition of some of the suggested supertribes, but we envision that the general taxonomic framework will likely not undergo radical changes in the future.

Specifically, two major changes are suggested. First, the establishment of two subfamilies to acknowledge the deepest split within the family (fig. 1, 3): Aethionemoideae and Brassicoideae (commonly referred to as ‘core Brassicaceae’). Aethionemoideae contains a single tribe (Aethionemeae) with a single genus, *Aethionema*. Instead, Brassicoideae contains the bulk of the family, i.e., the remaining 57 tribes (including 99% of the species). We propose to abandon the uninformative names for the main lineages that fall within this subfamily (commonly referred to as ‘lineages I–V’), and we suggest replacement by five supertribes: Camelinodae (lineage I), Brassicodae (lineage II), Hesperodae (lineage III), Arabodae (lineage IV), and Heliophilodae (lineage V). Second, in addition to the 53 Brassicaceae tribes previously accepted, we propose five new ones (see also Nikolov et al., 2019, supplementary table S10). These are Arabidopsideae (*Arabidopsis*, Camelinodae), Asperuginoideae (*Asperuginoides*, Arabodae), Hemilophieae (*Dipoma* and *Hemilophia*, Camelinodae), Schrenkielleae (*Schrenkiella*, Brassicodae), and Subularieae (*Idahoa* and *Subularia*, Heliophilodae). The isolated position of the genus *Arabidopsis* from the other genera that have traditionally been assigned to tribe Camelineae sensu Al-Shehbaz et al. (2006) has already been pointed out by various phylogenetic studies (Bailey et al. 2006; Clauss and Koch 2006; Koch et al. 2007; Lysak et al. 2009; Hohmann et al. 2015; Huang et al. 2016), raising *Arabidopsis* to the tribal level. The phylogenetic placement of *Asperuginoides* has been enigmatic for a long time (Al-Shehbaz 2012; Španiel et al. 2015; Walden et al. 2020), until Nikolov et al. (2019) and the present study showed a sister relationship to tribe Alysseae. *Dipoma* and *Hemilophia* were previously listed as unplaced (Al-Shehbaz 2012), until Nikolov et al. (2019) showed the two genera form a monophyletic clade unrelated to any tribe and suggested their placement in a new tribe, which is confirmed by our nuclear dataset and morphological insights (however, rejected by Walden et al., 2020, and our plastome phylogeny showing *Dipoma* to be affiliated with tribe Crucihimalayeae). The first robust, isolating position of *Schrenkiella*, sister to a clade including the tribes Fourraeeae-Brassiceae-Isatideae-Sisymbrieae-Thelypodieae, was shown by Walden et al. (2020) and fully supported by the present study, meriting tribal status. Nikolov et al. (2019) retrieved a sister relationship between the unplaced *Idahoa* and *Subularia*, later corroborated by Dogan et al. (2022), who proposed the placement of both genera into the resurrected tribe Subularieae DC., which is reconfirmed in the present study. The contentious phylogenetic placement of *Idahoa* and *Subularia* is best explained by two WGDs involving one or more shared parental genomes (Dogan et al. 2022).

Despite the number of recognized tribes, nearly all are monophyletic in both nuclear and plastome phylogenies with maximum or very high molecular support. Notable exceptions are tribes Camelineae (polyphyletic in both nuclear and plastome phylogenies), Iberideae (polyphyletic in the nuclear phylogeny only), and Subularieae (polyphyletic in the nuclear phylogeny). The South American CES clade (composed of tribes Cremolobeae, Eudemeae, and Schizopetaleae; Salariato et al. 2016), whose intertribal relations are well resolved in our nuclear phylogeny, were highly mixed in our plastome phylogeny, suggesting a yet unresolved mesopolyploid history of these tribes (Dogan et al. 2022).

### Toward a complete Brassicaceae species phylogeny

The ultimate goal of our Brassicaceae consortium is to build a complete Brassicaceae family phylogeny including all ~4,000 species. The methods that we applied here show that this is possible, and these can easily be scaled up. We solely applied herbarium material to extract genomic DNA from, something that has become only recently possible due to the advent of sophisticated genomic laboratory methods, such as target capture sequencing (Lemmon and Lemmon 2013; Brewer et al. 2019; Dodsworth et al. 2019). Many of the samples we used had a pre-1950 or even pre-1900 origin, which is considered very old in the light of genomic studies, but resulted in a high coverage of the targeted genes nonetheless (supplementary fig. S2). To our surprise, we found little or no effect of sample age, at least in terms of the number of genes we could retrieve for each sample. Interestingly, this means that using relatively old collection material can save on laboratory cost, as DNA extractions of older samples usually result in more fragmented DNA, therefore allowing to skip DNA fragmentation using sonication – one of the moneywise most expensive steps in the whole library preparation process. Instead, the way plants were dried upon collection is likely more important than the sample age, as shown empirically by Forrest et al. (2019). Finally, the influence of DNA damage on sequence read quality, expected to increase with sample age, needs further investigation.

We applied the ‘mixed baits’ approach of Hendriks et al. (2021), combining two bait sets in a single target capture reaction, with one set designed specifically for the Brassicaceae family (B764; Nikolov et al. 2019) and the other for general applications in angiosperms (A353; Johnson et al. 2019). Whereas both sets performed very well in terms of number of genes and total target length recovered, we found that the lineage-specific B764 bait set performed slightly better than the angiosperm-wide A353 in terms of recovery of genes proportion and target length (supplementary fig. S1). A similar result was found for subtribe Malinae (Rosaceae; Ufimov et al. 2021), but those results were compared after two separate target captures using their lineage-specific and the A353 bait sets. Contrary to Larridon et al. (2020), Ogutcen et al. (2021), and Ufimov et al. (2021), we found genes from the lineage-specific B764 bait set to be phylogenetically more informative than those recovered from the angiosperm-wide A353 bait set (supplementary fig. S12).

In conclusion, we provide the first global, complete calibrated nuclear and plastome BrassiToLs, both in terms of genus-level sampling and genome-wide data, which offers an important step forward in untangling the notoriously difficult phylogenetic relationships across Brassicaceae. Using HybPhaser, we applied the latest bioinformatic insights into the selection of most reliable genes to construct the species tree by removing all genes flagged as potential paralogs, thereby increasing topological accuracy and highlighting likely polyploid taxa. Our improved phylogenetic framework supports the reinstatement of two subfamilies (Aethionemoideae, Brassicoideae) with five supertribes in Brassicoideae (Arabodae, Brassicodae, Camelinodae, Heliophilodae and Hesperodae) that represent the former major lineages I–V, and five new (or re-established) tribes, including the monotypic Arabidopsideae, accounting for a total of 58 tribes in Brassicaceae. Because we relied solely on the rich collections of the many worldwide herbaria, our methods allow us to easily scale up sampling and analyses in the near future, which should ultimately lead to a complete ~4,000 species BrassiToL, an indispensable tool for the thousands of researchers working on this important model plant family.

## Materials and Methods

### Taxon sampling

To reconstruct the nuclear Brassicaceae phylogeny, we aimed at including at least one species from each of the 349 currently accepted genera –preferably each genus’ type species– along with one species of every non-Brassicaceae Brassicales families and all species needed for fossil calibration checks of vicariance and colonisation within lineages of Brassicaceae (Table 1). We followed BrassiBase (Kiefer et al. 2014) for taxonomic delimitation of species, added with more recent insights by taxonomic experts. Where possible, we sampled the type specimens of genera to support future taxonomic judgements. Some sampled vouchers were as much as 200 years old (to show the possibilities of the target capture methods with regard to natural history collections).

We generated new nuclear data for 365 samples (supplementary table S1) and added available sequences from 38 samples from Nikolov et al. (2019) (supplementary table S2). All new data were sequenced from dried herbarium specimens or silica tissue coming from 29 different herbarium collections across the world, with plants collected between 1807 and 2020 (including 35 pre-1900 and 64 pre-1950 samples). We used the original type material in 24 species (supplementary table S1).

New plastome data were generated from genome skimming (see below) for 237 samples (supplementary table S1). We used additional data for 196 samples from Walden et al. (2020), 60 plastid genomes downloaded from GenBank that have also been included in Walden et al. (2020) (supplementary table S7), and 31 samples from Nikolov et al. (2019) (supplementary table S2).

### Library preparation, target capture, and sequencing

Wet lab methods were described in Hendriks et al. (2021). Briefly, we extracted genomic DNA from 25 mg of dried leaf tissue (or less if not enough material was available; in case no leaf tissue was available from any herbarium voucher available to us, we used branches and/or flowers) using the DNeasy PowerPlant Pro Kit (Qiagen, Hilden, Germany), following the manufacturer’s protocol (but with a final elution time of 1 hr). DNA extracts with visible impurities (green or brown colour; ~25% of samples) were subsequently purified using the DNeasy PowerClean Pro Cleanup Kit (Qiagen, Hilden, Germany). Genomic DNA was stored in the DNA bank of Naturalis Biodiversity Center, Leiden, the Netherlands. Genomic libraries were generated using the NEBNext Ultra II FS kit (New England Biolabs, Ipswich, Massachusetts, USA) with sonication in an M220 Focused-ultrasonicator (Covaris, Woburn, Massachusetts, USA; only for libraries with fragment peak length >400 bp). Indexing was performed with 384 unique combinations from IDT 10 bp primers (Integrated DNA Technologies, Coralville, Iowa, USA), with protocol adjustments described by Hendriks et al. (2021). Target sequence capture was carried out on pools of 10-30 libraries each, using the ‘mixed baits approach’ described by Hendriks et al. (2021). This method targets putatively single-copy nuclear genes from two different bait sets in a single capture reaction: a Brassicaceae-specific set targeting 1,827 exons from 764 genes, using 40k probes (Nikolov et al. 2019; hereafter B764), and the now widely used Angiosperms353 v1 universal bait set, using 80k genes (Johnson et al. 2019; William J. Baker et al. 2021; hereafter A353) (both from Arbor Biosciences, Ann Arbor, Michigan, USA). To maintain the ratio of probes among the bait sets, we used an B764 : A353= 1 : 2 (v/v) mixture. To aid skimming of chloroplast gene reads during sequencing (Weitemier et al. 2014), we used genome spiking of each enriched library with its unenriched library at a ratio of 1 : 1 (M/M). Sequencing was performed on an Illumina HiSeq 2500 sequencer (Illumina, San Diego, California, USA) at BaseClear, the Netherlands, producing 150-bp paired-end reads, at a targeted 100✕ technical coverage. Raw sequence data files were uploaded to NCBI SRA under BioProjects PRJNA806513 and PRJNA678873. Target capture and sequencing for a sample were repeated if results from sequence assembly (described below) were poor (<500 genes recovered).

New ‘mixed baits’ data for 12 outgroup species were generated by the PAFTOL project at the Royal Botanic Gardens, Kew, UK, following methods described by Baker et al. (2021). Further samples were handled by the Bailey Lab, New Mexico State University, USA, and at Heidelberg University, Germany. To further increase sampling, we added previously published raw sequence data for 37 samples from Nikolov et a. (2019), available from NCBI SRA as BioProject PRJNA518905 (target capture for B764 only; supplementary table S2).

### Sequence assembly of target capture data

Raw sequence data (from our study, as well as from other sources; see taxon sampling) were quality controlled and trimmed with Trimmomatic v0.38 (Bolger et al. 2014) using the same parameters as Baker et al. (2021). Trimmed reads were mapped against two reference files (i.e., for B764 and A353 bait sets) using HybPiper v1.3.1 (Johnson et al. 2016) with BWA v0.7.16a (Li and Durbin 2009) and SPAdes v3.14.1 (Bankevich et al. 2012), and GNU Parallel (Tange 2011) to manage parallel computing of samples on the XSEDE Stampede2 HPC (Towns et al. 2014). We built a gene reference file for the B764 dataset from the 1,827 exon reference file of Nikolov et a. (2019) by concatenation of same-gene exons, resulting in a total target length of 919,712 bp. For the A353 dataset, we used the ‘mega353’ target file with the script ‘filter_megatarget.py’ to create a mustard family-specific reference (McLay et al. 2020) with a total target length of 263,894 bp. We identified a total of 36 genes with overlap among the two bait sets, which was possible because the two bait sets have been developed independently (supplementary table S8). To avoid studying the same genetic marker twice, we discarded this subset of genes from the B764 dataset (generally the shorter targets), leaving us with a final nuclear dataset of 1,081 (i.e., 764 + 353 - 36) genes.

### Sequence assembly of plastome

We used off-target reads from genome spiking (samples with new raw data) and genome skimming (raw data from previous studies listed above) to reconstruct as much of the plastome as possible. In order to integrate the newly assembled plastid data into an already existing plastid dataset containing 231 Brassicaceae species (plus 3 duplicates) with 60 gene-coding sequences for each species (Walden et al. 2020), we sequenced and assembled plastid genomes for 237 samples and 31 samples from Nikolov et al. (2019). After removal of gap columns, the final data matrix had a total target length of 29,120 bp and was 96.2% complete. Trimmed sequencing reads were mapped using BWA v0.7.17 using option ‘BWA-MEM’ (Li 2013) and the *Arabidopsis thaliana* plastid genome (GenBank accession number NC_000932) as reference. Prior to mapping, the second copy of the inverted repeat region was removed as identical regions lead to secondary alignments which are omitted by tools used in subsequent analysis. SAMtools v1.3.1 (Danecek et al. 2021) was used for enhancing mapping quality as well as sorting and indexing the bam files. Duplicates were removed using Picard tools (http://broadinstitute.github.io/picard/). Variant calling was performed using the GATK4 function ‘HaplotypeCaller’ setting ploidy to 1 and pcr-indel-model to none (Van der Auwera and O’Connor 2020). The GATK3 function ‘FastaAlternateReferenceMaker’ (McKenna et al. 2010) was used to generate sequences including the detected SNPs and indels. Regions of low coverage (<5) and low mapping quality (<30) were detected using GATK3 function ‘CallableLoci’. After having adjusted the positions of the regions to be masked using the inhouse script ‘masker.sh’ (Markus Kiefer, Heidelberg University), BEDTools function ‘maskfasta’ (Quinlan and Hall 2010) was used for masking. The annotation of genes was transferred by alignment using the inhouse script ‘cpanno.py’ (Markus Kiefer, Heidelberg University).

### Taxonomic verification

We performed ‘taxonomic verification’ on a preliminary species phylogeny. To construct this phylogeny, multiple sequence alignments were created for each gene using MAFFT v7.273 (Katoh and Standley 2013), with a quick gene tree inference using FastTree v2.1 (Price et al. 2010), both within the pipeline PASTA v1.8.6 (Mirarab et al. 2014). Unfiltered gene trees were used as input to ASTRAL-III v5.7.8 (Zhang et al. 2018). Data from samples marked as possible or likely errors (found in highly unlikely positions in the preliminary ASTRAL-III species tree) were removed, followed by either repetition of library preparations or resampling the species. This routine was repeated once more, such that all samples were verified by taxonomic experts. Subsequently, any data resulting from multiple DNA extractions and/or library preparations from the same voucher were merged after trimming and again mapped following the same routine (supplementary table S1).

### Allelic variation and paralog detection

We used HybPhaser v2.0 (Nauheimer et al. 2021), an extension to HybPiper (Johnson et al. 2016), to assess allelic variation and to detect possible paralogs in our nuclear dataset. In short, HybPhaser performs a re-mapping of raw sequence data, using each sample’s contigs (created by HybPiper) as a new reference. Whereas HybPiper by default constructs the most likely allele for a —supposedly single-copy— gene based on the relative nucleotide frequency of each heterozygous site, HybPhaser instead takes SNP variation into account using nucleotide ambiguity codes, and uses this to quantify divergence between gene variants to detect paralogy and hybridisation. Single genes with high SNP count are likely paralogs, while samples with high SNP count across all genes are likely hybrids or polyploids (Nauheimer et al. 2021). Putative paralogs, genes with high SNP count compared to other genes, can be removed from the dataset. Since there is no single threshold to define a paralog, we used four ‘routines’ (table 1). In the ‘inclusive’ routine (1,018 genes, 332 genera, 375 species), we retained as much data as possible (including all samples, and thus all genera for which we had any data) and only discarded poorly recovered genes (gene recovered for <10% of the samples and/or proportion of gene target length recovered <10% on average across all samples). We discarded putative paralogs in the ‘strict’ routine (1,013 genes, 317 genera, 356 species) by removing all ‘outlier’ genes, as defined by HybPhaser as outlier of a boxplot distribution of mean proportion of SNPs across all genes, and ‘superstrict’ routine (297 genes, 317 genera, 356 species) by removing all genes with a mean proportion of SNPs across the dataset of 0.02 or more. This acknowledged that relatively high values are indicative of genes representing paralogs that we wanted to exclude. Finally, after noticing large differences in mean SNP proportions among tribes within the mustard family, we applied a ‘superstrict by tribe’ routine (mean 1,031 genes, with gene choice varying by tribe, 317 genera, 356 species) in which we assessed and removed mean SNP proportions by tribe (supplementary fig. S3-S4). In all but the first routine, multiple species and genera were removed from the dataset, because in some samples too few genes were recovered to validly assess possible paralogy.

After removing putative paralogs, we used HybPhaser to detect possible hybrids by calculating each sample’s allele divergence (AD, percentage of SNPs across all genes) and locus heterozygosity (LH, percentage of genes with SNPs), two metrics that are useful in the detection of hybrids (Nauheimer et al. 2021). Because hybrids (whether polyploid or not) are expected to inherit multiple alleles from their different parent species, they are expected to show relatively high levels of LH and an AD that corresponds to the divergence of the parental lineages. Very high values for AD are expected in lineages with multiple polyploidisations (Nauheimer et al. 2021). With time polyploid lineages are expected to lose duplicated genes leading to a decrease in LH. Therefore, high AD combined with intermediate LH can indicate that samples are more ancient polyploids. While there is no universal definition of what values correspond to hybrids or other types of polyploids, these values can give a good indication on the history of hybridisation in samples. Here we broadly distinguish four classes: hybrid (high LH, medium AD), highly polyploid (high LH and high AD), old polyploid (medium LH, medium AD), and old and highly polyploid (medium LH, high AD).

### Nuclear phylogenomics

We applied four different phylogenomic approaches to analyse our nuclear dataset. We used default settings and parameters for all tools, unless specified.

First, we applied a network approach to visualise possible evolutionary reticulations, inferring a splits graph (based on uncorrected p-distances) with SplitsTree4 v4.17.1 (Huson and Bryant 2006). We used a nuclear supermatrix from the gene alignments from the ‘superstrict’ routine as input.

Second, we used a maximum likelihood (ML) supermatrix approach with IQ-TREE2 v2.1.3 (Minh, Schmidt, et al. 2020), using 1,000 ultrafast bootstraps (Hoang et al. 2018) and saving bootstrap replicates, a GTR+F+R model, and sample S1321 (*Synsepalum afzelii*) as outgroup. Contrary to our coalescent-based approach (see below), this analysis generated a phylogeny in which branch lengths were representative of evolutionary change (number of mutations), which was needed in subsequent phylogeny calibration. As an input we again used a nuclear supermatrix approach, but this time with sequence alignments from the ‘inclusive’ routine, thereby including *all* samples as poor samples removed in the stricter routines were often needed for fossil calibration. To reduce the influence of genes likely plagued by too many paralogs, we only included genes with a mean SNP proportion <0.02 in the supermatrix (thus cf. the ‘superstrict’ routine, but now keeping all samples). We again used IQ-TREE2 v2.1.3 with the 297 associated gene trees and alignments (inferred also with IQ-TREE v2.1.3; see next) from the ‘superstrict’ routine to calculate gene (gCF) and site concordance factors (sCF; parameter --scf 1,000) for all nodes (Minh, Hahn, et al. 2020).

Third, we applied a coalescent approach with ASTRAL-III (Zhang et al. 2018). As input, we took the consensus sequences from the four paralog detection routines in HybPhaser (see above) to infer gene trees that served as input for a coalescent analysis. For each of the routines, we started by running a ‘de-noising loop’: sequences were aligned using MAFFT v7.273 and trimmed using trimAl v1.2 (Capella-Gutiérrez et al. 2009) with parameters resoverlap 0.75, seqoverlap 0.90, and gt 0.90. Any remaining likely sequencing errors were masked using TAPER v1.0.0 (Zhang et al. 2021) with default parameters. Gene trees were inferred using IQ-TREE2 v2.1.3 (Minh, Schmidt, et al. 2020) inferring branch support using ultrafast bootstrapping (Hoang et al. 2018), with other parameters following Baker et al. (2021). We used TreeShrink v1.3.9 (Mai and Mirarab 2018) with default parameters on the complete set of gene trees to detect and remove outlier branches and update gene trees and alignments. Discordance among gene trees was scored using all gene trees associated with each routine and calculated using normalised quartet scores for the main topology, along with first and second alternatives.

Fourth, we applied another coalescent approach with ASTRAL-Pro v1.1.6 (Zhang et al. 2020). Contrary to ASTRAL-III, this version allows the input of multiple gene copies from each individual, acknowledging that multiple alleles from gene duplications (paralogs) may actually be informative in species tree inference (e.g., no a priori choice of a definitive homologous gene copy needs to be made). Gene trees were now collected from mapping done by HybPiper using the script ‘paralog_investigator.py’, which saves all possible alleles (Johnson et al. 2016). Genes were again aligned with MAFFT v7.273 and trimmed using trimAl v1.2 with parameters resoverlap 0.75, seqoverlap 0.90, and gt 0.90, with subsequent gene tree inference with IQ-TREE2 v2.1.3.

### Phylogenetic informativeness

To study any differences in support from different nuclear genes in inferring the nuclear species tree, we studied Townsend’s phylogenetic informativeness (Townsend 2007) for all genes included in the ‘strict’ routine. First, we calculated relative evolutionary rates for all sites in each gene’s multiple sequence alignment (constrained on the ML supermatrix approach species tree) using Rate4Site v3.2 (Mayrose et al. 2004). Second, we used R package PhyInformR v1.0 (Dornburg et al. 2016) to calculate and draw phylogenetic informativeness profiles for each gene, making a distinction between genes obtained from either the B764 and A353 bait sets and genes with a mean SNP proportion of <=0.02 and >0.02.

### Plastome phylogenetics

For generating a phylogeny based on information from the plastid genome, coding sequences as well as sequences encoding tRNAs and rRNAs, genes were extracted using BEDTools v2.27.1 function ‘getfasta’ (Quinlan and Hall 2010) from the reference-based plastid genomes assembled in this study using read data generated in this study, or from Nikolov et al. (2019). The gene set was reduced to the 60 loci which had been used by Walden et al. (2020) in phylogenetic reconstruction. Sequences were aligned along with the corresponding sequences from Walden et al. (2020) using MAFFT v7.45.3. Integration of the dataset from this study into the previous dataset from Walden et al. (2020) was performed to add missing genera, but also to confirm the fit of both datasets. In a last step, gap columns were deleted and alignments were concatenated using the script ‘catfasta2phyml’ (https://github.com/nylander/catfasta2phyml). Phylogenetic reconstruction was performed using IQ-TREE v1.6.12 (Nguyen et al. 2015), supplying the program with partition information from the alignment, defining the outgroup, and running 1,000 ultrafast bootstrap replicates (Hoang et al. 2018). A second phylogenetic reconstruction was performed using IQ-TREE v2.2.0 (Minh, Schmidt, et al. 2020) with the former topology as input, this time to calculate site concordance factors as a measure of support for the splits in the tree (Minh, Hahn, et al. 2020).

### Fossil calibration

For both the nuclear and the plastome dataset, we performed phylogenetic dating in treePL v1.0 (Smith and O’Meara 2012) using the Turonian *Dressiantha bicarpellata* fossil (Gandolfo et al. 1998), estimated at 89.3–93.6 mya, as a single calibration point at the stem of the order Brassicales (cf. Couvreur et al. 2010). For the nuclear dataset, we used the topology from the ML supermatrix approach as input species tree, and reran IQ-TREE using the gene alignments from the 20 most clock-like genes only to infer relative branch lengths, acknowledging that inclusion of too many genes can easily result in an artificial pushback in time of internal nodes. To do so, we first calculated the clock-likeness of all genes following (Vankan et al. 2022), who defined clock-likeness as the coefficient of variation of all root-to-tip distances in the gene tree. When running the priming analysis in treePL, the value for ‘opt’ was set to 2, and ‘optad’ set to 1. The ‘moredetail’ and ‘moredetailad’ options were in effect and ‘optcvad’ was set to 1. Cross validation analysis indicated 10 as the best smoothing value. We assessed node age estimates by repeating the treePL (using the above optimised settings) analysis for 1,000 bootstrap trees generated with IQ-TREE, this time fixing the topology of the species tree (but not branch lengths) and summarising with TreeAnnotator v2.4.7 (Bouckaert et al. 2014) to obtain 95% HPD confidence intervals.

For the plastid dataset, the tree and the 1,000 bootstrap replicates resulting from analysis by IQ-TREE v1.6.12 were used to determine divergence times in treePL. When running the priming analysis and later adjustments, the values for ‘opt’ and ‘optad’ were both set to 3. The ‘moredetail’ and ‘moredetailad’ options were in effect and ‘optcvad’ was set to 4. Cross validation analysis indicated 0.000001 as best smoothing value. We assessed node age estimates by repeating the treePL (using the above optimised settings) analysis for the 1,000 bootstrap replicates generated in IQ-TREE. Calibrated gene trees were again summarised using TreeAnnotator v2.6.7 (Bouckaert et al. 2019) to obtain 95% HPD confidence intervals.

We performed multi-evidence validation of our new results against four other fossils and five biogeographical dating events as suggested by Franzke et al. (2016) by comparing expected and recovered node ages (table 2).

### Species tree incongruence

We used a Procrustean Approach to Cophylogeny (PACo; Balbuena et al. 2013) to detect incongruences between nuclear and plastome-derived supermatrix ML species trees, following the R-based pipeline of Pérez-Escobar et al. (2016). In short, this method scores similarities between two phylogenies by comparing patristic distances (i.e., branch length sums) among all tree tips using a Procrustes rotation. Species phylogenies were first pruned to the tribe-level by selecting a random sample from each tribe as a representative. Only tribes present in both the nuclear and plastome-derived phylogenies could be retained, because of methodological reasons (all tip labels from one tree need to be associated with tip labels from the other tree), leaving 57 ingroup tribes (plus a distinction between Iberideae I and II, which did not form a single clade in the nuclear phylogeny), plus outgroup Cleomaceae. Topological incongruence was statistically quantified using the 1,000 bootstrap phylogenies from each of the two species phylogenies, saved from bootstrap calculations in IQ-TREE before (see above under sections Nuclear phylogenomics and Plastid phylogenomics).

## Acknowledgments

This work was supported by the German Research Foundation (DFG; grant numbers MU1137/17-1 to K.M. and KO2302/23-2 to M.A.K.), the Czech Science Foundation (grant numbers 21-03909S to M.A.L. and 21-06839S to T.M.), grants from the Calleva Foundation to the Plant and Fungal Trees of Life project at the Royal Botanic Gardens, Kew, and CONICYT PAI Subvención a la instalación en la Academia convocatoria 2019 (grant number N°77190055 to O.T.-N.). MNHN kindly provided V.I. access to the collections in the framework of the RECOLNAT national Research Infrastructure. This work used the Extreme Science and Engineering Discovery Environment (XSEDE, project BIO210079), which is supported by the National Science Foundation (grant number ACI-1548562). We thank Helen Barnes, Roxali Bijmoer, Manuel Benito Crespo, Ivalu Cacho, Cyrille Chatelain, Suzanne Cubey, Gabi Droege, Catherine Gallagher, Mary Korver, Alicia Marticorena, Pina Milne, Peter Sack, Marnel Scherrenberg, Jan Wieringa, and Hasan Yıldırım for assistance sampling herbarium material. Elza Duijm and Marina Ventayol Garcia kindly provided fruitful discussions on optimisation of laboratory techniques. David Alejandro Duchene Garzon provided valuable ideas on calculations of clock-likeness of gene trees. Raw Illumina sequence data are available from NCBI SRA BioProjects PRJNA678873 and PRJNA806513.

## Author contributions

F.L. and K.M. designed the study. F.L., K.M., K.P.H., I.A.A-S., C.D.B., C.K., A.H.H., D.A.G., M.A.K., O.T-N., B.O., V.R.I., O.M., N.M.H., M.T., M.D.W., I.R., S.S., B.N., R.V., C.B., H.R., S.B.J., M.S., A.F., A.G., S.Z., B.J.L., N.S., F.W.S., I.S., P.H., W.J.B., and F.F. contributed to the sampling process. K.P.H., A.H.H., C.K., N.W., C.D.B., and A.R.Z. performed laboratory work. K.PH., C.K., L.N., N.M.H., A.R.Z., and E.L. performed the various analyses. K.P.H., F.L., C.K., K.M., W.J.B., I.A.A-S., C.D.B., A.H.H., L.N., L.N., D.A.G., M.A.K., M.A.L., N.W. took the lead in writing the manuscript, with subsequent input from all co-authors.

## References

Al-Shehbaz IA. 2012. A Generic and Tribal Synopsis Of The Brassicaceae (Cruciferae). TAXON 61:931–954.

Al-Shehbaz IA, Beilstein MA, Kellogg EA. 2006. Systematics and Phylogeny of The Brassicaceae (Cruciferae): An Overview. Plant Syst. Evol. 259:89–120.

Alsos IG, Lavergne S, Merkel MKF, Boleda M, Lammers Y, Alberti A, Pouchon C, Denoeud F, Pitelkova I, Puşcaş M, et al. 2020. The Treasure Vault Can be Opened: Large-Scale Genome Skimming Works Well Using Herbarium and Silica Gel Dried Material. Plants 9:432.

Bailey CD, Koch MA, Mayer M, Mummenhoff K, O’Kane SL Jr, Warwick SI, Windham MD, Al-Shehbaz IA. 2006. Toward a Global Phylogeny of the Brassicaceae. Mol. Biol. Evol. 23:2142–2160.

Baker William J, Bailey P, Barber V, Barker A, Bellot S, Bishop D, Botigué LR, Brewer G, Carruthers T, Clarkson JJ, et al. 2021. A Comprehensive Phylogenomic Platform for Exploring the Angiosperm Tree of Life. Syst. Biol. 71.2:301–319. [Internet]. Available from: https://doi.org/10.1093/sysbio/syab035

Baker William J., Dodsworth S, Forest F, Graham SW, Johnson MG, McDonnell A, Pokorny L, Tate JA, Wicke S, Wickett NJ. 2021. Exploring Angiosperms353: An open, community toolkit for collaborative phylogenomic research on flowering plants. Am. J. Bot. 108:1059–1065.

Bakker FT, Antonelli A, Clarke JA, Cook JA, Edwards SV, Ericson PGP, Faurby S, Ferrand N, Gelang M, Gillespie RG, et al. 2020. The Global Museum: Natural History Collections And The Future Of Evolutionary Science And Public Education. PeerJ 8:e8225.

Balbuena JA, Míguez-Lozano R, Blasco-Costa I. 2013. PACo: A Novel Procrustes Application to Cophylogenetic Analysis. PLOS ONE 8:e61048.

Bankevich A, Nurk S, Antipov D, Gurevich AA, Dvorkin M, Kulikov AS, Lesin VM, Nikolenko SI, Pham S, Prjibelski AD, et al. 2012. SPAdes: A New Genome Assembly Algorithm and Its Applications to Single-Cell Sequencing. J. Comput. Biol. 19:455–477.

Becker HF. 1961. Oligocene Plants from the Upper Ruby River Basin, Southwestern Montana. Geological Society of America

Beilstein MA, Nagalingum NS, Clements MD, Manchester SR, Mathews S. 2010. Dated Molecular Phylogenies Indicate a Miocene Origin For *Arabidopsis thaliana*. Proc. Natl. Acad. Sci. 107:18724–18728.

Beric A, Mabry ME, Harkess AE, Brose J, Schranz ME, Conant GC, Edger PP, Meyers BC, Pires JC. 2021. Comparative Phylogenetics Of Repetitive Elements in A Diverse Order of Flowering Plants (Brassicales). G3 GenesGenomesGenetics 11:jkab140.

Bloesch Z, Nauheimer L, Elias Almeida T, Crayn D, Field AR. 2022. HybPhaser Identifies Hybrid Evolution in Australian Thelypteridaceae. Mol. Phylogenet. Evol. 173:107526.

Bolger AM, Lohse M, Usadel B. 2014. Trimmomatic: A Flexible Trimmer For Illumina Sequence Data. Bioinformatics 30:2114–2120.

Bouckaert R, Heled J, Kühnert D, Vaughan T, Wu C-H, Xie D, Suchard MA, Rambaut A, Drummond AJ. 2014. BEAST2: A Software Platform for Bayesian Evolutionary Analysis. PLoS Comput. Biol. 10:e1003537.

Bouckaert R, Vaughan TG, Barido-Sottani J, Duchêne S, Fourment M, Gavryushkina A, Heled J, Jones G, Kühnert D, Maio ND, et al. 2019. BEAST 2.5: An Advanced Software Platform for Bayesian Evolutionary Analysis. PLOS Comput. Biol. 15:e1006650.

Boussau B, Scornavacca C. 2020. Reconciling Gene trees with Species Trees. In: Scornavacca C, Delsuc F, Galtier N, editors. Phylogenetics in the Genomic Era. No commercial publisher | Authors open access book. p. 3.2:1–3.2:23. Available from: https://hal.archives-ouvertes.fr/hal-02535529

Brewer GE, Clarkson JJ, Maurin O, Zuntini AR, Barber V, Bellot S, Biggs N, Cowan RS, Davies NMJ, Dodsworth S, et al. 2019. Factors Affecting Targeted Sequencing Of 353 Nuclear Genes from Herbarium Specimens Spanning the Diversity of Angiosperms. Front. Plant Sci. 10:1102.

Capella-Gutiérrez S, Silla-Martínez JM, Gabaldón T. 2009. trimAl: A Tool for Automated Alignment Trimming in Large-Scale Phylogenetic Analyses. Bioinformatics 25:1972–1973.

Castañeda-Álvarez NP, Khoury CK, Achicanoy HA, Bernau V, Dempewolf H, Eastwood RJ, Guarino L, Harker RH, Jarvis A, Maxted N, et al. 2016. Global Conservation Priorities for Crop Wild Relatives. Nat. Plants 2:1–6.

Castillo-Lorenzo E, Finch-Savage WE, Seal CE, Pritchard HW. 2019. Adaptive Significance of Functional Germination Traits in Crop Wild Relatives of *Brassica*. Agric. For. Meteorol. 264:343–350.

Chandler M. 1962. Flora of the Pipe-Clay Series of Dorset (Lower Bagshot). The Lower Tertiary Floras of Southern England, II. Br. Mus. Nat. Hist. Lond.:1–176.

Clauss MJ, Koch MA. 2006. Poorly Known Relatives of *Arabidopsis thaliana*. Trends Plant Sci. 11:449–459.

Couvreur TLP, Franzke A, Al-Shehbaz IA, Bakker FT, Koch MA, Mummenhoff K. 2010. Molecular Phylogenetics, Temporal Diversification, and Principles of Evolution in the Mustard Family (Brassicaceae). Mol. Biol. Evol. 27:55–71.

Danecek P, Bonfield JK, Liddle J, Marshall J, Ohan V, Pollard MO, Whitwham A, Keane T, McCarthy SA, Davies RM, et al. 2021. Twelve years of SAMtools and BCFtools. GigaScience 10:giab008.

Degnan JH, Rosenberg NA. 2009. Gene Tree Discordance, Phylogenetic Inference and The Multispecies Coalescent. Trends Ecol. Evol. 24:332–340.

Dodsworth S, Pokorny L, Johnson MG, Kim JT, Maurin O, Wickett NJ, Forest F, Baker WJ. 2019. Hyb-Seq for Flowering Plant Systematics. Trends Plant Sci. 24:887–891.

Dogan M, Mandáková T, Guo X, Lysak MA. 2022. *Idahoa* and *Subularia*: Hidden Polyploid Origins of Two Enigmatic Genera of Crucifers. Am. J. Bot. 109:1273–1289.

Dogan M, Pouch M, Mandáková T, Hloušková P, Guo X, Winter P, Chumová Z, Van Niekerk A, Mummenhoff K, Al-Shehbaz IA, et al. 2021. Evolution of Tandem Repeats Is Mirroring Post-polyploid Cladogenesis in *Heliophila* (Brassicaceae). Front. Plant Sci. 11:607893.

Dornburg A, Fisk JN, Tamagnan J, Townsend JP. 2016. PhyInformR: Phylogenetic Experimental Design and Phylogenomic Data Exploration in R. BMC Evol. Biol. 16:262.

Edger PP, Heidel-Fischer HM, Bekaert M, Rota J, Glöckner G, Platts AE, Heckel DG, Der JP, Wafula EK, Tang M, et al. 2015. The Butterfly Plant Arms-Race Escalated by Gene and Genome Duplications. Proc. Natl. Acad. Sci. 112:8362–8366.

Folk RA, Kates HR, LaFrance R, Soltis DE, Soltis PS, Guralnick RP. 2021. High-Throughput Methods for Efficiently Building Massive Phylogenies from Natural History Collections. Appl. Plant Sci. 9:e11410.

Forrest LL, Hart ML, Hughes M, Wilson HP, Chung K-F, Tseng Y-H, Kidner CA. 2019. The limits of Hyb-Seq for Herbarium Specimens: Impact of Preservation Techniques. Front. Ecol. Evol. 7:439.

Forsythe ES, Nelson ADL, Beilstein MA. 2020. Biased Gene Retention in The Face of Introgression Obscures Species Relationships. Genome Biol. Evol. 12(9): 1646–1663.

Franzke A, German D, Al-Shehbaz IA, Mummenhoff K. 2009. *Arabidopsis* Family Ties: Molecular Phylogeny and Age Estimates in Brassicaceae. TAXON 58:425–437.

Franzke A, Hurka H, Janssen D, Neuffer B, Friesen N, Markov M, Mummenhoff K. 2004. Molecular Signals for Late Tertiary/Early Quaternary Range Splits of a Eurasian Steppe Plant: *Clausia aprica* (Brassicaceae). Mol. Ecol. 13:2789–2795.

Franzke A, Koch MA, Mummenhoff K. 2016. Turnip Time Travels: Age Estimates in Brassicaceae. Trends Plant Sci. 21:554–561.

Franzke A, Lysak MA, Al-Shehbaz IA, Koch MA, Mummenhoff K. 2011. Cabbage Family Affairs: The Evolutionary History of Brassicaceae. Trends Plant Sci. 16:108–116.

Gandolfo MA, Nixon KC, Crepet WL. 1998. A New Fossil Flower from the Turonian of New Jersey: Dressiantha bicarpellata gen. et sp. nov. (Capparales). Am. J. Bot. 85:964–974.

German DA, Friesen NW. 2014. Shehbazia (Shehbazieae, Cruciferae), a New Monotypic Genus and Tribe of Hybrid Origin from Tibet. Turczaninowia 17:17–23.

German DA, Grant JR, Lysak MA, Al-Shehbaz IA. 2011. Molecular Phylogeny and Systematics of the Tribe Chorisporeae (Brassicaceae). Plant Syst. Evol. 294:65–86.

Guo X, Mandáková T, Trachtová K, Özüdoğru B, Liu J, Lysak MA. 2020. Linked by Ancestral Bonds: Multiple Whole-Genome Duplications and Reticulate Evolution in a Brassicaceae Tribe. Mol. Biol. Evol. 38(5):1695–714.

Hart ML, Forrest LL, Nicholls JA, Kidner CA. 2016. Retrieval of Hundreds of Nuclear Loci From Herbarium Specimens. TAXON 65:1081–1092.

von Hayek A. 1911. Entwurf Eines Cruciferen-Systems auf Phylogenetischer Grundlage. Beih. Bot. Centralbl. 27:127–335.

Heenan PB, McGlone MS. 2013. Evolution of New Zealand Alpine and Open-Habitat Plant Species During the Late Cenozoic. N. Z. J. Ecol.:105–113.

Heenan PB, Mitchell AD. 2003. Phylogeny, Biogeography and Adaptive Radiation Of *Pachycladon* (Brassicaceae) in The Mountains of South Island, New Zealand. J. Biogeogr. 30:1737–1749.

Hendriks KP, Mandáková T, Hay NM, Ly E, Huysduynen AH van, Tamrakar R, Thomas SK, Toro-Núñez O, Pires JC, Nikolov LA, et al. 2021. The Best of Both Worlds: Combining Lineage-Specific and Universal Bait Sets in Target-Enrichment Hybridization Reactions. Appl. Plant Sci. 9:e11438.

Hoang DT, Chernomor O, von Haeseler A, Minh BQ, Vinh LS. 2018. UFBoot2: Improving the Ultrafast Bootstrap Approximation. Mol. Biol. Evol. 35:518–522.

Hohmann N, Wolf EM, Lysak MA, Koch MA. 2015. A Time-Calibrated Road Map of Brassicaceae Species Radiation and Evolutionary History. Plant Cell 27:2770–2784.

Hu Q, Ma Y, Mandáková T, Shi S, Chen C, Sun P, Zhang L, Feng L, Zheng Y, Feng X, et al. 2021. Genome Evolution of The Psammophyte *Pugionium* for Desert Adaptation and Further Speciation. Proc. Natl. Acad. Sci. 118.42: e2025711118.

Huang C-H, Sun R, Hu Y, Zeng L, Zhang N, Cai L, Zhang Q, Koch MA, Al-Shehbaz I, Edger PP, et al. 2016. Resolution of Brassicaceae Phylogeny Using Nuclear Genes Uncovers Nested Radiations and Supports Convergent Morphological Evolution. Mol. Biol. Evol. 33:394–412.

Huang X-C, German DA, Koch MA. 2020. Temporal Patterns of Diversification in Brassicaceae Demonstrate Decoupling of Rate Shifts and Mesopolyploidization Events. Ann. Bot. 125:29–47.

Huson DH, Bryant D. 2006. Application of Phylogenetic Networks in Evolutionary Studies. Mol. Biol. Evol. 23:254–267.

Jabeen N. 2020. Agricultural, Economic and Societal Importance of Brassicaceae Plants. In: Hasanuzzaman M, editor. The Plant Family Brassicaceae: Biology and Physiological Responses to Environmental Stresses. Singapore: Springer. p. 45–128.

Janchen E. 1942. Das System der Cruciferen. Österr. Bot. Z. 91:1–28.

Jeffroy O, Brinkmann H, Delsuc F, Philippe H. 2006. Phylogenomics: the Beginning of Incongruence? Trends Genet. 22:225–231.

Johnson MG, Gardner EM, Liu Y, Medina R, Goffinet B, Shaw AJ, Zerega NJC, Wickett NJ. 2016. HybPiper: Extracting Coding Sequence and Introns for Phylogenetics from High-Throughput Sequencing Reads Using Target Enrichment. Appl. Plant Sci. 4:1600016.

Johnson MG, Pokorny L, Dodsworth S, Botigué LR, Cowan RS, Devault A, Eiserhardt WL, Epitawalage N, Forest F, Kim JT, et al. 2019. A Universal Probe Set for Targeted Sequencing of 353 Nuclear Genes from Any Flowering Plant Designed Using k-Medoids Clustering. Syst. Biol. 68:594–606.

Karl R, Koch MA. 2013. A World-Wide Perspective on Crucifer Speciation and Evolution: Phylogenetics, Biogeography and Trait Evolution in Tribe Arabideae. Ann. Bot. 112:983–1001.

Kates HR, Doby JR, Siniscalchi CM, LaFrance R, Soltis DE, Soltis PS, Guralnick RP, Folk RA. 2021. The Effects of Herbarium Specimen Characteristics on Short-Read NGS Sequencing Success in Nearly 8000 Specimens: Old, Degraded Samples Have Lower DNA Yields but Consistent Sequencing Success. Front. Plant Sci. 12:1076.

Katoh K, Standley DM. 2013. MAFFT Multiple Sequence Alignment Software Version 7: Improvements in Performance and Usability. Mol. Biol. Evol. 30:772–780.

Kiefer M, Schmickl R, German DA, Mandáková T, Lysak MA, Al-Shehbaz IA, Franzke A, Mummenhoff K, Stamatakis A, Koch MA. 2014. BrassiBase: Introduction to a Novel Knowledge Database on Brassicaceae Evolution. Plant Cell Physiol. 55:e3.

Koch M, Al-Shehbaz IA, Mummenhoff K. 2003. Molecular Systematics, Evolution, and Population Biology in the Mustard Family (Brassicaceae). Ann. Mo. Bot. Gard. 90:151– 171.

Koch M, Haubold B, Mitchell-Olds T. 2001. Molecular Systematics of the Brassicaceae: Evidence from Coding Plastidic matK and Nuclear Chs Sequences. Am. J. Bot. 88:534–544.

Koch MA, Al-Shehbaz IA. 2009. Molecular Systematics and Evolution of “Wild” Crucifers (Brassicaceae or Cruciferae), SK Gupta, ed. In: Biology and Breeding of Crucifers. Taylor and Francis Group, Boca Raton, FL.

Koch MA, Dobeš C, Kiefer C, Schmickl R, Klimeš L, Lysak MA. 2007. Supernetwork Identifies Multiple Events of Plastid trnF_(GAA)_ Pseudogene Evolution in the Brassicaceae. Mol. Biol. Evol. 24:63–73.

Koch MA, German DA, Kiefer M, Franzke A. 2018. Database Taxonomics as Key to Modern Plant Biology. Trends Plant Sci. 23:4–6.

Koch MA, Mummenhoff K. 2006. Editorial: Evolution and Phylogeny of the Brassicaceae. Plant Syst. Evol. 259:81–83.

Larridon I, Villaverde T, Zuntini AR, Pokorny L, Brewer GE, Epitawalage N, Fairlie I, Hahn M, Kim J, Maguilla E, et al. 2020. Tackling Rapid Radiations with Targeted Sequencing. Front. Plant Sci. 10:1655.

Lemmon EM, Lemmon AR. 2013. High-Throughput Genomic Data in Systematics and Phylogenetics. Annu. Rev. Ecol. Evol. Syst. 44:99–121.

Li H. 2013. Aligning Sequence Reads, Clone Sequences and Assembly Contigs with BWA-MEM. Available from: http://arxiv.org/abs/1303.3997.

Li H, Durbin R. 2009. Fast and Accurate Short Read Alignment with Burrows–Wheeler Transform. Bioinformatics 25:1754–1760.

Lielke K, Manchester S, Meyer H. 2012. Reconstructing the Environment of the Northern Rocky Mountains During the Eocene/Oligocene Transition: Constraints from the Palaeobotany and Geology of South-Western Montana, Usa. Acta Palaeobot. 52:317–358.

Liu L-M, Du X-Y, Guo C, Li D-Z. 2020. Resolving Robust Phylogenetic Relationships of Core Brassicaceae Using Genome Skimming Data. J. Syst. Evol. 59(3):442–453.

Lysak MA, Cheung K, Kitschke M, Bureš P. 2007. Ancestral Chromosomal Blocks Are Triplicated in Brassiceae Species with Varying Chromosome Number and Genome Size. Plant Physiol. 145:402–410.

Lysak MA, Koch MA, Beaulieu JM, Meister A, Leitch IJ. 2009. The Dynamic Ups and Downs of Genome Size Evolution in Brassicaceae. Mol. Biol. Evol. 26:85–98.

Lysak MA, Koch MA, Pecinka A, Schubert I. 2005. Chromosome Triplication Found Across the Tribe Brassiceae. Genome Res. 15:516–525.

Mabry ME, Brose JM, Blischak PD, Sutherland B, Dismukes WT, Bottoms CA, Edger PP, Washburn JD, An H, Hall JC, et al. 2020. Phylogeny and Multiple Independent Whole-Genome Duplication Events in the Brassicales. Am. J. Bot. 107:1148–1164.

Maddison WP. 1997. Gene Trees in Species Trees. Syst. Biol. 46:523–536.

Mai U, Mirarab S. 2018. TreeShrink: Fast and Accurate Detection of Outlier Long Branches in Collections of Phylogenetic Trees. BMC Genomics 19:272.

Mandáková T, Guo X, Özüdoğru B, Mummenhoff K, Lysak MA. 2018. Hybridization-Facilitated Genome Merger and Repeated Chromosome Fusion After 8 Million Years. Plant J. 96:748–760.

Mandáková T, Li Z, Barker MS, Lysak MA. 2017. Diverse Genome Organization Following 13 Independent Mesopolyploid Events in Brassicaceae Contrasts with Convergent Patterns of Gene Retention. Plant J. 91:3–21.

Mandáková T, Lysak MA. 2018. Post-Polyploid Diploidization and Diversification Through Dysploid Changes. Curr. Opin. Plant Biol. 42:55–65.

Mandáková T, Pouch M, Harmanová K, Zhan SH, Mayrose I, Lysak MA. 2017. Multispeed Genome Diploidization and Diversification after an Ancient Allopolyploidization. Mol. Ecol. 26:6445–6462.

Mayrose I, Graur D, Ben-Tal N, Pupko T. 2004. Comparison of Site-Specific Rate-Inference Methods for Protein Sequences: Empirical Bayesian Methods Are Superior. Mol. Biol. Evol. 21:1781–1791.

McKenna A, Hanna M, Banks E, Sivachenko A, Cibulskis K, Kernytsky A, Garimella K, Altshuler D, Gabriel S, Daly M, et al. 2010. The Genome Analysis Toolkit: A MapReduce framework for Analyzing Next-Generation DNA Sequencing Data. Genome Res. 20:1297–1303.

McLay TG, Gunn BF, Ning W, Tate JA, Nauheimer L, Joyce EM, Simpson L, Schmidt-Lebuhn AN, Baker WJ, Forest F, et al. 2020. New Targets Acquired: Improving Locus Recovery from the Angiosperms353 Probe Set. bioRxiv:2020.10.04.325571.

Minh BQ, Hahn MW, Lanfear R. 2020. New Methods to Calculate Concordance Factors for Phylogenomic Datasets. Mol. Biol. Evol. 37(9): 2727–2733.

Minh BQ, Schmidt HA, Chernomor O, Schrempf D, Woodhams MD, von Haeseler A, Lanfear R. 2020. IQ-TREE 2: New Models and Efficient Methods for Phylogenetic Inference in the Genomic Era. Mol. Biol. Evol. 37:1530–1534.

Mirarab S, Nguyen N, Guo S, Wang L-S, Kim J, Warnow T. 2014. PASTA: Ultra-Large Multiple Sequence Alignment for Nucleotide and Amino-Acid Sequences. J. Comput. Biol. 22:377–386.

Mitchell-Olds T, Al-Shehbaz IA, Koch M, Sharbel TF. 2005. Crucifer Evolution in the Post-Genomic Era. In: Plant Diversity and Evolution: Genotypic and Phenotypic Variation in Higher Plants. CAB International Wallingford, Oxfordshire, United Kingdom. p. 119–137.

Nauheimer L, Weigner N, Joyce E, Crayn D, Clarke C, Nargar K. 2021. HybPhaser: A Workflow for the Detection and Phasing of Hybrids in Target Capture Data Sets. Appl. Plant Sci. 9.

Nguyen L-T, Schmidt HA, von Haeseler A, Minh BQ. 2015. IQ-TREE: A Fast and Effective Stochastic Algorithm for Estimating Maximum-Likelihood Phylogenies. Mol. Biol. Evol. 32:268–274.

Nikolov LA, Shushkov P, Nevado B, Gan X, Al-Shehbaz IA, Filatov D, Bailey CD, Tsiantis M. 2019. Resolving the Backbone of the Brassicaceae Phylogeny for Investigating Trait Diversity. New Phytol. 222:1638–1651.

Nikolov LA, Tsiantis M. 2017. Using Mustard Genomes to Explore the Genetic Basis of Evolutionary Change. Curr. Opin. Plant Biol. 36:119–128.

Ogutcen E, Christe C, Nishii K, Salamin N, Möller M, Perret M. 2021. Phylogenomics of Gesneriaceae Using Targeted Capture of Nuclear Genes. Mol. Phylogenet. Evol. 157:107068.

Özüdoğru B, Akaydın G, Erik S, Al-Shehbaz IA, Mummenhoff K. 2015. Phylogeny, Diversification and Biogeographic Implications of the Eastern Mediterranean Endemic Genus *Ricotia* (Brassicaceae). TAXON 64:727–740.

Parkin IAP, Gulden SM, Sharpe AG, Lukens L, Trick M, Osborn TC, Lydiate DJ. 2005. Segmental Structure of the *Brassica napus* Genome Based on Comparative Analysis with *Arabidopsis thaliana*. Genetics 171:765–781.

Pérez-Escobar OA, Balbuena JA, Gottschling M. 2016. Rumbling Orchids: How to Assess Divergent Evolution Between Chloroplast Endosymbionts and the Nuclear Host. Syst. Biol. 65:51–65.

Price MN, Dehal PS, Arkin AP. 2010. FastTree 2 – Approximately Maximum-Likelihood Trees for Large Alignments. PLOS ONE 5:e9490.

Quezada-Martinez D, Addo Nyarko CP, Schiessl SV, Mason AS. 2021. Using Wild Relatives and Related Species to Build Climate Resilience in *Brassica* Crops. Theor. Appl. Genet. 134:1711–1728.

Quinlan AR, Hall IM. 2010. BEDTools: A Flexible Suite of Utilities for Comparing Genomic Features. Bioinformatics 26:841–842.

Rahimi A, Karami O, Lestari AD, de Werk T, Amakorová P, Shi D, Novák O, Greb T, Offringa R. 2022. Control of Cambium Initiation and Activity in *Arabidopsis* by the Transcriptional Regulator AHL15. Curr. Biol. 32:1764–1775.e3.

Ramírez-Barahona S, Sauquet H, Magallón S. 2020. The Delayed and Geographically Heterogeneous Diversification of Flowering Plant Families. Nat. Ecol. Evol.:1–7.

Rannala B, Yang Z. 2008. Phylogenetic Inference Using Whole Genomes. Annu. Rev. Genomics Hum. Genet. 9:217–231.

Rokas A, Carroll SB. 2005. More Genes or More Taxa? The Relative Contribution of Gene Number and Taxon Number to Phylogenetic Accuracy. Mol. Biol. Evol. 22:1337–1344.

Romero EJ, Hickey LJ. 1976. A Fossil Leaf of Akaniaceae from Paleocene Beds in Argentina. Bull. Torrey Bot. Club:126–131.

Salariato DL, Zuloaga FO, Franzke A, Mummenhoff K, Al-Shehbaz IA. 2016. Diversification Patterns in the CES Clade (Brassicaceae Tribes Cremolobeae, Eudemeae, Schizopetaleae) in Andean South America. Bot. J. Linn. Soc. 181:543–566.

Särkinen T, Staats M, Richardson JE, Cowan RS, Bakker FT. 2012. How to Open the Treasure Chest? Optimising DNA Extraction from Herbarium Specimens. PLOS ONE 7:e43808.

Sayyari E, Mirarab S. 2016. Fast Coalescent-Based Computation of Local Branch Support from Quartet Frequencies. Mol. Biol. Evol. 33:1654–1668.

Schulz OE. 1919. Cruciferae-Brassiceae. Part 1. In: Engler A, editor. Pflanzenreich IV. 105 (Heft 70). Leipzig: Verlag von Wilhelm Engelmann. p. 1–290.

Selmeier A. 2005. *Capparidoxylon holleisii* nov. spec., a Silicified Capparis (Capparaceae) Wood with Insect Coprolites from the Neogene of Southern Germany. Zitteliana:199–209.

Smith SA, Brown JW, Walker JF. 2018. So Many Genes, So Little Time: A Practical Approach to Divergence-Time Estimation in the Genomic Era. PLOS ONE 13:e0197433.

Smith SA, O’Meara BC. 2012. treePL: Divergence Time Estimation Using Penalized Likelihood for Large Phylogenies. Bioinformatics 28:2689–2690.

Španiel S, Kempa M, Salmerón-Sánchez E, Fuertes-Aguilar J, Mota JF, Al-Shehbaz IA, German DA, Olšavská K, Šingliarová B, Zozomová-Lihová J, et al. 2015. AlyBase: Database of Names, Chromosome Numbers, and Ploidy Levels of Alysseae (Brassicaceae), with a New Generic Concept of the Tribe. Plant Syst. Evol. 301:2463–2491.

Sun J, Ni X, Bi S, Wu W, Ye J, Meng J, Windley BF. 2014. Synchronous Turnover of Flora, Fauna and Climate at the Eocene–Oligocene Boundary in Asia. Sci. Rep. 4:7463.

Tange O. 2011. GNU Parallel: The Command-Line Power Tool. USENIX Mag. [Internet]. Available from: https://www.usenix.org/publications/login/february-2011-volume-36-number-1/gnu-parallel-command-line-power-tool

Towns J, Cockerill T, Dahan M, Foster I, Gaither K, Grimshaw A, Hazlewood V, Lathrop S, Lifka D, Peterson GD. 2014. XSEDE: Accelerating Scientific Discovery. Comput. Sci. Eng. 16:62–74.

Townsend JP. 2007. Profiling Phylogenetic Informativeness. Syst. Biol. 56:222–231.

Ufimov R, Zeisek V, Píšová S, Baker WJ, Fér T, van Loo M, Dobeš C, Schmickl R. 2021. Relative Performance of Customized and Universal Probe Sets in Target Enrichment: A Case Study in Subtribe Malinae. Appl. Plant Sci. 9:e11442.

Van der Auwera GA, O’Connor BD. 2020. Genomics in the Cloud: Using Docker, GATK, and WDL in Terra. O’Reilly Media

Vankan M, Ho SYW, Duchêne DA. 2022. Evolutionary Rate Variation among Lineages in Gene Trees has a Negative Impact on Species-Tree Inference. Syst. Biol. 71:490–500.

Walden N, German DA, Wolf EM, Kiefer M, Rigault P, Huang X-C, Kiefer C, Schmickl R, Franzke A, Neuffer B, et al. 2020. Nested Whole-Genome Duplications Coincide with Diversification and High Morphological Disparity in Brassicaceae. Nat. Commun. 11:3795.

Warwick SI. 2011. Brassicaceae in Agriculture. In: Schmidt R, Bancroft I, editors. Genetics and Genomics of the Brassicaceae. Plant Genetics and Genomics: Crops and Models. New York, NY: Springer. p. 33–65.

Warwick SI, Mummenhoff K, Sauder CA, Koch MA, Al-Shehbaz IA. 2010. Closing the Gaps: Phylogenetic Relationships in the Brassicaceae Based on DNA Sequence Data of Nuclear Ribosomal ITS Region. Plant Syst. Evol. 285:209–232.

Weitemier K, Straub SCK, Cronn RC, Fishbein M, Schmickl R, McDonnell A, Liston A. 2014. Hyb-Seq: Combining Target Enrichment and Genome Skimming for Plant Phylogenomics. Appl. Plant Sci. 2:1400042.

Yang Q, Bi H, Yang W, Li T, Jiang J, Zhang L, Liu J, Hu Q. 2020. The Genome Sequence of Alpine *Megacarpaea delavayi* Identifies Species-Specific Whole-Genome Duplication. Front. Genet. 11:812.

Yue J-P, Sun H, Baum DA, Li J-H, Al-Shehbaz IA, Ree R. 2009. Molecular Phylogeny of *Solms-laubachia* (Brassicaceae) s.l., Based on Multiple Nuclear and Plastid DNA Sequences, and its Biogeographic Implications. J. Syst. Evol. 47:402–415.

Zanazzi A, Kohn MJ, MacFadden BJ, Terry DO. 2007. Large Temperature Drop Across the Eocene–Oligocene Transition in central North America. Nature 445:639–642.

Zhang C, Rabiee M, Sayyari E, Mirarab S. 2018. ASTRAL-III: Polynomial Time Species Tree Reconstruction from Partially Resolved Gene Trees. BMC Bioinformatics 19:153.

Zhang C, Scornavacca C, Molloy EK, Mirarab S. 2020. ASTRAL-Pro: Quartet-Based Species-Tree Inference Despite Paralogy. Mol. Biol. Evol. 37:3292–3307.

Zhang C, Zhao Y, Braun EL, Mirarab S. 2021. TAPER: Pinpointing Errors in Multiple Sequence Alignments Despite Varying Rates of Evolution. Methods Ecol. Evol. 12:2145–2158.

