## Supplementary Tables and Figures for "Global Phylogeny of the Brassicaceae Provides Important Insights into Gene Discordance"

Table S1. Sample details for samples newly sequenced in this study. Dat for samples with suffix 'sl' derive from data pooled from multiple extractions and/or library preparations from the same herbarium specimen after taxonomic verification of all data separately.

| Sample | Tribe | Species | Main lineage | Herbarium | Biomaterial provider | Accession code | Collection date | Type status |
|---|---|---|---|---|---|---|---|---|
| S0055 | Erysimeae | *Erysimum cheiranthoides* | I | L | Kasper P. Hendriks, Roxali Bijmoer, Marnel Scherrenberg, Jan J. Wieringa | L.4259397 | 07/04/2011 | |
| S0156 | Anastaticeae | *Cithareloma lehmannii* | | A | Kasper P. Hendriks, Klaus Mummenhoff | B. Shishkin & V. Gonin | 31/05/1913 | |
| S0196sl | Arabideae | *Arcyosperma primulifolium* | IV | A | Kasper P. Hendriks, Klaus Mummenhoff | Polunin, Sykes, and Williams No. 965 | 16/06/1952 | |
| S0268 | Boechereae | *Phoenicaulis cheiranthoides* | I | A | Kasper P. Hendriks, Klaus Mummenhoff | Bartholomew 4159 | 13/05/1988 | |
| S0275 | Physarieae | *Synthlipsis greggii* | I | A | Kasper P. Hendriks, Klaus Mummenhoff | Zika 8514 | 09/03/1985 | |
| S0277 | Eudemeae | *Aschersoniodoxa mandoniana* | V | A | Kasper P. Hendriks, Klaus Mummenhoff | Beck 2877 | 18/02/1980 | |
| S0278 | Euclidieae | *Catenulina hedysaroides* | III | A | Kasper P. Hendriks, Klaus Mummenhoff | see photo label | 18/04/1978 | |
| S0279 | Arabideae | *Athysanus pusillus* | IV | A | Kasper P. Hendriks, Klaus Mummenhoff | Ross 7469 | 12/04/1994 | |
| S0280 | Isatideae | *Glastaria glastifolia* | II | A | Kasper P. Hendriks, Klaus Mummenhoff | Al-Shehbaz 7685 | 27/04/1976 | |
| S0281 | Arabideae | *Aubrieta deltoidea* | IV | A | Kasper P. Hendriks, Klaus Mummenhoff | Nelson 5891 | 02/06/1905 | |
| S0282 | Boechereae | *Anelsonia eurycarpa* | I | A | Kasper P. Hendriks, Klaus Mummenhoff | Tiehm 8042 | 30/06/1983 | |
| S0283 | Aphragmeae | *Aphragmus obscurus* | II | A | Kasper P. Hendriks, Klaus Mummenhoff | Stainton, Sykes, & Williams, No. 6131 | 04/07/1954 | |
| S0284 | Cochlearieae | *Ionopsidium acaule* | | A | Kasper P. Hendriks, Klaus Mummenhoff | Baum 373 | 20/08/1999 | |
| S0285 | Microlepidieae | *Carinavalva glauca* | I | A | Kasper P. Hendriks, Klaus Mummenhoff | Ising, no number | 24/08/1955 | |
| S0286 | Thelypodieae | *Chlorocrambe hastata* | II | A | Kasper P. Hendriks, Klaus Mummenhoff | Tuhy 2478 | 10/08/1985 | |
| S0288 | Alysseae | *Clastopus vestitus* | IV | A | Kasper P. Hendriks, Klaus Mummenhoff | Koelz 17812 | 26/05/1941 | |
| S0289 | Euclidieae | *Rudolf-kamelinia korolkowii* | III | A | Kasper P. Hendriks, Klaus Mummenhoff | Bartholomew 8220 | 25/06/2001 | |
| S0292 | Thlaspideae | *Parlatoria cakiloidea* | II | A | Kasper P. Hendriks, Klaus Mummenhoff | Al-Shehbaz 76214 | 06/05/1976 | |
| S0293 | Boechereae | *Cusickiella douglasii* | I | A | Kasper P. Hendriks, Klaus Mummenhoff | Tiehm 5001 | 03/06/1979 | |
| S0294 | Physarieae | *Dimorphocarpa wislizenii* | I | A | Kasper P. Hendriks, Klaus Mummenhoff | Reed & Rollins 9383 | 26/03/1993 | |
| S0297 | Brassiceae | *Horwoodia dicksoniae* | II | A | Kasper P. Hendriks, Klaus Mummenhoff | Al-Shehbaz, Weinert, & Hilly, no number | 17/04/1974 | |
| S0298 | Cardamineae | *Iodanthus pinnatifidus* | I | A | Kasper P. Hendriks, Klaus Mummenhoff | Lowden 5048 | 29/05/1993 | |
| S0299 | Euclidieae | *Leptaleum filifolium* | III | A | Kasper P. Hendriks, Klaus Mummenhoff | Al-Shehbaz 7453 | 20/03/1974 | |
| S0300 | Physarieae | *Lyrocarpa coulteri* | I | A | Kasper P. Hendriks, Klaus Mummenhoff | Ross 7045 | 03/05/1993 | |
| S0301 | Boechereae | *Polyctenium fremontii* | I | A | Kasper P. Hendriks, Klaus Mummenhoff | Tiehm 12576 | 06/07/1998 | |
| S0303 | Alysseae | *Alyssoides utriculata* | IV | B | Kasper P. Hendriks, Klaus Mummenhoff | B 10 0184667 | 24/04/1990 | |
| S0305 | Brassiceae | *Ammosperma cinerea* | II | B | Kasper P. Hendriks, Klaus Mummenhoff | B 10 0024139 | 10/04/1999 | |
| S0306 | Microlepidieae | *Blennodia canescens* | I | B | Kasper P. Hendriks, Klaus Mummenhoff | B 10 0054312 | 24/08/1997 | |
| S0308 | Alysseae | *Bornmuellera tymphaea* | IV | B | Kasper P. Hendriks, Klaus Mummenhoff | B 10 0184642 | 12/07/1989 | |
| S0312 | Calepineae | *Calepina irregularis* | II | B | Kasper P. Hendriks, Klaus Mummenhoff | B 10 0073115 | 19/04/2001 | |
| S0315 | Brassiceae | *Didesmus aegyptius* | II | B | Kasper P. Hendriks, Klaus Mummenhoff | B 10 0184655 | 23/04/1991 | |
| S0316sl | Alyssopsideae | *Dielsiocharis kotschyi* | I | B | Kasper P. Hendriks, Klaus Mummenhoff | B 10 0184654 | 1975 (?) | |
| S0317 | Brassiceae | *Erucastrum gallicum* | II | B | Kasper P. Hendriks, Klaus Mummenhoff | B 10 0099137 | 06/09/2002 | |
| S0320 | Euclidieae | *Euclidium syriacum* | III | B | Kasper P. Hendriks, Klaus Mummenhoff | B 10 0184651 | 22/04/1988 | |
| S0322 | Alysseae | *Fibigia clypeata* | IV | B | Kasper P. Hendriks, Klaus Mummenhoff | B 10 0184661 | 24/04/1990 | |
| S0323 | Brassiceae | *Guiraoa arvensis* | II | B | Kasper P. Hendriks, Klaus Mummenhoff | B 10 0184649 | 25/06/1986 | |
| S0326-2 | Brassiceae | *Kremeriella cordylocarpus* | II | B | Kasper P. Hendriks, Klaus Mummenhoff | B 10 0184646 | 07/05/1993 | |
| S0327 | Euclidieae | *Lachnoloma lehmannii* | III | B | Kasper P. Hendriks, Klaus Mummenhoff | B 10 0184645 | 16/05/1987 | |
| S0329 | Biscutelleae | *Megadenia pygmaea* | | A | Kasper P. Hendriks, Klaus Mummenhoff | Ho, Bartholomew, & Gilbert 837 | 06/08/1993 | |
| S0331 | Brassiceae | *Morisia monanthos* | II | B | Kasper P. Hendriks, Klaus Mummenhoff | B 10 0184701 | 27/03/1985 | |
| S0335 | Isatideae | *Myagrum perfoliatum* | II | B | Kasper P. Hendriks, Klaus Mummenhoff | B 10 0184672 | 29/07/1993 | |
| S0345 | Sisymbrieae | *Sisymbrium burchellii* | II | MOBOT | Ihsan Al-Shehbaz | 2659911 | 13/10/2014 | |
| S0346 | Thelypodieae | *Romanschulzia arabiformis* | II | MOBOT | Ihsan Al-Shehbaz | McVaugh 13413A | 10/10/1952 | |
| S0355 | Alysseae | *Lutzia cretica* | IV | MOBOT | Ihsan Al-Shehbaz | Strudl & Pokorny ? | 16/04/1982 | |
| S0371 | Brassiceae | *Vella pseudocytisus* | II | B | Kasper P. Hendriks, Klaus Mummenhoff | B 10 0013694 | 17/05/1986 | |
| S0373 | Brassiceae | *Succowia balearica* | II | B | Kasper P. Hendriks, Klaus Mummenhoff | B 10 0184684 | 28/04/1979 | |
| S0376 | Cardamineae | *Sisymbrella aspera* | I | B | Kasper P. Hendriks, Klaus Mummenhoff | B 10 0184692 | 30/06/1980 | |
| S0377 | Brassiceae | *Sinapidendron frutescens* | II | B | Kasper P. Hendriks, Klaus Mummenhoff | B 10 0184697 | 13/06/1997 | |
| S0380 | Brassiceae | *Psychine stylosa* | II | B | Kasper P. Hendriks, Klaus Mummenhoff | B 10 0184630 | 06/05/1990 | |
| S0381 | Thlaspideae | *Pseudovesicaria digitata* | II | B | Kasper P. Hendriks, Klaus Mummenhoff | B 10 0122158 | 03/07/2002 | |
| S0383 | Physarieae | *Physaria didymocarpa* | I | B | Kasper P. Hendriks, Klaus Mummenhoff | B 10 0184670 | 14/05/1981 | |
| S0384 | Thlaspideae | *Peltariopsis planisiliqua* | II | B | Kasper P. Hendriks, Klaus Mummenhoff | B 10 0184632 | 20/07/1981 | |
| S0385 | Brassiceae | *Nasturtiopsis coronopifolia* | II | B | Kasper P. Hendriks, Klaus Mummenhoff | B 10 0000850 | 26/03/1989 | |
| S0400 | Boechereae | *Nevada holmgrenii* | I | MOBOT | Ihsan Al-Shehbaz | 5188435 | 15/06/1999 | |
| S0403sl | Anastaticeae | *Maresia nana* | | MOBOT | Ihsan Al-Shehbaz | 5557384 | 19/05/2000 | |
| S0405 | Halimolobeae | *Mancoa hispida* | I | MOBOT | Ihsan Al-Shehbaz | 5304549 | 30/01/2000 | |
| S0407sl | Euclidieae | *Lepidostemon pedunculosus* | III | MOBOT | Ihsan Al-Shehbaz | 5731991 | 2000 (?) | |
| S0412sl | Descurainieae | *Ianhedgea minutiflora* | I | MOBOT | Ihsan Al-Shehbaz | 5736813 | 02/08/2002 | |
| S0416 | Eutremeae | *Eutrema edwardsii* | II | MOBOT | Ihsan Al-Shehbaz | 4925243 | 30/06/1998 | |
| S0417 | Euclidieae | *Christolea crassifolia* | III | GH | Michaela Schmull | Bartholomew 8237 | 25/06/2001 | |
| S0419 | Arabideae | *Acirostrum alaschanicum* | IV | MOBOT | Ihsan Al-Shehbaz | 4505376 | 29/06/1996 | |
| S0420sl | Iberideae II | *Teesdalia nudicaulis* | | MOBOT | Ihsan Al-Shehbaz | 5688349 | 20/05/2002 | |
| S0421 | Euclidieae | *Streptoloma desertorum* | III | MOBOT | Ihsan Al-Shehbaz | 3149466 | 21/01/1972 | |
| S0424 | Euclidieae | *Spryginia winkleri* | III | MOBOT | Ihsan Al-Shehbaz | 5836372 | 17/04/2003 | |
| S0427 | Cardamineae | *Planodes virginicum* | I | MOBOT | Ihsan Al-Shehbaz | 4931575 | 18/04/1998 | |
| S0428sl | Euclidieae | *Pycnoplinthus uniflora* | III | MOBOT | Ihsan Al-Shehbaz | 4890785 | 02/08/1993 | |
| S0430 | Alysseae | *Physoptychis caspica* | IV | MOBOT | Ihsan Al-Shehbaz | 4347306 | 17/06/1977 | |
| S0435 | Microlepidieae | *Pachycladon novae-zelandiae* | I | MOBOT | Ihsan Al-Shehbaz | 5324756 | 15/12/2000 | |
| S0436 | Anastaticeae | *Parolinia ornata* | | MOBOT | Ihsan Al-Shehbaz | 5324756 | 08/02/1969 | |
| S0450 | Anastaticeae | *Anastatica hierochuntica* | | MOBOT | Ihsan Al-Shehbaz | Thomas 2030 | 10/04/1983 | |
| S0464 | Brassiceae | *Fortuynia garcinii* | II | RMNH | Frederic Lens | tbd | tbd | |
| S0478 | Euclidieae | *Leiospora exscapa* | III | OSBU | Barbara Neuffer | 10410 | tbd | |

### Table S1. Continued.

| Sample | Tribe | Species | Main lineage | Herbarium | Biomaterial provider | Accession code | Collection date | Type status |
|---|---|---|---|---|---|---|---|---|
| S0509 | Microlepidieae | *Arabidella trisecta* | I | MOBOT | Ihsan Al-Shehbaz | Brewer 328 | 08/12/2009 | |
| S0512sl | Thelypodieae | *Chaunanthus mexicanus* | II | MOBOT | Ihsan Al-Shehbaz | 100281147 | ??/09/1909 | |
| S0521 | Thelypodieae | *Stanleya pinnata* | II | MOBOT | Ihsan Al-Shehbaz | Beilstein 01-47 | 14/06/2001 | |
| S0547 | Lepidieae | *Lepidium virginicum* | I | WAG | Kasper P. Hendriks, Roxali Bijmoer, Marnel Scherrenberg, Jan J. Wieringa | WAG.8003906 | 27/05/2005 | |
| S0552sl | Coluteocarpeae | *Noccaea rotundifolia* | II | WAG | Kasper P. Hendriks, Roxali Bijmoer, Marnel Scherrenberg, Jan J. Wieringa | WAG.8003723 | 29/07/2006 | |
| S0555 | Anastaticeae | *Lobularia maritima* | | WAG | Kasper P. Hendriks, Roxali Bijmoer, Marnel Scherrenberg, Jan J. Wieringa | WAG.8003708 | 07/08/2006 | |
| S0558 | Descurainieae | *Hornungia petraea* | I | WAG | Kasper P. Hendriks, Roxali Bijmoer, Marnel Scherrenberg, Jan J. Wieringa | WAG.8003379 | 11/05/2008 | |
| S0560 | Biscutelleae | *Biscutella didyma* | | WAG | Kasper P. Hendriks, Roxali Bijmoer, Marnel Scherrenberg, Jan J. Wieringa | WAG.8002349 | 03/05/2002 | |
| S0562 | Coluteocarpeae | *Microthlaspi perfoliatum* | II | WAG | Kasper P. Hendriks, Roxali Bijmoer, Marnel Scherrenberg, Jan J. Wieringa | WAG.8002357 | 06/05/2002 | |
| S0577 | Heliophileae | *Heliophila elongata* | V | L | Kasper P. Hendriks, Roxali Bijmoer, Marnel Scherrenberg, Jan J. Wieringa | L.3774413 | 15/09/1897 | |
| S0583 | Brassiceae | *Raphanus sativus* | II | L | Kasper P. Hendriks, Roxali Bijmoer, Marnel Scherrenberg, Jan J. Wieringa | L.3774403 | 07/05/1985 | |
| S0590 | Brassiceae | *Hirschfeldia incana* | II | WAG | Kasper P. Hendriks, Roxali Bijmoer, Marnel Scherrenberg, Jan J. Wieringa | L.3767817 | 18/01/1983 | |
| S0591 | Lepidieae | *Lepidium latifolium* | I | L | Kasper P. Hendriks, Roxali Bijmoer, Marnel Scherrenberg, Jan J. Wieringa | WAG.8001002 | 09/07/2015 | |
| S0593 | Cardamineae | *Nasturtium officinale* | I | WAG | Kasper P. Hendriks, Roxali Bijmoer, Marnel Scherrenberg, Jan J. Wieringa | L.3767836 | 28/07/1955 | |
| S0612 | Oreophytoneae | *Oreophyton falcatum* | I | WAG | Kasper P. Hendriks, Roxali Bijmoer, Marnel Scherrenberg, Jan J. Wieringa | WAG.0143200 | 20/06/1966 | |
| S0613sl | Coluteocarpeae | *Noccidium hastulatum* | II | WAG | Kasper P. Hendriks, Roxali Bijmoer, Marnel Scherrenberg, Jan J. Wieringa | WAG.0143230 | unknown | |
| S0616 | Alysseae | *Meniocus linifolius* | IV | WAG | Kasper P. Hendriks, Roxali Bijmoer, Marnel Scherrenberg, Jan J. Wieringa | WAG.0143205 | 28/04/1976 | |
| S0617 | Brassiceae | *Hemicrambe fruticulosa* | II | WAG | Kasper P. Hendriks, Roxali Bijmoer, Marnel Scherrenberg, Jan J. Wieringa | WAG.0071640 | 20/04/1967 | |
| S0620 | Alysseae | *Resetnikia triquetra* | IV | WAG | Kasper P. Hendriks, Roxali Bijmoer, Marnel Scherrenberg, Jan J. Wieringa | WAG.0143243 | unknown | |
| S0624 | Euclidieae | *Cymatocarpus pilosissimus* | III | WAG | Kasper P. Hendriks, Roxali Bijmoer, Marnel Scherrenberg, Jan J. Wieringa | WAG.0143248 | 01/05/1976 | |
| S0629-2 | Isatideae | *Chartoloma platycarpum* | II | WAG | Kasper P. Hendriks, Roxali Bijmoer, Marnel Scherrenberg, Jan J. Wieringa | WAG.0143229 | 21/04/1957 | |
| S0641 | Camelineae I | *Neslia paniculata* | I | OSBU | Barbara Neuffer | 18197 | ??/??/2006 | |
| S0642 | Anastaticeae | *Notoceras bicorne* | | OSBU | Barbara Neuffer | 19678 | 07/04/2009 | |
| S0643 | Halimolobeae | *Pennellia longifolia* | I | OSBU | Barbara Neuffer | 14955 | ??/??/2002 | |
| S0645sl | Euclidieae | *Tetracme quadricornis* | III | OSBU | Barbara Neuffer | 15902 | ??/??/2009 | |
| S0647-2 | Thelypodieae | *Thysanocarpus curvipes* | II | OSBU | Barbara Neuffer | 19058 | ??/??/2005 | |
| S0654 | Anastaticeae | *Eremobium aegyptiacum* | | | Klaus Mummenhoff | KM2250 | | |
| S0655sl | Anastaticeae | *Lachnocapsa spathulata* | | B | Klaus Mummenhoff | B 10 0181495 | | |
| S0656sl | Anchonieae | *Micrantha multicaulis* | III | | Klaus Mummenhoff | KM1625 | 16/08/1995 | |
| S0657 | Anchonieae | *Sterigmostemum incanum* | III | | Klaus Mummenhoff | KM2316 | 16/06/2005 | |
| S0660sl | Brassiceae | *Ceratocnemum rapistroides* | II | | Klaus Mummenhoff | KM2193 | | |
| S0662 | Brassiceae | *Enarthrocarpus arcuatus* | II | | Klaus Mummenhoff | KM2270 | | |
| S0664 | Brassiceae | *Erucaria hispanica* | II | | Klaus Mummenhoff | KM1405 | | |
| S0667 | Brassiceae | *Raffenaldia primuloides* | II | | Klaus Mummenhoff | KM2257 | ??/??/2007 | |
| S0669 | Brassiceae | *Schouwia purpurea* | II | | Klaus Mummenhoff | KM1720 | | |
| S0670 | Camelineae I | *Capsella bursa-pastoris* | I | | Klaus Mummenhoff | KM2223 | ??/??/2007 | |
| S0671 | Cardamineae | *Armoracia rusticana* | I | | Klaus Mummenhoff | KM1535 | | |
| S0672 | Cardamineae | *Rorippa sylvestris* | I | OSBU | Barbara Neuffer | 18646 | 10/07/2008 | |
| S0673 | Chorisporeae | *Diptychocarpus strictus* | III | TUH | Klaus Mummenhoff | TUH35369 | 23/05/2004 | |
| S0677 | Coluteocarpeae | *Kotschyella stenocarpa* | II | | Klaus Mummenhoff | KM2187 | ??/??/2005 | |
| S0688 | Megacarpaeeae | *Megacarpaea megalocarpa* | | OSBU | Barbara Neuffer | 14776 | ??/??/2002 | |
| S0689 | Smelowskieae | *Smelowskia alba* | I | OSBU | Barbara Neuffer | 18001 | ??/??/2006 | |
| S0690 | Boechereae | *Borodinia macrophylla* | I | OSBU | Barbara Neuffer | 04-0123-50-00 | | |
| S0691 | Anastaticeae | *Farsetia aegyptia* | | BR | Steven Janssens | BR30302136 | | |
| S0717 | Stevenieae | *Stevenia alyssoides* | IV | OSBU | Barbara Neuffer | 04-0291-50-00 | | |
| S0718 | Camelineae I | *Catolobus pendulus* | I | OSBU | Barbara Neuffer | 13514 | | |
| S0720 | Arabideae | *Pachyneurum grandiflorum* | IV | OSBU | Barbara Neuffer | 23553 | ??/??/2014 | |
| S0721sl | Camelineae III | *Pseudoarabidopsis toxophylla* | I | OSBU | Barbara Neuffer | 24601 | ??/??/2015 | |
| S0722 | Stevenieae | *Macropodium nivale* | IV | OSBU | Barbara Neuffer | 18806 | ??/??/2008 | |
| S0724sl | Alysseae | *Acuston lunarioides* | IV | OSBU | Barbara Neuffer | 26802 | ??/??/2018 | |
| S0726 | Brassiceae | *Eremophyton chevallieri* | II | OSBU | Barbara Neuffer | 13643 | ??/??/2012 | |
| S0727 | Thelypodieae | *Dryopetalon runcinatum* | II | OSBU | Barbara Neuffer | 902 | ??/??/1992 | |
| S0728sl | Microlepidieae | *Drabastrum alpestre* | I | OSBU | Barbara Neuffer | 6124 | ??/??/1992 | |
| S0731sl | Brassiceae | *Crambe maritima* | II | OSBU | Barbara Neuffer | 26278 | ??/??/2015 | |
| S0732 | Alysseae | *Clypeola jonthlaspi* | IV | OSBU | Barbara Neuffer | 26794 | ??/??/2018 | |
| S0733sl | Cardamineae | *Cardamine pratensis* | I | OSBU | Barbara Neuffer | 11501 | ??/??/1966 | |
| S0734sl | Cardamineae | *Barbarea vulgaris* | I | OSBU | Barbara Neuffer | 27005 | ??/??/2018 | |
| S0735 | Aethionemeae | *Aethionema saxatile* | basal | OSBU | Barbara Neuffer | 5814 | ??/??/1994 | |
| S0738sl | Thlaspideae | *Elburzia fenestrata* | II | TUH | Atena Eslami | TUH12238 | | |
| S0739 | Alysseae | *Odontarrhena obovata* | IV | OSBU | Barbara Neuffer | 12823 | ??/??/2002 | |
| S0740 | Alysseae | *Alyssum montanum* | IV | OSBU | Barbara Neuffer | 27096 | ??/??/2018 | |
| S0741 | Microlepidieae | *Microlepidium pilosulum* | I | | Klaus Mummenhoff | KM1185 | 16/09/2001 | |
| S0742 | Descurainieae | *Robeschia schimperi* | I | HEID | Marcus Koch, Peter Sack | 8000390 | | |
| S0743 | Brassiceae | *Cakile maritima* | II | HEID | Marcus Koch, Peter Sack | 8009813 | | |
| S0744sl | Thlaspideae | *Graellsia saxifragifolia* | II | HEID | Marcus Koch, Peter Sack | 8003280 | | |
| S0745 | Thlaspideae | *Pseudocamelina glaucophylla* | II | HEID | Marcus Koch, Peter Sack | 8002362 | | |

Table S1. Continued.

| Sample | Tribe | Species | Main lineage | Herbarium | Biomaterial provider | Accession code | Collection date | Type status |
|---|---|---|---|---|---|---|---|---|
| S0746 | Biscutelleae | *Ricotia lunaria* | | HEID | Marcus Koch, Peter Sack | 8002379 | | |
| S0747sl | Brassiceae | *Zahora ait-atta* | II | HEID | Marcus Koch, Peter Sack | 8009872 | | |
| S0748 | Alysseae | *Degenia velebitica* | IV | HEID | Marcus Koch, Peter Sack | 8009797 | | |
| S0751 | Lepidieae | *Delpinophytum patagonicum* | I | HEID | Marcus Koch, Peter Sack | 8000248 | | |
| S0752 | Brassiceae | *Otocarpus virgatus* | II | HEID | Marcus Koch, Peter Sack | 8002342 | | |
| S0753 | Cremolobeae | *Menonvillea linearis* | V | HEID | Marcus Koch, Peter Sack | 8000343 | | |
| S0754 | Thelypodieae | *Parodiodoxa chionophila* | II | HEID | Marcus Koch, Peter Sack | 8000369 | | |
| S0756 | Crucihimalayeae | *Ladakiella klimesii* | I | HEID | Marcus Koch, Peter Sack | 8000319 | | |
| S0757 | Calepineae | *Goldbachia laevigata* | II | HEID | Marcus Koch, Peter Sack | 8002205 | | |
| S0758 | Coluteocarpeae | *Callothlaspi lilacinum* | II | HEID | Marcus Koch, Peter Sack | 8003525 | | |
| S0759 | Thelypodieae | *Sarcodraba karraikensis* | II | HEID | Marcus Koch, Peter Sack | 8000397 | | |
| S0760 | Arabideae | *Scapiarabis saxicola* | IV | HEID | Marcus Koch, Peter Sack | 8000399 | | |
| S0761sl | Shehbazieae | *Shehbazia tibetica* | III | HEID | Marcus Koch, Peter Sack | 8000404 | | |
| S0762-2 | Descurainieae | *Tropidocarpum gracile* | I | HEID | Marcus Koch, Peter Sack | 8000435 | | |
| S0763 | Boechereae | *Yosemitea repanda* | I | HEID | Marcus Koch, Peter Sack | 8000447 | | |
| S0764 | Physarieae | *Nerisyrenia linearifolia* | I | HEID | Marcus Koch, Peter Sack | 8000471 | | |
| S0765 | Arabideae | *Baimashania pulvinata* | IV | HEID | Marcus Koch, Peter Sack | 8000801 | | |
| S0766 | Thlaspideae | *Didymophysa aucheri* | II | HEID | Marcus Koch, Peter Sack | 8000804 | | |
| S0768 | Brassiceae | *Carrichtera annua* | II | HEID | Marcus Koch, Peter Sack | 8002120 | | |
| S0770 | Arabideae | *Botschantzevia karatavica* | IV | HEID | Marcus Koch, Peter Sack | 8003497 | | |
| S0771 | Coluteocarpeae | *Ihsanalshehbazia granatensis* | II | HEID | Marcus Koch, Peter Sack | 8002312 | | |
| S0772 | Brassiceae | *Foleyola billotii* | II | HEID | Marcus Koch, Peter Sack | 8002200 | | |
| S0791 | Anastaticeae | *Morettia canescens* | | OSBU | Barbara Neuffer | 13669 | 29/03/2002 | |
| S0795 | Arabideae | *Dendroarabis fruticulosa* | IV | OSBU | Barbara Neuffer | 14725 | ??/??/2003 | |
| S0809 | Alyssopsideae | *Olimarabidopsis pumila* | I | OSBU | Barbara Neuffer | 14719 | ??/??/1990 | |
| S0827 | Anchonieae | *Dvorakia alaica* | III | OSBU | Barbara Neuffer | 15699 | ??/??/2004 | |
| S0834 | Arabideae | *Abdra brachycarpa* | IV | HEID | Marcus Koch, Peter Sack | 8003196 | | |
| S0836 | Arabideae | *Tomostima reptans* | IV | HEID | Marcus Koch, Peter Sack | 8003232 | | |
| S0851 | Brassiceae | *Diplotaxis harra* | II | BR | Steven Janssens | BR976564 | | |
| S0888 | Alysseae | *Lepidotrichum uechtritzianum* | IV | L | Kasper P. Hendriks, Roxali Bijmoer, Marnel Scherrenberg, Jan J. Wieringa | L.1850016 | 13/11/1963 | |
| S0890 | Aethionemeae | *Aethionema erinaceum* | basal | WAG | Kasper P. Hendriks, Roxali Bijmoer, Marnel Scherrenberg, Jan J. Wieringa | WAG.0004278 | 20/07/1842 | |
| S0892 | Heliophileae | *Heliophila africana* | V | WAG | Kasper P. Hendriks, Roxali Bijmoer, Marnel Scherrenberg, Jan J. Wieringa | WAG.1751584 | ??/??/1959 | |
| S0893 | Cardamineae | *Selenia aurea* | I | WAG | Kasper P. Hendriks, Roxali Bijmoer, Marnel Scherrenberg, Jan J. Wieringa | WAG.1471065 | 04/08/1969 | |
| S0894 | Chorisporeae | *Parrya arctica* | III | L | Kasper P. Hendriks, Roxali Bijmoer, Marnel Scherrenberg, Jan J. Wieringa | L.1855288 | unknown | |
| S0895 | Microlepidieae | *Phlegmatospermum cochlearinum* | I | L | Kasper P. Hendriks, Roxali Bijmoer, Marnel Scherrenberg, Jan J. Wieringa | L.1855167 | 30/08/1960 | |
| S0896 | Microlepidieae | *Harmsiodoxa blennodioides* | I | L | Kasper P. Hendriks, Roxali Bijmoer, Marnel Scherrenberg, Jan J. Wieringa | L.1827529 | ??/09/1971 | |
| S0897 | Brassiceae | *Quidproquo confusum* | II | L | Kasper P. Hendriks, Roxali Bijmoer, Marnel Scherrenberg, Jan J. Wieringa | L.1855113 | 28/02/1912 | |
| S0899 | Anchonieae | *Petiniotia purpurascens* | III | L | Kasper P. Hendriks, Roxali Bijmoer, Marnel Scherrenberg, Jan J. Wieringa | L.1844261 | ??/04/1859 | |
| S0901 | Arabideae | *Drabella muralis* | IV | WAG | Kasper P. Hendriks, Roxali Bijmoer, Marnel Scherrenberg, Jan J. Wieringa | WAG.0314767 | 29/04/2008 | |
| S0902 | Arabideae | *Draba incana* | IV | WAG | Kasper P. Hendriks, Roxali Bijmoer, Marnel Scherrenberg, Jan J. Wieringa | WAG.1751055 | 07/08/1970 | |
| S0903 | Physarieae | *Dithyrea californica* | I | WAG | Kasper P. Hendriks, Roxali Bijmoer, Marnel Scherrenberg, Jan J. Wieringa | WAG.1751139 | 06/04/1952 | |
| S0904 | Cardamineae | *Aplanodes doidgeana* | I | WAG | Kasper P. Hendriks, Roxali Bijmoer, Marnel Scherrenberg, Jan J. Wieringa | WAG.1240239 | 13/12/1998 | |
| S0906 | Kernereae | *Rhizobotrya alpina* | II | L | Kasper P. Hendriks, Roxali Bijmoer, Marnel Scherrenberg, Jan J. Wieringa | L.1825875 | 17/08/1889 | |
| S0941 | Microlepidieae | *Menkea australis* | I | Z | Alessia Guggisberg | Z-000161620 | ??/??/1978 | |
| S0946 | Microlepidieae | *Scambopus curvipes* | I | MEL | Pina Milne, Catherine Gallagher, Helen Barnes | MEL 2375340 | 11/08/2008 | |
| S0947sl | Microlepidieae | *Irenepharsus phasmatodes* | I | MEL | Pina Milne, Catherine Gallagher, Helen Barnes | MEL 2375096 | 08/08/2008 | |
| S0949 | Microlepidieae | *Pachymitus cardaminoides* | I | MEL | Pina Milne, Catherine Gallagher, Helen Barnes | MEL 2346248 | 25/08/2010 | |
| S0951 | Euclidieae | *Solms-laubachia pulcherrima* | III | W | Christian Bräuchler, Heimo Rainer | W0112412 | unknown | |
| S0952 | Alysseae | *Phyllolepidum rupestre* | IV | W | Christian Bräuchler, Heimo Rainer | W0112415 | 06/07/2002 | |
| S0954 | Brassiceae | *Douepea tortuosa* | II | W | Christian Bräuchler, Heimo Rainer | W0112431 | ??/??/2000 | |
| S0956 | Coluteocarpeae | *Thlaspiceras rechingeri* | II | W | Christian Bräuchler, Heimo Rainer | W0112409 | unknown | |
| S0957 | Megacarpaeeae | *Pugionium cornutum* | | W | Christian Bräuchler, Heimo Rainer | W0112413 | 15/07/1922 | |
| S0959 | Coluteocarpeae | *Neurotropis orbiculata* | II | W | Christian Bräuchler, Heimo Rainer | W0112468 | 06/08/2004 | |
| S0960 | Coluteocarpeae | *Pseudosempervivum aucheri* | II | W | Christian Bräuchler, Heimo Rainer | W0112496 | | |
| S0963 | Euclidieae | *Braya alpina* | III | W | Christian Bräuchler, Heimo Rainer | W0112433 | 19/07/1990 | |
| S0964 | Coluteocarpeae | *Masmenia rosularis* | II | W | Christian Bräuchler, Heimo Rainer | W0112466 | ??/07/1906 | |
| S0989 | Eudemeae | *Brayopsis calycina* | V | L | Kasper P. Hendriks, Roxali Bijmoer, Marnel Scherrenberg, Jan J. Wieringa | L.3768319 | 18/04/2006 | |
| S1011sl | Brassiceae | *Sinapis alba* | II | L | Kasper P. Hendriks, Roxali Bijmoer, Marnel Scherrenberg, Jan J. Wieringa | L.1834496 | 05/12/1807 | |
| S1021 | Brassiceae | *Crambella teretifolia* | II | WAG | Kasper P. Hendriks, Roxali Bijmoer, Marnel Scherrenberg, Jan J. Wieringa | WAG.1897861 | 13/04/1993 | |
| S1038 | Eudemeae | *Eudema rupestris* | V | MOBOT | Ihsan Al-Shehbaz | C. Wloq , et al. 1290 | unknown | |
| S1039 | Thelypodieae | *Englerocharis peruviana* | II | MOBOT | Ihsan Al-Shehbaz | A. Cano , et al. 20058 | unknown | |
| S1040 | Thelypodieae | *Sibara angelorum* | II | MOBOT | Ihsan Al-Shehbaz | P. Tenorio & C. Romero 12993 | unknown | |
| S1043sl | Eudemeae | *Dactylocardamum imbricatifolium* | V | MOBOT | Ihsan Al-Shehbaz | A. Cano 21546 | unknown | |
| S1044 | Thelypodieae | *Chilocardamum patagonicum* | II | MOBOT | Ihsan Al-Shehbaz | A. Donat 172 | unknown | |
| S1045 | Eudemeae | *Alshehbazia friesii* | V | MOBOT | Ihsan Al-Shehbaz | H. Trinidad HT-2017 | unknown | |
| S1049sl | Thelypodieae | *Thelypodiopsis elegans* | II | MOBOT | Ihsan Al-Shehbaz | C. Reed & R.C. Rollins 1933 | unknown | |

### Table S1. Continued.

| Sample | Tribe | Species | Main lineage | Herbarium | Biomaterial provider | Accession code | Collection date | Type status |
|---|---|---|---|---|---|---|---|---|
| S1050 | Thelypodieae | *Phravenia viereckii* | II | MOBOT | Ihsan Al-Shehbaz | J.C. Solomon 2743 | unknown | |
| S1052sl | Physarieae | *Paysonia lescurii* | I | MOBOT | Ihsan Al-Shehbaz | R. Kral 52490 | unknown | |
| S1053sl | Thelypodieae | *Streptanthus maculatus* | II | MOBOT | Ihsan Al-Shehbaz | D.E. Boufford , et al. 25509 | unknown | |
| S1069 | Heliophileae | *Heliophila macra* | V | L | Kasper P. Hendriks, Roxali Bijmoer, Marnel Scherrenberg, Jan J. Wieringa | L.1827376 | 29/11/1896 | Isosyntype |
| S1072 | Thelypodieae | *Hesperidanthus argillaceus* | II | MOBOT | Ihsan Al-Shehbaz | M.B. Franklin 7472 | unknown | |
| S1074 | Biscutelleae | *Ricotia sinuata* | | HUB | Barış Özüdoğru | Özüdoğru 3501 | 23/05/2014 | |
| S1077 | Biscutelleae | *Ricotia aucheri* | | HUB | Barış Özüdoğru | Özüdoğru 3559 | 13/06/2014 | |
| S1083 | Arabideae | *Arabis kennedyae* | IV | HEID | Marcus Koch, Peter Sack | unknown | ??/??/2019 | |
| S1089 | Schizopetaleae | *Mathewsia foliosa* | V | CONC | Oscar Toro, Alicia Marticorena | 185156 | ??/10/2014 | |
| S1093sl | Cremolobeae | *Aimara rollinsii* | V | CONC | Oscar Toro, Alicia Marticorena | 138611 | ??/04/1997 | |
| S1095 | Schizopetaleae | *Atacama nivea* | V | CONC | Oscar Toro, Alicia Marticorena | 172935 | ??/3/2007 | |
| S1096 | Thelypodieae | *Mostacillastrum pectinifolium* | II | CONC | Oscar Toro, Alicia Marticorena | 183635 | ??/11/2011 | |
| S1097 | Thelypodieae | *Ivania cremnophila* | II | K | William Baker, Sue Zmarzty (PAFTOL) | K000485233 | 08/11/1925 | Isotype |
| S1098 | Thelypodieae | *Neuontobotrys linearifolia* | II | K | William Baker, Sue Zmarzty (PAFTOL) | E. Wedermann 1115 | ??/03/1926 | |
| S1100 | Aethionemeae | *Aethionema maraschicum* | basal | K | William Baker, Sue Zmarzty (PAFTOL) | K000484211 | 04/05/1957 | Isotype |
| S1102 | Cremolobeae | *Cremolobus suffruticosus* | V | P | Vanessa Invernon | P00730023 | unknown | Syntype |
| S1103 | Plagiolobeae | *Plagioloba meyeri* | II | P | Vanessa Invernon | P05415915 | unknown | |
| S1106sl | Thelypodieae | *Phlebolobium maclovianum* | II | P | Vanessa Invernon | P05035742 | ??/??/1842 | |
| S1107-2 | Heliophileae | *Heliophila brachycarpa* | V | P | Vanessa Invernon | P05445367 | ??/??/1847 | |
| S1110 | Alysseae | *Aurinia gionae* | IV | P | Vanessa Invernon | P05426427 | 13/07/1896 | |
| S1111 | Alysseae | *Cuprella homalocarpa* | IV | P | Vanessa Invernon | P05330155 | 28/03/1931 | |
| S1113 | Arabideae | *Draba funckii* | IV | P | Vanessa Invernon | P02272466 | ??/06/1847 | Type |
| S1114-2 | Anastaticeae | *Diceratella floccosa* | | P | Vanessa Invernon | P04626322 | ??/??/1961 | |
| S1116 | Microlepidieae | *Cuphonotus humistratus* | I | CANB | Brendan Lepschi | CANB919176 | 05/07/2020 | |
| S1117 | Thelypodieae | *Weberbauera densifolia* | II | G | Fred Stauffer, Cyrille Chatelain | G00383991 | unknown | Syntype |
| S1118 | Thelypodieae | *Ivania cremnophila* | II | G | Fred Stauffer, Cyrille Chatelain | G00002279 | 08/11/1925 | Isotype |
| S1119 | Cremolobeae | *Cremolobus suffruticosus* | V | G | Fred Stauffer, Cyrille Chatelain | G00399238 | ??/??/1827 | |
| S1120 | Alysseae | *Alyssum lassiticum* | IV | G | Fred Stauffer, Cyrille Chatelain | G00389045 | 01/07/1899 | Isotype |
| S1121 | Arabideae | *Draba funckii* | IV | G | Fred Stauffer, Cyrille Chatelain | G00441353 | ??/06/1847 | Isotype |
| S1128 | Coluteocarpeae | *Eunomia oppositifolia* | II | HUB | Barış Özüdoğru | Özüdoğru 4482 | 11/06/2016 | |
| S1129sl | Coluteocarpeae | *Vania campylophylla* | II | HUB | Barış Özüdoğru | Altınözlü s.n. | ??/??/1900 | |
| S1148 | Aethionemeae | *Aethionema acarii* | basal | EGE | Barış Özüdoğru | EGE-26772 | 17/06/1993 | Holotype |
| S1168 | Brassiceae | *Physorhynchus brahuicus* | II | E | Alan Forrest, Suzanne Cubey | E00372120 | 08/03/1977 | |
| S1176 | Alysseae | *Odontarrhena tavolarae* | IV | E | Alan Forrest, Suzanne Cubey | E00016572 | 21/05/1885 | Type |
| S1177 | Arabideae | *Parryodes axilliflora* | IV | E | Alan Forrest, Suzanne Cubey | E00170538 | 14/06/1949 | Isotype |
| S1180 | Aethionemeae | *Aethionema munzurense* | basal | E | Alan Forrest, Suzanne Cubey | E00199859 | 17/07/1957 | Holotype |
| S1181-2 | Lepidieae | *Cyphocardamum aretioides* | I | E | Alan Forrest, Suzanne Cubey | E00039815 | 08/06/1965 | Isotype |
| S1182 | Anastaticeae | *Eigia longistyla* | | E | Alan Forrest, Suzanne Cubey | E00083373 | 13/03/1986 | |
| S1184 | Aethionemeae | *Aethionema demirizii* | basal | E | Alan Forrest, Suzanne Cubey | E00199849 | 28/05/1951 | Holotype |
| S1187sl | Dontostemoneae | *Clausia aprica - western group* | III | OSBU | Barbara Neuffer | 01-0076-50-00 | 16/07/1968 | |
| S1188 | Dontostemoneae | *Clausia aprica - eastern group* | III | OSBU | Barbara Neuffer | 12362 | 19/06/2001 | |
| S1189 | Coluteocarpeae | *Friedrichkarlmeyeria umbellata* | II | LE | Dmitry A. German | Grigorian s.n. | 07/05/1960 | |
| S1201sl | Schrenkielleae trib. nov. | *Schrenkiella parvula* | II | L | Kasper P. Hendriks, Roxali Bijmoer, Marnel Scherrenberg, Jan J. Wieringa | L.1832130 | ??/??/1954 | |
| S1204 | Camelineae I | *Camelina sativa* | I | L | Kasper P. Hendriks, Roxali Bijmoer, Marnel Scherrenberg, Jan J. Wieringa | L.1819735 | 05/07/1962 | |
| S1208sl | Brassiceae | *Brassica oleracea - nominate* | II | L | Kasper P. Hendriks, Roxali Bijmoer, Marnel Scherrenberg, Jan J. Wieringa | L.1834667 | 05/05/1983 | |
| S1210 | Alysseae | *Aurinia saxatilis* | IV | L | Kasper P. Hendriks, Roxali Bijmoer, Marnel Scherrenberg, Jan J. Wieringa | L.1826985 | unknown | |
| S1211 | Arabidopsideae trib. nov. | *Arabidopsis thaliana* | I | L | Kasper P. Hendriks, Roxali Bijmoer, Marnel Scherrenberg, Jan J. Wieringa | L.1832016 | 29/06/1914 | |
| S1213 | Heliophileae | *Heliophila filicaulis* | V | L | Kasper P. Hendriks, Roxali Bijmoer, Marnel Scherrenberg, Jan J. Wieringa | L.0035385 | 28/10/1962 | Paratype |
| S1214sl | Thlaspideae | *Mummenhoffia alliacea* | II | WAG | Kasper P. Hendriks, Roxali Bijmoer, Marnel Scherrenberg, Jan J. Wieringa | WAG.0100905 | 30/09/2003 | |
| S1220 | Brassiceae | *Brassica villosa* | II | WAG | Kasper P. Hendriks, Roxali Bijmoer, Marnel Scherrenberg, Jan J. Wieringa | WAG.1749486 | 21/04/1855 | |
| S1221sl | Brassiceae | *Guenthera souliei* | II | WAG | Kasper P. Hendriks, Roxali Bijmoer, Marnel Scherrenberg, Jan J. Wieringa | WAG.1749474 | 17/05/1852 | |
| S1223 | Aethionemeae | *Aethionema stenopterum* | basal | WAG | Kasper P. Hendriks, Roxali Bijmoer, Marnel Scherrenberg, Jan J. Wieringa | WAG.1410999 | 08/06/1843 | |
| S1227 | Boechereae | *Boechera holboellii* | I | L | Kasper P. Hendriks, Roxali Bijmoer, Marnel Scherrenberg, Jan J. Wieringa | L.1817752 | 11/05/1966 | |
| S1229-2 | Alysseae | *Odontarrhena cf. pterocarpa* | IV | WAG | Kasper P. Hendriks, Roxali Bijmoer, Marnel Scherrenberg, Jan J. Wieringa | WAG.0004235 | ??/??/1845 | Isosyntype |
| S1230 | Euclidieae | *Strigosella cabulica* | III | LE | Dmitry A. German | Lamond 1444 | 20/05/1956 | |
| S1232sl | Euclidieae | *Synstemon petrovii* | III | LE | Dmitry A. German | M.P. Petrov s.n. | 18/06/1958 | Type |
| S1234 | Anchonieae | *Iskandera hissarica* | III | LE | Dmitry A. German | Yunusov 6329 | 28/07/1962 | |
| S1235sl | Alyssopsideae | *Calymmatium draboides* | I | LE | Dmitry A. German | Tzvelev 874 | 28/07/1958 | |
| S1238sl | Calepineae | *Leiocarpaea cochlearioides* | II | LE | Dmitry A. German | Korotkiy & Lebedeva 653 | unknown | |
| S1239 | Euclidieae | *Sisymbriopsis schugnana* | III | LE | Dmitry A. German | Ikonnikov 9743 | 25/07/1958 | |
| S1241 | Euclidieae | *Atelanthera perpusilla* | III | LE | Dmitry A. German | Tzvelev 878 | 28/07/1958 | |
| S1245 | Chorisporeae | *Parrya maidantalica* | III | LE | Dmitry A. German | Fedtschenko s.n. | 31/07/1897 | |
| S1246 | Chorisporeae | *Parrya darvazica* | III | LE | Dmitry A. German | Korshinsky 1629 | 16/06/1897 | |
| S1249 | Alysseae | *Pterygostemon spathulatus* | IV | ALTB | Dmitry A. German | SRAE2007416 | 13/07/2007 | |
| S1250sl | Anchonieae | *Microstigma deflexum* | III | ALTB | Dmitry A. German | German s.n. | 03/07/2003 | |
| S1251 | Euclidieae | *Cryptospora falcata* | III | ALTB | Dmitry A. German | Smirnov, German & Antonyuk s.n. | 10/05/2001 | |
| S1258-2 | Thelypodieae | *Warea amplexifolia* | II | Z | Alessia Guggisberg | Z-000170252 | ??/??/1897 | |
| S1260 | Alysseae | *Odontarrhena nebrodensis* | IV | Z | Alessia Guggisberg | Z-000107630 | ??/??/1889 | |

### Table S1. Continued.

| Sample | Tribe | Species | Main lineage | Herbarium | Biomaterial provider | Accession code | Collection date | Type status |
|---|---|---|---|---|---|---|---|---|
| S1261 | Aethionemeae | *Aethionema orbiculatum* | basal | Z | Alessia Guggisberg | Z-000034855 | ??/??/1891 | |
| S1262 | Alysseae | *Alyssum taygeteum* | IV | Z | Alessia Guggisberg | Z-000004365 | ??/??/1897 | |
| S1266sl | Alysseae | *Irania pendula* | IV | K | William Baker, Sue Zmarzty (PAFTOL) | K000484541 | unknown | Type |
| S1269 | Alysseae | *Irania multicaulis* | IV | K | William Baker, Sue Zmarzty (PAFTOL) | K000484535 | 15/07/1843 | |
| S1270 | Brassiceae | *Vella castrilensis* | II | ABH | Manuel Benito Crespo Villalba, Vanessa Invernon | ABH43664 | 12/07/2000 | Isotype |
| S1276 | Cardamineae | *Leavenworthia aurea* | I | NMC | Donovan Bailey | 83019 | 17/03/2008 | |
| S1278 | Halimolobeae | *Sphaerocardamum nesliiforme* | I | BH | Donovan Bailey | Bailey & Ochoterena 156 | 11/09/1998 | |
| S1279 | Halimolobeae | *Exhalimolobos palmeri* | I | BH | Donovan Bailey | Bailey & Ochoterena 159 | 11/09/1998 | |
| S1314 | Brassiceae | *Moricandia arvensis* | II | WAG | Kasper P. Hendriks, Roxali Bijmoer, Marnel Scherrenberg, Jan J. Wieringa | WAG.1470250 | 14/07/2008 | |
| S1315 | Alysseae | *Berteroa incana* | IV | WAG | Kasper P. Hendriks, Roxali Bijmoer, Marnel Scherrenberg, Jan J. Wieringa | WAG.1241192 | 23/09/1989 | |
| S1318-2 | Camelineae III | *Chrysochamela velutina* | I | L | Kasper P. Hendriks, Roxali Bijmoer, Marnel Scherrenberg, Jan J. Wieringa | L.1826190 | 22/03/1888 | |
| S1319 | Thelypodieae | *Polypsecadium harmsianum* | II | U | Kasper P. Hendriks, Roxali Bijmoer, Marnel Scherrenberg, Jan J. Wieringa | U.1166651 | 22/03/1904 | |
| S1328sl | Hesperideae | *Hesperis matronalis* | III | L | Kasper P. Hendriks, Roxali Bijmoer, Marnel Scherrenberg, Jan J. Wieringa | L.1827168 | 01/07/1964 | |
| S1331 | Brassiceae | *Pseuderucaria clavata* | II | L | Kasper P. Hendriks, Roxali Bijmoer, Marnel Scherrenberg, Jan J. Wieringa | L.1829429 | 11/03/1938 | |
| S1332 | Sisymbrieae | *Sisymbrium altissimum* | II | L | Kasper P. Hendriks, Roxali Bijmoer, Marnel Scherrenberg, Jan J. Wieringa | L.1842627 | 20/06/1953 | |
| S1336 | Eutremeae | *Eutrema edwardsii* | II | L | Kasper P. Hendriks, Roxali Bijmoer, Marnel Scherrenberg, Jan J. Wieringa | L.1827708 | ??/??/1839 | |
| S1339 | Brassiceae | *Savignya parviflora* | II | L | Kasper P. Hendriks, Roxali Bijmoer, Marnel Scherrenberg, Jan J. Wieringa | L.1842080 | 10/02/1956 | |
| S1341 | Iberideae II | *Teesdalia coronopifolia* | | L | Kasper P. Hendriks, Roxali Bijmoer, Marnel Scherrenberg, Jan J. Wieringa | L.1844030 | 12/04/1971 | |
| S1342 | Iberideae II | *Teesdalia nudicaulis* | | L | Kasper P. Hendriks, Roxali Bijmoer, Marnel Scherrenberg, Jan J. Wieringa | L.1844059 | 17/05/1972 | |
| S1354 | Iberideae I | *Iberis fontqueri* | | W | Christian Bräuchler, Heimo Rainer | W0112419 | ??/??/2016 | |
| S1365 | Euclidieae | *Rhammatophyllum pachyrhizum* | III | LE | Dmitry A. German | Karamysheva 4057 | 27/05/1966 | |
| S1366 | Thlaspideae | *Sobolewskia sibirica* | II | LE | Dmitry A. German | Van'kov s.n. | 12/05/1913 | |
| S1368 | Euclidieae | *Anzhenxia yechengnica* | III | MOBOT | Ihsan Al-Shehbaz | Bartholomew 9569 | 04/08/2004 | |
| S1369 | Boechereae | *Boechera yorkii* | I | MOBOT | Donovan Bailey | York & Davis 2611 | 16/05/2001 | Isotype |
| S1370 | Halimolobeae | *Halimolobos lasiolobus* | I | GH | Donovan Bailey | Rollins & Tryon 58251 | | |
| S1371 | Hemilophieae trib. nov. | *Dipoma iberideum* | I | U | Kasper P. Hendriks, Roxali Bijmoer, Marnel Scherrenberg, Jan J. Wieringa | U.1175109 | ??/??/1930 | |
| S1374 | Asperuginoideae trib. nov. | *Asperuginoides axillaris* | IV | L | Kasper P. Hendriks, Roxali Bijmoer, Marnel Scherrenberg, Jan J. Wieringa | L.1819349 | 28/04/1902 | |
| S1375 | Malcolmieae | *Malcolmia maritima* | I | L | Kasper P. Hendriks, Roxali Bijmoer, Marnel Scherrenberg, Jan J. Wieringa | L.1828371 | 18/07/1928 | |
| S1376 | Thlaspideae | *Thlaspi arvense* | II | L | Kasper P. Hendriks, Roxali Bijmoer, Marnel Scherrenberg, Jan J. Wieringa | L.1844745 | 25/07/1973 | |
| S1378 | Chamireae | *Chamira circaeoides* | V | WAG | Kasper P. Hendriks, Roxali Bijmoer, Marnel Scherrenberg, Jan J. Wieringa | WAG.1739894 | 27/08/1896 | |
| S1381 | Arabidopsideae trib. nov. | *Arabidopsis halleri* | I | WAG | Kasper P. Hendriks, Roxali Bijmoer, Marnel Scherrenberg, Jan J. Wieringa | WAG.1739932 | ??/06/1853 | |
| S1400 | Microlepidieae | *Pachycladon fastigiata* | I | | Ines Schönberger, Mary Korver | CHR 518460 | 21/01/1999 | |
| S1401 | Brassiceae | *Coincya monensis* | II | WAG | Kasper P. Hendriks, Roxali Bijmoer, Marnel Scherrenberg, Jan J. Wieringa | WAG.1739679 | 26/06/2006 | |
| S1405 | Buniadeae | *Bunias erucago* | III | WAG | Kasper P. Hendriks, Roxali Bijmoer, Marnel Scherrenberg, Jan J. Wieringa | WAG.1749423 | 01/05/1847 | |
| S1407 | Descurainieae | *Descurainia sophia* | I | WAG | Kasper P. Hendriks, Roxali Bijmoer, Marnel Scherrenberg, Jan J. Wieringa | WAG.1750597 | 13/06/2006 | |
| S1408 | Brassiceae | *Eruca vesicaria* | II | WAG | Kasper P. Hendriks, Roxali Bijmoer, Marnel Scherrenberg, Jan J. Wieringa | WAG.1752430 | 30/03/1970 | |
| S1409 | Alysseae | *Irania umbellata* | IV | WAG | Kasper P. Hendriks, Roxali Bijmoer, Marnel Scherrenberg, Jan J. Wieringa | WAG.1751666 | 08/07/1842 | Isotype |
| S1412 | Brassiceae | *Orychophragmus violaceus* | II | WAG | Kasper P. Hendriks, Roxali Bijmoer, Marnel Scherrenberg, Jan J. Wieringa | WAG.1470449 | 18/03/1986 | |
| S1417 | Thelypodieae | *Pringlea antiscorbutica* | II | U | Kasper P. Hendriks, Roxali Bijmoer, Marnel Scherrenberg, Jan J. Wieringa | U.1176017 | 02/11/1874 | |
| S1419 | Thlaspideae | *Alliaria petiolata* | II | L | Kasper P. Hendriks, Roxali Bijmoer, Marnel Scherrenberg, Jan J. Wieringa | L.4262125 | 08/02/1965 | |
| S1421 | Brassiceae | *Rapistrum rugosum* | II | L | Kasper P. Hendriks, Roxali Bijmoer, Marnel Scherrenberg, Jan J. Wieringa | L.1829750 | 31/07/1962 | |
| S1422 | Thlaspideae | *Pachyphragma macrophyllum* | II | L | Kasper P. Hendriks, Roxali Bijmoer, Marnel Scherrenberg, Jan J. Wieringa | L.1855278 | 05/06/1959 | |
| S1424 | Microlepidieae | *Stenopetalum lineare* | I | L | Kasper P. Hendriks, Roxali Bijmoer, Marnel Scherrenberg, Jan J. Wieringa | L.1844232 | 09/09/1964 | |
| S1428 | Anastaticeae | *Marcus-kochia littorea* | | L | Kasper P. Hendriks, Roxali Bijmoer, Marnel Scherrenberg, Jan J. Wieringa | L.1828479 | 16/05/1957 | |
| S1442 | Brassiceae | *Henophyton deserti* | II | L | Kasper P. Hendriks, Roxali Bijmoer, Marnel Scherrenberg, Jan J. Wieringa | L.1855276 | 03/05/1963 | |
| S1449 | Thlaspideae | *Lysakia rostrata* | II | MOBOT | Ihsan Al-Shehbaz | American-Iranian Botanical Delegation 34056 | | |
| S1451 | Euclidieae | *Dichasianthus subtilissimus* | III | MOBOT | Ihsan Al-Shehbaz | Al-Shehbaz 200212 | | |
| S1458 | Brassiceae | *Quezeliantha tibestica* | II | B | Robert Vogt, Gabi Droege | B 10 1154379 | 14/02/1965 | Holotype |
| S1459 | Thelypodieae | *Zuloagocardamum jujuyensis* | II | MOBOT | Ihsan Al-Shehbaz | MO-6609516 | 16/01/2012 | Isotype |
| S1460 | Alyssopsideae | *Alyssopsis mollis* | I | B | Klaus Mummenhoff | B 10 0184647 | 21/06/1974 | |
| S1462 | Arabideae | *Sinoarabis setosifolia* | IV | HEID | Marcus Koch, Peter Sack | 8000565 | | |
| S1464 | Thelypodieae | *Hollermayera valdiviana* | II | CONC | Oscar Toro, Alicia Marticorena, Ivalu Cacho | 166901 | 10/01/2004 | |
| S1474 | Euclidieae | *Dilophia salsa* | III | LE | Dmitry A. German | Tambovtseva & Popova s. n. | 16/08/1964 | |
| S1475 | Euclidieae | *Metashangrilaia forrestii* | III | W | Christian Bräuchler, Heimo Rainer | W0112467 | 18/05/1914 | |
| S1481sl | Thelypodieae | *Dictyophragmus englerianus* | II | MOBOT | Ihsan Al-Shehbaz | C.R. Worth & J.L. Morrison 15762 | | |
| S1488 | Eudemeae | *Xerodraba lycopodioides* | V | MOBOT | Ihsan Al-Shehbaz | F.O. Zuloaga , et al. 14023 | 23/11/2012 | |

### Table S1. Continued.

| Sample | Tribe | Species | Main lineage | Herbarium | Biomaterial provider | Accession code | Collection date | Type status |
|---|---|---|---|---|---|---|---|---|
| S1498 | Arabideae | *Arabis montbretiana* | IV | WAG | Kasper P. Hendriks, Roxali Bijmoer, Marnel Scherrenberg, Jan J. Wieringa | WAG.1240684 | 14/03/1842 | |
| S1506 | Brassiceae | *Zilla spinosa* | II | WAG | Kasper P. Hendriks, Roxali Bijmoer, Marnel Scherrenberg, Jan J. Wieringa | WAG.1415644 | 08/05/1897 | |
| S1509 | Brassiceae | *Rhamphospermum arvense* | II | AMD | Kasper P. Hendriks, Roxali Bijmoer, Marnel Scherrenberg, Jan J. Wieringa | AMD.39011 | ??/06/1886 | |
| S1519 | Alysseae | *Hormathophylla reverchonii* | IV | AMD | Kasper P. Hendriks, Roxali Bijmoer, Marnel Scherrenberg, Jan J. Wieringa | AMD.49780 | ??/05/1907 | |
| S1539 | Chorisporeae | *Litwinowia tenuissima* | III | MOBOT | Klaus Mummenhoff, Ihsan Al-Shehbaz | 4933741 | 24/05/1994 | |
| S1541sl | Cardamineae | *Ornithocarpa fimbriata* | I | MOBOT | Klaus Mummenhoff, Ihsan Al-Shehbaz | 4084118 | 25/07/1990 | |
| S1561 | Lepidieae | *Lepidium arbuscula* | I | US | Klaus Mummenhoff | 3297605 | 01/03/1994 | |
| S1579 | Notothlaspideae | *Notothlaspi rosulatum* | V | L | Kasper P. Hendriks, Roxali Bijmoer, Marnel Scherrenberg, Jan J. Wieringa | L.1855260 | ??/02/1909 | |
| 8000070 | Arabideae | *Arabis scabra* | IV | | | | | |
| 8008999 | Arabideae | *Arabis blepharophylla* | IV | | | | | |
| 8009952 | Arabideae | *Pseudodraba hystrix* | IV | HEID | Marcus Koch, Peter Sack, Christiane Kiefer | 8009952 | | |
| S_670 | Boechereae | *Sandbergia whitedii* | I | | Donovan Bailey, Nikolas Hay | | missing | |

**Details of outgroup samples.**

| Sample | Family | Species | Main lineage | Herbarium | Biomaterial provider | Accession code | Collection date | Type status |
|---|---|---|---|---|---|---|---|---|
| S0564sl | Pentadipland-raceae | *Pentadiplandra brazzeana* | | L | Kasper P. Hendriks, Roxali Bijmoer, Marnel Scherrenberg, Jan J. Wieringa | WAG.8002007 | 18/11/2011 | |
| S1321 | Sapotaceae | *Synsepalum afzelii* | | WAG | Kasper P. Hendriks, Roxali Bijmoer, Marnel Scherrenberg, Jan J. Wieringa | WAG.1829925 | 20/11/2007 | |
| S1384 | Akaniaceae | *Bretschneidera sinensis* | | L | Kasper P. Hendriks, Roxali Bijmoer, Marnel Scherrenberg, Jan J. Wieringa | L.2295728 | 03/10/1989 | |
| S1386 | Capparaceae | *Cappariocordis tweedieana* | | L | Kasper P. Hendriks, Roxali Bijmoer, Marnel Scherrenberg, Jan J. Wieringa | L.1851636 | 22/04/1960 | |
| S1387 | Capparaceae | *Cappariocordis crotonoides* | | L | Kasper P. Hendriks, Roxali Bijmoer, Marnel Scherrenberg, Jan J. Wieringa | L.1846033 | 06/08/1980 | |
| S1388 | Capparaceae | *Capparis spinosa* | | L | Kasper P. Hendriks, Roxali Bijmoer, Marnel Scherrenberg, Jan J. Wieringa | L.1855831 | 20/06/1973 | |
| S1389 | Capparaceae | *Capparis tomentosa* | | L | Kasper P. Hendriks, Roxali Bijmoer, Marnel Scherrenberg, Jan J. Wieringa | L.3976741 | 26/09/2013 | |
| S1398 | Emblingiaceae | *Emblingia calceoliflora* | | L | Kasper P. Hendriks, Roxali Bijmoer, Marnel Scherrenberg, Jan J. Wieringa | L.1851247 | 13/09/1973 | |
| PAFTOL_005410 | Koeberliniaceae | *Koeberlinia spinosa* | | | PAFTOL | | | |
| PAFTOL_005411 | Bataceae | *Batis maritima* | | | PAFTOL | | | |
| PAFTOL_008053 | Tiganophytaceae | *Tiganophyton karense* | | | PAFTOL | | | |
| PAFTOL_014331 | Malvaceae | *Macrohasseltia macroterantha* | | K | PAFTOL | K000591631 | 22/04/1905 | |
| PAFTOL_019361 | Cleomaceae | *Cleome albescens* | | K | PAFTOL | K000370566 | 04/06/1905 | |
| PAFTOL_019459 | Caricaceae | *Vasconcellea quercifolia* | | RB | PAFTOL | RB01055668 | 07/07/1905 | |
| PAFTOL_023477 | Moringaceae | *Moringa oleifera* | | RB | PAFTOL | RB00591000 | | |
| PAFTOL_023503 | Salvadoraceae | *Salvadora persica* | | RB | PAFTOL | RB00370741 | | |
| PAFTOL_023511 | Setchellanthaceae | *Setchellanthus caeruleus* | | K | PAFTOL | K000630390 | 08/06/1905 | |
| PAFTOL_023513 | Tovariaceae | *Tovaria pendula* | | K | PAFTOL | K001275782 | 26/06/1905 | |
| PAFTOL_023515 | Resedaceae | *Forchhammeria pallida* | | K | PAFTOL | K001275784 | 17/06/1905 | |
| PAFTOL_023519 | Gyrostemonaceae | *Tersonia cyathiflora* | | | PAFTOL | | | |

Table S2. Sample data taken from Nikolov et al. (2019) for this study's nuclear and/or plastome phylogeny. Raw data were downloaded and reanalysed (including new mapping) using this study's bioinformatic pipeline.

| **Sample (= SRA accession)** | **Tribe** | **Species** | **Main lineage** |
|---|---|---|---|
| SRR11470309 | Sisymbrieae | *Sisymbrium irio* | II |
| SRR11470320 | Arabideae | *Arabis alpina* | IV |
| SRR8528336 | Thlaspideae | *Peltaria alliacea* | II |
| SRR8528338 | Biscutelleae | *Heldreichia bupleurifolia* | |
| SRR8528339 | Dontostemoneae | *Dontostemon integrifolius* | III |
| SRR8528341 | Fourraeeae | *Fourraea alpina* | II |
| SRR8528342 | Microlepidieae | *Geococcus pusillus* | I |
| SRR8528343 | Crucihimalayeae | *Crucihimalaya himalaica* | I |
| SRR8528347 | Euclidieae | *Neotorularia torulosa* | III |
| SRR8528350 | Oreophytoneae | *Murbeckiella pinnatifida* | I |
| SRR8528354 | Anchonieae | *Matthiola incana* | III |
| SRR8528356 | Subularieae trib. nov. | *Subularia aquatica* | V |
| SRR8528357 | Turritideae | *Turritis glabra* | I |
| SRR8528358 | Anchonieae | *Veselskya griffithiana* | III |
| SRR8528359 | Yinshanieae | *Yinshania acutangula* subsp. *acutangula* | I |
| SRR8528361 | Sisymbrieae | *Sisymbrium aegyptiacum* | II |
| SRR8528364 | Cremolobeae | *Cremolobus peruvianus* | V |
| SRR8528365 | Isatideae | *Iljinskaea planisiliqua* | II |
| SRR8528370 | Chorisporeae | *Chorispora tenella* | III |
| SRR8528372 | Conringieae | *Conringia orientalis* | II |
| SRR8528373 | Cochlearieae | *Cochlearia officinalis* | |
| SRR8528376 | Hemilophieae trib. nov. | *Hemilophia rockii* | I |
| SRR8528379 | Kernereae | *Kernera saxatilis* | II |
| SRR8528380 | Isatideae | *Isatis tinctoria* | II |
| SRR8528381 | Subularieae trib. nov. | *Idahoa scapigera* | V |
| SRR8528382 | Biscutelleae | *Lunaria rediviva* | |
| SRR8528384 | Arabideae | *Pseudoturritis turrita* | IV |
| SRR8528385 | Asteae | *Scoliaxon mexicanus* | V |
| SRR8528386 | Brassiceae II | *Bivonaea lutea* | II |
| SRR8528388 | Thelypodieae | *Raphanorhyncha crassa* | II |
| SRR8528389 | Fourraeeae | *Hurkaea josiae* | II |
| SRR8528391 | Cardamineae | *Andrzeiowskia cardamine* | I |
| SRR8528394 | Asteae | *Asta schaffneri* var. *pringlei* | V |
| SRR8528397 | Schizopetaleae | *Schizopetalon walkeri* | V |
| SRR8528398 | Sisymbrieae | *Sisymbrium leucoclada* | II |
| SRR8528399 | Biscutelleae | *Ricotia cretica* | |
| SRR8528400 | Kernereae | *Petrocallis pyrenaica* | II |

Table S3. Overview of currently accepted 349 genera in Brassicaceae and their sampling in this study.

| Tribe | Genus name used in this study (following BrassiBase) | Currently accepted name (in case different from name used) | Overall sampling | Coalescent approach in ASTRAL | | | | | Supermatrix approach in IQ-TREE | |
|---|---|---|---|---|---|---|---|---|---|---|
| | | | | inclusive | strict | super-strict | super-strict by tribe | Pro | nuclear | plastome |
| Aethionemeae | *Aethionema* | | included | × | × | × | × | × | × | × |
| Alysseae | *Acuston* | *Pevalekia* | included | × | × | × | × | × | × | × |
| Alysseae | *Alyssoides* | | included | × | × | × | × | × | × | - |
| Alysseae | *Alyssum* | | included | × | × | × | × | × | × | × |
| Alysseae | *Aurinia* | | included | × | × | × | × | × | × | × |
| Alysseae | *Berteroa* | | included | × | × | × | × | × | × | × |
| Alysseae | *Bornmuellera* | | included | × | × | × | × | × | × | × |
| Alysseae | *Brachypus* | | missing | - | - | - | - | - | - | - |
| Alysseae | *Clastopus* | | included | × | × | × | × | × | × | - |
| Alysseae | *Clypeola* | | included | × | × | × | × | × | × | × |
| Alysseae | *Cuprella* | | included | × | × | × | × | × | × | - |
| Alysseae | *Degenia* | | included | × | × | × | × | × | × | × |
| Alysseae | *Fibigia* | | included | × | × | × | × | × | × | - |
| Alysseae | *Galitzkya* | | missing | - | - | - | - | - | - | - |
| Alysseae | *Hormathophylla* | | included | × | × | × | × | × | × | × |
| Alysseae | *Irania* | | included | × | × | × | × | × | × | × |
| Alysseae | *Lepidotrichum* | | included | × | × | × | × | × | × | × |
| Alysseae | *Lutzia* | | included | × | × | × | × | × | × | - |
| Alysseae | *Meniocus* | | included | × | × | × | × | × | × | × |
| Alysseae | *Odontarrhena* | | included | × | × | × | × | × | × | × |
| Alysseae | *Phyllolepidum* | | included | × | × | × | × | × | × | × |
| Alysseae | *Physoptychis* | | included | × | × | × | × | × | × | - |
| Alysseae | *Pterygostemon* | | included | × | × | × | × | × | × | × |
| Alysseae | *Resetnikia* | | included | × | × | × | × | × | × | × |
| Alysseae | *Takhtajaniella* | | missing | - | - | - | - | - | - | - |
| Alyssopsideae | *Alyssopsis* | | included | × | × | × | × | - | × | × |
| Alyssopsideae | *Calymmatium* | | included | × | × | × | × | - | × | × |
| Alyssopsideae | *Dielsiocharis* | | included | × | - | - | - | - | × | - |
| Alyssopsideae | *Olimarabidopsis* | | included | × | × | × | × | - | × | × |
| Anastaticeae | *Anastatica* | | included | × | × | × | × | × | × | × |
| Anastaticeae | *Cithareloma* | | included | × | × | × | × | × | × | × |
| Anastaticeae | *Diceratella* | | included | × | × | × | × | × | × | × |
| Anastaticeae | *Eigia* | | included | × | × | × | × | × | × | × |
| Anastaticeae | *Eremobium* | | included | × | × | × | × | × | × | × |
| Anastaticeae | *Farsetia* | | included | × | × | × | × | × | × | × |
| Anastaticeae | *Lachnocapsa* | | included | × | × | × | × | × | × | × |
| Anastaticeae | *Lobularia* | | included | × | × | × | × | × | × | × |
| Anastaticeae | *Marcus-kochia* | | included | × | × | × | × | × | × | × |
| Anastaticeae | *Maresia* | | included | × | × | × | × | × | × | × |
| Anastaticeae | *Morettia* | | included | × | × | × | × | × | - | × |
| Anastaticeae | *Notoceras* | | included | × | × | × | × | × | × | × |
| Anastaticeae | *Parolinia* | | included | × | × | × | × | × | × | × |
| Anchonieae | *Dvorakia* | | included | × | × | × | × | × | × | - |
| Anchonieae | *Eremoblastus* | | missing | - | - | - | - | - | - | - |
| Anchonieae | *Iskandera* | | included | × | × | × | × | × | × | × |
| Anchonieae | *Matthiola* | | included | × | × | × | × | × | - | × |
| Anchonieae | *Micrantha* | | included | × | × | × | × | × | × | × |
| Anchonieae | *Microstigma* | | included | × | × | × | × | × | × | × |
| Anchonieae | *Petiniotia* | | included | × | × | × | × | × | × | × |
| Anchonieae | *Sterigmostemum* | | included | × | × | × | × | × | × | × |
| Anchonieae | *Veselskya* | | included | × | × | × | × | × | - | × |
| Aphragmeae | *Aphragmus* | | included | × | × | × | × | × | × | × |
| Arabideae | *Abdra* | | included | × | × | × | × | × | × | - |
| Arabideae | *Acirostrum* | | included | × | × | × | × | × | × | - |
| Arabideae | *Arabis* | | included | × | × | × | × | × | × | × |
| Arabideae | *Arcyosperma* | | included | × | × | × | × | × | × | × |
| Arabideae | *Athysanus* | | included | × | × | × | × | × | × | × |
| Arabideae | *Aubrieta* | | included | × | × | × | × | × | × | × |
| Arabideae | *Baimashania* | | included | × | × | × | × | × | × | - |
| Arabideae | *Botschantzevia* | | included | × | × | × | × | × | × | - |
| Arabideae | *Dendroarabis* | | included | × | × | × | × | × | × | - |
| Arabideae | *Draba* | | included | × | × | × | × | × | × | × |
| Arabideae | *Drabella* | | included | × | × | × | × | × | × | × |
| Arabideae | *Pachyneurum* | | included | × | × | × | × | × | × | × |
| Arabideae | *Parryodes* | | included | × | × | × | × | × | × | × |
| Arabideae | *Pseudodraba* | | included | × | - | - | - | × | × | - |
| Arabideae | *Pseudoturritis* | | included | × | × | × | × | × | × | × |
| Arabideae | *Scapiarabis* | | included | × | × | × | × | × | × | × |
| Arabideae | *Sinoarabis* | | included | × | × | × | × | × | × | × |
| Arabideae | *Tomostima* | | included | × | × | × | × | × | × | × |
| Arabidopsideae trib. nov. | *Arabidopsis* | | included | × | × | × | × | × | × | × |
| Asperuginoideae trib. nov. | *Asperuginoides* | | included | × | × | × | × | × | × | × |
| Asteae | *Asta* | | included | × | × | × | × | × | × | × |
| Asteae | *Scoliaxon* | | included | × | × | × | × | × | × | × |
| Biscutelleae | *Biscutella* | | included | × | × | × | × | × | × | × |
| Biscutelleae | *Heldreichia* | | included | × | × | × | × | × | × | × |
| Biscutelleae | *Lunaria* | | included | × | × | × | × | × | × | × |
| Biscutelleae | *Megadenia* | | included | × | × | × | × | × | × | × |
| Biscutelleae | *Ricotia* | | included | × | × | × | × | × | × | × |

Table S3. Continued.

| Tribe | Genus name used in this study (following BrassiBase) | Currently accepted name (in case different from name used) | Overall sampling | Coalescent approach in ASTRAL | | | | | Supermatrix approach in IQ-TREE | |
|---|---|---|---|---|---|---|---|---|---|---|
| | | | | inclusive | strict | super-strict | super-strict by tribe | Pro | nuclear | plastome |
| Boechereae | *Anelsonia* | | included | × | × | × | × | × | × | × |
| Boechereae | *Boechera* | | included | × | × | × | × | × | × | × |
| Boechereae | *Borodinia* | | included | × | × | × | × | × | × | - |
| Boechereae | *Cusickiella* | | included | × | × | × | × | × | × | - |
| Boechereae | *Nevada* | | included | × | × | × | × | × | × | × |
| Boechereae | *Phoenicaulis* | | included | × | × | × | × | × | × | × |
| Boechereae | *Polyctenium* | | included | × | × | × | × | × | × | × |
| Boechereae | *Sandbergia* | | included | × | × | × | × | × | × | × |
| Boechereae | *Yosemitea* | | included | × | × | × | × | × | × | × |
| Brassiceae | *Ammosperma* | | included | × | × | × | × | × | × | - |
| Brassiceae | *Bivonaea* | | included | × | - | - | - | × | × | × |
| Brassiceae | *Brassica* | | included | × | × | × | × | × | × | × |
| Brassiceae | *Cakile* | | included | × | × | × | × | × | × | × |
| Brassiceae | *Carrichtera* | | included | × | × | × | × | × | × | × |
| Brassiceae | *Ceratocnemum* | | included | × | × | × | × | × | × | × |
| Brassiceae | *Coincya* | | included | × | × | × | × | × | × | × |
| Brassiceae | *Cordylocarpus* | | missing | - | - | - | - | - | - | - |
| Brassiceae | *Crambe* | | included | × | × | × | × | × | × | × |
| Brassiceae | *Crambella* | | included | × | × | × | × | × | × | × |
| Brassiceae | *Didesmus* | | included | × | × | × | × | × | × | - |
| Brassiceae | *Diplotaxis* | | included | × | × | × | × | × | × | × |
| Brassiceae | *Douepea* | | included | × | × | × | × | × | × | × |
| Brassiceae | *Enarthrocarpus* | | included | × | × | × | × | × | × | - |
| Brassiceae | *Eremophyton* | | included | × | × | × | × | × | × | × |
| Brassiceae | *Eruca* | | included | × | × | × | × | × | × | × |
| Brassiceae | *Erucaria* | | included | × | × | × | × | × | × | - |
| Brassiceae | *Erucastrum* | | included | × | × | × | × | × | × | × |
| Brassiceae | *Fezia* | | missing | - | - | - | - | - | - | - |
| Brassiceae | *Foleyola* | | included | × | × | × | × | × | × | - |
| Brassiceae | *Fortuynia* | | included | × | × | × | × | × | × | × |
| Brassiceae | *Guenthera* | | included | × | × | × | × | × | × | × |
| Brassiceae | *Guiraoa* | | included | × | × | × | × | × | × | × |
| Brassiceae | *Guiraoa* | | included | × | × | × | × | × | × | × |
| Brassiceae | *Hemicrambe* | | included | × | × | × | × | × | × | - |
| Brassiceae | *Henophyton* | | included | × | × | × | × | × | × | × |
| Brassiceae | *Hirschfeldia* | | included | × | × | × | × | × | × | × |
| Brassiceae | *Horwoodia* | | included | × | × | × | × | × | × | × |
| Brassiceae | *Kremeriella* | | included | × | - | - | - | × | × | × |
| Brassiceae | *Moricandia* | | included | × | × | × | × | × | × | × |
| Brassiceae | *Morisia* | | included | × | × | × | × | × | × | - |
| Brassiceae | *Muricaria* | | missing | - | - | - | - | - | - | - |
| Brassiceae | *Nasturtiopsis* | | included | × | × | × | × | × | × | - |
| Brassiceae | *Orychophragmus* | | included | × | - | - | - | × | × | × |
| Brassiceae | *Otocarpus* | | included | × | × | × | × | × | × | × |
| Brassiceae | *Physorhynchus* | | included | × | × | × | × | × | × | × |
| Brassiceae | *Pseuderucaria* | | included | × | × | × | × | × | × | × |
| Brassiceae | *Psychine* | | included | × | × | × | × | × | × | × |
| Brassiceae | *Quezeliantha* | | included | × | × | × | × | × | × | × |
| Brassiceae | *Quidproquo* | | included | × | × | × | × | × | × | - |
| Brassiceae | *Raffenaldia* | | included | × | × | × | × | × | × | × |
| Brassiceae | *Raphanus* | | included | × | × | × | × | × | × | × |
| Brassiceae | *Rapistrum* | | included | × | × | × | × | × | × | × |
| Brassiceae | *Rhamphospermum* | *Mutarda* | included | × | × | × | × | × | × | × |
| Brassiceae | *Rytidocarpus* | | missing | - | - | - | - | - | - | - |
| Brassiceae | *Savignya* | | included | × | × | × | × | × | × | × |
| Brassiceae | *Schouwia* | | included | × | × | × | × | × | × | × |
| Brassiceae | *Sinalliaria* | | included | × | × | × | × | × | × | × |
| Brassiceae | *Sinapidendron* | | included | × | × | × | × | × | × | - |
| Brassiceae | *Sinapis* | | included | × | × | × | × | × | × | × |
| Brassiceae | *Succowia* | | included | × | × | × | × | × | × | - |
| Brassiceae | *Vella* | | included | × | × | × | × | × | × | × |
| Brassiceae | *Zahora* | | included | × | × | × | × | × | × | × |
| Brassiceae | *Zilla* | | included | × | × | × | × | × | × | × |
| Buniadeae | *Bunias* | | included | × | × | × | × | × | × | × |
| Calepineae | *Calepina* | | included | × | × | × | × | × | × | × |
| Calepineae | *Goldbachia* | | included | × | × | × | × | × | × | × |
| Calepineae | *Leiocarpaea* | | included | × | × | × | × | × | × | × |
| Camelineae | *Camelina* | | included | × | × | × | × | × | × | × |
| Camelineae | *Capsella* | | included | × | × | × | × | × | × | × |
| Camelineae | *Catolobus* | | included | × | × | × | × | × | × | - |
| Camelineae | *chrysochamela* | | included | × | × | × | × | - | × | × |
| Camelineae | *Neslia* | | included | × | × | × | × | × | × | - |
| Camelineae | *Pseudoarabidopsis* | | included | × | × | × | × | × | × | × |
| Cardamineae | *Andrzeiowskia* | | included | × | × | × | × | × | × | × |
| Cardamineae | *Aplanodes* | | included | × | × | × | × | × | - | × |
| Cardamineae | *Armoracia* | | included | × | × | × | × | × | × | × |
| Cardamineae | *Barbarea* | | included | × | - | - | - | × | × | × |
| Cardamineae | *Bengt-jonsellia* | | missing | - | - | - | - | - | - | - |
| Cardamineae | *Cardamine* | | included | × | × | × | × | × | × | × |

Table S3. Continued.

| Tribe | Genus name used in this study (following BrassiBase) | Currently accepted name (in case different from name used) | Overall sampling | Coalescent approach in ASTRAL | | | | | Supermatrix approach in IQ-TREE | |
|---|---|---|---|---|---|---|---|---|---|---|
| | | | | inclusive | strict | super-strict | super-strict by tribe | Pro | nuclear | plastome |
| Cardamineae | *Iodanthus* | | included | × | × | × | × | × | × | × |
| Cardamineae | *Leavenworthia* | | included | × | × | × | × | × | × | × |
| Cardamineae | *Nasturtium* | | included | × | × | × | × | × | × | × |
| Cardamineae | *Ornithocarpa* | | included | × | × | × | × | × | × | × |
| Cardamineae | *Planodes* | | included | × | × | × | × | × | × | - |
| Cardamineae | *Rorippa* | | included | × | × | × | × | × | × | × |
| Cardamineae | *Selenia* | | included | × | × | × | × | × | × | - |
| Cardamineae | *Sisymbrella* | | included | × | × | × | × | × | × | - |
| Chamireae | *Chamira* | | included | × | × | × | × | × | × | × |
| Chorisporeae | *Chorispora* | | included | × | × | × | × | × | × | × |
| Chorisporeae | *Diptychocarpus* | | included | × | × | × | × | × | × | × |
| Chorisporeae | *Litwinowia* | | included | × | × | × | × | × | × | × |
| Chorisporeae | *Parrya* | | included | × | × | × | × | × | × | × |
| Cochlearieae | *Cochlearia* | | included | × | × | × | × | × | × | × |
| Cochlearieae | *Ionopsidium* | | included | × | × | × | × | × | × | × |
| Coluteocarpeae | *Eunomia* | | included | × | × | × | × | × | × | × |
| Coluteocarpeae | *Noccaea* | | included | × | × | × | × | × | × | × |
| Conringieae | *Conringia* | | included | × | × | × | × | × | × | × |
| Cremolobeae | *Aimara* | | included | × | × | × | × | × | × | × |
| Cremolobeae | *Cremolobus* | | included | × | × | × | × | × | × | × |
| Cremolobeae | *Menonvillea* | | included | × | × | × | × | × | × | × |
| Cremolobeae | *Yunkia* | | missing | - | - | - | - | - | - | - |
| Crucihimalayeae | *Crucihimalaya* | | included | × | × | × | × | × | × | × |
| Crucihimalayeae | *Ladakiella* | | included | × | × | × | × | × | × | × |
| Crucihimalayeae | *Pulvinatusia* | | missing | - | - | - | - | - | - | - |
| Descurainieae | *Descurainia* | | included | × | × | × | × | × | × | × |
| Descurainieae | *Hornungia* | | included | × | × | × | × | × | × | × |
| Descurainieae | *Ianhedgea* | | included | × | × | × | × | × | × | - |
| Descurainieae | *Robeschia* | | included | × | × | × | × | × | × | × |
| Descurainieae | *Trichotolinum* | | missing | - | - | - | - | - | - | - |
| Descurainieae | *Tropidocarpum* | | included | × | × | × | × | × | × | × |
| Dontostemoneae | *Clausia* | | included | × | × | × | × | × | × | × |
| Dontostemoneae | *Dontostemon* | | included | × | × | × | × | × | × | × |
| Erysimeae | *Erysimum* | | included | × | × | × | × | × | × | × |
| Euclidieae | *Anzhenxia* | | included | × | × | × | × | × | × | × |
| Euclidieae | *Atelanthera* | | included | × | × | × | × | × | × | × |
| Euclidieae | *Braya* | | included | × | × | × | × | × | × | × |
| Euclidieae | *Catenulina* | | included | × | × | × | × | × | × | × |
| Euclidieae | *Christolea* | | included | × | × | × | × | × | × | × |
| Euclidieae | *Cryptospora* | | included | × | × | × | × | × | × | × |
| Euclidieae | *Cymatocarpus* | | included | × | × | × | × | × | × | - |
| Euclidieae | *Dichasianthus* | | included | × | × | × | × | × | × | × |
| Euclidieae | *Dilophia* | | included | × | × | × | × | × | × | × |
| Euclidieae | *Euclidium* | | included | × | × | × | × | × | × | × |
| Euclidieae | *Lachnoloma* | | included | × | × | × | × | × | × | - |
| Euclidieae | *Leiospora* | | included | × | × | × | × | × | × | × |
| Euclidieae | *Lepidostemon* | | included | × | × | × | × | × | × | - |
| Euclidieae | *Leptaleum* | | included | × | × | × | × | × | × | × |
| Euclidieae | *Metashangrilaia* | | included | × | × | × | × | × | × | × |
| Euclidieae | *Neotorularia* | | included | × | × | × | × | × | - | × |
| Euclidieae | *Octoceras* | | missing | - | - | - | - | - | - | - |
| Euclidieae | *Pycnoplinthopsis* | | missing | - | - | - | - | - | - | - |
| Euclidieae | *Pycnoplinthus* | | included | × | × | × | × | × | × | × |
| Euclidieae | *Rhammatophyllum* | | included | × | × | × | × | × | × | × |
| Euclidieae | *Rudolf-kamelinia* | | included | × | × | × | × | × | × | × |
| Euclidieae | *Shangrilaia* | | missing | - | - | - | - | - | - | - |
| Euclidieae | *Sisymbriopsis* | | included | × | × | × | × | × | × | × |
| Euclidieae | *Solms-laubachia* | | included | × | × | × | × | × | × | × |
| Euclidieae | *Spryginia* | | included | × | × | × | × | × | × | - |
| Euclidieae | *Streptoloma* | | included | × | × | × | × | × | × | - |
| Euclidieae | *Strigosella* | | included | × | × | × | × | × | × | × |
| Euclidieae | *Synstemon* | | included | × | × | × | × | × | × | × |
| Euclidieae | *Tetracme* | | included | × | × | × | × | × | × | × |
| Euclidieae | *Vvedenskyella* | | missing | - | - | - | - | - | - | - |
| Eudemeae | *Alshehbazia* | | included | × | × | × | × | × | × | × |
| Eudemeae | *Aschersoniodoxa* | | included | × | × | × | × | × | × | - |
| Eudemeae | *Dactylocardamum* | | included | × | × | × | × | × | × | × |
| Eudemeae | *Eudema* | | included | × | × | × | × | × | × | × |
| Eudemeae | *Gongylis* | | missing | - | - | - | - | - | - | - |
| Eudemeae | *Onuris* | | included | - | - | - | - | - | - | × |
| Eudemeae | *Xerodraba* | | included | × | - | - | - | × | × | - |
| Eutremeae | *Eutrema* | | included | × | × | × | × | × | × | × |
| Fourraeeae | *Fourraea* | | included | × | × | × | × | × | × | × |
| Fourraeeae | *Hurkaea* | | included | × | × | × | × | × | × | × |
| Halimolobeae | *Exhalimolobos* | | included | × | × | × | × | × | × | × |
| Halimolobeae | *Halimolobos* | | included | × | × | × | × | × | × | × |
| Halimolobeae | *Mancoa* | | included | × | × | × | × | × | × | × |
| Halimolobeae | *Pennellia* | | included | × | × | × | × | × | × | × |
| Halimolobeae | *Sphaerocardamum* | | included | × | × | × | × | × | × | × |

Table S3. Continued.

| Tribe | Genus name used in this study (following BrassiBase) | Currently accepted name (in case different from name used) | Overall sampling | Coalescent approach in ASTRAL | | | | | Supermatrix approach in IQ-TREE | |
|---|---|---|---|---|---|---|---|---|---|---|
| | | | | inclusive | strict | super-strict | super-strict by tribe | Pro | nuclear | plastome |
| Heliophileae | *Heliophila* | | included | × | × | × | × | × | × | × |
| Hemilophieae trib. nov. | *Dipoma* | | included | × | × | × | × | × | × | × |
| Hemilophieae trib. nov. | *Hemilophia* | | included | × | × | × | × | × | × | × |
| Hesperideae | *Hesperis* | | included | × | × | × | × | × | × | × |
| Hillielleae | *Hilliella* | | included | - | - | - | - | - | - | × |
| Iberideae | *Iberis* | | included | × | × | × | × | × | × | × |
| Iberideae | *Teesdalia* | | included | × | × | × | × | × | × | × |
| Isatideae | *Chartoloma* | | included | × | - | - | - | × | × | × |
| Isatideae | *Glastaria* | | included | × | × | × | × | × | × | - |
| Isatideae | *Iljinskaea* | | included | × | × | × | × | × | × | × |
| Isatideae | *Isatis* | | included | × | × | × | × | × | × | × |
| Isatideae | *Myagrum* | | included | × | × | × | × | × | × | × |
| Isatideae | *Schimpera* | | included | × | × | × | × | × | × | × |
| Kernereae | *Kernera* | | included | × | × | × | × | × | × | × |
| Kernereae | *Petrocallis* | | included | × | × | × | × | × | × | × |
| Kernereae | *Rhizobotrya* | | included | × | × | × | × | × | × | × |
| Lepidieae | *Delpinophytum* | | included | × | × | × | × | × | × | × |
| Lepidieae | *Lepidium* | | included | × | × | × | × | × | × | × |
| Lepidieae | *Lithodraba* | *Lepidium* | missing | - | - | - | - | - | - | - |
| Malcolmieae | *Malcolmia* | | included | × | × | × | × | × | × | × |
| Megacarpaeeae | *Megacarpaea* | | included | × | × | × | × | × | × | × |
| Megacarpaeeae | *Pugionium* | | included | × | × | × | × | × | × | × |
| Microlepidieae | *Arabidella* | | included | × | × | × | × | × | × | × |
| Microlepidieae | *Ballantinia* | | included | × | × | × | × | × | × | - |
| Microlepidieae | *Blennodia* | | included | × | × | × | × | × | × | - |
| Microlepidieae | *Carinavalva* | | included | × | × | × | × | × | × | × |
| Microlepidieae | *Cuphonotus* | *Lemphoria* | included | × | × | × | × | × | × | × |
| Microlepidieae | *Drabastrum* | | included | × | × | × | × | × | × | - |
| Microlepidieae | *Geococcus* | | included | × | × | × | × | × | × | × |
| Microlepidieae | *Harmsiodoxa* | | included | × | × | × | × | × | × | - |
| Microlepidieae | *Irenepharsus* | | included | × | × | × | × | × | × | × |
| Microlepidieae | *Menkea* | | included | × | × | × | × | × | × | - |
| Microlepidieae | *Microlepidium* | | included | × | × | × | × | × | × | - |
| Microlepidieae | *Pachycladon* | | included | × | × | × | × | × | × | × |
| Microlepidieae | *Pachymitus* | | included | × | × | × | × | × | × | × |
| Microlepidieae | *Phlegmatospermum* | | included | × | × | × | × | × | × | - |
| Microlepidieae | *Scambopus* | | included | × | × | × | × | × | × | × |
| Microlepidieae | *Stenopetalum* | | included | × | × | × | × | × | × | × |
| Notothlaspideae | *Notothlaspi* | | included | × | × | × | × | × | × | × |
| Oreophytoneae | *Murbeckiella* | | included | × | × | × | × | × | - | × |
| Oreophytoneae | *Oreophyton* | | included | × | × | × | × | × | × | × |
| Physarieae | *Dimorphocarpa* | | included | × | × | × | × | × | × | × |
| Physarieae | *Dithyrea* | | included | × | × | × | × | × | × | × |
| Physarieae | *Lyrocarpa* | | included | × | × | × | × | × | × | × |
| Physarieae | *Nerisyrenia* | | included | × | × | × | × | × | × | × |
| Physarieae | *Paysonia* | | included | × | × | × | × | - | × | × |
| Physarieae | *Physaria* | | included | × | × | × | × | - | × | × |
| Physarieae | *Synthlipsis* | | included | × | × | × | × | × | × | × |
| Plagiolobeae | *Plagioloba* | | included | × | × | × | × | × | × | × |
| Schizopetaleae | *Atacama* | | included | × | × | × | × | × | × | × |
| Schizopetaleae | *Mathewsia* | | included | × | × | × | × | × | × | × |
| Schizopetaleae | *Schizopetalon* | | included | × | × | × | × | × | × | × |
| Schrenkielleae trib. nov. | *Schrenkiella* | | included | × | × | × | × | × | × | × |
| Shehbazieae | *Shehbazia* | | included | × | - | - | - | × | × | × |
| Sisymbrieae | *Sisymbrium* | | included | × | × | × | × | × | × | × |
| Smelowskieae | *Smelowskia* | | included | × | × | × | × | × | × | × |
| Stevenieae | *Macropodium* | | included | × | × | × | × | × | × | × |
| Stevenieae | *Stevenia* | | included | × | × | × | × | × | × | × |
| Subularieae trib. nov. | *Idahoa* | | included | × | × | × | × | × | × | × |
| Subularieae trib. nov. | *Subularia* | | included | × | × | × | × | × | - | × |
| Thelypodieae | *Caulanthus* | | missing | - | - | - | - | - | - | - |
| Thelypodieae | *Chaunanthus* | | included | × | × | × | × | × | × | × |
| Thelypodieae | *Chilocardamum* | | included | × | × | × | × | × | × | × |
| Thelypodieae | *Chlorocrambe* | | included | × | × | × | × | × | × | - |
| Thelypodieae | *Dictyophragmus* | | included | × | - | - | - | × | × | - |
| Thelypodieae | *Dryopetalon* | | included | × | × | × | × | × | × | × |
| Thelypodieae | *Englerocharis* | | included | × | × | × | × | × | × | - |
| Thelypodieae | *Guillenia* | | missing | - | - | - | - | - | - | - |
| Thelypodieae | *Hesperidanthus* | | included | × | × | × | × | × | × | × |
| Thelypodieae | *Hollermayera* | | included | × | × | × | × | × | × | × |
| Thelypodieae | *Ivania* | | included | × | × | × | × | × | × | × |
| Thelypodieae | *Machaerophorus* | | missing | - | - | - | - | - | - | - |
| Thelypodieae | *Mostacillastrum* | | included | × | × | × | × | × | × | × |
| Thelypodieae | *Neuontobotrys* | | included | × | × | × | × | × | × | × |
| Thelypodieae | *Parodiodoxa* | | included | × | × | × | × | × | × | × |
| Thelypodieae | *Petroravenia* | | missing | - | - | - | - | - | - | - |
| Thelypodieae | *Phlebolobium* | | included | × | × | × | × | × | × | - |
| Thelypodieae | *Phravenia* | | included | × | × | × | × | × | × | × |
| Thelypodieae | *Polypsecadium* | | included | × | × | × | × | × | × | × |

Table S3. Continued.

| Tribe | Genus name used in this study (following BrassiBase) | Currently accepted name (in case different from name used) | Overall sampling | Coalescent approach in ASTRAL | | | | | Supermatrix approach in IQ-TREE | |
|---|---|---|---|---|---|---|---|---|---|---|
| | | | | inclusive | strict | super-strict | super-strict by tribe | Pro | nuclear | plastome |
| Thelypodieae | *Pringlea* | | included | × | - | - | - | × | × | × |
| Thelypodieae | *Pterygiosperma* | | missing | - | - | - | - | - | - | - |
| Thelypodieae | *Raphanorhyncha* | | included | × | × | × | × | × | × | × |
| Thelypodieae | *Romanschulzia* | | included | × | × | × | × | × | × | × |
| Thelypodieae | *Sarcodraba* | | included | × | - | - | - | × | × | × |
| Thelypodieae | *Sibara* | | included | × | × | × | × | × | × | × |
| Thelypodieae | *Sibaropsis* | | missing | - | - | - | - | - | - | - |
| Thelypodieae | *Stanleya* | | included | × | × | × | × | × | × | × |
| Thelypodieae | *Stenodraba* | | missing | - | - | - | - | - | - | - |
| Thelypodieae | *Streptanthella* | | missing | - | - | - | - | - | - | - |
| Thelypodieae | *Streptanthus* | | included | × | × | × | × | × | × | × |
| Thelypodieae | *Terraria* | | missing | - | - | - | - | - | - | - |
| Thelypodieae | *Thelypodiopsis* | | included | × | - | - | - | × | - | - |
| Thelypodieae | *Thelypodium* | | included | - | - | - | - | - | - | × |
| Thelypodieae | *Thysanocarpus* | | included | × | × | × | × | × | × | × |
| Thelypodieae | *Warea* | | included | × | × | × | × | × | × | - |
| Thelypodieae | *Weberbauera* | | included | × | × | × | × | × | × | × |
| Thelypodieae | *Zuloagocardamum* | | included | × | × | × | × | × | × | × |
| Thlaspideae | *Alliaria* | | included | × | × | × | × | × | × | × |
| Thlaspideae | *Didymophysa* | | included | × | × | × | × | × | × | × |
| Thlaspideae | *Graellsia* | | included | × | - | - | - | × | × | × |
| Thlaspideae | *Lysakia* | | included | × | × | × | × | × | × | × |
| Thlaspideae | *Mummenhoffia* | | included | × | × | × | × | × | × | × |
| Thlaspideae | *Pachyphragma* | | included | × | × | × | × | × | × | × |
| Thlaspideae | *Parlatoria* | | included | × | × | × | × | × | × | × |
| Thlaspideae | *Peltaria* | | included | × | × | × | × | × | × | × |
| Thlaspideae | *Peltariopsis* | | included | × | × | × | × | × | × | - |
| Thlaspideae | *Pseudocamelina* | | included | × | × | × | × | × | × | × |
| Thlaspideae | *Pseudovesicaria* | | included | × | × | × | × | × | × | - |
| Thlaspideae | *Sobolewskia* | | included | × | × | × | × | × | × | × |
| Thlaspideae | *Thlaspi* | | included | × | × | × | × | × | × | × |
| Turritideae | *Turritis* | | included | × | × | × | × | × | - | × |
| Yinshanieae | *Yinshania* | | included | × | × | × | × | × | - | × |
| **Additional genera sampled following BrassiBase, but not currently accepted.** | | | | | | | | | | |
| Coluteocarpeae | *Callothlaspi* | *Noccaea* | included | × | × | × | × | × | × | × |
| Coluteocarpeae | *Friedrichkarlmeyeria* | *Noccaea* | included | × | × | × | × | × | × | × |
| Coluteocarpeae | *Ihsanalshehbazia* | *Noccaea* | included | × | × | × | × | × | × | - |
| Coluteocarpeae | *Kotschyella* | *Noccaea* | included | × | × | × | × | × | × | × |
| Coluteocarpeae | *Masmenia* | *Noccaea* | included | × | × | × | × | × | × | × |
| Coluteocarpeae | *Microthlaspi* | *Noccaea* | included | × | × | × | × | × | × | × |
| Coluteocarpeae | *Neurotropis* | *Noccaea* | included | × | × | × | × | × | × | × |
| Coluteocarpeae | *Noccidium* | *Noccaea* | included | × | × | × | × | × | × | × |
| Coluteocarpeae | *Pseudosempervivum* | *Noccaea* | included | × | × | × | × | × | × | - |
| Coluteocarpeae | *Thlaspiceras* | *Noccaea* | included | × | × | × | × | × | × | - |
| Coluteocarpeae | *Vania* | *Noccaea* | included | × | × | × | × | × | × | × |
| Eudemeae | *Brayopsis* | *Eudema* | included | × | × | × | × | × | × | × |
| Lepidieae | *Cyphocardamum* | *Lepidium* | included | × | × | × | × | × | × | × |
| Thlaspideae | *Elburzia* | *Didymophysa* | included | × | - | - | - | × | × | - |
| **Genera not yet included in our sampling, to be published shortly.** | | | | | | | | | | |
| Eudemeae | *Ancashia* | | | | | | | | | |
| Eudemeae | *Borealandina* | | | | | | | | | |
| Thelypodieae | *Stenodrabopsis* | | | | | | | | | |

Table S4. HybPhaser output per sample, with sequencing and mapping success (total sequence length and number of genes recovered) and metrics on allele divergence (AD) and locus heterozygosity (LH). Values per sample and based on the total dataset (i.e. from all genes) and split by bait set (A353: Angiosperms353; B764: Brassicaceae-specific set).

| SAMPLE DETAILS | | | | ASTRAL TAXON OCCUPANCY | | | | | RECOVERED TOTAL LENGTH (BP) | | | RECOVERED TOTAL LENGTH OF TARGET LENGTH (%) | | | # PARALOG CALLS | | | # LOCI RECOVERED | | | ALLELE DIVERGENCE | | | | LOCUS HETEROZYGOSITY | | | |
|---|---|---|---|---|---|---|---|---|---|---|---|---|---|---|---|---|---|---|---|---|---|---|---|---|---|---|---|---|
| Main lineage | Tribe | Species | Sample | inclusive | strict | superstrict | superstrict by tribe | A-Pro | Total | B764 | A353 | Total | B764 | A353 | Total | B764 | A353 | Total | B764 | A353 | Total (%) | B764 (%) | A353(%) | B764/A353 | Total (%) | B764 (%) | A353(%) | B764/A353 |
| basal | Aethionemeae | *Aethionema acarii* | S1148 | 924 | 889 | 265 | 666 | 5,856 | 1,033,344 | 781,404 | 251,940 | 87.5 | 88 | 86.3 | 68 | 41 | 27 | 935 | 627 | 308 | 0.67 | 0.67 | 0.67 | 1.00 | 80.53 | 80.70 | 80.19 | 1.01 |
| basal | Aethionemeae | *Aethionema demirizii* | S1184 | 842 | 803 | 233 | 500 | 5,513 | 922,092 | 695,094 | 226,998 | 78.1 | 78.2 | 77.8 | 56 | 32 | 24 | 856 | 564 | 292 | 1.51 | 1.53 | 1.44 | 1.06 | 92.52 | 93.79 | 90.07 | 1.04 |
| basal | Aethionemeae | *Aethionema erinaceum* | S0890 | 693 | 666 | 184 | 111 | 4,482 | 613,185 | 481,137 | 132,048 | 52 | 54.2 | 45.2 | 23 | 11 | 12 | 727 | 487 | 240 | 1.89 | 1.82 | 2.15 | 0.85 | 78.40 | 80.49 | 74.17 | 1.09 |
| basal | Aethionemeae | *Aethionema maraschicum* | S1100 | 898 | 860 | 249 | 741 | 5,747 | 988,170 | 749,481 | 238,689 | 83.7 | 84.4 | 81.8 | 69 | 38 | 31 | 901 | 605 | 296 | 0.28 | 0.27 | 0.29 | 0.94 | 52.39 | 51.90 | 53.38 | 0.97 |
| basal | Aethionemeae | *Aethionema munzurense* | S1180 | 907 | 877 | 257 | 541 | 5,714 | 1,011,651 | 768,816 | 242,835 | 85.7 | 86.5 | 83.2 | 96 | 59 | 37 | 886 | 599 | 287 | 0.25 | 0.25 | 0.26 | 0.93 | 64.11 | 66.11 | 59.93 | 1.10 |
| basal | Aethionemeae | *Aethionema orbiculatum* | S1261 | 818 | 784 | 220 | 267 | 5,187 | 862,221 | 677,376 | 184,845 | 73 | 76.2 | 63.3 | 25 | 16 | 9 | 857 | 577 | 280 | 3.72 | 3.59 | 4.23 | 0.85 | 85.41 | 85.62 | 85.00 | 1.01 |
| basal | Aethionemeae | *Aethionema saxatile* | S0735 | 933 | 895 | 273 | 238 | 5,936 | 1,029,714 | 764,757 | 264,957 | 87.2 | 86.1 | 90.8 | 62 | 29 | 33 | 945 | 631 | 314 | 0.17 | 0.16 | 0.17 | 0.94 | 46.14 | 44.37 | 49.68 | 0.89 |
| basal | Aethionemeae | *Aethionema stenopterum* | S1223 | 886 | 845 | 247 | 459 | 5,680 | 964,179 | 734,088 | 230,091 | 81.7 | 82.6 | 78.8 | 72 | 44 | 28 | 879 | 593 | 286 | 0.42 | 0.42 | 0.39 | 1.09 | 76.91 | 78.41 | 73.78 | 1.06 |
| I | Alyssopsideae | *Alyssopsis mollis* | S1460 | 1012 | 1011 | 296 | 595 | 5,674 | 1,133,415 | 859,797 | 273,618 | 96 | 96.8 | 93.7 | 68 | 32 | 36 | 1006 | 694 | 312 | 0.49 | 0.48 | 0.54 | 0.88 | 77.73 | 75.65 | 82.37 | 0.92 |
| I | Alyssopsideae | *Calymmatium draboides* | S1235sl | 1007 | 1005 | 292 | 527 | 5,560 | 1,133,901 | 858,933 | 274,968 | 96.1 | 96.7 | 94.2 | 44 | 19 | 25 | 1026 | 705 | 321 | 0.04 | 0.02 | 0.11 | 0.21 | 23.88 | 15.18 | 42.99 | 0.35 |
| I | Alyssopsideae | *Dielsiocharis kotschyi* | S0316sl | 504 | - | - | - | 2,912 | 227,094 | 168,816 | 58,278 | 19.2 | 19 | 20 | 8 | 3 | 5 | 594 | 386 | 208 | 0.10 | 0.11 | 0.10 | 1.04 | 17.00 | 18.65 | 13.94 | 1.34 |
| I | Alyssopsideae | *Olimarabidopsis pumila* | S0809 | 1001 | 997 | 287 | 178 | 5,930 | 1,066,074 | 801,546 | 264,528 | 90.3 | 90.2 | 90.6 | 39 | 18 | 21 | 1028 | 704 | 324 | 2.76 | 2.78 | 2.70 | 1.03 | 99.61 | 99.43 | 100 | 0.99 |
| I | Arabidopsideae trib. nov. | *Arabidopsis halleri* | S1381 | 1002 | 1001 | 292 | 145 | 5,910 | 1,123,200 | 860,694 | 262,506 | 95.2 | 96.9 | 89.9 | 70 | 27 | 43 | 994 | 690 | 304 | 0.54 | 0.49 | 0.67 | 0.74 | 70.22 | 67.83 | 75.66 | 0.90 |
| I | Arabidopsideae trib. nov. | *Arabidopsis thaliana* | S1211 | 999 | 993 | 287 | 485 | 5,883 | 1,130,118 | 859,833 | 270,285 | 95.7 | 96.8 | 92.6 | 29 | 8 | 21 | 1032 | 707 | 325 | 0.04 | 0.04 | 0.07 | 0.50 | 14.05 | 11.74 | 19.08 | 0.62 |
| I | Boechereae | *Anelsonia eurycarpa* | S0282 | 995 | 995 | 289 | 256 | 5,634 | 1,131,672 | 860,355 | 271,317 | 95.9 | 96.8 | 92.9 | 74 | 36 | 38 | 985 | 680 | 305 | 0.13 | 0.12 | 0.17 | 0.69 | 36.45 | 30.74 | 49.18 | 0.63 |
| I | Boechereae | *Boechera holboellii* | S1227 | 1001 | 999 | 293 | 532 | 5,747 | 1,124,112 | 855,378 | 268,734 | 95.2 | 96.3 | 92.1 | 37 | 16 | 21 | 1027 | 702 | 325 | 0.17 | 0.13 | 0.31 | 0.41 | 28.72 | 22.65 | 41.85 | 0.54 |
| I | Boechereae | *Boechera yorkii* | S1369 | 1005 | 1002 | 291 | 311 | 5,851 | 1,131,015 | 857,286 | 273,729 | 95.8 | 96.5 | 93.8 | 50 | 29 | 21 | 1018 | 693 | 325 | 0.54 | 0.45 | 0.84 | 0.53 | 56.29 | 51.08 | 67.38 | 0.76 |
| I | Boechereae | *Borodinia macrophylla* | S0690 | 927 | 916 | 255 | 62 | 5,229 | 911,871 | 703,953 | 207,918 | 77.3 | 79.2 | 71.2 | 92 | 43 | 49 | 905 | 625 | 280 | 0.23 | 0.23 | 0.26 | 0.88 | 67.40 | 66.88 | 68.57 | 0.98 |
| I | Boechereae | *Cusickiella douglasii* | S0293 | 952 | 934 | 272 | 240 | 5,394 | 914,424 | 693,531 | 220,893 | 77.5 | 78.1 | 75.7 | 100 | 55 | 45 | 916 | 623 | 293 | 0.44 | 0.43 | 0.45 | 0.96 | 77.51 | 76.40 | 79.86 | 0.96 |
| I | Boechereae | *Nevada holmgrenii* | S0400 | 960 | 957 | 274 | 509 | 5,479 | 1,076,889 | 828,903 | 247,986 | 91.2 | 93.3 | 84.9 | 62 | 29 | 33 | 963 | 660 | 303 | 0.10 | 0.07 | 0.17 | 0.43 | 38.84 | 31.52 | 54.79 | 0.58 |
| I | Boechereae | *Phoenicaulis cheiranthoides* | S0268 | 973 | 971 | 280 | 250 | 5,550 | 1,115,238 | 849,438 | 265,800 | 94.5 | 95.6 | 91 | 81 | 38 | 43 | 953 | 655 | 298 | 0.65 | 0.65 | 0.65 | 1.00 | 91.61 | 91.30 | 92.28 | 0.99 |
| I | Boechereae | *Polyctenium fremontii* | S0301 | 1003 | 1000 | 289 | 260 | 5,661 | 1,104,399 | 859,023 | 245,376 | 93.6 | 96.7 | 84.1 | 76 | 46 | 30 | 990 | 680 | 310 | 0.54 | 0.54 | 0.52 | 1.04 | 86.67 | 86.62 | 86.77 | 1.00 |
| I | Boechereae | *Sandbergia whitedii* | S_670 | 1014 | 1011 | 296 | 263 | 5,773 | 1,128,732 | 857,628 | 271,104 | 95.6 | 96.5 | 92.9 | 76 | 40 | 36 | 1000 | 688 | 312 | 0.65 | 0.64 | 0.69 | 0.93 | 83.90 | 82.85 | 86.22 | 0.96 |
| I | Boechereae | *Yosemitea repanda* | S0763 | 973 | 973 | 278 | 250 | 5,571 | 1,126,308 | 852,651 | 273,657 | 95.4 | 96 | 93.7 | 42 | 17 | 25 | 994 | 675 | 319 | 0.12 | 0.09 | 0.23 | 0.37 | 27.06 | 21.48 | 38.87 | 0.55 |
| I | Camelineae I | *Camelina sativa* | S1204 | 997 | 994 | 287 | 483 | 5,649 | 1,088,991 | 829,599 | 259,392 | 92.3 | 93.4 | 88.9 | 31 | 19 | 12 | 1031 | 699 | 332 | 3.42 | 3.46 | 3.29 | 1.05 | 99.13 | 99.00 | 99.40 | 1.00 |
| I | Camelineae I | *Capsella bursa-pastoris* | S0670 | 1000 | 996 | 289 | 77 | 5,571 | 1,093,731 | 833,265 | 260,466 | 92.7 | 93.8 | 89.2 | 45 | 21 | 24 | 1019 | 697 | 322 | 1.84 | 1.85 | 1.80 | 1.03 | 98.43 | 98.13 | 99.07 | 0.99 |
| I | Camelineae I | *Catolobus pendulus* | S0718 | 999 | 998 | 294 | 77 | 5,852 | 1,075,941 | 817,524 | 258,417 | 91.2 | 92 | 88.5 | 37 | 17 | 20 | 1029 | 703 | 326 | 2.80 | 2.88 | 2.55 | 1.13 | 98.15 | 97.87 | 98.77 | 0.99 |
| I | Camelineae I | *Neslia paniculata* | S0641 | 1000 | 997 | 289 | 538 | 5,670 | 1,135,269 | 860,682 | 274,587 | 96.2 | 96.9 | 94.1 | 43 | 17 | 26 | 1019 | 700 | 319 | 0.05 | 0.03 | 0.11 | 0.31 | 26.20 | 18.14 | 43.89 | 0.41 |
| I | Camelineae III | *Chrysochamela velutina* | S1318-2 | 1000 | 991 | 287 | 292 | 5,864 | 1,085,445 | 838,713 | 246,732 | 92 | 94.4 | 84.5 | 44 | 19 | 25 | 1021 | 698 | 323 | 1.38 | 1.37 | 1.44 | 0.95 | 96.08 | 96.28 | 95.67 | 1.01 |
| I | Camelineae III | *Pseudoarabidopsis toxophylla* | S0721sl | 966 | 966 | 278 | 67 | 5,703 | 1,108,434 | 842,394 | 266,040 | 93.9 | 94.8 | 91.1 | 64 | 34 | 30 | 967 | 653 | 314 | 1.51 | 1.47 | 1.64 | 0.90 | 96.17 | 95.87 | 96.82 | 0.99 |
| I | Cardamineae | *Andrzeiowskia cardamine* | SRR8528391 | 723 | 729 | 199 | 218 | 4,555 | 919,329 | 883,404 | 35,925 | 77.9 | 99.4 | 12.3 | 30 | 28 | 2 | 742 | 700 | 42 | 0.04 | 0.04 | 0.10 | 0.42 | 25.34 | 24.57 | 38.10 | 0.64 |
| I | Cardamineae | *Aplanodes doidgeana* | S0904 | 654 | 650 | 178 | 293 | 4,748 | 755,355 | 755,355 | 0 | 64 | 85 | 0 | 7 | 7 | 0 | 693 | 693 | 0 | 4.31 | 4.31 | NA | NA | 98.56 | 98.56 | NA | NA |
| I | Cardamineae | *Armoracia rusticana* | S0671 | 970 | 964 | 279 | 67 | 5,743 | 1,063,902 | 809,772 | 254,130 | 90.1 | 91.2 | 87.1 | 17 | 9 | 8 | 1023 | 695 | 328 | 3.99 | 4.04 | 3.84 | 1.05 | 96.58 | 96.40 | 96.95 | 0.99 |
| I | Cardamineae | *Barbarea vulgaris* | S0734sl | 380 | - | - | - | 2,169 | 204,897 | 155,724 | 49,173 | 17.4 | 17.5 | 16.8 | 9 | 5 | 4 | 426 | 286 | 140 | 0.16 | 0.14 | 0.24 | 0.57 | 22.30 | 22.73 | 21.43 | 1.06 |
| I | Cardamineae | *Cardamine pratensis* | S0733sl | 922 | 906 | 254 | 230 | 5,219 | 881,328 | 679,386 | 201,942 | 74.7 | 76.5 | 69.2 | 73 | 41 | 32 | 915 | 623 | 292 | 1.11 | 1.10 | 1.12 | 0.98 | 89.40 | 89.25 | 89.73 | 0.99 |
| I | Cardamineae | *Iodanthus pinnatifidus* | S0298 | 975 | 973 | 282 | 255 | 5,880 | 1,058,793 | 798,540 | 260,253 | 89.7 | 89.9 | 89.1 | 13 | 9 | 4 | 1036 | 695 | 341 | 5.46 | 5.57 | 5.12 | 1.09 | 98.94 | 99.28 | 98.24 | 1.01 |
| I | Cardamineae | *Leavenworthia aurea* | S1276 | 969 | 947 | 271 | 511 | 5,661 | 1,056,444 | 791,907 | 264,537 | 89.5 | 89.1 | 90.6 | 51 | 36 | 15 | 985 | 658 | 327 | 4.01 | 4.09 | 3.79 | 1.08 | 96.35 | 95.90 | 97.25 | 0.99 |
| I | Cardamineae | *Nasturtium officinale* | S0593 | 952 | 943 | 265 | 506 | 5,435 | 976,086 | 740,646 | 235,440 | 82.7 | 83.4 | 80.6 | 7 | 4 | 3 | 1024 | 691 | 333 | 3.59 | 3.67 | 3.33 | 1.10 | 87.50 | 87.41 | 87.69 | 1.00 |
| I | Cardamineae | *Ornithocarpa fimbriata* | S1541sl | 982 | 977 | 280 | 537 | 5,770 | 1,034,760 | 795,549 | 239,211 | 87.7 | 89.5 | 81.9 | 1 | 0 | 0 | 1059 | 715 | 344 | 4.35 | 4.47 | 3.96 | 1.13 | 93.86 | 94.13 | 93.31 | 1.01 |
| I | Cardamineae | *Planodes virginicum* | S0427 | 994 | 992 | 285 | 519 | 5,616 | 1,131,333 | 859,617 | 271,716 | 95.8 | 96.8 | 93.1 | 50 | 19 | 31 | 1007 | 693 | 314 | 0.05 | 0.04 | 0.10 | 0.36 | 23.54 | 17.46 | 36.94 | 0.47 |
| I | Cardamineae | *Rorippa sylvestris* | S0672 | 940 | 939 | 264 | 63 | 5,406 | 1,096,467 | 830,808 | 265,659 | 92.9 | 93.5 | 91 | 50 | 21 | 29 | 952 | 643 | 309 | 2.18 | 2.19 | 2.17 | 1.01 | 97.16 | 97.05 | 97.41 | 1.00 |
| I | Cardamineae | *Selenia aurea* | S0893 | 661 | 650 | 177 | 308 | 3,884 | 749,085 | 564,819 | 184,266 | 63.5 | 63.6 | 63.1 | 15 | 4 | 11 | 703 | 452 | 251 | 2.08 | 1.97 | 2.42 | 0.81 | 90.33 | 90.49 | 90.04 | 1.00 |
| I | Cardamineae | *Sisymbrella aspera* | S0376 | 930 | 926 | 258 | 234 | 5,326 | 935,715 | 728,748 | 206,967 | 79.3 | 82 | 70.9 | 63 | 35 | 28 | 931 | 659 | 272 | 0.24 | 0.23 | 0.27 | 0.84 | 63.48 | 62.06 | 66.91 | 0.93 |
| I | Crucihimalayeae | *Crucihimalaya himalaica* | SRR8528343 | 719 | 724 | 198 | 688 | 4,551 | 916,290 | 882,558 | 33,732 | 77.6 | 99.3 | 11.6 | 35 | 31 | 4 | 737 | 697 | 40 | 0.06 | 0.05 | 0.13 | 0.40 | 25.64 | 24.68 | 42.50 | 0.58 |
| I | Crucihimalayeae | *Ladakiella klimesii* | S0756 | 1001 | 999 | 291 | 256 | 5,704 | 1,133,634 | 860,334 | 273,300 | 96 | 96.8 | 93.6 | 30 | 15 | 15 | 1033 | 702 | 331 | 0.28 | 0.20 | 0.51 | 0.39 | 30.49 | 21.94 | 48.64 | 0.45 |
| I | Descurainieae | *Descurainia sophia* | S1407 | 978 | 973 | 280 | 813 | 5,914 | 974,628 | 729,216 | 245,412 | 82.6 | 82.1 | 84.1 | 26 | 14 | 12 | 1024 | 697 | 327 | 4.05 | 4.14 | 3.78 | 1.09 | 100 | 100 | 100 | 1.00 |
| I | Descurainieae | *Hornungia petraea* | S0558 | 1003 | 992 | 287 | 526 | 6,010 | 1,131,447 | 857,850 | 273,597 | 95.9 | 96.6 | 93.7 | 37 | 20 | 17 | 1029 | 702 | 327 | 0.03 | 0.02 | 0.06 | 0.38 | 15.65 | 9.12 | 29.66 | 0.31 |
| I | Descurainieae | *Ianhedgea minutiflora* | S0412sl | 974 | 960 | 279 | 499 | 5,731 | 1,007,868 | 769,536 | 238,332 | 85.4 | 86.6 | 81.6 | 76 | 39 | 37 | 966 | 659 | 307 | 0.32 | 0.31 | 0.35 | 0.88 | 66.25 | 66.01 | 66.78 | 0.99 |
| I | Descurainieae | *Robeschia schimperi* | S0742 | 982 | 979 | 283 | 247 | 5,707 | 1,109,106 | 845,040 | 264,066 | 94 | 95.1 | 90.5 | 70 | 30 | 40 | 975 | 673 | 302 | 0.11 | 0.10 | 0.13 | 0.72 | 50.67 | 48.59 | 55.30 | 0.88 |
| I | Descurainieae | *Tropidocarpum gracile* | S0762-2 | 999 | 993 | 288 | 254 | 5,938 | 1,086,567 | 838,200 | 248,367 | 92.1 | 94.4 | 85.1 | 48 | 25 | 23 | 1019 | 698 | 321 | 2.00 | 2.06 | 1.79 | 1.15 | 98.43 | 98.57 | 98.13 | 1.00 |
| I | Erysimeae | *Erysimum cheiranthoides* | S0055 | 997 | 994 | 286 | 254 | 5,856 | 1,129,920 | 857,718 | 272,202 | 95.7 | 96.5 | 93.2 | 65 | 27 | 38 | 996 | 690 | 306 | 0.22 | 0.19 | 0.29 | 0.66 | 44.78 | 39.42 | 56.86 | 0.69 |
| I | Halimolobeae | *Exhalimolobos palmeri* | S1279 | 970 | 967 | 275 | 273 | 5,536 | 1,080,300 | 823,989 | 256,311 | 91.5 | 92.8 | 87.8 | 40 | 24 | 16 | 992 | 675 | 317 | 1.86 | 1.95 | 1.57 | 1.24 | 98.29 | 98.37 | 98.11 | 1.00 |
| I | Halimolobeae | *Halimolobos lasiolobus* | S1370 | 1004 | 1004 | 295 | 313 | 5,882 | 1,117,704 | 845,277 | 272,427 | 94.7 | 95.1 | 93.3 | 65 | 35 | 30 | 1007 | 690 | 317 | 1.11 | 1.09 | 1.17 | 0.94 | 76.76 | 73.77 | 83.28 | 0.89 |
| I | Halimolobeae | *Mancoa hispida* | S0405 | 983 | 982 | 282 | 514 | 5,584 | 1,124,853 | 855,174 | 269,679 | 95.3 | 96.3 | 92.4 | 87 | 38 | 49 | 958 | 664 | 294 | 0.30 | 0.29 | 0.30 | 0.97 | 80.69 | 80.87 | 80.27 | 1.01 |
| I | Halimolobeae | *Pennellia longifolia* | S0643 | 950 | 946 | 274 | 513 | 5,424 | 1,007,223 | 759,516 | 247,707 | 85.3 | 85.5 | 84.9 | 97 | 53 | 44 | 918 | 621 | 297 | 0.29 | 0.29 | 0.28 | 1.03 | 68.95 | 67.63 | 71.72 | 0.94 |
| I | Halimolobeae | *Sphaerocardamum nesliiforme* | S1278 | 990 | 988 | 286 | 284 | 5,611 | 1,121,169 | 851,022 | 270,147 | 95 | 95.8 | 92.5 | 10 | 5 | 5 | 1044 | 708 | 336 | 0.14 | 0.11 | 0.24 | 0.45 | 10.73 | 8.19 | 16.07 | 0.51 |
| I | Hemilophieae trib. nov. | *Dipoma iberideum* | S1371 | 948 | 943 | 266 | 607 | 5,722 | 1,018,074 | 773,601 | 244,473 | 86.3 | 87.1 | 83.7 | 10 | 7 | 3 | 1006 | 675 | 331 | 6.33 | 6.54 | 5.66 | 1.16 | 97.81 | 97.63 | 98.19 | 0.99 |
| I | Hemilophieae trib. nov. | *Hemilophia rockii* | SRR8528376 | 709 | 712 | 195 | 220 | 4,757 | 889,383 | 851,007 | 38,376 | 75.4 | 95.8 | 13.1 | 0 | 0 | 0 | 757 | 711 | 46 | 2.91 | 2.86 | 3.90 | 0.74 | 90.89 | 91.98 | 73.91 | 1.24 |
| I | Lepidieae | *Cyphocardamum aretioides* | S1181-2 | 970 | 956 | 265 | 588 | 5,339 | 992,763 | 753,561 | 239,202 | 84.1 | 84.8 | 81.9 | 24 | 20 | 4 | 1023 | 684 | 339 | 4.13 | 4.17 | 3.99 | 1.05 | 97.65 | 97.66 | 97.64 | 1.00 |
| I | Lepidieae | *Delpinophytum patagonicum* | S0751 | 905 | 903 | 255 | 225 | 6,300 | 1,025,874 | 765,984 | 259,890 | 86.9 | 86.2 | 89 | 1 | 0 | 0 | 981 | 642 | 339 | 7.43 | 7.58 | 6.98 | 1.09 | 97.35 | 97.04 | 97.94 | 0.99 |
| I | Lepidieae | *Lepidium arbuscula* | S1561 | 974 | 961 | 270 | 657 | 5,393 | 1,024,143 | 771,651 | 252,492 | 86.8 | 86.9 | 86.5 | 10 | 7 | 3 | 1040 | 700 | 340 | 4.90 | 4.96 | 4.72 | 1.05 | 95.48 | 95.00 | 96.47 | 0.98 |
| I | Lepidieae | *Lepidium latifolium* | S0591 | 906 | 903 | 246 | 481 | 4,919 | 1,073,634 | 811,779 | 261,855 | 91 | 91.4 | 89.7 | 70 | 36 | 34 | 896 | 600 | 296 | 1.73 | 1.77 | 1.58 | 1.12 | 97.54 | 97.50 | 97.64 | 1.00 |
| I | Lepidieae | *Lepidium virginicum* | S0547 | 924 | 912 | 255 | 481 | 5,319 | 1,024,086 | 767,853 | 256,233 | 86.8 | 86.4 | 87.8 | 28 | 17 | 11 | 971 | 643 | 328 | 4.09 | 4.19 | 3.79 | 1.10 | 96.19 | 95.33 | 97.87 | 0.97 |
| I | Malcolmieae | *Malcolmia maritima* | S1375 | 1013 | 1010 | 294 | 656 | 5,958 | 1,125,492 | 854,925 | 270,567 | 95.4 | 96.2 | 92.7 | 57 | 35 | 22 | 1017 | 691 | 326 | 1.00 | 0.94 | 1.17 | 0.81 | 71.58 | 67.73 | 79.75 | 0.85 |
| I | Microlepidieae | *Arabidella trisecta* | S0509 | 978 | 975 | 280 | 512 | 6,257 | 1,088,763 | 827,238 | 261,525 | 92.2 | 93.1 | 89.6 | 0 | 0 | 0 | 1044 | 709 | 335 | 4.59 | 4.51 | 4.81 | 0.94 | 97.70 | 97.60 | 97.91 | 1.00 |
| I | Microlepidieae | *Ballantinia antipoda* | ERR2789774 | 328 | 319 | 103 | 324 | 1,235 | - | - | - | - | - | - | - | - | - | - | - | - | - | - | - | - | - | - | - | - |
| I | Microlepidieae | *Blennodia canescens* | S0306 | 475 | 476 | 127 | 104 | 3,217 | 680,112 | 662,823 | 17,289 | 57.6 | 74.6 | 5.9 | 1 | 0 | 0 | 503 | 482 | 21 | 3.43 | 3.43 | 3.71 | 0.92 | 92.25 | 91.91 | 100 | 0.92 |
| I | Microlepidieae | *Carinavalva glauca* | S0285 | 837 | 827 | 223 | 210 | 4,855 | 966,723 | 734,868 | 231,855 | 81.9 | 82.7 | 79.4 | 1 | 0 | 0 | 890 | 582 | 308 | 3.35 | 3.29 | 3.54 | 0.93 | 80.34 | 80.07 | 80.84 | 0.99 |
| I | Microlepidieae | *Cuphonotus humistratus* | S1116 | 967 | 963 | 276 | 890 | 5,927 | 1,077,405 | 811,341 | 266,064 | 91.3 | 91.3 | 91.1 | 0 | 0 | 0 | 1031 | 689 | 342 | 3.59 | 3.68 | 3.30 | 1.12 | 81.57 | 81.28 | 82.16 | 0.99 |
| I | Microlepidieae | *Drabastrum alpestre* | S0728sl | 890 | 865 | 243 | 57 | 5,098 | 773,553 | 599,889 | 173,664 | 65.5 | 67.5 | 59.5 | 0 | 0 | 0 | 963 | 646 | 317 | 2.95 | 2.88 | 3.22 | 0.89 | 86.60 | 86.53 | 86.75 | 1.00 |
| I | Microlepidieae | *Geococcus pusillus* | SRR8528342 | 705 | 709 | 194 | 488 | 4,888 | 890,859 | 852,720 | 38,139 | 75.5 | 96 | 13.1 | 0 | 0 | 0 | 755 | 715 | 40 | 3.14 | 3.15 | 3.00 | 1.05 | 56.29 | 56.50 | 52.50 | 1.08 |
| I | Microlepidieae | *Harmsiodoxa blennodioides* | S0896 | 905 | 900 | 263 | 433 | 5,345 | 986,319 | 757,929 | 228,390 | 83.6 | 85.3 | 78.2 | 1 | 0 | 0 | 960 | 669 | 291 | 4.80 | 4.81 | 4.76 | 1.01 | 99.38 | 99.55 | 98.97 | 1.01 |
| I | Microlepidieae | *Irenepharsus phasmatodes* | S0947sl | 971 | 960 | 280 | 796 | 6,444 | 1,029,801 | 769,899 | 259,902 | 87.2 | 86.7 | 89 | 1 | 0 | 0 | 1049 | 704 | 345 | 5.10 | 5.29 | 4.52 | 1.17 | 89.42 | 88.49 | 91.30 | 0.97 |
| I | Microlepidieae | *Menkea australis* | S0941 | 870 | 857 | 243 | 416 | 5,093 | 949,509 | 741,312 | 208,197 | 80.4 | 83.4 | 71.3 | 6 | 3 | 3 | 928 | 616 | 312 | 3.45 | 3.40 | 3.61 | 0.94 | 96.55 | 96.59 | 96.47 | 1.00 |
| I | Microlepidieae | *Microlepidium pilosulum* | S0741 | 957 | 952 | 277 | 243 | 5,862 | 1,074,492 | 808,398 | 266,094 | 91 | 91 | 91.1 | 0 | 0 | 0 | 1022 | 683 | 339 | 2.62 | 2.62 | 2.61 | 1.00 | 63.21 | 62.23 | 65.19 | 0.95 |
| I | Microlepidieae | *Pachycladon fastigiata* | S1400 | 954 | 949 | 268 | 142 | 6,708 | 1,035,669 | 775,929 | 259,740 | 87.7 | 87.3 | 89 | 15 | 12 | 3 | 1020 | 680 | 340 | 5.76 | 5.89 | 5.35 | 1.10 | 95.98 | 96.32 | 95.29 | 1.01 |
| I | Microlepidieae | *Pachycladon novae-zelandiae* | S0435 | 958 | 954 | 275 | 501 | 6,916 | 1,010,373 | 751,461 | 258,912 | 85.6 | 84.6 | 88.7 | 15 | 8 | 7 | 1022 | 688 | 334 | 5.78 | 5.96 | 5.26 | 1.13 | 96.87 | 97.38 | 95.81 | 1.02 |
| I | Microlepidieae | *Pachymitus cardaminoides* | S0949 | 975 | 972 | 286 | 803 | 5,994 | 1,093,686 | 827,964 | 265,722 | 92.7 | 93.2 | 91 | 0 | 0 | 0 | 1038 | 693 | 345 | 4.16 | 4.22 | 3.96 | 1.07 | 93.83 | 93.80 | 93.91 | 1.00 |
| I | Microlepidieae | *Phlegmatospermum cochlearinum* | S0895 | 945 | 940 | 267 | 461 | 5,727 | 1,055,946 | 805,947 | 249,999 | 89.5 | 90.7 | 85.6 | 0 | 0 | 0 | 1019 | 686 | 333 | 4.20 | 4.27 | 3.97 | 1.08 | 96.27 | 95.77 | 97.30 | 0.98 |
| I | Microlepidieae | *Scambopus curvipes* | S0946 | 890 | 887 | 246 | 740 | 5,347 | 1,030,875 | 774,975 | 255,900 | 87.3 | 87.2 | 87.7 | 2 | 2 | 0 | 947 | 626 | 321 | 4.14 | 4.15 | 4.12 | 1.01 | 95.35 | 94.41 | 97.20 | 0.97 |
| I | Microlepidieae | *Stenopetalum lineare* | S1424 | 987 | 985 | 287 | 876 | 6,287 | 1,092,081 | 821,184 | 270,897 | 92.5 | 92.4 | 92.8 | 0 | 0 | 0 | 1057 | 713 | 344 | 3.34 | 3.37 | 3.26 | 1.03 | 67.27 | 62.97 | 76.16 | 0.83 |
| I | Oreophytoneae | *Murbeckiella pinnatifida* | SRR8528350 | 719 | 724 | 201 | 504 | 4,636 | 919,314 | 881,292 | 38,022 | 77.9 | 99.2 | 13 | 31 | 29 | 2 | 739 | 698 | 41 | 0.04 | 0.03 | 0.09 | 0.39 | 21.11 | 19.77 | 43.90 | 0.45 |
| I | Oreophytoneae | *Oreophyton falcatum* | S0612 | 974 | 968 | 278 | 527 | 5,766 | 1,078,518 | 825,561 | 252,957 | 91.4 | 92.9 | 86.6 | 36 | 15 | 21 | 1003 | 686 | 317 | 2.10 | 2.14 | 1.97 | 1.08 | 98.40 | 98.54 | 98.11 | 1.00 |
| I | Physarieae | *Dimorphocarpa wislizenii* | S0294 | 926 | 920 | 259 | 236 | 5,613 | 1,088,445 | 820,920 | 267,525 | 92.2 | 92.4 | 91.6 | 29 | 22 | 7 | 953 | 626 | 327 | 2.07 | 2.16 | 1.81 | 1.19 | 84.78 | 85.30 | 83.79 | 1.02 |
| I | Physarieae | *Dithyrea californica* | S0903 | 973 | 964 | 276 | 462 | 5,802 | 1,101,267 | 836,262 | 265,005 | 93.3 | 94.1 | 90.8 | 195 | 145 | 50 | 841 | 553 | 288 | 0.79 | 0.75 | 0.89 | 0.84 | 94.29 | 92.95 | 96.88 | 0.96 |
| I | Physarieae | *Lyrocarpa coulteri* | S0300 | 908 | 910 | 239 | 221 | 5,507 | 1,062,858 | 824,583 | 238,275 | 90 | 92.8 | 81.6 | 31 | 26 | 5 | 942 | 652 | 290 | 3.58 | 3.67 | 3.29 | 1.11 | 97.88 | 98.47 | 96.55 | 1.02 |
| I | Physarieae | *Nerisyrenia linearifolia* | S0764 | 982 | 974 | 280 | 173 | 5,882 | 1,107,282 | 839,373 | 267,909 | 93.8 | 94.5 | 91.8 | 93 | 77 | 16 | 959 | 629 | 330 | 1.47 | 1.30 | 2.02 | 0.64 | 86.03 | 82.99 | 91.82 | 0.90 |
| I | Physarieae | *Paysonia lescurii* | S1052sl | 922 | 882 | 248 | 134 | 5,131 | 818,094 | 613,842 | 204,252 | 69.3 | 69.1 | 70 | 45 | 38 | 7 | 950 | 623 | 327 | 1.10 | 1.07 | 1.20 | 0.89 | 71.68 | 71.91 | 71.25 | 1.01 |
| I | Physarieae | *Physaria didymocarpa* | S0383 | 838 | 818 | 221 | 426 | 4,987 | 979,611 | 774,678 | 204,933 | 83 | 87.2 | 70.2 | 19 | 16 | 3 | 876 | 606 | 270 | 3.17 | 3.26 | 2.85 | 1.14 | 91.10 | 93.07 | 86.67 | 1.07 |
| I | Physarieae | *Synthlipsis greggii* | S0275 | 957 | 940 | 270 | 243 | 5,659 | 1,077,342 | 818,361 | 258,981 | 91.3 | 92.1 | 88.7 | 32 | 26 | 6 | 989 | 658 | 331 | 3.69 | 3.70 | 3.66 | 1.01 | 96.76 | 96.35 | 97.58 | 0.99 |
| I | Smelowskieae | *Smelowskia alba* | S0689 | 934 | 938 | 263 | 71 | 5,478 | 1,084,656 | 864,681 | 219,975 | 91.9 | 97.3 | 75.4 | 71 | 43 | 28 | 924 | 681 | 243 | 0.70 | 0.68 | 0.78 | 0.87 | 82.47 | 79.30 | 91.36 | 0.87 |
| I | Turritideae | *Turritis glabra* | SRR8528357 | 976 | 973 | 284 | 677 | 5,783 | 1,093,320 | 826,911 | 266,409 | 92.6 | 93.1 | 91.3 | 23 | 9 | 14 | 1013 | 684 | 329 | 0.03 | 0.03 | 0.02 | 2.07 | 10.07 | 12.72 | 4.56 | 2.79 |
| I | Yinshanieae | *Yinshania acutangula subsp. acutangula* | SRR8528359 | 719 | 727 | 201 | 434 | 4,598 | 916,848 | 881,772 | 35,076 | 77.7 | 99.3 | 12 | 30 | 25 | 5 | 741 | 703 | 38 | 0.05 | 0.05 | 0.13 | 0.38 | 23.62 | 22.62 | 42.11 | 0.54 |
| II | Aphragmeae | *Aphragmus obscurus* | S0283 | 927 | 917 | 263 | 234 | 5,716 | 988,761 | 776,196 | 212,565 | 83.8 | 87.4 | 72.8 | 42 | 19 | 23 | 953 | 647 | 306 | 1.92 | 1.91 | 1.95 | 0.98 | 96.64 | 97.37 | 95.10 | 1.02 |
| II | Brassiceae | *Ammosperma cinerea* | S0305 | 725 | 715 | 181 | 154 | 4,039 | 832,335 | 777,810 | 54,525 | 70.5 | 87.6 | 18.7 | 0 | 0 | 0 | 787 | 661 | 126 | 4.48 | 4.55 | 3.45 | 1.32 | 94.03 | 97.88 | 73.81 | 1.33 |
| II | Brassiceae | *Brassica oleracea - nominate* | S1208sl | 780 | 739 | 205 | 376 | 3,931 | 756,906 | 555,771 | 201,135 | 64.1 | 62.6 | 68.9 | 5 | 4 | 1 | 836 | 536 | 300 | 2.71 | 2.62 | 2.98 | 0.88 | 81.22 | 79.85 | 83.67 | 0.95 |
| II | Brassiceae | *Brassica villosa* | S1220 | 791 | 759 | 202 | 374 | 3,970 | 780,819 | 602,976 | 177,843 | 66.2 | 67.9 | 60.9 | 2 | 2 | 0 | 864 | 573 | 291 | 2.84 | 2.83 | 2.87 | 0.99 | 77.31 | 77.14 | 77.66 | 0.99 |
| II | Brassiceae | *Cakile maritima* | S0743 | 912 | 908 | 254 | 227 | 4,692 | 1,038,927 | 780,153 | 258,774 | 88 | 87.8 | 88.6 | 0 | 0 | 0 | 982 | 648 | 334 | 2.93 | 3.12 | 2.36 | 1.32 | 72.20 | 72.07 | 72.46 | 0.99 |
| II | Brassiceae | *Carrichtera annua* | S0768 | 816 | 806 | 218 | 135 | 4,353 | 953,457 | 719,157 | 234,300 | 80.8 | 81 | 80.3 | 0 | 0 | 0 | 875 | 567 | 308 | 2.63 | 2.77 | 2.22 | 1.25 | 62.63 | 64.02 | 60.06 | 1.07 |
| II | Brassiceae | *Ceratocnemum rapistroides* | S0660sl | 754 | 717 | 191 | 405 | 3,749 | 593,523 | 442,281 | 151,242 | 50.3 | 49.8 | 51.8 | 14 | 7 | 7 | 810 | 532 | 278 | 2.68 | 2.53 | 3.10 | 0.82 | 88.40 | 87.41 | 90.29 | 0.97 |
| II | Brassiceae | *Coincya monensis* | S1401 | 981 | 970 | 280 | 139 | 5,128 | 1,072,311 | 809,181 | 263,130 | 90.8 | 91.1 | 90.1 | 1 | 0 | 0 | 1047 | 704 | 343 | 5.01 | 5.22 | 4.35 | 1.20 | 96.94 | 96.45 | 97.96 | 0.98 |
| II | Brassiceae | *Crambe maritima* | S0731sl | 925 | 922 | 254 | 68 | 5,058 | 1,062,081 | 802,923 | 259,158 | 90 | 90.4 | 88.8 | 0 | 0 | 0 | 990 | 658 | 332 | 4.11 | 4.39 | 3.23 | 1.36 | 88.69 | 88.60 | 88.86 | 1.00 |
| II | Brassiceae | *Crambella teretifolia* | S1021 | 977 | 969 | 279 | 144 | 5,171 | 1,067,931 | 806,271 | 261,660 | 90.5 | 90.8 | 89.6 | 1 | 0 | 0 | 1046 | 704 | 342 | 6.09 | 6.29 | 5.50 | 1.14 | 97.61 | 97.59 | 97.66 | 1.00 |
| II | Brassiceae | *Didesmus aegyptius* | S0315 | 981 | 971 | 281 | 248 | 5,046 | 1,064,427 | 803,379 | 261,048 | 90.2 | 90.4 | 89.4 | 2 | 2 | 0 | 1046 | 703 | 343 | 4.70 | 4.91 | 4.04 | 1.22 | 90.54 | 91.04 | 89.50 | 1.02 |
| II | Brassiceae | *Diplotaxis harra* | S0851 | 945 | 935 | 266 | 158 | 4,998 | 1,020,387 | 764,232 | 256,155 | 86.4 | 86 | 87.7 | 0 | 0 | 0 | 1023 | 682 | 341 | 4.55 | 4.65 | 4.26 | 1.09 | 91.10 | 91.06 | 91.20 | 1.00 |
| II | Brassiceae | *Douepea tortuosa* | S0954 | 943 | 927 | 265 | 769 | 5,051 | 1,044,741 | 791,304 | 253,437 | 88.5 | 89.1 | 86.8 | 0 | 0 | 0 | 1011 | 677 | 334 | 3.83 | 3.95 | 3.45 | 1.14 | 89.71 | 89.36 | 90.42 | 0.99 |
| II | Brassiceae | *Enarthrocarpus arcuatus* | S0662 | 910 | 890 | 255 | 59 | 4,824 | 1,034,346 | 773,616 | 260,730 | 87.6 | 87.1 | 89.3 | 0 | 0 | 0 | 976 | 645 | 331 | 3.69 | 3.95 | 2.95 | 1.34 | 79.71 | 80.93 | 77.34 | 1.05 |
| II | Brassiceae | *Eremophyton chevallieri* | S0726 | 852 | 837 | 227 | 60 | 4,440 | 965,637 | 739,506 | 226,131 | 81.8 | 83.2 | 77.5 | 0 | 0 | 0 | 917 | 609 | 308 | 4.82 | 4.94 | 4.40 | 1.12 | 93.46 | 93.92 | 92.53 | 1.02 |
| II | Brassiceae | *Eruca vesicaria* | S1408 | 917 | 894 | 252 | 744 | 4,759 | 1,001,691 | 750,285 | 251,406 | 84.9 | 84.5 | 86.1 | 0 | 0 | 0 | 990 | 653 | 337 | 4.73 | 4.86 | 4.32 | 1.13 | 95.05 | 95.10 | 94.96 | 1.00 |
| II | Brassiceae | *Erucaria hispanica* | S0664 | 940 | 931 | 260 | 64 | 4,933 | 1,062,378 | 801,558 | 260,820 | 90 | 90.2 | 89.3 | 0 | 0 | 0 | 1010 | 670 | 340 | 5.51 | 5.74 | 4.79 | 1.20 | 98.91 | 98.66 | 99.41 | 0.99 |
| II | Brassiceae | *Erucastrum gallicum* | S0317 | 955 | 931 | 264 | 239 | 5,133 | 1,052,646 | 789,525 | 263,121 | 89.2 | 88.9 | 90.1 | 0 | 0 | 0 | 1024 | 682 | 342 | 4.91 | 5.15 | 4.18 | 1.23 | 95.90 | 95.89 | 95.91 | 1.00 |
| II | Brassiceae | *Foleyola billotii* | S0772 | 925 | 913 | 252 | 162 | 4,834 | 1,034,973 | 786,666 | 248,307 | 87.7 | 88.5 | 85.1 | 3 | 3 | 0 | 990 | 663 | 327 | 5.15 | 5.30 | 4.67 | 1.14 | 98.59 | 98.34 | 99.08 | 0.99 |
| II | Brassiceae | *Fortuynia garcinii* | S0464 | 919 | 912 | 254 | 484 | 5,002 | 1,039,605 | 783,987 | 255,618 | 88.1 | 88.2 | 87.6 | 0 | 0 | 0 | 988 | 659 | 329 | 4.44 | 4.65 | 3.78 | 1.23 | 92.00 | 92.56 | 90.88 | 1.02 |
| II | Brassiceae | *Guenthera souliei* | S1221sl | 959 | 917 | 267 | 449 | 4,988 | 996,063 | 753,810 | 242,253 | 84.4 | 84.9 | 83 | 14 | 11 | 3 | 1020 | 686 | 334 | 3.83 | 3.90 | 3.63 | 1.07 | 96.27 | 96.21 | 96.41 | 1.00 |
| II | Brassiceae | *Guiraoa arvensis* | S0323 | 930 | 921 | 262 | 232 | 4,924 | 1,033,359 | 774,672 | 258,687 | 87.5 | 87.2 | 88.6 | 0 | 0 | 0 | 995 | 661 | 334 | 4.83 | 5.09 | 4.05 | 1.26 | 94.87 | 95.76 | 93.11 | 1.03 |
| II | Brassiceae | *Hemicrambe fruticulosa* | S0617 | 735 | 726 | 184 | 398 | 4,224 | 823,833 | 752,406 | 71,427 | 69.8 | 84.7 | 24.5 | 0 | 0 | 0 | 807 | 659 | 148 | 9.44 | 9.78 | 5.87 | 1.67 | 96.16 | 98.33 | 86.49 | 1.14 |
| II | Brassiceae | *Henophyton deserti* | S1442 | 887 | 880 | 244 | 580 | 4,740 | 963,390 | 731,316 | 232,074 | 81.6 | 82.3 | 79.5 | 4 | 4 | 0 | 952 | 629 | 323 | 3.11 | 3.13 | 3.04 | 1.03 | 88.34 | 87.92 | 89.16 | 0.99 |
| II | Brassiceae | *Hirschfeldia incana* | S0590 | 709 | 684 | 185 | 365 | 3,456 | 796,824 | 581,520 | 215,304 | 67.5 | 65.5 | 73.8 | 2 | 1 | 1 | 760 | 472 | 288 | 4.81 | 4.82 | 4.77 | 1.01 | 91.32 | 90.89 | 92.01 | 0.99 |
| II | Brassiceae | *Horwoodia dicksoniae* | S0297 | 745 | 736 | 193 | 177 | 3,921 | 871,581 | 668,034 | 203,547 | 73.8 | 75.2 | 69.7 | 3 | 3 | 0 | 802 | 525 | 277 | 3.18 | 3.26 | 2.91 | 1.12 | 89.90 | 91.81 | 86.28 | 1.06 |
| II | Brassiceae | *Kremeriella cordylocarpus* | S0326-2 | 216 | - | - | - | 977 | 71,709 | 52,350 | 19,359 | 6.1 | 5.9 | 6.6 | 1 | 0 | 0 | 254 | 169 | 85 | 0.51 | 0.55 | 0.41 | 1.33 | 20.47 | 22.49 | 16.47 | 1.37 |
| II | Brassiceae | *Moricandia arvensis* | S1314 | 924 | 919 | 262 | 244 | 4,964 | 1,020,168 | 766,872 | 253,296 | 86.4 | 86.3 | 86.8 | 0 | 0 | 0 | 991 | 654 | 337 | 4.35 | 4.53 | 3.82 | 1.19 | 82.54 | 82.57 | 82.49 | 1.00 |
| II | Brassiceae | *Morisia monanthos* | S0331 | 655 | 642 | 170 | 158 | 3,189 | 775,746 | 588,567 | 187,179 | 65.7 | 66.3 | 64.1 | 2 | 1 | 1 | 711 | 452 | 259 | 4.08 | 4.26 | 3.50 | 1.22 | 93.11 | 94.91 | 89.96 | 1.06 |
| II | Brassiceae | *Nasturtiopsis coronopifolia* | S0385 | 946 | 935 | 268 | 473 | 5,094 | 1,017,549 | 766,230 | 251,319 | 86.2 | 86.2 | 86.1 | 1 | 0 | 0 | 1020 | 681 | 339 | 4.09 | 4.16 | 3.87 | 1.08 | 92.35 | 92.80 | 91.45 | 1.01 |
| II | Brassiceae | *Orychophragmus violaceus* | S1412 | 260 | - | - | - | 1,300 | 99,240 | 70,794 | 28,446 | 8.4 | 8 | 9.7 | 5 | 4 | 1 | 283 | 185 | 98 | 1.43 | 1.26 | 1.84 | 0.68 | 59.01 | 56.76 | 63.27 | 0.90 |
| II | Brassiceae | *Otocarpus virgatus* | S0752 | 956 | 941 | 265 | 237 | 5,250 | 1,060,074 | 793,155 | 266,919 | 89.8 | 89.3 | 91.4 | 0 | 0 | 0 | 1019 | 677 | 342 | 5.29 | 5.41 | 4.93 | 1.10 | 94.21 | 93.94 | 94.74 | 0.99 |

### Table S4. Continued.

| SAMPLE DETAILS | | | | ASTRAL TAXON OCCUPANCY | | | | | RECOVERED TOTAL LENGTH (BP) | | | RECOVERED TOTAL LENGTH OF TARGET LENGTH (%) | | | # PARALOG CALLS | | | # LOCI RECOVERED | | | ALLELE DIVERGENCE | | | | LOCUS HETEROZYGOSITY | | | |
|---|---|---|---|---|---|---|---|---|---|---|---|---|---|---|---|---|---|---|---|---|---|---|---|---|---|---|---|---|
| Main lineage | Tribe | Species | Sample | inclusive | strict | superstrict | superstrict by tribe | A-Pro | Total | B764 | A353 | Total | B764 | A353 | Total | B764 | A353 | Total | B764 | A353 | Total (%) | B764 (%) | A353(%) | B764/A353 | Total (%) | B764 (%) | A353(%) | B764/A353 |
| II | Brassiceae | *Physorhynchus brahuicus* | S1168 | 914 | 909 | 262 | 662 | 4,954 | 1,032,996 | 781,734 | 251,262 | 87.5 | 88 | 86.1 | 0 | 0 | 0 | 979 | 650 | 329 | 4.07 | 4.20 | 3.64 | 1.15 | 87.84 | 87.54 | 88.45 | 0.99 |
| II | Brassiceae | *Pseuderucaria clavata* | S1331 | 921 | 914 | 249 | 801 | 4,885 | 1,010,850 | 772,455 | 238,395 | 85.6 | 86.9 | 81.7 | 4 | 3 | 1 | 986 | 660 | 326 | 3.84 | 4.00 | 3.31 | 1.21 | 92.49 | 92.88 | 91.72 | 1.01 |
| II | Brassiceae | *Psychine stylosa* | S0380 | 953 | 942 | 265 | 488 | 5,142 | 1,047,051 | 786,249 | 260,802 | 88.7 | 88.5 | 89.3 | 0 | 0 | 0 | 1017 | 677 | 340 | 4.31 | 4.59 | 3.48 | 1.32 | 89.38 | 89.96 | 88.24 | 1.02 |
| II | Brassiceae | *Quezeliantha tibestica* | S1458 | 720 | 701 | 175 | 270 | 3,535 | 581,838 | 454,002 | 127,836 | 49.3 | 51.1 | 43.8 | 13 | 6 | 7 | 771 | 518 | 253 | 4.50 | 4.42 | 4.78 | 0.92 | 92.61 | 92.86 | 92.09 | 1.01 |
| II | Brassiceae | *Quidproquo confusum* | S0897 | 778 | 758 | 207 | 368 | 3,803 | 801,945 | 598,761 | 203,184 | 67.9 | 67.4 | 69.6 | 3 | 1 | 2 | 840 | 535 | 305 | 3.55 | 3.60 | 3.40 | 1.06 | 91.79 | 92.52 | 90.49 | 1.02 |
| II | Brassiceae | *Raffenaldia primuloides* | S0667 | 821 | 813 | 222 | 52 | 4,320 | 932,940 | 691,191 | 241,749 | 79 | 77.8 | 82.8 | 1 | 0 | 0 | 878 | 562 | 316 | 4.71 | 4.92 | 4.10 | 1.20 | 95.22 | 94.31 | 96.84 | 0.97 |
| II | Brassiceae | *Raphanus sativus* | S0583 | 906 | 885 | 252 | 482 | 4,864 | 1,010,538 | 754,068 | 256,470 | 85.6 | 84.9 | 87.9 | 1 | 0 | 0 | 967 | 632 | 335 | 4.39 | 4.53 | 3.99 | 1.13 | 86.14 | 86.23 | 85.97 | 1.00 |
| II | Brassiceae | *Rapistrum rugosum* | S1421 | 807 | 795 | 216 | 696 | 4,188 | 885,777 | 656,862 | 228,915 | 75 | 73.9 | 78.4 | 0 | 0 | 0 | 860 | 555 | 305 | 4.63 | 4.65 | 4.57 | 1.02 | 91.86 | 92.07 | 91.48 | 1.01 |
| II | Brassiceae | *Rhamphospermum arvense* | S1509 | 927 | 891 | 260 | 325 | 4,765 | 979,785 | 734,967 | 244,818 | 83 | 82.7 | 83.9 | 1 | 0 | 0 | 1000 | 661 | 339 | 4.10 | 4.14 | 3.95 | 1.05 | 88.20 | 86.99 | 90.56 | 0.96 |
| II | Brassiceae | *Savignya parviflora* | S1339 | 861 | 851 | 227 | 59 | 4,586 | 954,735 | 709,437 | 245,298 | 80.9 | 79.9 | 84 | 0 | 0 | 0 | 924 | 599 | 325 | 2.67 | 2.77 | 2.38 | 1.16 | 57.68 | 57.10 | 58.77 | 0.97 |
| II | Brassiceae | *Schouwia purpurea* | S0669 | 950 | 940 | 268 | 64 | 5,250 | 1,068,459 | 805,122 | 263,337 | 90.5 | 90.6 | 90.2 | 0 | 0 | 0 | 1014 | 678 | 336 | 3.47 | 3.61 | 3.04 | 1.19 | 60.65 | 60.91 | 60.12 | 1.01 |
| II | Brassiceae | *Sinalliaria limprichtiana* | SRR6441723 | 330 | 323 | 107 | 329 | 1,253 | - | - | - | - | - | - | - | - | - | - | - | - | - | - | - | - | - | - | - | - |
| II | Brassiceae | *Sinapidendron frutescens* | S0377 | 953 | 906 | 265 | 477 | 5,011 | 1,009,221 | 760,554 | 248,667 | 85.5 | 85.6 | 85.2 | 0 | 0 | 0 | 1018 | 690 | 328 | 3.80 | 3.86 | 3.60 | 1.07 | 86.94 | 86.09 | 88.72 | 0.97 |
| II | Brassiceae | *Sinapis alba* | S1011sl | 637 | 623 | 163 | 89 | 3,124 | 633,858 | 466,566 | 167,292 | 53.7 | 52.5 | 57.3 | 7 | 5 | 2 | 681 | 428 | 253 | 5.89 | 5.92 | 5.79 | 1.02 | 98.38 | 98.13 | 98.81 | 0.99 |
| II | Brassiceae | *Succowia balearica* | S0373 | 939 | 933 | 266 | 241 | 5,297 | 1,070,856 | 807,516 | 263,340 | 90.7 | 90.9 | 90.2 | 2 | 2 | 0 | 1012 | 676 | 336 | 3.68 | 3.82 | 3.23 | 1.18 | 66.40 | 66.12 | 66.96 | 0.99 |
| II | Brassiceae | *Vella castrilensis* | S1270 | 837 | 827 | 229 | 465 | 4,432 | 945,612 | 720,144 | 225,468 | 80.1 | 81.1 | 77.2 | 3 | 3 | 0 | 897 | 588 | 309 | 4.97 | 5.01 | 4.83 | 1.04 | 98.22 | 99.15 | 96.44 | 1.03 |
| II | Brassiceae | *Vella pseudocytisus* | S0371 | 610 | 608 | 159 | 133 | 3,744 | 804,216 | 776,400 | 27,816 | 68.1 | 87.4 | 9.5 | 1 | 0 | 0 | 652 | 621 | 31 | 4.60 | 4.60 | 4.79 | 0.96 | 95.09 | 95.33 | 90.32 | 1.06 |
| II | Brassiceae | *Zahora ait-atta* | S0747sl | 978 | 963 | 274 | 246 | 5,307 | 1,097,010 | 824,037 | 272,973 | 92.9 | 92.8 | 93.5 | 0 | 0 | 0 | 1046 | 702 | 344 | 4.27 | 4.54 | 3.44 | 1.32 | 84.51 | 83.62 | 86.34 | 0.97 |
| II | Brassiceae | *Zilla spinosa* | S1506 | 735 | 724 | 184 | 234 | 3,742 | 689,271 | 538,146 | 151,125 | 58.4 | 60.6 | 51.8 | 4 | 3 | 1 | 784 | 521 | 263 | 2.86 | 2.76 | 3.22 | 0.86 | 86.99 | 87.91 | 85.17 | 1.03 |
| II | Brassiceae II | *Bivonaea lutea* | SRR8528386 | 517 | - | - | - | 3,139 | 411,882 | 395,148 | 16,734 | 34.9 | 44.5 | 5.7 | 2 | 2 | 0 | 578 | 546 | 32 | 1.49 | 1.46 | 2.24 | 0.65 | 63.84 | 63.92 | 62.50 | 1.02 |
| II | Calepineae | *Calepina irregularis* | S0312 | 999 | 997 | 289 | 256 | 6,056 | 1,064,712 | 810,483 | 254,229 | 90.2 | 91.2 | 87.1 | 37 | 19 | 18 | 1029 | 702 | 327 | 2.31 | 2.34 | 2.20 | 1.07 | 98.64 | 98.58 | 98.78 | 1.00 |
| II | Calepineae | *Goldbachia laevigata* | S0757 | 974 | 960 | 276 | 246 | 6,017 | 1,038,822 | 790,293 | 248,529 | 88 | 89 | 85.1 | 49 | 28 | 21 | 993 | 671 | 322 | 3.17 | 3.28 | 2.82 | 1.16 | 98.79 | 98.96 | 98.45 | 1.01 |
| II | Calepineae | *Leiocarpaea cochlearioides* | S1238sl | 872 | 868 | 238 | 764 | 5,437 | 1,013,028 | 781,899 | 231,129 | 85.8 | 88 | 79.2 | 53 | 27 | 26 | 871 | 587 | 284 | 0.18 | 0.17 | 0.19 | 0.91 | 50.63 | 49.57 | 52.82 | 0.94 |
| II | Coluteocarpeae | *Callothlaspi lilacinum* | S0758 | 693 | 695 | 200 | 176 | 4,357 | 816,069 | 605,637 | 210,432 | 69.1 | 68.2 | 72.1 | 38 | 13 | 25 | 703 | 453 | 250 | 0.63 | 0.59 | 0.75 | 0.79 | 70.70 | 67.55 | 76.40 | 0.88 |
| II | Coluteocarpeae | *Eunomia oppositifolia* | S1128 | 949 | 942 | 270 | 696 | 5,921 | 1,081,626 | 828,186 | 253,440 | 91.6 | 93.2 | 86.8 | 68 | 36 | 32 | 945 | 643 | 302 | 0.72 | 0.71 | 0.76 | 0.94 | 81.38 | 81.34 | 81.46 | 1.00 |
| II | Coluteocarpeae | *Friedrichkarlmeyeria umbellata* | S1189 | 986 | 982 | 281 | 594 | 6,081 | 1,116,546 | 849,777 | 266,769 | 94.6 | 95.7 | 91.4 | 69 | 41 | 28 | 979 | 665 | 314 | 0.17 | 0.16 | 0.23 | 0.69 | 43.31 | 39.10 | 52.23 | 0.75 |
| II | Coluteocarpeae | *Ihsanalshehbazia granatensis* | S0771 | 902 | 897 | 258 | 154 | 5,614 | 1,075,467 | 810,138 | 265,329 | 91.1 | 91.2 | 90.9 | 34 | 15 | 19 | 925 | 612 | 313 | 0.03 | 0.02 | 0.06 | 0.38 | 16.65 | 13.40 | 23.00 | 0.58 |
| II | Coluteocarpeae | *Kotschyella stenocarpa* | S0677 | 890 | 881 | 256 | 55 | 5,502 | 1,023,936 | 775,884 | 248,052 | 86.8 | 87.3 | 85 | 51 | 32 | 19 | 895 | 588 | 307 | 1.34 | 1.30 | 1.45 | 0.90 | 94.86 | 94.73 | 95.11 | 1.00 |
| II | Coluteocarpeae | *Masmenia rosularis* | S0964 | 831 | 829 | 237 | 709 | 5,221 | 976,938 | 741,102 | 235,836 | 82.8 | 83.4 | 80.8 | 66 | 32 | 34 | 824 | 543 | 281 | 0.32 | 0.30 | 0.37 | 0.83 | 70.75 | 68.88 | 74.38 | 0.93 |
| II | Coluteocarpeae | *Microthlaspi perfoliatum* | S0562 | 979 | 972 | 282 | 525 | 6,039 | 1,038,480 | 788,790 | 249,690 | 88 | 88.8 | 85.5 | 12 | 8 | 4 | 1040 | 698 | 342 | 4.71 | 4.69 | 4.77 | 0.98 | 98.75 | 98.57 | 99.12 | 0.99 |
| II | Coluteocarpeae | *Neurotropis orbiculata* | S0959 | 935 | 935 | 274 | 765 | 5,903 | 1,058,631 | 818,004 | 240,627 | 89.7 | 92.1 | 82.4 | 57 | 25 | 32 | 943 | 674 | 269 | 0.10 | 0.07 | 0.21 | 0.32 | 29.48 | 20.62 | 51.67 | 0.40 |
| II | Coluteocarpeae | *Noccaea rotundifolia* | S0552sl | 987 | 985 | 286 | 530 | 6,076 | 1,112,832 | 845,691 | 267,141 | 94.3 | 95.2 | 91.5 | 113 | 62 | 51 | 936 | 643 | 293 | 0.39 | 0.38 | 0.42 | 0.92 | 76.18 | 76.21 | 76.11 | 1.00 |
| II | Coluteocarpeae | *Noccidium hastulatum* | S0613sl | 959 | 953 | 273 | 507 | 5,943 | 1,071,222 | 823,002 | 248,220 | 90.8 | 92.6 | 85 | 43 | 19 | 24 | 978 | 662 | 316 | 1.17 | 1.13 | 1.31 | 0.86 | 90.29 | 89.58 | 91.77 | 0.98 |
| II | Coluteocarpeae | *Pseudosempervivum aucheri* | S0960 | 965 | 958 | 278 | 800 | 5,989 | 1,066,029 | 821,835 | 244,194 | 90.3 | 92.5 | 83.6 | 76 | 38 | 38 | 955 | 651 | 304 | 0.43 | 0.42 | 0.47 | 0.90 | 79.69 | 79.72 | 79.61 | 1.00 |
| II | Coluteocarpeae | *Thlaspiceras rechingeri* | S0956 | 828 | 829 | 228 | 707 | 5,133 | 982,062 | 747,849 | 234,213 | 83.2 | 84.2 | 80.2 | 79 | 38 | 41 | 806 | 533 | 273 | 0.43 | 0.41 | 0.48 | 0.85 | 78.29 | 77.11 | 80.59 | 0.96 |
| II | Coluteocarpeae | *Vania campylophylla* | S1129sl | 1005 | 993 | 289 | 733 | 6,232 | 1,121,439 | 851,748 | 269,691 | 95 | 95.9 | 92.4 | 55 | 29 | 26 | 1013 | 692 | 321 | 0.86 | 0.81 | 1.03 | 0.78 | 79.76 | 78.47 | 82.55 | 0.95 |
| II | Conringieae | *Conringia orientalis* | SRR8528372 | 716 | 721 | 199 | 217 | 4,953 | 915,384 | 880,677 | 34,707 | 77.6 | 99.1 | 11.9 | 30 | 24 | 6 | 736 | 701 | 35 | 0.03 | 0.03 | 0.10 | 0.31 | 19.84 | 18.26 | 51.43 | 0.36 |
| II | Eutremeae | *Eutrema edwardsii* | S0416 | 980 | 978 | 283 | 513 | 6,043 | 1,107,846 | 844,512 | 263,334 | 93.9 | 95.1 | 90.2 | 102 | 52 | 50 | 939 | 646 | 293 | 0.87 | 0.89 | 0.81 | 1.10 | 92.23 | 92.11 | 92.49 | 1.00 |
| II | Eutremeae | *Eutrema edwardsii* | S1336 | 941 | 934 | 265 | 837 | 5,958 | 1,037,508 | 805,707 | 231,801 | 87.9 | 90.7 | 79.4 | 26 | 12 | 14 | 981 | 663 | 318 | 2.25 | 2.22 | 2.34 | 0.95 | 93.58 | 93.36 | 94.03 | 0.99 |
| II | Fourraeeae | *Fourraea alpina* | SRR8528341 | 722 | 728 | 199 | 503 | 4,165 | 923,916 | 885,300 | 38,616 | 78.3 | 99.7 | 13.2 | 32 | 29 | 3 | 740 | 699 | 41 | 0.08 | 0.08 | 0.14 | 0.56 | 25.81 | 24.75 | 43.90 | 0.56 |
| II | Fourraeeae | *Hurkaea josiae* | SRR8528389 | 720 | 726 | 198 | 218 | 4,157 | 912,003 | 874,284 | 37,719 | 77.3 | 98.4 | 12.9 | 31 | 27 | 4 | 738 | 698 | 40 | 0.05 | 0.04 | 0.08 | 0.55 | 22.36 | 21.20 | 42.50 | 0.50 |
| II | Isatideae | *Chartoloma platycarpum* | S0629-2 | 199 | - | - | - | 1,025 | 62,733 | 44,019 | 18,714 | 5.3 | 5 | 6.4 | 2 | 0 | 2 | 224 | 148 | 76 | 0.20 | 0.19 | 0.22 | 0.87 | 17.41 | 16.22 | 19.74 | 0.82 |
| II | Isatideae | *Glastaria glastifolia* | S0280 | 948 | 945 | 272 | 242 | 5,334 | 1,093,593 | 834,672 | 258,921 | 92.7 | 94 | 88.7 | 53 | 27 | 26 | 954 | 646 | 308 | 0.82 | 0.77 | 0.98 | 0.79 | 87.00 | 85.60 | 89.94 | 0.95 |
| II | Isatideae | *Iljinskaea planisiliqua* | SRR8528365 | 721 | 725 | 198 | 218 | 4,456 | 914,925 | 878,841 | 36,084 | 77.5 | 98.9 | 12.4 | 25 | 23 | 2 | 744 | 704 | 40 | 0.03 | 0.03 | 0.11 | 0.27 | 17.61 | 16.19 | 42.50 | 0.38 |
| II | Isatideae | *Isatis tinctoria* | SRR8528380 | 715 | 722 | 198 | 217 | 4,430 | 913,977 | 878,943 | 35,034 | 77.4 | 98.9 | 12 | 18 | 17 | 1 | 745 | 709 | 36 | 1.67 | 1.68 | 1.19 | 1.41 | 96.11 | 96.47 | 88.89 | 1.09 |
| II | Isatideae | *Myagrum perfoliatum* | S0335 | 927 | 923 | 269 | 243 | 5,200 | 1,092,819 | 829,722 | 263,097 | 92.6 | 93.4 | 90.1 | 50 | 19 | 31 | 939 | 632 | 307 | 0.13 | 0.12 | 0.19 | 0.61 | 29.61 | 25.63 | 37.79 | 0.68 |
| II | Isatideae | *Schimpera arabica* | ERR4210213 | 263 | 261 | 79 | 265 | 997 | - | - | - | - | - | - | - | - | - | - | - | - | - | - | - | - | - | - | - | - |
| II | Kernereae | *Kernera saxatilis* | SRR8528379 | 717 | 723 | 202 | 219 | 4,881 | 920,193 | 882,501 | 37,692 | 78 | 99.3 | 12.9 | 33 | 30 | 3 | 735 | 693 | 42 | 0.04 | 0.04 | 0.10 | 0.36 | 22.18 | 21.07 | 40.48 | 0.52 |
| II | Kernereae | *Petrocallis pyrenaica* | SRR8528400 | 719 | 727 | 199 | 220 | 4,903 | 919,041 | 881,352 | 37,689 | 77.9 | 99.2 | 12.9 | 42 | 37 | 5 | 727 | 688 | 39 | 0.18 | 0.18 | 0.21 | 0.84 | 61.07 | 61.05 | 61.54 | 0.99 |
| II | Kernereae | *Rhizobotrya alpina* | S0906 | 924 | 924 | 255 | 447 | 5,759 | 1,078,497 | 823,140 | 255,357 | 91.4 | 92.7 | 87.5 | 55 | 28 | 27 | 932 | 625 | 307 | 0.17 | 0.17 | 0.18 | 0.96 | 40.34 | 37.76 | 45.60 | 0.83 |
| II | Plagiolobeae | *Plagioloba meyeri* | S1103 | 957 | 952 | 272 | 887 | 5,960 | 1,071,882 | 814,512 | 257,370 | 90.8 | 91.7 | 88.2 | 40 | 22 | 18 | 983 | 659 | 324 | 1.69 | 1.66 | 1.77 | 0.94 | 96.34 | 95.75 | 97.53 | 0.98 |
| II | Schrenkielleae trib. nov. | *Schrenkiella parvula* | S1201sl | 1007 | 1005 | 291 | 614 | 5,768 | 1,127,076 | 855,948 | 271,128 | 95.5 | 96.3 | 92.9 | 38 | 14 | 24 | 1031 | 708 | 323 | 0.04 | 0.03 | 0.08 | 0.39 | 20.17 | 12.57 | 36.84 | 0.34 |
| II | Sisymbrieae | *Sisymbrium aegyptiacum* | SRR8528361 | 712 | 718 | 195 | 244 | 4,170 | 908,238 | 871,599 | 36,639 | 76.9 | 98.1 | 12.6 | 28 | 27 | 1 | 732 | 694 | 38 | 0.54 | 0.55 | 0.51 | 1.07 | 69.67 | 68.88 | 84.21 | 0.82 |
| II | Sisymbrieae | *Sisymbrium altissimum* | S1332 | 1003 | 998 | 288 | 870 | 5,228 | 1,125,120 | 855,183 | 269,937 | 95.3 | 96.3 | 92.5 | 47 | 15 | 32 | 1018 | 704 | 314 | 0.27 | 0.27 | 0.28 | 0.95 | 40.67 | 38.07 | 46.50 | 0.82 |
| II | Sisymbrieae | *Sisymbrium burchellii* | S0345 | 682 | 669 | 171 | 169 | 3,441 | 603,180 | 464,970 | 138,210 | 51.1 | 52.3 | 47.3 | 36 | 11 | 25 | 702 | 471 | 231 | 0.15 | 0.12 | 0.23 | 0.54 | 32.91 | 29.09 | 40.69 | 0.71 |
| II | Sisymbrieae | *Sisymbrium irio* | SRR11470309 | 403 | - | - | - | 1,828 | 464,547 | 325,941 | 138,606 | 39.4 | 36.7 | 47.5 | 0 | 0 | 0 | 452 | 276 | 176 | 0.09 | 0.02 | 0.26 | 0.08 | 3.10 | 2.17 | 4.55 | 0.48 |
| II | Sisymbrieae | *Sisymbrium leucoclada* | SRR8528398 | 698 | 703 | 194 | 209 | 4,119 | 886,830 | 850,515 | 36,315 | 75.1 | 95.7 | 12.4 | 59 | 57 | 2 | 685 | 649 | 36 | 0.88 | 0.89 | 0.71 | 1.26 | 87.15 | 88.14 | 69.44 | 1.27 |
| II | Thelypodieae | *Chaunanthus mexicanus* | S0512sl | 681 | 664 | 175 | 361 | 3,264 | 594,045 | 444,156 | 149,889 | 50.3 | 50 | 51.3 | 21 | 8 | 13 | 708 | 454 | 254 | 2.49 | 2.22 | 3.28 | 0.68 | 91.95 | 90.31 | 94.88 | 0.95 |
| II | Thelypodieae | *Chilocardamum patagonicum* | S1044 | 966 | 965 | 270 | 142 | 5,424 | 1,049,589 | 797,400 | 252,189 | 88.9 | 89.8 | 86.4 | 26 | 16 | 10 | 1010 | 681 | 329 | 5.84 | 5.99 | 5.36 | 1.12 | 98.81 | 98.68 | 99.09 | 1.00 |
| II | Thelypodieae | *Chlorocrambe hastata* | S0286 | 881 | 880 | 244 | 222 | 4,859 | 980,781 | 751,047 | 229,734 | 83.1 | 84.5 | 78.7 | 17 | 10 | 7 | 936 | 625 | 311 | 4.82 | 4.94 | 4.40 | 1.12 | 95.73 | 95.52 | 96.14 | 0.99 |
| II | Thelypodieae | *Dictyophragmus englerianus* | S1481sl | 303 | - | - | - | 1,602 | 109,851 | 85,494 | 24,357 | 9.3 | 9.6 | 8.3 | 14 | 10 | 4 | 351 | 250 | 101 | 0.37 | 0.37 | 0.40 | 0.91 | 29.34 | 30.80 | 25.74 | 1.20 |
| II | Thelypodieae | *Dryopetalon runcinatum* | S0727 | 867 | 867 | 243 | 71 | 5,433 | 1,003,680 | 814,929 | 188,751 | 85 | 91.7 | 64.7 | 7 | 5 | 2 | 918 | 693 | 225 | 5.85 | 5.89 | 5.68 | 1.04 | 97.28 | 96.97 | 98.22 | 0.99 |
| II | Thelypodieae | *Englerocharis peruviana* | S1039 | 955 | 949 | 266 | 140 | 5,421 | 1,025,676 | 781,020 | 244,656 | 86.9 | 87.9 | 83.8 | 23 | 11 | 12 | 1005 | 680 | 325 | 6.03 | 6.27 | 5.30 | 1.18 | 98.81 | 99.26 | 97.85 | 1.01 |
| II | Thelypodieae | *Hesperidanthus argillaceus* | S1072 | 970 | 966 | 272 | 824 | 5,938 | 1,040,364 | 780,117 | 260,247 | 88.1 | 87.8 | 89.1 | 16 | 8 | 8 | 1025 | 691 | 334 | 5.95 | 6.15 | 5.36 | 1.15 | 97.85 | 97.97 | 97.60 | 1.00 |
| II | Thelypodieae | *Hollermayera valdiviana* | S1464 | 971 | 965 | 278 | 571 | 6,085 | 1,013,715 | 761,976 | 251,739 | 85.9 | 85.8 | 86.2 | 28 | 13 | 15 | 1008 | 681 | 327 | 5.26 | 5.40 | 4.83 | 1.12 | 98.02 | 97.80 | 98.47 | 0.99 |
| II | Thelypodieae | *Ivania cremnophila* | S1097 | 911 | 904 | 252 | 782 | 4,841 | 966,312 | 745,755 | 220,557 | 81.9 | 83.9 | 75.6 | 19 | 10 | 9 | 961 | 646 | 315 | 4.31 | 4.37 | 4.09 | 1.07 | 95.01 | 95.36 | 94.29 | 1.01 |
| II | Thelypodieae | *Ivania cremnophila* | S1118 | 847 | 839 | 228 | 793 | 4,505 | 906,816 | 687,807 | 219,009 | 76.8 | 77.4 | 75 | 17 | 6 | 11 | 892 | 590 | 302 | 4.01 | 3.99 | 4.08 | 0.98 | 96.52 | 95.93 | 97.68 | 0.98 |
| II | Thelypodieae | *Mostacillastrum pectinifolium* | S1096 | 989 | 985 | 283 | 834 | 6,417 | 1,059,954 | 792,033 | 267,921 | 89.8 | 89.2 | 91.8 | 21 | 9 | 12 | 1030 | 697 | 333 | 5.57 | 5.73 | 5.10 | 1.12 | 98.35 | 98.42 | 98.20 | 1.00 |
| II | Thelypodieae | *Neuontobotrys linearifolia* | S1098 | 944 | 942 | 260 | 807 | 5,629 | 1,006,716 | 759,168 | 247,548 | 85.3 | 85.5 | 84.8 | 20 | 11 | 9 | 991 | 662 | 329 | 5.39 | 5.53 | 4.96 | 1.12 | 99.09 | 99.70 | 97.87 | 1.02 |
| II | Thelypodieae | *Parodiodoxa chionophila* | S0754 | 949 | 946 | 265 | 248 | 5,952 | 1,034,973 | 773,937 | 261,036 | 87.7 | 87.1 | 89.4 | 19 | 11 | 8 | 996 | 668 | 328 | 5.27 | 5.45 | 4.72 | 1.15 | 94.78 | 94.46 | 95.43 | 0.99 |
| II | Thelypodieae | *Phlebolobium maclovianum* | S1106sl | 976 | 969 | 279 | 895 | 5,656 | 1,001,601 | 760,338 | 241,263 | 84.9 | 85.6 | 82.6 | 15 | 7 | 8 | 1043 | 706 | 337 | 4.68 | 4.79 | 4.34 | 1.10 | 96.74 | 96.74 | 96.74 | 1.00 |
| II | Thelypodieae | *Phravenia viereckii* | S1050 | 942 | 931 | 259 | 143 | 5,426 | 991,782 | 762,348 | 229,434 | 84 | 85.8 | 78.6 | 15 | 8 | 7 | 1008 | 686 | 322 | 5.33 | 5.52 | 4.70 | 1.17 | 96.53 | 96.06 | 97.52 | 0.99 |
| II | Thelypodieae | *Polypsecadium harmsianum* | S1319 | 919 | 911 | 254 | 819 | 5,232 | 989,169 | 750,372 | 238,797 | 83.8 | 84.5 | 81.8 | 30 | 16 | 14 | 960 | 643 | 317 | 4.56 | 4.65 | 4.27 | 1.09 | 96.67 | 97.05 | 95.90 | 1.01 |
| II | Thelypodieae | *Pringlea antiscorbutica* | S1417 | 430 | - | - | - | 2,139 | 206,940 | 167,022 | 39,918 | 17.5 | 18.8 | 13.7 | 18 | 12 | 6 | 470 | 337 | 133 | 3.16 | 3.07 | 3.54 | 0.87 | 92.34 | 92.58 | 91.73 | 1.01 |
| II | Thelypodieae | *Raphanorhyncha crassa* | SRR8528388 | 712 | 711 | 198 | 217 | 4,579 | 843,405 | 809,883 | 33,522 | 71.5 | 91.2 | 11.5 | 2 | 2 | 0 | 763 | 713 | 50 | 5.70 | 5.74 | 4.73 | 1.21 | 96.33 | 98.18 | 70.00 | 1.40 |
| II | Thelypodieae | *Romanschulzia arabiformis* | S0346 | 967 | 964 | 272 | 251 | 6,352 | 1,029,711 | 761,865 | 267,846 | 87.2 | 85.8 | 91.8 | 34 | 14 | 20 | 1000 | 676 | 324 | 5.24 | 5.36 | 4.90 | 1.09 | 96.60 | 96.01 | 97.84 | 0.98 |
| II | Thelypodieae | *Sarcodraba karraikensis* | S0759 | 131 | - | - | - | 673 | 41,775 | 33,294 | 8,481 | 3.5 | 3.7 | 2.9 | 1 | 0 | 0 | 153 | 114 | 39 | 0.40 | 0.36 | 0.52 | 0.70 | 28.10 | 28.95 | 25.64 | 1.13 |
| II | Thelypodieae | *Sibara angelorum* | S1040 | 938 | 930 | 257 | 139 | 5,542 | 1,010,019 | 763,191 | 246,828 | 85.6 | 85.9 | 84.6 | 18 | 7 | 11 | 991 | 667 | 324 | 5.37 | 5.48 | 5.03 | 1.09 | 97.38 | 97.60 | 96.91 | 1.01 |
| II | Thelypodieae | *Stanleya pinnata* | S0521 | 939 | 930 | 260 | 494 | 5,898 | 1,040,835 | 780,693 | 260,142 | 88.2 | 87.9 | 89.1 | 19 | 12 | 7 | 989 | 661 | 328 | 6.25 | 6.46 | 5.62 | 1.15 | 97.78 | 98.03 | 97.26 | 1.01 |
| II | Thelypodieae | *Streptanthus maculatus* | S1053sl | 869 | 863 | 240 | 120 | 4,523 | 887,493 | 693,918 | 193,575 | 75.2 | 78.1 | 66.3 | 17 | 8 | 9 | 914 | 608 | 306 | 3.89 | 3.85 | 4.05 | 0.95 | 97.16 | 97.86 | 95.75 | 1.02 |
| II | Thelypodieae | *Thelypodiopsis elegans* | S1049sl | 5 | - | - | - | 36 | 2,016 | 1,863 | 153 | 0.2 | 0.2 | 0.1 | 0 | 0 | 0 | 8 | 7 | 1 | 0.00 | 0.00 | 0.00 | NA | 0.00 | 0.00 | 0.00 | NA |
| II | Thelypodieae | *Thysanocarpus curvipes* | S0647-2 | 982 | 972 | 287 | 531 | 5,414 | 1,055,976 | 804,804 | 251,172 | 89.5 | 90.6 | 86 | 2 | 2 | 0 | 1057 | 716 | 341 | 6.33 | 6.57 | 5.58 | 1.18 | 97.82 | 97.63 | 98.24 | 0.99 |
| II | Thelypodieae | *Warea amplexifolia* | S1258-2 | 972 | 957 | 274 | 350 | 5,118 | 1,021,326 | 791,466 | 229,860 | 86.5 | 89.1 | 78.7 | 3 | 3 | 0 | 1042 | 702 | 340 | 5.06 | 5.20 | 4.58 | 1.14 | 95.59 | 95.87 | 95.00 | 1.01 |
| II | Thelypodieae | *Weberbauera densifolia* | S1117 | 944 | 942 | 266 | 883 | 5,166 | 1,002,324 | 771,300 | 231,024 | 84.9 | 86.8 | 79.1 | 15 | 10 | 5 | 1001 | 680 | 321 | 6.01 | 6.20 | 5.36 | 1.16 | 99.20 | 99.41 | 98.75 | 1.01 |
| II | Thelypodieae | *Zuloagocardamum jujuyensis* | S1459 | 961 | 959 | 269 | 390 | 4,944 | 967,557 | 751,680 | 215,877 | 82 | 84.6 | 73.9 | 22 | 9 | 13 | 1011 | 686 | 325 | 4.46 | 4.51 | 4.27 | 1.06 | 97.82 | 98.25 | 96.92 | 1.01 |
| II | Thlaspideae | *Alliaria petiolata* | S1419 | 965 | 962 | 275 | 865 | 5,992 | 1,068,411 | 817,131 | 251,280 | 90.5 | 92 | 86.1 | 46 | 28 | 18 | 984 | 660 | 324 | 2.65 | 2.66 | 2.63 | 1.01 | 99.19 | 99.24 | 99.07 | 1.00 |
| II | Thlaspideae | *Didymophysa aucheri* | S0766 | 917 | 919 | 264 | 155 | 5,753 | 1,049,361 | 801,753 | 247,608 | 88.9 | 90.2 | 84.8 | 23 | 13 | 10 | 957 | 635 | 322 | 2.08 | 2.01 | 2.31 | 0.87 | 90.60 | 90.87 | 90.06 | 1.01 |
| II | Thlaspideae | *Elburzia fenestrata* | S0738sl | 492 | - | - | - | 2,974 | 265,287 | 189,384 | 75,903 | 22.5 | 21.3 | 26 | 14 | 3 | 11 | 541 | 337 | 204 | 0.28 | 0.25 | 0.37 | 0.69 | 40.11 | 39.17 | 41.67 | 0.94 |
| II | Thlaspideae | *Graellsia saxifragifolia* | S0744sl | 568 | - | - | - | 3,531 | 368,097 | 268,995 | 99,102 | 31.2 | 30.3 | 33.9 | 21 | 12 | 9 | 617 | 399 | 218 | 0.32 | 0.30 | 0.38 | 0.79 | 43.60 | 44.86 | 41.28 | 1.09 |
| II | Thlaspideae | *Lysakia rostrata* | S1449 | 1004 | 1003 | 293 | 654 | 6,090 | 1,125,441 | 855,630 | 269,811 | 95.4 | 96.3 | 92.4 | 51 | 27 | 24 | 1016 | 695 | 321 | 0.50 | 0.47 | 0.58 | 0.82 | 66.93 | 62.59 | 76.32 | 0.82 |
| II | Thlaspideae | *Mummenhoffia alliacea* | S1214sl | 978 | 973 | 283 | 477 | 6,061 | 1,114,395 | 851,019 | 263,376 | 94.4 | 95.8 | 90.2 | 41 | 17 | 24 | 996 | 683 | 313 | 0.17 | 0.16 | 0.20 | 0.82 | 25.80 | 22.25 | 33.55 | 0.66 |
| II | Thlaspideae | *Pachyphragma macrophyllum* | S1422 | 990 | 988 | 286 | 881 | 6,005 | 1,078,089 | 822,375 | 255,714 | 91.3 | 92.6 | 87.6 | 88 | 44 | 44 | 967 | 666 | 301 | 0.19 | 0.18 | 0.22 | 0.82 | 62.67 | 62.01 | 64.12 | 0.97 |
| II | Thlaspideae | *Parlatoria cakiloidea* | S0292 | 977 | 975 | 280 | 248 | 6,028 | 1,115,907 | 850,359 | 265,548 | 94.5 | 95.7 | 91 | 102 | 49 | 53 | 938 | 649 | 289 | 0.50 | 0.50 | 0.48 | 1.05 | 83.58 | 83.82 | 83.04 | 1.01 |
| II | Thlaspideae | *Peltaria alliacea* | SRR8528336 | 716 | 719 | 199 | 659 | 4,864 | 916,722 | 880,545 | 36,177 | 77.7 | 99.1 | 12.4 | 64 | 63 | 1 | 700 | 662 | 38 | 0.19 | 0.19 | 0.19 | 0.98 | 62.43 | 62.54 | 60.53 | 1.03 |
| II | Thlaspideae | *Peltariopsis planisiliqua* | S0384 | 830 | 822 | 234 | 437 | 5,208 | 1,011,165 | 764,166 | 246,999 | 85.7 | 86 | 84.6 | 50 | 28 | 22 | 833 | 546 | 287 | 0.87 | 0.84 | 0.97 | 0.87 | 90.40 | 90.48 | 90.24 | 1.00 |
| II | Thlaspideae | *Pseudocamelina glaucophylla* | S0745 | 1000 | 999 | 290 | 254 | 6,104 | 1,118,172 | 850,632 | 267,540 | 94.7 | 95.7 | 91.6 | 54 | 31 | 23 | 1008 | 687 | 321 | 1.11 | 1.10 | 1.13 | 0.97 | 92.16 | 92.43 | 91.59 | 1.01 |
| II | Thlaspideae | *Pseudovesicaria digitata* | S0381 | 870 | 866 | 239 | 463 | 5,342 | 862,629 | 658,152 | 204,477 | 73.1 | 74.1 | 70 | 59 | 26 | 33 | 879 | 599 | 280 | 0.69 | 0.67 | 0.76 | 0.88 | 89.08 | 88.81 | 89.64 | 0.99 |
| II | Thlaspideae | *Sobolewskia sibirica* | S1366 | 921 | 912 | 259 | 268 | 5,726 | 1,035,282 | 782,709 | 252,573 | 87.7 | 88.1 | 86.5 | 55 | 29 | 26 | 928 | 621 | 307 | 1.03 | 1.03 | 1.05 | 0.98 | 78.12 | 77.94 | 78.50 | 0.99 |
| II | Thlaspideae | *Thlaspi arvense* | S1376 | 968 | 963 | 278 | 618 | 5,988 | 1,095,810 | 837,975 | 257,835 | 92.8 | 94.3 | 88.3 | 43 | 25 | 18 | 987 | 669 | 318 | 0.26 | 0.23 | 0.34 | 0.69 | 35.06 | 34.08 | 37.11 | 0.92 |
| III | Anchonieae | *Dvorakia alaica* | S0827 | 815 | 810 | 223 | 138 | 4,991 | 962,811 | 730,830 | 231,981 | 81.6 | 82.3 | 79.5 | 61 | 35 | 26 | 810 | 538 | 272 | 0.45 | 0.39 | 0.63 | 0.62 | 60.25 | 57.81 | 65.07 | 0.89 |
| III | Anchonieae | *Iskandera hissarica* | S1234 | 933 | 921 | 262 | 490 | 5,899 | 1,059,444 | 801,675 | 257,769 | 89.8 | 90.2 | 88.3 | 44 | 23 | 21 | 956 | 648 | 308 | 2.09 | 2.09 | 2.06 | 1.02 | 84.62 | 82.87 | 88.31 | 0.94 |
| III | Anchonieae | *Matthiola incana* | SRR8528354 | 704 | 693 | 193 | 471 | 4,826 | 875,262 | 841,200 | 34,062 | 74.2 | 94.7 | 11.7 | 20 | 19 | 1 | 731 | 694 | 37 | 0.35 | 0.32 | 0.90 | 0.36 | 27.36 | 26.51 | 43.24 | 0.61 |
| III | Anchonieae | *Micrantha multicaulis* | S0656sl | 835 | 827 | 232 | 452 | 5,163 | 947,475 | 713,445 | 234,030 | 80.3 | 80.3 | 80.2 | 84 | 43 | 41 | 815 | 544 | 271 | 0.77 | 0.74 | 0.85 | 0.87 | 83.56 | 83.82 | 83.03 | 1.01 |
| III | Anchonieae | *Microstigma deflexum* | S1250sl | 926 | 912 | 253 | 325 | 5,690 | 1,027,044 | 780,483 | 246,561 | 87 | 87.9 | 84.5 | 70 | 39 | 31 | 919 | 621 | 298 | 0.88 | 0.81 | 1.11 | 0.73 | 74.21 | 74.24 | 74.16 | 1.00 |
| III | Anchonieae | *Petiniotia purpurascens* | S0899 | 855 | 843 | 236 | 405 | 5,215 | 995,142 | 763,887 | 231,255 | 84.3 | 86 | 79.2 | 49 | 25 | 24 | 860 | 575 | 285 | 1.51 | 1.47 | 1.66 | 0.88 | 77.33 | 77.22 | 77.54 | 1.00 |
| III | Anchonieae | *Sterigmostemum incanum* | S0657 | 835 | 830 | 240 | 458 | 5,294 | 957,024 | 714,591 | 242,433 | 81.1 | 80.4 | 83 | 51 | 27 | 24 | 841 | 544 | 297 | 2.85 | 2.72 | 3.23 | 0.84 | 92.98 | 91.91 | 94.95 | 0.97 |
| III | Anchonieae | *Veselskya griffithiana* | SRR8528358 | 708 | 702 | 192 | 409 | 4,900 | 877,200 | 846,582 | 30,618 | 74.3 | 95.3 | 10.5 | 36 | 32 | 4 | 721 | 687 | 34 | 1.46 | 1.46 | 1.40 | 1.04 | 89.32 | 89.37 | 88.24 | 1.01 |
| III | Buniadeae | *Bunias erucago* | S1405 | 874 | 857 | 240 | 702 | 5,351 | 950,877 | 724,710 | 226,167 | 80.6 | 81.6 | 77.5 | 56 | 32 | 24 | 876 | 579 | 297 | 1.03 | 0.91 | 1.40 | 0.65 | 81.39 | 80.48 | 83.16 | 0.97 |
| III | Chorisporeae | *Chorispora tenella* | SRR8528370 | 699 | 688 | 190 | 205 | 4,795 | 877,629 | 842,370 | 35,259 | 74.4 | 94.8 | 12.1 | 32 | 29 | 3 | 715 | 678 | 37 | 0.03 | 0.03 | 0.06 | 0.51 | 18.18 | 16.96 | 40.54 | 0.42 |
| III | Chorisporeae | *Diptychocarpus strictus* | S0673 | 861 | 849 | 237 | 58 | 5,205 | 1,011,954 | 753,411 | 258,543 | 85.7 | 84.8 | 88.6 | 29 | 14 | 15 | 889 | 581 | 308 | 0.11 | 0.08 | 0.20 | 0.41 | 19.01 | 17.04 | 22.73 | 0.75 |
| III | Chorisporeae | *Litwinowia tenuissima* | S1539 | 982 | 960 | 280 | 523 | 6,012 | 1,092,636 | 823,623 | 269,013 | 92.6 | 92.7 | 92.1 | 86 | 41 | 45 | 962 | 664 | 298 | 0.12 | 0.11 | 0.15 | 0.71 | 42.62 | 36.60 | 56.04 | 0.65 |
| III | Chorisporeae | *Parrya arctica* | S0894 | 860 | 845 | 237 | 420 | 5,102 | 954,063 | 719,430 | 234,633 | 80.8 | 81 | 80.4 | 93 | 44 | 49 | 820 | 543 | 277 | 0.42 | 0.40 | 0.49 | 0.80 | 77.20 | 77.72 | 76.17 | 1.02 |
| III | Chorisporeae | *Parrya darvazica* | S1246 | 872 | 863 | 245 | 744 | 5,268 | 943,725 | 707,688 | 236,037 | 80 | 79.7 | 80.9 | 78 | 38 | 40 | 856 | 570 | 286 | 0.38 | 0.36 | 0.43 | 0.84 | 67.87 | 68.77 | 66.08 | 1.04 |
| III | Chorisporeae | *Parrya maidantalica* | S1245 | 949 | 935 | 266 | 805 | 5,817 | 1,072,611 | 808,017 | 264,594 | 90.9 | 91 | 90.6 | 78 | 41 | 37 | 935 | 632 | 303 | 0.62 | 0.57 | 0.76 | 0.76 | 71.44 | 70.73 | 72.94 | 0.97 |
| III | Dontostemoneae | *Clausia aprica - eastern group* | S1188 | 888 | 881 | 248 | 549 | 5,497 | 985,407 | 742,500 | 242,907 | 83.5 | 83.6 | 83.2 | 40 | 23 | 17 | 912 | 604 | 308 | 2.37 | 2.23 | 2.80 | 0.80 | 92.76 | 92.38 | 93.51 | 0.99 |
| III | Dontostemoneae | *Clausia aprica - western group* | S1187sl | 579 | 564 | 150 | 357 | 3,449 | 476,727 | 344,151 | 132,576 | 40.4 | 38.7 | 45.4 | 18 | 8 | 10 | 617 | 385 | 232 | 1.27 | 1.09 | 1.73 | 0.63 | 73.10 | 70.91 | 76.72 | 0.92 |
| III | Dontostemoneae | *Dontostemon integrifolius* | SRR8528339 | 697 | 695 | 189 | 505 | 4,789 | 878,466 | 842,856 | 35,610 | 74.4 | 94.9 | 12.2 | 38 | 38 | 0 | 713 | 672 | 41 | 1.57 | 1.56 | 1.75 | 0.89 | 90.74 | 90.77 | 90.24 | 1.01 |
| III | Euclidieae | *Anzhenxia yechengnica* | S1368 | 872 | 870 | 245 | 259 | 5,474 | 988,698 | 746,853 | 241,845 | 83.8 | 84.1 | 82.8 | 34 | 21 | 13 | 902 | 595 | 307 | 3.02 | 2.96 | 3.22 | 0.92 | 91.35 | 90.59 | 92.83 | 0.98 |
| III | Euclidieae | *Atelanthera perpusilla* | S1241 | 928 | 920 | 259 | 807 | 6,555 | 1,046,685 | 791,679 | 255,006 | 88.7 | 89.1 | 87.4 | 2 | 1 | 1 | 998 | 666 | 332 | 7.03 | 7.10 | 6.82 | 1.04 | 94.69 | 94.29 | 95.48 | 0.99 |
| III | Euclidieae | *Braya alpina* | S0963 | 785 | 780 | 214 | 648 | 4,768 | 889,008 | 672,942 | 216,066 | 75.3 | 75.7 | 74 | 29 | 15 | 14 | 817 | 528 | 289 | 3.10 | 2.95 | 3.55 | 0.83 | 97.43 | 97.92 | 96.54 | 1.01 |
| III | Euclidieae | *Catenulina hedysaroides* | S0278 | 799 | 778 | 227 | 195 | 4,935 | 906,372 | 682,275 | 224,097 | 76.8 | 76.8 | 76.8 | 20 | 10 | 10 | 838 | 544 | 294 | 0.30 | 0.20 | 0.58 | 0.35 | 27.57 | 21.14 | 39.46 | 0.54 |
| III | Euclidieae | *Christolea crassifolia* | S0417 | 918 | 906 | 263 | 486 | 5,628 | 1,050,822 | 807,525 | 243,297 | 89 | 90.9 | 83.3 | 97 | 58 | 39 | 880 | 613 | 267 | 0.41 | 0.40 | 0.46 | 0.86 | 72.84 | 71.94 | 74.91 | 0.96 |
| III | Euclidieae | *Cryptospora falcata* | S1251 | 907 | 881 | 256 | 311 | 5,552 | 1,000,569 | 745,851 | 254,718 | 84.8 | 84 | 87.3 | 25 | 14 | 11 | 944 | 626 | 318 | 0.68 | 0.61 | 0.89 | 0.69 | 38.77 | 36.10 | 44.03 | 0.82 |
| III | Euclidieae | *Cymatocarpus pilosissimus* | S0624 | 877 | 860 | 248 | 464 | 5,348 | 973,923 | 729,972 | 243,951 | 82.5 | 82.2 | 83.6 | 48 | 22 | 26 | 881 | 582 | 299 | 0.60 | 0.54 | 0.77 | 0.70 | 53.58 | 51.20 | 58.19 | 0.88 |
| III | Euclidieae | *Dichasianthus subtilissimus* | S1451 | 892 | 875 | 251 | 337 | 5,489 | 969,591 | 738,012 | 231,579 | 82.1 | 83.1 | 79.3 | 32 | 15 | 17 | 922 | 613 | 309 | 0.15 | 0.11 | 0.29 | 0.38 | 28.09 | 22.19 | 39.81 | 0.56 |

### Table S4. Continued.

| SAMPLE DETAILS | | | | ASTRAL TAXON OCCUPANCY | | | | | RECOVERED TOTAL LENGTH (BP) | | | RECOVERED TOTAL LENGTH OF TARGET LENGTH (%) | | | # PARALOG CALLS | | | # LOCI RECOVERED | | | ALLELE DIVERGENCE | | | | LOCUS HETEROZYGOSITY | | | |
|---|---|---|---|---|---|---|---|---|---|---|---|---|---|---|---|---|---|---|---|---|---|---|---|---|---|---|---|---|
| Main lineage | Tribe | Species | Sample | inclusive | strict | superstrict | superstrict by tribe | A-Pro | Total | B764 | A353 | Total | B764 | A353 | Total | B764 | A353 | Total | B764 | A353 | Total (%) | B764 (%) | A353(%) | B764/A353 | Total (%) | B764 (%) | A353(%) | B764/A353 |
| III | Euclidieae | *Dilophia salsa* | S1474 | 788 | 781 | 223 | 740 | 4,926 | 904,224 | 681,459 | 222,765 | 76.6 | 76.7 | 76.3 | 54 | 27 | 27 | 793 | 516 | 277 | 1.88 | 1.79 | 2.16 | 0.83 | 95.96 | 95.74 | 96.39 | 0.99 |
| III | Euclidieae | *Euclidium syriacum* | S0320 | 840 | 829 | 243 | 212 | 5,074 | 889,899 | 667,041 | 222,858 | 75.4 | 75.1 | 76.3 | 51 | 27 | 24 | 844 | 556 | 288 | 0.18 | 0.15 | 0.29 | 0.51 | 45.02 | 38.31 | 57.99 | 0.66 |
| III | Euclidieae | *Lachnoloma lehmannii* | S0327 | 835 | 824 | 232 | 209 | 5,115 | 963,912 | 728,652 | 235,260 | 81.7 | 82 | 80.6 | 20 | 9 | 11 | 874 | 577 | 297 | 0.49 | 0.33 | 0.98 | 0.34 | 40.50 | 35.01 | 51.18 | 0.68 |
| III | Euclidieae | *Leiospora exscapa* | S0478 | 795 | 795 | 226 | 440 | 5,032 | 946,518 | 720,633 | 225,885 | 80.2 | 81.1 | 77.4 | 69 | 33 | 36 | 781 | 529 | 252 | 0.32 | 0.29 | 0.43 | 0.68 | 60.82 | 61.06 | 60.32 | 1.01 |
| III | Euclidieae | *Lepidostemon pedunculosus* | S0407sl | 758 | 744 | 213 | 409 | 4,586 | 792,306 | 578,622 | 213,684 | 67.1 | 65.1 | 73.2 | 33 | 15 | 18 | 783 | 497 | 286 | 0.41 | 0.33 | 0.64 | 0.51 | 44.70 | 42.45 | 48.60 | 0.87 |
| III | Euclidieae | *Leptaleum filifolium* | S0299 | 940 | 893 | 256 | 231 | 5,622 | 918,516 | 693,342 | 225,174 | 77.8 | 78 | 77.1 | 79 | 46 | 33 | 925 | 626 | 299 | 0.34 | 0.35 | 0.33 | 1.04 | 60.32 | 60.70 | 59.53 | 1.02 |
| III | Euclidieae | *Metashangrilaia forrestii* | S1475 | 795 | 776 | 222 | 742 | 4,818 | 824,835 | 617,850 | 206,985 | 69.9 | 69.5 | 70.9 | 46 | 24 | 22 | 810 | 524 | 286 | 0.57 | 0.42 | 1.00 | 0.42 | 56.79 | 53.63 | 62.59 | 0.86 |
| III | Euclidieae | *Neotorularia torulosa* | SRR8528347 | 699 | 688 | 190 | 641 | 4,792 | 876,360 | 843,321 | 33,039 | 74.2 | 94.9 | 11.3 | 16 | 14 | 2 | 734 | 698 | 36 | 0.20 | 0.19 | 0.43 | 0.45 | 22.21 | 20.92 | 47.22 | 0.44 |
| III | Euclidieae | *Pycnoplinthus uniflora* | S0428sl | 600 | 589 | 171 | 329 | 3,699 | 579,891 | 423,423 | 156,468 | 49.1 | 47.7 | 53.6 | 44 | 20 | 24 | 609 | 376 | 233 | 0.61 | 0.53 | 0.84 | 0.63 | 60.43 | 59.31 | 62.23 | 0.95 |
| III | Euclidieae | *Rhammatophyllum pachyrhizum* | S1365 | 848 | 843 | 243 | 56 | 5,152 | 937,371 | 700,449 | 236,922 | 79.4 | 78.8 | 81.2 | 46 | 24 | 22 | 860 | 565 | 295 | 1.93 | 1.80 | 2.30 | 0.78 | 93.37 | 92.74 | 94.58 | 0.98 |
| III | Euclidieae | *Rudolf-kamelinia korolkowii* | S0289 | 965 | 953 | 274 | 248 | 5,817 | 1,078,293 | 810,945 | 267,348 | 91.4 | 91.3 | 91.6 | 35 | 18 | 17 | 995 | 673 | 322 | 0.72 | 0.69 | 0.82 | 0.83 | 70.05 | 68.35 | 73.60 | 0.93 |
| III | Euclidieae | *Sisymbriopsis schugnana* | S1239 | 965 | 950 | 271 | 814 | 5,898 | 1,086,171 | 820,146 | 266,025 | 92 | 92.3 | 91.1 | 34 | 22 | 12 | 997 | 669 | 328 | 0.59 | 0.59 | 0.60 | 0.98 | 45.34 | 43.80 | 48.48 | 0.90 |
| III | Euclidieae | *Solms-laubachia pulcherrima* | S0951 | 523 | 509 | 144 | 423 | 3,109 | 518,286 | 369,348 | 148,938 | 43.9 | 41.6 | 51 | 18 | 6 | 12 | 551 | 338 | 213 | 0.67 | 0.45 | 1.20 | 0.38 | 51.72 | 46.75 | 59.62 | 0.78 |
| III | Euclidieae | *Spryginia winkleri* | S0424 | 646 | 641 | 170 | 352 | 4,519 | 828,318 | 802,140 | 26,178 | 70.2 | 90.3 | 9 | 28 | 28 | 0 | 663 | 631 | 32 | 1.25 | 1.23 | 1.86 | 0.66 | 87.48 | 87.80 | 81.25 | 1.08 |
| III | Euclidieae | *Streptoloma desertorum* | S0421 | 949 | 929 | 262 | 484 | 5,751 | 1,050,723 | 793,521 | 257,202 | 89 | 89.3 | 88.1 | 48 | 31 | 17 | 966 | 645 | 321 | 1.72 | 1.68 | 1.84 | 0.91 | 93.69 | 93.18 | 94.70 | 0.98 |
| III | Euclidieae | *Strigosella cabulica* | S1230 | 971 | 952 | 279 | 492 | 5,977 | 1,068,606 | 802,557 | 266,049 | 90.5 | 90.3 | 91.1 | 29 | 20 | 9 | 1009 | 672 | 337 | 1.69 | 1.64 | 1.82 | 0.91 | 89.00 | 87.95 | 91.10 | 0.97 |
| III | Euclidieae | *Synstemon petrovii* | S1232sl | 951 | 942 | 263 | 495 | 5,799 | 1,051,383 | 796,422 | 254,961 | 89.1 | 89.6 | 87.3 | 27 | 14 | 13 | 988 | 662 | 326 | 1.59 | 1.50 | 1.86 | 0.81 | 88.36 | 88.07 | 88.96 | 0.99 |
| III | Euclidieae | *Tetracme quadricornis* | S0645sl | 967 | 943 | 279 | 504 | 5,904 | 1,070,835 | 802,239 | 268,596 | 90.7 | 90.3 | 92 | 84 | 45 | 39 | 951 | 648 | 303 | 0.15 | 0.13 | 0.24 | 0.52 | 48.58 | 45.06 | 56.11 | 0.80 |
| III | Hesperideae | *Hesperis matronalis* | S1328sl | 889 | 868 | 247 | 767 | 5,494 | 942,552 | 714,171 | 228,381 | 79.9 | 80.4 | 78.2 | 49 | 22 | 27 | 902 | 605 | 297 | 1.49 | 1.38 | 1.83 | 0.75 | 87.58 | 86.28 | 90.24 | 0.96 |
| III | Shehbazieae | *Shehbazia tibetica* | S0761sl | 372 | - | - | - | 2,226 | 244,836 | 173,046 | 71,790 | 20.7 | 19.5 | 24.6 | 11 | 6 | 5 | 408 | 253 | 155 | 0.50 | 0.42 | 0.70 | 0.60 | 37.25 | 33.60 | 43.23 | 0.78 |
| IV | Alysseae | *Acuston lunarioides* | S0724sl | 907 | 893 | 257 | 55 | 4,938 | 1,020,387 | 777,411 | 242,976 | 86.4 | 87.5 | 83.2 | 24 | 18 | 6 | 952 | 631 | 321 | 1.14 | 1.03 | 1.48 | 0.69 | 63.87 | 61.65 | 68.22 | 0.90 |
| IV | Alysseae | *Alyssoides utriculata* | S0303 | 768 | 764 | 225 | 196 | 4,189 | 878,307 | 660,867 | 217,440 | 74.4 | 74.4 | 74.5 | 56 | 36 | 20 | 769 | 495 | 274 | 0.43 | 0.40 | 0.53 | 0.75 | 61.90 | 61.41 | 62.77 | 0.98 |
| IV | Alysseae | *Alyssum lassiticum* | S1120 | 880 | 850 | 248 | 632 | 4,955 | 939,312 | 717,345 | 221,967 | 79.6 | 80.7 | 76 | 39 | 22 | 17 | 907 | 598 | 309 | 1.32 | 1.24 | 1.56 | 0.79 | 80.37 | 78.93 | 83.17 | 0.95 |
| IV | Alysseae | *Alyssum montanum* | S0740 | 949 | 924 | 265 | 231 | 5,412 | 1,074,849 | 805,647 | 269,202 | 91.1 | 90.7 | 92.2 | 41 | 27 | 14 | 974 | 647 | 327 | 2.79 | 2.67 | 3.15 | 0.85 | 97.64 | 97.06 | 98.78 | 0.98 |
| IV | Alysseae | *Alyssum taygeteum* | S1262 | 854 | 837 | 239 | 463 | 4,848 | 914,772 | 696,417 | 218,355 | 77.5 | 78.4 | 74.8 | 16 | 10 | 6 | 902 | 591 | 311 | 3.99 | 3.96 | 4.10 | 0.97 | 90.35 | 89.51 | 91.96 | 0.97 |
| IV | Alysseae | *Aurinia gionae* | S1110 | 793 | 784 | 224 | 714 | 4,483 | 871,524 | 660,618 | 210,906 | 73.8 | 74.4 | 72.2 | 61 | 33 | 28 | 796 | 521 | 275 | 0.58 | 0.52 | 0.76 | 0.69 | 73.12 | 71.02 | 77.09 | 0.92 |
| IV | Alysseae | *Aurinia saxatilis* | S1210 | 875 | 866 | 240 | 407 | 4,892 | 976,113 | 739,635 | 236,478 | 82.7 | 83.3 | 81 | 29 | 18 | 11 | 908 | 603 | 305 | 0.48 | 0.46 | 0.56 | 0.82 | 31.94 | 29.19 | 37.38 | 0.78 |
| IV | Alysseae | *Berteroa incana* | S1315 | 966 | 943 | 270 | 261 | 5,390 | 1,071,423 | 810,111 | 261,312 | 90.8 | 91.2 | 89.5 | 48 | 33 | 15 | 977 | 650 | 327 | 1.21 | 1.13 | 1.44 | 0.79 | 79.22 | 79.08 | 79.51 | 0.99 |
| IV | Alysseae | *Bornmuellera tymphaea* | S0308 | 944 | 920 | 265 | 229 | 5,102 | 1,078,263 | 826,320 | 251,943 | 91.4 | 93 | 86.3 | 62 | 41 | 21 | 946 | 643 | 303 | 0.37 | 0.34 | 0.47 | 0.73 | 58.99 | 55.68 | 66.01 | 0.84 |
| IV | Alysseae | *Clastopus vestitus* | S0288 | 591 | 588 | 150 | 134 | 3,764 | 719,061 | 695,859 | 23,202 | 60.9 | 78.3 | 7.9 | 38 | 35 | 3 | 597 | 564 | 33 | 1.04 | 1.04 | 1.16 | 0.90 | 81.07 | 82.09 | 63.64 | 1.29 |
| IV | Alysseae | *Clypeola jonthlaspi* | S0732 | 981 | 965 | 277 | 72 | 5,437 | 1,077,678 | 815,817 | 261,861 | 91.3 | 91.8 | 89.7 | 49 | 26 | 23 | 994 | 675 | 319 | 1.75 | 1.75 | 1.75 | 1.00 | 97.59 | 96.59 | 99.69 | 0.97 |
| IV | Alysseae | *Cuprella homalocarpa* | S1111 | 959 | 948 | 272 | 878 | 5,780 | 1,044,261 | 777,423 | 266,838 | 88.5 | 87.5 | 91.4 | 9 | 6 | 3 | 1021 | 682 | 339 | 6.19 | 6.45 | 5.40 | 1.19 | 91.58 | 91.20 | 92.33 | 0.99 |
| IV | Alysseae | *Degenia velebitica* | S0748 | 933 | 922 | 260 | 234 | 5,082 | 1,081,515 | 814,992 | 266,523 | 91.6 | 91.7 | 91.3 | 86 | 44 | 42 | 909 | 618 | 291 | 0.35 | 0.34 | 0.37 | 0.92 | 59.96 | 60.36 | 59.11 | 1.02 |
| IV | Alysseae | *Fibigia clypeata* | S0322 | 827 | 819 | 226 | 203 | 4,442 | 918,585 | 697,998 | 220,587 | 77.8 | 78.6 | 75.6 | 18 | 12 | 6 | 865 | 568 | 297 | 0.78 | 0.66 | 1.16 | 0.57 | 55.14 | 51.76 | 61.62 | 0.84 |
| IV | Alysseae | *Hormathophylla reverchonii* | S1519 | 918 | 890 | 258 | 323 | 4,989 | 948,954 | 719,415 | 229,539 | 80.4 | 81 | 78.6 | 19 | 11 | 8 | 972 | 647 | 325 | 5.31 | 5.29 | 5.36 | 0.99 | 97.22 | 96.91 | 97.85 | 0.99 |
| IV | Alysseae | *Irania multicaulis* | S1269 | 849 | 838 | 236 | 460 | 4,578 | 937,005 | 704,565 | 232,440 | 79.4 | 79.3 | 79.6 | 35 | 20 | 15 | 873 | 566 | 307 | 1.10 | 0.99 | 1.40 | 0.71 | 73.88 | 73.67 | 74.27 | 0.99 |
| IV | Alysseae | *Irania pendula* | S1266sl | 864 | 851 | 244 | 461 | 4,725 | 964,821 | 728,940 | 235,881 | 81.7 | 82.1 | 80.8 | 42 | 26 | 16 | 888 | 582 | 306 | 1.23 | 1.09 | 1.68 | 0.65 | 79.84 | 80.07 | 79.41 | 1.01 |
| IV | Alysseae | *Irania umbellata* | S1409 | 794 | 784 | 225 | 651 | 4,344 | 906,132 | 681,414 | 224,718 | 76.8 | 76.7 | 77 | 23 | 16 | 7 | 825 | 531 | 294 | 1.47 | 1.34 | 1.87 | 0.72 | 86.79 | 86.44 | 87.41 | 0.99 |
| IV | Alysseae | *Lepidotrichum uechtritzianum* | S0888 | 969 | 952 | 272 | 165 | 5,392 | 1,097,223 | 828,177 | 269,046 | 93 | 93.2 | 92.2 | 91 | 48 | 43 | 937 | 639 | 298 | 0.49 | 0.46 | 0.58 | 0.79 | 67.02 | 64.63 | 72.15 | 0.90 |
| IV | Alysseae | *Lutzia cretica* | S0355 | 541 | 540 | 142 | 117 | 3,403 | 773,805 | 745,335 | 28,470 | 65.6 | 83.9 | 9.8 | 17 | 17 | 0 | 560 | 530 | 30 | 0.59 | 0.57 | 1.02 | 0.56 | 40.00 | 39.62 | 46.67 | 0.85 |
| IV | Alysseae | *Meniocus linifolius* | S0616 | 852 | 841 | 234 | 445 | 4,831 | 991,047 | 741,402 | 249,645 | 84 | 83.5 | 85.5 | 42 | 27 | 15 | 865 | 562 | 303 | 0.23 | 0.17 | 0.39 | 0.44 | 38.27 | 33.10 | 47.85 | 0.69 |
| IV | Alysseae | *Odontarrhena cf. pterocarpa* | S1229-2 | 983 | 969 | 280 | 496 | 5,475 | 1,069,209 | 810,618 | 258,591 | 90.6 | 91.2 | 88.6 | 70 | 37 | 33 | 975 | 661 | 314 | 0.82 | 0.78 | 0.96 | 0.80 | 86.87 | 87.29 | 85.99 | 1.02 |
| IV | Alysseae | *Odontarrhena nebrodensis* | S1260 | 955 | 940 | 272 | 329 | 5,305 | 1,054,803 | 811,167 | 243,636 | 89.4 | 91.3 | 83.5 | 43 | 25 | 18 | 973 | 660 | 313 | 1.22 | 1.18 | 1.38 | 0.85 | 81.81 | 81.36 | 82.75 | 0.98 |
| IV | Alysseae | *Odontarrhena obovata* | S0739 | 976 | 967 | 278 | 243 | 5,392 | 1,097,808 | 831,345 | 266,463 | 93 | 93.6 | 91.3 | 59 | 26 | 33 | 981 | 673 | 308 | 0.89 | 0.90 | 0.88 | 1.02 | 86.54 | 86.03 | 87.66 | 0.98 |
| IV | Alysseae | *Odontarrhena tavolarae* | S1176 | 885 | 881 | 251 | 635 | 4,946 | 972,132 | 734,652 | 237,480 | 82.4 | 82.7 | 81.3 | 40 | 23 | 17 | 904 | 597 | 307 | 1.38 | 1.30 | 1.64 | 0.79 | 91.92 | 92.46 | 90.88 | 1.02 |
| IV | Alysseae | *Phyllolepidum rupestre* | S0952 | 835 | 834 | 237 | 687 | 4,673 | 989,850 | 761,706 | 228,144 | 83.9 | 85.7 | 78.2 | 53 | 34 | 19 | 844 | 564 | 280 | 0.47 | 0.39 | 0.73 | 0.53 | 52.25 | 48.58 | 59.64 | 0.81 |
| IV | Alysseae | *Physoptychis caspica* | S0430 | 837 | 825 | 234 | 429 | 4,534 | 950,982 | 724,392 | 226,590 | 80.6 | 81.5 | 77.6 | 62 | 39 | 23 | 824 | 550 | 274 | 0.51 | 0.47 | 0.64 | 0.73 | 69.78 | 69.27 | 70.80 | 0.98 |
| IV | Alysseae | *Pterygostemon spathulatus* | S1249 | 843 | 834 | 234 | 710 | 4,492 | 928,044 | 689,349 | 238,695 | 78.6 | 77.6 | 81.8 | 30 | 19 | 11 | 875 | 567 | 308 | 0.95 | 0.86 | 1.24 | 0.69 | 65.83 | 64.73 | 67.86 | 0.95 |
| IV | Alysseae | *Resetnikia triquetra* | S0620 | 511 | 496 | 131 | 266 | 2,675 | 555,399 | 407,454 | 147,945 | 47.1 | 45.9 | 50.7 | 18 | 13 | 5 | 529 | 322 | 207 | 0.63 | 0.51 | 0.95 | 0.54 | 60.30 | 58.39 | 63.29 | 0.92 |
| IV | Arabideae | *Abdra brachycarpa* | S0834 | 897 | 887 | 240 | 153 | 4,603 | 862,434 | 674,340 | 188,094 | 73.1 | 75.9 | 64.4 | 17 | 10 | 7 | 957 | 639 | 318 | 4.53 | 4.45 | 4.82 | 0.92 | 97.28 | 97.65 | 96.54 | 1.01 |
| IV | Arabideae | *Acirostrum alaschanicum* | S0419 | 886 | 885 | 247 | 477 | 5,462 | 1,038,441 | 784,869 | 253,572 | 88 | 88.3 | 86.9 | 80 | 38 | 42 | 860 | 576 | 284 | 0.17 | 0.18 | 0.15 | 1.17 | 47.44 | 49.48 | 43.31 | 1.14 |
| IV | Arabideae | *Arabis alpina* | SRR11470320 | 798 | 784 | 217 | 683 | 4,675 | 882,930 | 629,400 | 253,530 | 74.8 | 70.8 | 86.8 | 22 | 10 | 12 | 840 | 529 | 311 | 0.02 | 0.03 | 0.02 | 1.37 | 13.45 | 16.26 | 8.68 | 1.87 |
| IV | Arabideae | *Arabis blepharophylla* | 8008999 | 145 | - | - | - | 868 | 40,347 | 25,998 | 14,349 | 3.4 | 2.9 | 4.9 | 7 | 1 | 6 | 162 | 97 | 65 | 0.13 | 0.13 | 0.13 | 1.05 | 22.22 | 22.68 | 21.54 | 1.05 |
| IV | Arabideae | *Arabis kennedyae* | S1083 | 993 | 984 | 285 | 808 | 5,829 | 1,119,813 | 848,331 | 271,482 | 94.9 | 95.5 | 93 | 15 | 7 | 8 | 1042 | 705 | 337 | 0.27 | 0.21 | 0.44 | 0.48 | 24.47 | 20.14 | 33.53 | 0.60 |
| IV | Arabideae | *Arabis montbretiana* | S1498 | 983 | 974 | 280 | 916 | 5,776 | 1,082,658 | 827,232 | 255,426 | 91.7 | 93.1 | 87.5 | 66 | 35 | 31 | 980 | 669 | 311 | 0.28 | 0.26 | 0.36 | 0.72 | 51.22 | 47.68 | 58.84 | 0.81 |
| IV | Arabideae | *Arabis scabra* | 8000070 | 123 | - | - | - | 696 | 37,821 | 25,497 | 12,324 | 3.2 | 2.9 | 4.2 | 1 | 0 | 0 | 145 | 90 | 55 | 0.15 | 0.14 | 0.16 | 0.85 | 14.48 | 16.67 | 10.91 | 1.53 |
| IV | Arabideae | *Arcyosperma primulifolium* | S0196sl | 873 | 862 | 237 | 209 | 5,312 | 760,761 | 567,135 | 193,626 | 64.5 | 63.8 | 66.3 | 54 | 26 | 28 | 886 | 591 | 295 | 0.14 | 0.13 | 0.15 | 0.84 | 36.91 | 37.06 | 36.61 | 1.01 |
| IV | Arabideae | *Athysanus pusillus* | S0279 | 959 | 956 | 271 | 250 | 5,810 | 1,029,906 | 767,910 | 261,996 | 87.3 | 86.4 | 89.7 | 17 | 12 | 5 | 1025 | 686 | 339 | 5.53 | 5.62 | 5.25 | 1.07 | 95.90 | 95.19 | 97.35 | 0.98 |
| IV | Arabideae | *Aubrieta deltoidea* | S0281 | 946 | 942 | 268 | 238 | 5,462 | 1,085,556 | 825,747 | 259,809 | 92 | 92.9 | 89 | 64 | 37 | 27 | 946 | 635 | 311 | 0.90 | 0.90 | 0.90 | 1.00 | 71.25 | 71.65 | 70.42 | 1.02 |
| IV | Arabideae | *Baimashania pulvinata* | S0765 | 791 | 791 | 228 | 131 | 4,879 | 976,737 | 735,300 | 241,437 | 82.8 | 82.8 | 82.7 | 66 | 36 | 30 | 775 | 502 | 273 | 0.15 | 0.13 | 0.19 | 0.70 | 51.48 | 50.00 | 54.21 | 0.92 |
| IV | Arabideae | *Botschantzevia karatavica* | S0770 | 931 | 931 | 267 | 157 | 5,760 | 1,093,668 | 828,864 | 264,804 | 92.7 | 93.3 | 90.7 | 80 | 41 | 39 | 912 | 617 | 295 | 0.23 | 0.23 | 0.24 | 0.97 | 61.62 | 61.59 | 61.69 | 1.00 |
| IV | Arabideae | *Dendroarabis fruticulosa* | S0795 | 912 | 910 | 260 | 160 | 5,489 | 1,073,853 | 809,436 | 264,417 | 91 | 91.1 | 90.6 | 66 | 31 | 35 | 904 | 605 | 299 | 0.62 | 0.61 | 0.68 | 0.90 | 71.35 | 71.40 | 71.24 | 1.00 |
| IV | Arabideae | *Draba funckii* | S1113 | 876 | 866 | 252 | 809 | 4,593 | 916,863 | 718,422 | 198,441 | 77.7 | 80.9 | 68 | 17 | 11 | 6 | 935 | 623 | 312 | 3.30 | 3.29 | 3.32 | 0.99 | 98.40 | 99.20 | 96.79 | 1.02 |
| IV | Arabideae | *Draba funckii* | S1121 | 968 | 964 | 276 | 718 | 5,096 | 1,020,879 | 781,257 | 239,622 | 86.5 | 87.9 | 82.1 | 19 | 15 | 4 | 1020 | 684 | 336 | 4.38 | 4.52 | 3.91 | 1.16 | 98.43 | 99.12 | 97.02 | 1.02 |
| IV | Arabideae | *Draba incana* | S0902 | 808 | 798 | 224 | 386 | 4,193 | 806,214 | 625,059 | 181,155 | 68.3 | 70.4 | 62.1 | 41 | 15 | 26 | 837 | 557 | 280 | 1.90 | 1.77 | 2.32 | 0.77 | 95.34 | 95.69 | 94.64 | 1.01 |
| IV | Arabideae | *Drabella muralis* | S0901 | 935 | 920 | 262 | 437 | 5,400 | 1,047,900 | 788,169 | 259,731 | 88.8 | 88.7 | 89 | 20 | 12 | 8 | 981 | 650 | 331 | 3.60 | 3.58 | 3.65 | 0.98 | 98.57 | 98.77 | 98.19 | 1.01 |
| IV | Arabideae | *Pachyneurum grandiflorum* | S0720 | 948 | 946 | 270 | 68 | 5,721 | 1,102,809 | 836,082 | 266,727 | 93.4 | 94.1 | 91.4 | 86 | 38 | 48 | 926 | 635 | 291 | 0.53 | 0.53 | 0.56 | 0.94 | 73.22 | 71.50 | 76.98 | 0.93 |
| IV | Arabideae | *Parryodes axilliflora* | S1177 | 897 | 894 | 255 | 672 | 5,564 | 1,018,218 | 768,090 | 250,128 | 86.3 | 86.5 | 85.7 | 20 | 11 | 9 | 934 | 614 | 320 | 0.31 | 0.27 | 0.43 | 0.64 | 26.34 | 23.13 | 32.50 | 0.71 |
| IV | Arabideae | *Pseudodraba hystrix* | 8009952 | 240 | - | - | - | 1,516 | 81,036 | 57,996 | 23,040 | 6.9 | 6.5 | 7.9 | 3 | 2 | 1 | 283 | 177 | 106 | 0.09 | 0.08 | 0.12 | 0.64 | 14.49 | 15.25 | 13.21 | 1.15 |
| IV | Arabideae | *Pseudoturritis turrita* | SRR8528384 | 719 | 725 | 198 | 218 | 4,917 | 922,671 | 884,268 | 38,403 | 78.2 | 99.5 | 13.2 | 22 | 22 | 0 | 746 | 703 | 43 | 0.16 | 0.16 | 0.22 | 0.74 | 19.84 | 18.21 | 46.51 | 0.39 |
| IV | Arabideae | *Scapiarabis saxicola* | S0760 | 966 | 965 | 272 | 243 | 5,978 | 1,113,492 | 842,526 | 270,966 | 94.3 | 94.8 | 92.8 | 33 | 18 | 15 | 992 | 666 | 326 | 0.20 | 0.16 | 0.31 | 0.50 | 26.21 | 21.02 | 36.81 | 0.57 |
| IV | Arabideae | *Sinoarabis setosifolia* | S1462 | 1004 | 1002 | 293 | 592 | 6,137 | 1,125,639 | 854,232 | 271,407 | 95.4 | 96.2 | 93 | 47 | 27 | 20 | 1017 | 690 | 327 | 0.14 | 0.09 | 0.27 | 0.35 | 29.11 | 21.30 | 45.57 | 0.47 |
| IV | Arabideae | *Tomostima reptans* | S0836 | 910 | 903 | 257 | 156 | 5,114 | 993,036 | 750,771 | 242,265 | 84.1 | 84.5 | 83 | 8 | 4 | 4 | 974 | 649 | 325 | 4.83 | 4.99 | 4.31 | 1.16 | 91.68 | 92.30 | 90.46 | 1.02 |
| IV | Asperuginoideae trib. nov. | *Asperuginoides axillaris* | S1374 | 972 | 968 | 277 | 622 | 5,919 | 1,041,528 | 781,452 | 260,076 | 88.2 | 88 | 89.1 | 51 | 35 | 16 | 987 | 658 | 329 | 4.07 | 4.10 | 3.96 | 1.04 | 99.39 | 99.09 | 100 | 0.99 |
| IV | Stevenieae | *Macropodium nivale* | S0722 | 948 | 938 | 267 | 70 | 6,963 | 1,054,740 | 786,348 | 268,392 | 89.4 | 88.5 | 91.9 | 0 | 0 | 0 | 1022 | 680 | 342 | 5.15 | 5.27 | 4.81 | 1.10 | 91.49 | 91.47 | 91.52 | 1.00 |
| IV | Stevenieae | *Stevenia alyssoides* | S0717 | 905 | 897 | 253 | 64 | 5,763 | 1,044,036 | 784,776 | 259,260 | 88.5 | 88.3 | 88.8 | 6 | 6 | 0 | 964 | 627 | 337 | 3.35 | 3.44 | 3.09 | 1.11 | 92.01 | 92.03 | 91.99 | 1.00 |
| V | Asteae | *Asta schaffneri var. pringlei* | SRR8528394 | 700 | 706 | 193 | 214 | 4,833 | 870,966 | 841,362 | 29,604 | 73.8 | 94.7 | 10.1 | 0 | 0 | 0 | 748 | 712 | 36 | 3.29 | 3.32 | 2.53 | 1.31 | 91.98 | 92.28 | 86.11 | 1.07 |
| V | Asteae | *Scoliaxon mexicanus* | SRR8528385 | 702 | 705 | 190 | 215 | 4,889 | 880,491 | 848,793 | 31,698 | 74.6 | 95.5 | 10.9 | 8 | 8 | 0 | 749 | 712 | 37 | 3.78 | 3.77 | 3.87 | 0.97 | 95.73 | 95.65 | 97.30 | 0.98 |
| V | Chamireae | *Chamira circaeoides* | S1378 | 852 | 823 | 231 | 96 | 5,408 | 970,155 | 728,355 | 241,800 | 82.2 | 82 | 82.8 | 0 | 0 | 0 | 918 | 601 | 317 | 1.79 | 1.91 | 1.40 | 1.36 | 39.11 | 38.60 | 40.06 | 0.96 |
| V | Cremolobeae | *Aimara rollinsii* | S1093sl | 984 | 982 | 286 | 832 | 6,327 | 1,080,276 | 826,425 | 253,851 | 91.5 | 93 | 87 | 2 | 2 | 0 | 1051 | 709 | 342 | 2.89 | 3.05 | 2.38 | 1.28 | 89.44 | 89.99 | 88.30 | 1.02 |
| V | Cremolobeae | *Cremolobus peruvianus* | SRR8528364 | 698 | 699 | 193 | 236 | 5,018 | 886,608 | 855,426 | 31,182 | 75.1 | 96.3 | 10.7 | 1 | 0 | 0 | 750 | 716 | 34 | 2.42 | 2.43 | 2.09 | 1.17 | 79.47 | 79.05 | 88.24 | 0.90 |
| V | Cremolobeae | *Cremolobus suffruticosus* | S1102 | 863 | 858 | 244 | 738 | 5,435 | 946,386 | 713,556 | 232,830 | 80.2 | 80.3 | 79.8 | 33 | 26 | 7 | 889 | 575 | 314 | 1.28 | 1.29 | 1.23 | 1.05 | 64.00 | 64.17 | 63.69 | 1.01 |
| V | Cremolobeae | *Cremolobus suffruticosus* | S1119 | 886 | 881 | 242 | 824 | 5,425 | 921,864 | 696,540 | 225,324 | 78.1 | 78.4 | 77.2 | 5 | 4 | 1 | 957 | 624 | 333 | 6.03 | 6.12 | 5.74 | 1.07 | 91.12 | 89.90 | 93.39 | 0.96 |
| V | Cremolobeae | *Menonvillea linearis* | S0753 | 608 | 597 | 164 | 159 | 3,540 | 697,914 | 534,480 | 163,434 | 59.1 | 60.2 | 56 | 43 | 33 | 10 | 606 | 378 | 228 | 1.03 | 1.05 | 0.97 | 1.08 | 75.58 | 76.72 | 73.68 | 1.04 |
| V | Eudemeae | *Alshehbazia friesii* | S1045 | 993 | 988 | 288 | 151 | 6,573 | 1,110,147 | 838,065 | 272,082 | 94.1 | 94.3 | 93.2 | 0 | 0 | 0 | 1056 | 713 | 343 | 2.96 | 3.16 | 2.36 | 1.34 | 81.25 | 79.94 | 83.97 | 0.95 |
| V | Eudemeae | *Aschersoniodoxa mandoniana* | S0277 | 913 | 896 | 247 | 231 | 5,503 | 923,904 | 715,800 | 208,104 | 78.3 | 80.6 | 71.3 | 45 | 36 | 9 | 935 | 617 | 318 | 4.04 | 4.06 | 3.96 | 1.02 | 94.65 | 95.62 | 92.77 | 1.03 |
| V | Eudemeae | *Brayopsis calycina* | S0989 | 661 | 653 | 176 | 554 | 3,738 | 647,439 | 488,178 | 159,261 | 54.9 | 55 | 54.6 | 20 | 12 | 8 | 689 | 446 | 243 | 3.74 | 3.67 | 3.94 | 0.93 | 96.37 | 95.74 | 97.53 | 0.98 |
| V | Eudemeae | *Dactylocardamum imbricatifolium* | S1043sl | 853 | 849 | 234 | 118 | 5,159 | 950,418 | 721,005 | 229,413 | 80.5 | 81.2 | 78.6 | 27 | 24 | 3 | 886 | 574 | 312 | 4.45 | 4.43 | 4.50 | 0.98 | 94.24 | 94.08 | 94.55 | 1.00 |
| V | Eudemeae | *Eudema rupestris* | S1038 | 910 | 907 | 258 | 134 | 5,691 | 957,567 | 728,268 | 229,299 | 81.1 | 82 | 78.5 | 18 | 14 | 4 | 967 | 646 | 321 | 3.48 | 3.47 | 3.49 | 0.99 | 76.94 | 77.09 | 76.64 | 1.01 |
| V | Eudemeae | *Xerodraba lycopodioides* | S1488 | 111 | - | - | - | 674 | 58,548 | 38,001 | 20,547 | 5 | 4.3 | 7 | 8 | 2 | 6 | 118 | 67 | 51 | 0.94 | 0.77 | 1.26 | 0.61 | 50.00 | 47.76 | 52.94 | 0.90 |
| V | Heliophileae | *Heliophila africana* | S0892 | 869 | 828 | 233 | 409 | 5,515 | 970,383 | 727,467 | 242,916 | 82.2 | 81.9 | 83.2 | 0 | 0 | 0 | 937 | 616 | 321 | 1.99 | 2.03 | 1.89 | 1.07 | 55.82 | 54.22 | 58.88 | 0.92 |
| V | Heliophileae | *Heliophila brachycarpa* | S1107-2 | 417 | - | - | - | 2,453 | 192,141 | 135,390 | 56,751 | 16.3 | 15.2 | 19.4 | 12 | 7 | 5 | 469 | 302 | 167 | 1.30 | 1.13 | 1.69 | 0.67 | 50.53 | 50.00 | 51.50 | 0.97 |
| V | Heliophileae | *Heliophila elongata* | S0577 | 794 | 786 | 210 | 420 | 4,895 | 743,445 | 563,904 | 179,541 | 63 | 63.5 | 61.5 | 9 | 6 | 3 | 851 | 556 | 295 | 2.31 | 2.25 | 2.50 | 0.90 | 84.49 | 84.71 | 84.07 | 1.01 |
| V | Heliophileae | *Heliophila filicaulis* | S1213 | 950 | 937 | 266 | 448 | 6,002 | 1,061,661 | 800,289 | 261,372 | 89.9 | 90.1 | 89.5 | 2 | 2 | 0 | 1018 | 683 | 335 | 3.37 | 3.42 | 3.23 | 1.06 | 93.91 | 94.00 | 93.73 | 1.00 |
| V | Heliophileae | *Heliophila macra* | S1069 | 974 | 958 | 275 | 146 | 6,232 | 1,083,090 | 818,859 | 264,231 | 91.8 | 92.2 | 90.5 | 0 | 0 | 0 | 1037 | 698 | 339 | 3.11 | 3.17 | 2.93 | 1.08 | 82.93 | 82.95 | 82.89 | 1.00 |
| V | Notothlaspideae | *Notothlaspi rosulatum* | S1579 | 898 | 889 | 250 | 611 | 5,632 | 972,162 | 739,365 | 232,797 | 82.4 | 83.2 | 79.7 | 15 | 12 | 3 | 949 | 618 | 331 | 1.94 | 2.02 | 1.69 | 1.20 | 83.35 | 83.82 | 82.48 | 1.02 |
| V | Schizopetaleae | *Atacama nivea* | S1095 | 925 | 923 | 259 | 797 | 6,364 | 1,060,497 | 798,210 | 262,287 | 89.8 | 89.8 | 89.8 | 0 | 0 | 0 | 1001 | 667 | 334 | 4.55 | 4.80 | 3.79 | 1.27 | 86.71 | 86.06 | 88.02 | 0.98 |
| V | Schizopetaleae | *Mathewsia foliosa* | S1089 | 902 | 901 | 247 | 777 | 5,889 | 1,016,886 | 767,877 | 249,009 | 86.2 | 86.4 | 85.3 | 0 | 0 | 0 | 974 | 647 | 327 | 3.58 | 3.71 | 3.19 | 1.16 | 79.26 | 78.36 | 81.04 | 0.97 |
| V | Schizopetaleae | *Schizopetalon walkeri* | SRR8528397 | 700 | 703 | 192 | 210 | 4,914 | 874,113 | 841,269 | 32,844 | 74.1 | 94.7 | 11.3 | 0 | 0 | 0 | 752 | 712 | 40 | 2.98 | 3.02 | 2.07 | 1.46 | 63.43 | 63.76 | 57.50 | 1.11 |
| V | Subularieae | *Idahoa scapigera* | SRR8528381 | 666 | 642 | 173 | 197 | 4,545 | 808,320 | 772,863 | 35,457 | 68.5 | 87 | 12.1 | 60 | 59 | 1 | 660 | 620 | 40 | 0.33 | 0.33 | 0.28 | 1.17 | 47.73 | 47.90 | 45.00 | 1.06 |
| V | Subularieae | *Subularia aquatica* | SRR8528356 | 682 | 658 | 183 | 439 | 4,715 | 838,503 | 804,369 | 34,134 | 71 | 90.5 | 11.7 | 41 | 38 | 3 | 696 | 659 | 37 | 3.87 | 3.90 | 3.15 | 1.24 | 96.12 | 96.97 | 81.08 | 1.20 |
| Unplaced | Anastaticeae | *Anastatica hierochuntica* | S0450 | 970 | 962 | 271 | 500 | 6,134 | 1,092,834 | 831,537 | 261,297 | 92.6 | 93.6 | 89.5 | 0 | 0 | 0 | 1030 | 691 | 339 | 1.61 | 1.68 | 1.40 | 1.20 | 51.07 | 47.90 | 57.52 | 0.83 |
| Unplaced | Anastaticeae | *Cithareloma lehmannii* | S0156 | 706 | 704 | 188 | 174 | 4,461 | 859,500 | 642,912 | 216,588 | 72.8 | 72.4 | 74.2 | 5 | 2 | 3 | 761 | 483 | 278 | 1.18 | 1.20 | 1.13 | 1.06 | 48.49 | 45.34 | 53.96 | 0.84 |
| Unplaced | Anastaticeae | *Diceratella floccosa* | S1114-2 | 708 | 695 | 188 | 631 | 4,273 | 805,548 | 607,266 | 198,282 | 68.2 | 68.4 | 67.9 | 30 | 19 | 11 | 728 | 465 | 263 | 2.07 | 1.98 | 2.35 | 0.84 | 86.26 | 85.59 | 87.45 | 0.98 |
| Unplaced | Anastaticeae | *Eigia longistyla* | S1182 | 881 | 840 | 238 | 524 | 5,456 | 954,561 | 716,793 | 237,768 | 80.9 | 80.7 | 81.4 | 104 | 81 | 23 | 840 | 540 | 300 | 0.63 | 0.61 | 0.69 | 0.89 | 80.00 | 80.93 | 78.33 | 1.03 |
| Unplaced | Anastaticeae | *Eremobium aegyptiacum* | S0654 | 804 | 807 | 225 | 441 | 5,081 | 920,358 | 696,585 | 223,773 | 78 | 78.4 | 76.7 | 102 | 76 | 26 | 764 | 493 | 271 | 1.66 | 1.60 | 1.86 | 0.86 | 91.36 | 91.08 | 91.88 | 0.99 |
| Unplaced | Anastaticeae | *Farsetia aegyptia* | S0691 | 981 | 970 | 279 | 72 | 6,145 | 1,084,671 | 823,818 | 260,853 | 91.9 | 92.7 | 89.4 | 61 | 50 | 11 | 981 | 650 | 331 | 3.93 | 3.90 | 4.03 | 0.97 | 98.17 | 98.00 | 98.49 | 1.00 |
| Unplaced | Anastaticeae | *Lachnocapsa spathulata* | S0655sl | 958 | 948 | 267 | 507 | 5,926 | 1,010,001 | 769,593 | 240,408 | 85.6 | 86.6 | 82.4 | 26 | 18 | 8 | 999 | 668 | 331 | 3.65 | 3.66 | 3.61 | 1.01 | 88.29 | 86.98 | 90.94 | 0.96 |
| Unplaced | Anastaticeae | *Lobularia maritima* | S0555 | 908 | 865 | 249 | 477 | 5,706 | 1,019,286 | 760,638 | 258,648 | 86.4 | 85.6 | 88.6 | 87 | 69 | 18 | 888 | 575 | 313 | 1.28 | 1.30 | 1.21 | 1.07 | 66.44 | 64.70 | 69.65 | 0.93 |
| Unplaced | Anastaticeae | *Marcus-kochia littorea* | S1428 | 938 | 890 | 256 | 595 | 5,864 | 1,015,605 | 762,456 | 253,149 | 86 | 85.8 | 86.7 | 93 | 71 | 22 | 911 | 596 | 315 | 0.89 | 0.86 | 0.98 | 0.88 | 71.57 | 69.13 | 76.19 | 0.91 |
| Unplaced | Anastaticeae | *Maresia nana* | S0403sl | 897 | 854 | 243 | 457 | 5,494 | 858,663 | 636,702 | 221,961 | 72.7 | 71.7 | 76 | 98 | 75 | 23 | 871 | 563 | 308 | 0.19 | 0.18 | 0.20 | 0.91 | 50.52 | 48.49 | 54.22 | 0.89 |
| Unplaced | Anastaticeae | *Morettia canescens* | S0791 | 624 | 623 | 162 | 105 | 4,435 | 824,925 | 824,925 | 0 | 69.9 | 92.9 | 0 | 20 | 20 | 0 | 652 | 652 | 0 | 3.52 | 3.52 | NA | NA | 93.71 | 93.71 | NA | NA |
| Unplaced | Anastaticeae | *Notoceras bicorne* | S0642 | 899 | 855 | 239 | 471 | 5,585 | 1,041,390 | 784,779 | 256,611 | 88.2 | 88.3 | 87.9 | 3 | 2 | 1 | 957 | 630 | 327 | 1.42 | 1.52 | 1.10 | 1.38 | 49.32 | 47.62 | 52.60 | 0.91 |
| Unplaced | Anastaticeae | *Parolinia ornata* | S0436 | 928 | 917 | 258 | 478 | 5,888 | 1,032,504 | 797,802 | 234,702 | 87.5 | 89.8 | 80.4 | 46 | 41 | 5 | 947 | 628 | 319 | 1.92 | 1.91 | 1.95 | 0.98 | 84.27 | 85.67 | 81.50 | 1.05 |
| Unplaced | Biscutelleae | *Biscutella didyma* | S0560 | 783 | 768 | 205 | 410 | 4,939 | 935,385 | 747,168 | 188,217 | 79.2 | 84.1 | 64.5 | 0 | 0 | 0 | 835 | 577 | 258 | 2.37 | 2.44 | 2.09 | 1.16 | 57.49 | 57.54 | 57.36 | 1.00 |
| Unplaced | Biscutelleae | *Heldreichia bupleurifolia* | SRR8528338 | 731 | 735 | 199 | 548 | 4,896 | 926,364 | 881,997 | 44,367 | 78.5 | 99.3 | 15.2 | 33 | 33 | 0 | 747 | 694 | 53 | 0.24 | 0.23 | 0.30 | 0.77 | 63.45 | 64.27 | 52.83 | 1.22 |
| Unplaced | Biscutelleae | *Lunaria rediviva* | SRR8528382 | 699 | 697 | 188 | 206 | 5,871 | 855,945 | 820,356 | 35,589 | 72.5 | 92.3 | 12.2 | 1 | 0 | 0 | 749 | 706 | 43 | 4.66 | 4.66 | 4.79 | 0.97 | 79.71 | 80.31 | 69.77 | 1.15 |
| Unplaced | Biscutelleae | *Megadenia pygmaea* | S0329 | 907 | 907 | 259 | 235 | 5,552 | 1,050,561 | 806,997 | 243,564 | 89 | 90.8 | 83.4 | 39 | 17 | 22 | 927 | 628 | 299 | 0.07 | 0.06 | 0.12 | 0.50 | 24.16 | 19.59 | 33.78 | 0.58 |
| Unplaced | Biscutelleae | *Ricotia aucheri* | S1077 | 911 | 905 | 254 | 770 | 5,942 | 1,035,372 | 794,331 | 241,041 | 87.7 | 89.4 | 82.6 | 168 | 131 | 37 | 814 | 529 | 285 | 1.13 | 1.11 | 1.20 | 0.92 | 92.01 | 93.19 | 89.82 | 1.04 |
| Unplaced | Biscutelleae | *Ricotia cretica* | SRR8528399 | 700 | 701 | 194 | 212 | 5,024 | 867,075 | 834,822 | 32,253 | 73.5 | 94 | 11 | 0 | 0 | 0 | 748 | 709 | 39 | 3.09 | 3.10 | 2.80 | 1.11 | 87.43 | 87.73 | 82.05 | 1.07 |
| Unplaced | Biscutelleae | *Ricotia lunaria* | S0746 | 951 | 949 | 269 | 234 | 6,325 | 1,087,629 | 816,915 | 270,714 | 92.1 | 92 | 92.7 | 0 | 0 | 0 | 1012 | 672 | 340 | 3.35 | 3.45 | 3.05 | 1.13 | 77.27 | 75.89 | 80.00 | 0.95 |
| Unplaced | Biscutelleae | *Ricotia sinuata* | S1074 | 974 | 968 | 277 | 825 | 6,445 | 1,089,504 | 822,882 | 266,622 | 92.3 | 92.6 | 91.3 | 0 | 0 | 0 | 1037 | 697 | 340 | 3.88 | 3.96 | 3.61 | 1.10 | 91.42 | 91.25 | 91.76 | 0.99 |
| Unplaced | Cochlearieae | *Cochlearia officinalis* | SRR8528373 | 683 | 662 | 180 | 205 | 4,813 | 854,547 | 816,405 | 38,142 | 72.4 | 91.9 | 13.1 | 105 | 100 | 5 | 634 | 593 | 41 | 0.98 | 1.00 | 0.69 | 1.44 | 88.64 | 89.38 | 78.05 | 1.15 |
| Unplaced | Cochlearieae | *Ionopsidium acaule* | S0284 | 695 | 666 | 188 | 169 | 4,324 | 742,776 | 538,674 | 204,102 | 62.9 | 60.6 | 69.9 | 75 | 53 | 22 | 684 | 414 | 270 | 0.99 | 0.97 | 1.05 | 0.92 | 67.69 | 65.94 | 70.37 | 0.94 |
| Unplaced | Iberideae I | *Iberis fontqueri* | S1354 | 985 | 959 | 283 | 78 | 6,241 | 1,074,246 | 813,507 | 260,739 | 91 | 91.6 | 89.3 | 110 | 85 | 25 | 940 | 621 | 319 | 2.15 | 2.13 | 2.23 | 0.96 | 95.21 | 95.17 | 95.30 | 1.00 |

### Table S4. Continued.

| SAMPLE DETAILS | | | | ASTRAL TAXON OCCUPANCY | | | | | RECOVERED TOTAL LENGTH (BP) | | | RECOVERED TOTAL LENGTH OF TARGET LENGTH (%) | | | # PARALOG CALLS | | | # LOCI RECOVERED | | | ALLELE DIVERGENCE | | | | LOCUS HETEROZYGOSITY | | | |
|---|---|---|---|---|---|---|---|---|---|---|---|---|---|---|---|---|---|---|---|---|---|---|---|---|---|---|---|---|
| Main lineage | Tribe | Species | Sample | inclusive | strict | superstrict | superstrict by tribe | A-Pro | Total | B764 | A353 | Total | B764 | A353 | Total | B764 | A353 | Total | B764 | A353 | Total (%) | B764 (%) | A353(%) | B764/A353 | Total (%) | B764 (%) | A353(%) | B764/A353 |
| Unplaced | Iberideae II | *Teesdalia coronopifolia* | S1341 | 922 | 882 | 252 | 74 | 5,423 | 934,737 | 680,832 | 253,905 | 79.2 | 76.6 | 87 | 24 | 20 | 4 | 972 | 643 | 329 | 7.32 | 7.20 | 7.64 | 0.94 | 96.30 | 94.87 | 99.09 | 0.96 |
| Unplaced | Iberideae II | *Teesdalia nudicaulis* | S0420sl | 878 | 843 | 243 | 453 | 4,885 | 865,800 | 628,407 | 237,393 | 73.4 | 70.7 | 81.3 | 55 | 42 | 13 | 904 | 589 | 315 | 3.20 | 3.20 | 3.21 | 1.00 | 96.68 | 96.77 | 96.51 | 1.00 |
| Unplaced | Iberideae II | *Teesdalia nudicaulis* | S1342 | 827 | 792 | 227 | 62 | 4,519 | 797,019 | 569,739 | 227,280 | 67.5 | 64.1 | 77.9 | 51 | 36 | 15 | 846 | 540 | 306 | 3.50 | 3.39 | 3.78 | 0.90 | 95.51 | 95.19 | 96.08 | 0.99 |
| Unplaced | Megacarpaeeae | *Megacarpaea megalocarpa* | S0688 | 841 | 838 | 238 | 55 | 5,412 | 995,919 | 756,300 | 239,619 | 84.4 | 85.1 | 82.1 | 0 | 0 | 0 | 903 | 594 | 309 | 2.31 | 2.36 | 2.13 | 1.11 | 76.63 | 77.27 | 75.40 | 1.02 |
| Unplaced | Megacarpaeeae | *Pugionium cornutum* | S0957 | 836 | 831 | 229 | 695 | 5,362 | 967,467 | 734,325 | 233,142 | 82 | 82.7 | 79.9 | 46 | 38 | 8 | 850 | 551 | 299 | 1.58 | 1.66 | 1.31 | 1.27 | 78.82 | 80.40 | 75.92 | 1.06 |
| Outgroup | Akaniaceae | *Akania bidwillii* | HYZL | 289 | 303 | 103 | 311 | 1,506 | - | - | - | - | - | - | - | - | - | - | - | - | - | - | - | - | - | - | - | - |
| Outgroup | Akaniaceae | *Bretschneidera sinensis* | S1384 | 266 | - | - | - | 1,495 | 187,809 | 35,064 | 152,745 | 15.9 | 3.9 | 52.3 | 8 | 0 | 8 | 346 | 72 | 274 | 2.74 | 2.45 | 2.80 | 0.88 | 75.72 | 75.00 | 75.91 | 0.99 |
| Outgroup | Bataceae | *Batis maritima* | PAFTOL_005411 | 65 | - | - | - | 389 | 30,015 | 3,732 | 26,283 | 2.5 | 0.4 | 9 | 1 | 0 | 0 | 99 | 12 | 87 | 0.09 | 0.05 | 0.10 | 0.57 | 17.17 | 16.67 | 17.24 | 0.97 |
| Outgroup | Capparaceae | *Capparicordis crotonoides* | S1387 | 501 | - | - | - | 3,444 | 453,507 | 277,383 | 176,124 | 38.4 | 31.2 | 60.3 | 20 | 17 | 3 | 588 | 308 | 280 | 0.59 | 0.72 | 0.39 | 1.85 | 33.50 | 34.09 | 32.86 | 1.04 |
| Outgroup | Capparaceae | *Capparicordis tweedieana* | S1386 | 501 | - | - | - | 3,319 | 436,806 | 266,772 | 170,034 | 37 | 30 | 58.2 | 35 | 29 | 6 | 560 | 294 | 266 | 0.65 | 0.69 | 0.59 | 1.17 | 55.18 | 52.38 | 58.27 | 0.90 |
| Outgroup | Capparaceae | *Capparis spinosa* | S1388 | 329 | - | - | - | 2,313 | 217,962 | 125,298 | 92,664 | 18.5 | 14.1 | 31.7 | 12 | 6 | 6 | 398 | 211 | 187 | 0.66 | 0.81 | 0.45 | 1.80 | 42.46 | 43.13 | 41.71 | 1.03 |
| Outgroup | Capparaceae | *Capparis tomentosa* | S1389 | 373 | - | - | - | 2,528 | 313,962 | 185,631 | 128,331 | 26.6 | 20.9 | 44 | 3 | 3 | 0 | 431 | 221 | 210 | 1.67 | 1.75 | 1.56 | 1.12 | 61.25 | 59.28 | 63.33 | 0.94 |
| Outgroup | Caricaceae | *Vasconcellea quercifolia* | PAFTOL_019459 | 391 | - | - | - | 2,629 | 347,163 | 145,887 | 201,276 | 29.4 | 16.4 | 68.9 | 26 | 9 | 17 | 501 | 195 | 306 | 0.10 | 0.05 | 0.14 | 0.34 | 34.33 | 18.97 | 44.12 | 0.43 |
| Outgroup | Cleomaceae | *Cleome albescens* | PAFTOL_019361 | 703 | 732 | 218 | 202 | 5,013 | 720,399 | 497,535 | 222,864 | 61 | 56 | 76.3 | 39 | 18 | 21 | 808 | 502 | 306 | 0.16 | 0.14 | 0.21 | 0.68 | 47.15 | 41.24 | 56.86 | 0.73 |
| Outgroup | Emblingiaceae | *Emblingia calceoliflora* | S1398 | 340 | - | - | - | 1,993 | 261,717 | 97,455 | 164,262 | 22.2 | 11 | 56.3 | 14 | 4 | 10 | 412 | 147 | 265 | 0.15 | 0.15 | 0.14 | 1.05 | 37.62 | 32.65 | 40.38 | 0.81 |
| Outgroup | Gyrostemonaceae | *Tersonia cyathiflora* | PAFTOL_023519 | 449 | - | - | - | 3,096 | 374,646 | 200,205 | 174,441 | 31.7 | 22.5 | 59.8 | 48 | 26 | 22 | 526 | 241 | 285 | 0.85 | 0.78 | 0.93 | 0.83 | 84.41 | 79.67 | 88.42 | 0.90 |
| Outgroup | Koeberliniaceae | *Koeberlinia spinosa* | PAFTOL_005410 | 192 | - | - | - | 930 | 137,220 | 17,145 | 120,075 | 11.6 | 1.9 | 41.1 | 7 | 0 | 7 | 243 | 33 | 210 | 0.62 | 0.62 | 0.62 | 1.00 | 30.04 | 30.30 | 30.00 | 1.01 |
| Outgroup | Limnanthaceae | *Limnanthes douglasii* | CRNC | 255 | 275 | 90 | 279 | 1,322 | - | - | - | - | - | - | - | - | - | - | - | - | - | - | - | - | - | - | - | - |
| Outgroup | Malvaceae | *Macrohasseltia macroterantha* | PAFTOL_014331 | 251 | - | - | - | 1,292 | 158,751 | 17,538 | 141,213 | 13.4 | 2 | 48.4 | 3 | 1 | 2 | 356 | 47 | 309 | 3.56 | 3.91 | 3.51 | 1.11 | 78.65 | 85.11 | 77.67 | 1.10 |
| Outgroup | Moringaceae | *Moringa oleifera* | PAFTOL_023477 | 225 | - | - | - | 1,274 | 101,721 | 22,503 | 79,218 | 8.6 | 2.5 | 27.1 | 6 | 0 | 6 | 302 | 75 | 227 | 0.14 | 0.14 | 0.14 | 1.01 | 24.50 | 26.67 | 23.79 | 1.12 |
| Outgroup | Pentadiplandraceae | *Pentadiplandra brazzeana* | S0564sl | 434 | - | - | - | 2,892 | 425,604 | 222,090 | 203,514 | 36.1 | 25 | 69.7 | 30 | 15 | 15 | 526 | 231 | 295 | 0.30 | 0.26 | 0.35 | 0.73 | 61.41 | 54.98 | 66.44 | 0.83 |
| Outgroup | Resedaceae | *Forchhammeria pallida* | PAFTOL_023515 | 438 | - | - | - | 3,231 | 379,038 | 192,777 | 186,261 | 32.1 | 21.7 | 63.8 | 37 | 25 | 12 | 548 | 248 | 300 | 0.15 | 0.12 | 0.17 | 0.70 | 32.66 | 21.77 | 41.67 | 0.52 |
| Outgroup | Salvadoraceae | *Salvadora persica* | PAFTOL_023503 | 116 | - | - | - | 684 | 33,786 | 12,450 | 21,336 | 2.9 | 1.4 | 7.3 | 3 | 2 | 1 | 149 | 48 | 101 | 0.59 | 0.79 | 0.47 | 1.66 | 25.50 | 27.08 | 24.75 | 1.09 |
| Outgroup | Sapotaceae | *Synsepalum afzelii* | S1321 | 301 | - | - | - | 1,689 | 238,494 | 49,227 | 189,267 | 20.2 | 5.5 | 64.8 | 36 | 9 | 27 | 371 | 79 | 292 | 0.36 | 0.26 | 0.39 | 0.67 | 56.33 | 45.57 | 59.25 | 0.77 |
| Outgroup | Setchellanthaceae | *Setchellanthus caeruleus* | PAFTOL_023511 | 290 | - | - | - | 1,645 | 198,336 | 65,709 | 132,627 | 16.8 | 7.4 | 45.4 | 32 | 14 | 18 | 358 | 111 | 247 | 0.25 | 0.19 | 0.28 | 0.66 | 49.44 | 39.64 | 53.85 | 0.74 |
| Outgroup | Tiganophytaceae | *Tiganophyton karense* | PAFTOL_008053 | 249 | - | - | - | 1,296 | 188,535 | 21,945 | 166,590 | 16 | 2.5 | 57.1 | 15 | 2 | 13 | 315 | 47 | 268 | 0.23 | 0.20 | 0.23 | 0.84 | 57.14 | 46.81 | 58.96 | 0.79 |
| Outgroup | Tovariaceae | *Tovaria pendula* | PAFTOL_023513 | 440 | - | - | - | 3,079 | 296,688 | 151,809 | 144,879 | 25.1 | 17.1 | 49.6 | 1 | 0 | 0 | 562 | 261 | 301 | 1.54 | 1.67 | 1.41 | 1.18 | 43.42 | 42.91 | 43.85 | 0.98 |
| Outgroup | Tropaeolaceae | *Tropaeolum peregrinum* | MYZV | 286 | 302 | 100 | 308 | 1,501 | - | - | - | - | - | - | - | - | - | - | - | - | - | - | - | - | - | - | - | - |

Table S5. Overview of tribal node support and age estimates in mya (with 95% HPD), ordered by main lineage (in bold). Results for the nuclear datasets are from the ML supermatrix approach in IQ-TREE and coalescent approach on the 'strict' routine in ASTRAL-III using only the 20 most clocklike genes to estimate branch lengths. Results for the plastome dataset are from the ML supermatrix approach in IQ-TREE. BS: bootstrap support from 100 bootstraps, gCF: gene concordance factor (only for nuclear dataset), sCF: site concordance factor, LPP: local posterior probability, Q1: frequency of most common quartet tree. * indicates maximum support (100% in BS, 1 in LPP). Note that it is impossible to calculate clade support and crown ages for tribes represented by a single sample in our analysis, and such cases are indicated by a hyphen.

| | Clade support | | | | | | | Age estimates (MYA with 95% CI) | | | |
|---|---|---|---|---|---|---|---|---|---|---|---|
| | Nuclear data | | | | | | | | | | |
| | Supermatrix | | | Coalescent | | Plastome data | | Nuclear data | | Plastome data | |
| Main lineage/tribe | BS | gCF | sCF | LPP | Q1 | BS | sCF | Crown | Stem | Crown | Stem |
| **Basal** | ***** | **75** | **65** | **1** | **0.94** | * | 87 | **15.4 (16.4-14.3)** | **24.5 (25.7-23.1)** | **8.6 (12.4-5.6)** | **19.7 (29-13)** |
| Aethionemeae | * | 75 | 65 | 1 | 0.94 | * | 87 | 15.4 (16.4-14.3) | 24.5 (25.7-23.1) | 8.6 (12.4-5.6) | 19.7 (29-13) |
| **I** | ***** | **5** | **52** | **1** | **0.44** | ***** | **52** | **16.3 (17.4-15.4)** | **20.4 (21.7-19.3)** | **12.5 (18.5-8.3)** | **16.4 (24.3-10.2)** |
| Alyssopsideae | * | 54 | 73 | 1 | 0.92 | * | 96 | 6.6 (7.5-5.9) | 8 (8.7-7.2) | 1 (1.5-0.7) | 6.2 (9.3-3.9) |
| Arabidopsideae trib. nov. | * | 69 | 68 | 1 | 0.88 | * | 70 | 6.1 (6.8-5.5) | 12.2 (13.1-11.4) | 4.3 (6.4-2.6) | 5.5 (8.6-3.6) |
| Boechereae | * | 25 | 57 | 1 | 0.64 | * | 67 | 3.6 (4-3.3) | 4 (4.4-3.6) | 6.4 (9.1-4) | 8.1 (12-5.2) |
| Camelineae I | * | 40 | 63 | 1 | 0.75 | * | 42 | 6.5 (7.3-6) | 9.4 (10-8.6) | 5.2 (8.1-3.3) | 5.5 (8.6-3.6) |
| Camelineae III | - | - | - | - | - | - | - | - | 8.5 (9.2-7.6) | - | 3.3 (5-2.1) |
| Cardamineae | * | 53 | 70 | 1 | 0.89 | * | 72 | 10.5 (11.4-9.8) | 16.3 (17.4-15.4) | 8.1 (11.9-5.2) | 11.7 (17.1-7.6) |
| Crucihimalayeae | * | 43 | 62 | 1 | 0.72 | * | 71 | 4.9 (6.1-3.7) | 6.1 (6.9-5.4) | 5.6 (8.1-3.5) | 7.1 (10.2-4.4) |
| Descurainieae | * | 21 | 48 | 1 | 0.69 | * | 49 | 11.9 (12.6-10.8) | 14 (15.1-13.3) | 10.2 (14.7-6.7) | 11 (16.1-7.4) |
| Erysimeae | - | - | - | - | - | * | 95 | - | 6.4 (7-5.6) | 1.4 (2.1-1) | 5.2 (7.7-3.3) |
| Halimolobeae | * | 38 | 60 | 1 | 0.70 | * | 71 | 2.9 (3.3-2.6) | 4 (4.4-3.6) | 5.7 (8.4-3.7) | 8.1 (12-5.2) |
| Hemilophieae trib. nov. | * | 32 | 50 | 1 | 0.48 | * | 88 | 4.9 (6.9-3.1) | 7.2 (7.9-6.7) | 4.5 (6.7-2.8) | 7.8 (11.5-4.9) |
| Lepidieae | * | 51 | 60 | 1 | 0.77 | * | 79 | 11.5 (12.3-10.4) | 15.5 (16.4-14.6) | 7.8 (11.3-5.1) | 11.8 (17.3-7.7) |
| Malcolmieae | - | - | - | - | - | * | 93 | - | 6.4 (7-5.6) | 0.4 (0.6-0.3) | 5.2 (7.7-3.3) |
| Microlepidieae | 49 | 8 | 47 | 1 | 0.50 | * | 66 | 6.2 (6.6-5.7) | 7.2 (7.9-6.7) | 4.2 (6.4-2.7) | 7.5 (11-4.8) |
| Oreophytoneae | * | 89 | 96 | 1 | 0.95 | * | 90 | 0.2 (0.5-0) | 10 (10.9-9.1) | 1.2 (1.8-0.8) | 5.6 (8.6-3.5) |
| Physarieae | * | 74 | 71 | 1 | 0.87 | * | 84 | 8.8 (9.5-8.3) | 12.8 (13.5-11.6) | 6.8 (9.8-4.4) | 10.5 (15.2-6.7) |
| Smelowskieae | - | - | - | - | - | * | 72 | - | 8.6 (11.2-6) | 5.8 (8.3-3.8) | 8.1 (11.6-5.2) |
| Turritideae | - | - | - | - | - | * | 90 | - | 9 (9.7-8.1) | - | 3.3 (5-2.1) |
| Yinshanieae | - | - | - | - | - | * | 92 | - | 8.6 (11.2-6) | 1 (1.5-0.6) | 8.1 (11.6-5.2) |
| **II** | ***** | **1** | **38** | **1** | **0.44** | - | - | **19.1 (20.3-17.6)** | **20.1 (21.3-19)** | **polyphyletic** | **polyphyletic** |
| Aphragmeae | - | - | - | - | - | 99 | 66 | - | 19.1 (20.3-17.6) | 9.7 (15.4-5.2) | 10.9 (17.9-6) |
| Brassiceae | 78 | 7 | 28 | 0.524 | 0.35 | * | 33 | 12.8 (13.8-11.8) | 13.1 (14-12) | 6.5 (10-4.1) | 6.6 (10.2-4.2) |
| Calepineae | * | 66 | 76 | 1 | 0.91 | * | 73 | 6.9 (7.8-6.2) | 16.7 (17.7-15.4) | 6.4 (9.9-4.2) | 10.3 (15.4-6.4) |
| Coluteocarpeae | * | 26 | 39 | 1 | 0.55 | * | 62 | 9.3 (9.9-8.4) | 9.8 (10.5-8.8) | 6.9 (11-3.8) | 7.6 (12.3-4.2) |
| Conringieae | - | - | - | - | - | * | 95 | - | 11.7 (13.2-10.1) | - | 7.6 (12.3-4.2) |
| Eutremeae | * | 88 | 99 | 1 | 0.96 | * | 87 | 0.7 (0.9-0.5) | 14.7 (15.7-13.4) | 5.6 (8.1-3.3) | 10.3 (15.4-6.4) |
| Fourraeeae | * | 30 | 79 | 1 | 0.83 | * | 92 | 2.5 (4.1-1.3) | 13.4 (14.6-12.3) | 2.2 (3.3-1.3) | 7.4 (11.1-4.8) |
| Isatideae | * | 78 | 73 | 1 | 0.95 | * | 86 | 6.8 (7.6-5.8) | 12.6 (13.5-11.6) | 3.7 (5.8-2.5) | 7.2 (11.1-4.6) |
| Kernereae | * | 80 | 80 | 1 | 0.93 | * | 94 | 5.6 (7.6-3.9) | 18.2 (19.1-16.7) | 2.1 (3.2-1.3) | 6.7 (10.1-4.4) |
| Plagiolobeae | - | - | - | - | - | - | - | - | 9.8 (10.5-8.8) | - | 3 (4.8-1.9) |
| Schrenkielleae trib. nov. | - | - | - | - | - | * | 96 | - | 14 (15.1-13) | 0.3 (0.4-0.2) | 7.4 (11.1-4.8) |
| Sisymbrieae | * | 37 | 53 | 1 | 0.78 | * | 80 | 6.6 (7.3-5.9) | 9.3 (10.2-8.4) | 2.2 (3.2-1.3) | 3.8 (5.7-2.3) |
| Thelypodieae | * | 2 | 66 | 1 | 0.42 | * | 67 | 8.3 (9.1-7.5) | 9.3 (10.2-8.4) | 2.8 (4.3-1.5) | 3.8 (5.7-2.3) |
| Thlaspideae | * | 51 | 71 | 1 | 0.90 | * | 68 | 10.8 (11.7-10) | 14.7 (15.7-13.4) | 8.1 (12.9-5.3) | 11.2 (17.3-6.8) |
| **III** | ***** | **40** | **58** | **1** | **0.91** | ***** | **56** | **14.4 (15.4-13.5)** | **21.2 (22.4-19.9)** | **15.3 (22.8-9.8)** | **16.1 (23.8-10)** |
| Anchonieae | * | 55 | 66 | 1 | 0.82 | * | 69 | 6.4 (7.1-5.8) | 10.7 (11.6-10.1) | 6.5 (10.7-3.7) | 9.3 (13.9-6) |
| Buniadeae | - | - | - | - | - | * | 84 | - | 10.7 (11.6-10.1) | 4.5 (6.5-2.6) | 9.8 (14.6-6.4) |
| Chorisporeae | * | 50 | 58 | 1 | 0.91 | * | 57 | 8.5 (9.8-7.5) | 9.2 (10.1-8.2) | 10.2 (15-6.6) | 11.1 (16.3-7.1) |
| Dontostemoneae | * | 73 | 76 | 1 | 0.83 | * | 85 | 5.6 (7.2-4.1) | 12.3 (13-11.2) | 5.9 (8.4-3.8) | 14.8 (22.2-9.7) |
| Euclidieae | * | 25 | 43 | 1 | 0.61 | * | 50 | 11.1 (11.9-10.6) | 12.3 (13.1-11.6) | 10.9 (16-7.2) | 11.9 (17.4-7.9) |
| Hesperideae | - | - | - | - | - | * | 84 | - | 13 (13.9-12.3) | 4.4 (6.5-2.8) | 9.3 (13.9-6) |
| Shehbazieae | - | - | - | - | - | * | 94 | - | 9.2 (10.1-8.2) | - | 11.1 (16.3-7.1) |
| **IV** | **98** | **3** | **36** | **0.843** | **0.38** | - | - | **19.9 (21.3-18.7)** | **20.5 (21.6-19.2)** | **polyphyletic** | **polyphyletic** |
| Alysseae | * | 63 | 72 | 1 | 0.95 | * | 73 | 13.2 (13.8-12) | 17.7 (18.7-16.4) | 11.4 (19.4-6.4) | 12.9 (20.5-7.5) |
| Arabideae | * | 32 | 52 | 1 | 0.76 | NA | NA | 13.6 (15.1-12.2) | 17.3 (18.8-16.3) | 10.8 (17.1-6.7) | 12.1 (18.7-7) |
| Asperuginoideae trib. nov. | - | - | - | - | - | * | 98 | - | 17.7 (18.7-16.4) | - | 14.8 (22.4-9.6) |
| Stevenieae | * | 37 | 59 | 1 | 0.56 | * | 71 | 9.9 (10.7-8.7) | 17.3 (18.8-16.3) | 7.8 (11.7-5.3) | 11.4 (16.4-7.3) |
| **V** | ***** | **1** | **37** | **1** | **0.40** | - | - | **19.3 (20.5-18.2)** | **19.8 (21-18.6)** | **polyphyletic** | **polyphyletic** |
| Asteae | * | 26 | 49 | 1 | 0.51 | * | 66 | 10.3 (13.7-8.3) | 15.7 (17.7-14.1) | 6.4 (10-3.4) | 8.3 (13.4-4.3) |
| Chamireae | - | - | - | - | - | * | 98 | - | 16.9 (18.1-15.9) | - | 9.8 (15.8-5.6) |
| Cremolobeae | * | 19 | 35 | 1 | 0.44 | * | 78 | 10.2 (11.2-9.4) | 10.7 (11.7-9.9) | 3.6 (5.4-2.3) | 6.1 (9.3-3.7) |
| Eudemeae | * | 50 | 62 | 1 | 0.69 | * | 81 | 9.9 (11-9.1) | 14.9 (16.1-13.9) | 5.4 (8.1-3.2) | 7.6 (11.2-4) |
| Heliophileae | * | 66 | 67 | 1 | 0.82 | * | 85 | 11.5 (12.3-10.7) | 16.9 (18.1-15.9) | 6.4 (9.5-3.8) | 9.8 (15.8-5.6) |
| Notothlaspideae | - | - | - | - | - | * | 87 | - | 17.5 (18.8-16.4) | 3.7 (5.8-2) | 10.1 (16.2-5.6) |
| Schizopetaleae | * | 37 | 68 | 1 | 0.57 | * | 85 | 5.4 (6.3-4.3) | 10.7 (11.7-9.9) | 2.4 (3.6-1.4) | 6.7 (10.1-4.4) |
| Subularieae | * | 7 | 43 | 1 | 0.40 | * | 43 | 18.1 (19.5-16.6) | 18.9 (20.1-17.6) | 7.8 (13.4-4.1) | 8.3 (13.4-4.3) |
| **Rogue tribes** | | | | | | | | | | | |
| Anastaticeae | * | 65 | 63 | 1 | 0.85 | * | 71 | 12.4 (13.3-11.8) | 17.1 (18.4-16.2) | 9.2 (14.6-5.4) | 11.3 (19.4-6.8) |
| Biscutelleae | * | 2 | 39 | 1 | 0.42 | * | 54 | 16.8 (17.9-15.4) | 18.7 (19.6-17.5) | 13 (19.5-8.6) | 14.8 (22.4-9.6) |
| Cochlearieae | * | 89 | 80 | 1 | 0.92 | * | 88 | 7.5 (9-6.2) | 16.9 (18.1-15.9) | 6.4 (9.8-4.2) | 12.1 (19.5-6.8) |
| Hillielleae | - | - | - | - | - | - | - | - | - | - | 11.2 (18.9-6.3) |
| Iberideae I | - | - | - | - | - | * | 94 | - | 17.1 (18.4-16.2) | 2.8 (4.5-1.6) | 10.9 (18.5-6.3) |
| Iberideae II | * | 84 | 92 | 1 | 0.90 | * | 99 | 1.9 (2.2-1.7) | 16.8 (18.4-15.4) | - | 10.9 (18.5-6.3) |
| Megacarpaeeae | * | 73 | 69 | 1 | 0.78 | * | 75 | 11.5 (12.7-10.6) | 19.2 (20.4-18.1) | 7.4 (11.9-4.5) | 11.1 (18.7-6.2) |

Table S6. By-tribe mean locus heterozygosity (LH) and allele divergence (AD). Sorted in descending order based on LH. Sample minimum and maximum values given in parentheses. *n* is sample size per tribe.

| | | Locus heterozygosity [%] | | | Allele divergence [%] | | |
|---|---|---|---|---|---|---|---|
| **Tribe** | **n** | **All genes** | **B764** | **A353** | **All genes** | **B764** | **A353** |
| Asperuginoideae trib. nov. | 1 | 99.4 (99.4 - 99.4) | 99.1 (99.1 - 99.1) | 100 (100 - 100) | 4.0 (4.0 - 4.0) | 4.0 (4.0 - 4.0) | 3.9 (3.9 - 3.9) |
| Thelypodieae | 24 | 97.0 (92.2 - 99.2) | 97.0 (90.4 - 99.7) | 95.9 (70.0 - 99.1) | 5.1 (2.5 - 6.3) | 5.3 (2.2 - 6.6) | 4.7 (3.3 - 5.6) |
| Lepidieae | 5 | 96.8 (95.5 - 97.7) | 96.5 (94.9 - 97.6) | 97.6 (96.7 - 97.9) | 4.4 (1.7 - 7.4) | 4.5 (1.8 - 7.6) | 4.2 (1.6 - 6.9) |
| Aphragmeae | 1 | 96.6 (96.6 - 96.6) | 97.4 (97.4 - 97.4) | 95.0 (95.0 - 95.0) | 1.9 (1.9 - 1.9) | 1.9 (1.9 - 1.9) | 2.0 (2.0 - 2.0) |
| Plagiolobeae | 1 | 96.3 (96.3 - 96.3) | 95.7 (95.7 - 95.7) | 97.5 (97.5 - 97.5) | 1.7 (1.7 - 1.7) | 1.7 (1.7 - 1.7) | 1.8 (1.8 - 1.8) |
| Iberideae II | 3 | 96.2 (95.6 - 96.9) | 95.7 (95.0 - 96.9) | 97.3 (96.3 - 99.1) | 4.7 (3.2 - 7.3) | 4.6 (3.2 - 7.2) | 4.9 (3.2 - 7.7) |
| Camelineae III | 2 | 96.1 (96.0 - 96.2) | 96.0 (95.8 - 96.2) | 96.3 (95.6 - 97.0) | 1.4 (1.4 - 1.5) | 1.4 (1.4 - 1.5) | 1.5 (1.4 - 1.6) |
| Iberideae I | 1 | 95.3 (95.3 - 95.3) | 95.3 (95.3 - 95.3) | 95.4 (95.4 - 95.4) | 2.1 (2.1 - 2.1) | 2.1 (2.1 - 2.1) | 2.2 (2.2 - 2.2) |
| Hemilophieae trib. nov. | 2 | 94.4 (90.9 - 97.8) | 94.8 (92.0 - 97.6) | 86.0 (73.9 - 98.1) | 4.6 (2.9 - 6.3) | 4.7 (2.9 - 6.5) | 4.8 (3.9 - 5.6) |
| Asteae | 2 | 93.8 (91.9 - 95.7) | 93.9 (92.2 - 95.6) | 91.7 (86.1 - 97.3) | 3.5 (3.3 - 3.7) | 3.5 (3.3 - 3.7) | 3.2 (2.5 - 3.9) |
| Eutremeae | 2 | 93.0 (92.2 - 93.7) | 92.7 (92.0 - 93.3) | 93.6 (92.7 - 94.5) | 1.6 (0.9 - 2.2) | 1.6 (0.9 - 2.2) | 1.6 (0.8 - 2.3) |
| Stevenieae | 2 | 91.6 (91.3 - 91.8) | 91.7 (91.4 - 92.0) | 91.4 (91.2 - 91.7) | 4.2 (3.3 - 5.1) | 4.3 (3.4 - 5.2) | 3.8 (2.9 - 4.8) |
| Physarieae | 7 | 88.6 (71.3 - 97.8) | 88.4 (71.7 - 98.4) | 89.0 (70.4 - 97.5) | 2.1 (0.8 - 3.6) | 2.1 (0.8 - 3.6) | 2.1 (0.9 - 3.5) |
| Eudemeae | 5 | 88.6 (76.9 - 96.3) | 88.5 (77.1 - 95.7) | 88.9 (76.5 - 97.5) | 3.7 (2.9 - 4.4) | 3.7 (3.1 - 4.4) | 3.6 (2.3 - 4.5) |
| Brassiceae | 47 | 88.2 (56.8 - 98.9) | 88.5 (56.6 - 99.1) | 87.2 (57.2 - 99.4) | 4.2 (2.6 - 9.4) | 4.4 (2.5 - 9.8) | 3.7 (2.1 - 5.8) |
| Hesperideae | 1 | 87.5 (87.5 - 87.5) | 86.2 (86.2 - 86.2) | 90.2 (90.2 - 90.2) | 1.5 (1.5 - 1.5) | 1.4 (1.4 - 1.4) | 1.8 (1.8 - 1.8) |
| Microlepidieae | 16 | 86.7 (56.0 - 99.4) | 86.2 (56.2 - 99.6) | 87.8 (52.5 - 100) | 4.0 (2.6 - 5.8) | 4.0 (2.6 - 6.0) | 3.9 (2.5 - 5.3) |
| Dontostemoneae | 3 | 85.5 (72.9 - 92.9) | 84.7 (70.9 - 92.5) | 86.8 (76.5 - 93.7) | 1.7 (1.2 - 2.4) | 1.6 (1.1 - 2.2) | 2.1 (1.7 - 2.8) |
| Notothlaspideae | 1 | 83.2 (83.2 - 83.2) | 83.6 (83.6 - 83.6) | 82.4 (82.4 - 82.4) | 1.9 (1.9 - 1.9) | 2.0 (2.0 - 2.0) | 1.6 (1.6 - 1.6) |
| Calepineae | 3 | 82.6 (50.3 - 98.8) | 82.3 (49.4 - 99.0) | 83.2 (52.3 - 98.8) | 1.9 (0.2 - 3.2) | 1.9 (0.2 - 3.3) | 1.7 (0.2 - 2.8) |
| Smelowskieae | 1 | 82.5 (82.5 - 82.5) | 79.4 (79.4 - 79.4) | 91.2 (91.2 - 91.2) | 0.7 (0.7 - 0.7) | 0.7 (0.7 - 0.7) | 0.8 (0.8 - 0.8) |
| Buniadeae | 1 | 81.1 (81.1 - 81.1) | 80.2 (80.2 - 80.2) | 82.8 (82.8 - 82.8) | 1.0 (1.0 - 1.0) | 0.9 (0.9 - 0.9) | 1.4 (1.4 - 1.4) |
| Camelineae I | 4 | 80.4 (25.8 - 99.1) | 78.2 (17.9 - 99.0) | 85.2 (43.4 - 99.4) | 2.0 (0.0 - 3.4) | 2.0 (0.0 - 3.5) | 2.0 (0.1 - 3.3) |
| Cardamineae | 12 | 80.1 (23.5 - 98.9) | 79.4 (17.5 - 99.3) | 81.2 (36.8 - 98.2) | 2.6 (0.0 - 5.5) | 2.6 (0.0 - 5.6) | 2.4 (0.1 - 5.1) |
| Cremolobeae | 5 | 79.8 (63.7 - 91.1) | 79.9 (64.0 - 89.9) | 81.3 (63.2 - 93.5) | 2.7 (1.0 - 6.0) | 2.8 (1.0 - 6.1) | 2.5 (1.0 - 5.7) |
| Heliophileae | 4 | 79.1 (55.2 - 93.8) | 78.9 (54.0 - 93.9) | 79.4 (57.6 - 93.5) | 2.7 (2.0 - 3.3) | 2.7 (2.0 - 3.4) | 2.5 (1.8 - 3.1) |
| Cochlearieae | 2 | 78.1 (67.7 - 88.6) | 77.7 (66.0 - 89.3) | 73.9 (70.3 - 77.5) | 1.0 (1.0 - 1.0) | 1.0 (1.0 - 1.0) | 0.8 (0.7 - 1.0) |
| Megacarpaeeae | 2 | 77.4 (76.8 - 78.1) | 78.5 (77.3 - 79.7) | 75.5 (75.3 - 75.6) | 1.8 (1.2 - 2.3) | 1.8 (1.2 - 2.3) | 1.6 (1.2 - 2.1) |
| Schizopetaleae | 3 | 76.2 (63.1 - 86.5) | 75.9 (63.4 - 85.9) | 75.2 (57.5 - 87.6) | 3.7 (3.0 - 4.5) | 3.8 (3.0 - 4.8) | 3.0 (2.1 - 3.7) |
| Anchonieae | 8 | 73.8 (27.5 - 92.9) | 73.0 (26.6 - 91.8) | 77.1 (43.2 - 94.8) | 1.3 (0.3 - 2.8) | 1.2 (0.3 - 2.7) | 1.5 (0.6 - 3.2) |
| Anastaticeae | 13 | 73.6 (47.8 - 98.1) | 72.5 (44.9 - 98.0) | 74.3 (52.4 - 98.4) | 1.8 (0.2 - 3.9) | 1.8 (0.2 - 3.9) | 1.6 (0.2 - 4.0) |
| Thlaspideae | 12 | 72.9 (25.0 - 99.2) | 72.2 (21.2 - 99.2) | 74.5 (33.2 - 99.1) | 0.8 (0.1 - 2.6) | 0.8 (0.1 - 2.6) | 0.9 (0.2 - 2.6) |
| Alysseae | 29 | 72.3 (31.3 - 97.6) | 71.1 (28.4 - 97.0) | 74.4 (37.1 - 99.7) | 1.4 (0.2 - 6.2) | 1.3 (0.2 - 6.4) | 1.5 (0.3 - 5.4) |
| Aethionemeae | 8 | 72.1 (46.0 - 92.4) | 72.8 (44.2 - 93.7) | 70.8 (49.5 - 89.8) | 1.1 (0.2 - 3.7) | 1.1 (0.2 - 3.6) | 1.2 (0.2 - 4.3) |
| Subularieae | 2 | 71.8 (47.5 - 96.2) | 72.4 (47.6 - 97.1) | 63.0 (45.0 - 81.1) | 2.1 (0.3 - 3.9) | 2.1 (0.3 - 3.9) | 1.7 (0.3 - 3.1) |
| Malcolmieae | 1 | 71.5 (71.5 - 71.5) | 67.7 (67.7 - 67.7) | 79.6 (79.6 - 79.6) | 1.0 (1.0 - 1.0) | 0.9 (0.9 - 0.9) | 1.2 (1.2 - 1.2) |
| Biscutelleae | 8 | 71.4 (24.0 - 91.7) | 71.1 (19.3 - 92.9) | 69.4 (34.1 - 91.5) | 2.3 (0.1 - 4.6) | 2.3 (0.1 - 4.6) | 2.2 (0.1 - 4.8) |
| Coluteocarpeae | 13 | 69.9 (16.5 - 98.7) | 68.0 (13.5 - 98.6) | 74.1 (22.6 - 99.1) | 0.9 (0.0 - 4.7) | 0.8 (0.0 - 4.7) | 0.9 (0.1 - 4.8) |
| Halimolobeae | 5 | 66.9 (10.3 - 98.3) | 65.7 ( 8.2 - 98.4) | 69.7 (14.7 - 98.1) | 0.7 (0.1 - 1.9) | 0.7 (0.1 - 1.9) | 0.7 (0.2 - 1.6) |
| Alyssopsideae | 3 | 66.8 (23.3 - 99.6) | 63.2 (14.8 - 99.4) | 74.9 (42.4 - 100) | 1.1 (0.0 - 2.8) | 1.1 (0.0 - 2.8) | 1.1 (0.1 - 2.7) |
| Descurainieae | 5 | 66.1 (15.3 - 100) | 64.3 ( 8.7 - 100) | 69.9 (29.8 - 100) | 1.3 (0.0 - 4.1) | 1.3 (0.0 - 4.2) | 1.2 (0.1 - 3.8) |
| Euclidieae | 26 | 63.7 (22.1 - 97.4) | 61.7 (20.8 - 97.9) | 68.4 (39.2 - 96.5) | 1.2 (0.1 - 7.0) | 1.1 (0.1 - 7.1) | 1.4 (0.2 - 6.8) |
| Arabideae | 21 | 60.6 (13.6 - 98.6) | 59.7 (16.3 - 99.2) | 63.9 ( 8.9 - 98.1) | 1.5 (0.0 - 5.5) | 1.5 (0.0 - 5.6) | 1.5 (0.0 - 5.2) |
| Oreophytoneae | 2 | 59.8 (21.2 - 98.4) | 59.2 (19.8 - 98.5) | 71.5 (45.0 - 98.1) | 1.1 (0.0 - 2.1) | 1.1 (0.0 - 2.1) | 1.1 (0.1 - 2.0) |
| Boechereae | 10 | 59.2 (26.4 - 91.6) | 56.0 (21.3 - 91.2) | 66.2 (37.2 - 92.2) | 0.3 (0.1 - 0.7) | 0.3 (0.1 - 0.6) | 0.4 (0.2 - 0.8) |
| Isatideae | 4 | 57.5 (17.8 - 96.1) | 56.0 (16.3 - 96.4) | 64.7 (37.4 - 89.8) | 0.7 (0.0 - 1.7) | 0.6 (0.0 - 1.7) | 0.6 (0.1 - 1.2) |
| Sisymbrieae | 4 | 57.5 (33.0 - 87.0) | 56.0 (29.3 - 88.0) | 60.0 (40.5 - 84.2) | 0.5 (0.1 - 0.9) | 0.4 (0.1 - 0.9) | 0.4 (0.2 - 0.7) |
| Chorisporeae | 6 | 49.2 (18.2 - 77.1) | 47.8 (16.6 - 77.5) | 55.6 (21.1 - 76.4) | 0.3 (0.0 - 0.6) | 0.3 (0.0 - 0.6) | 0.3 (0.1 - 0.8) |
| Erysimeae | 1 | 44.2 (44.2 - 44.2) | 38.9 (38.9 - 38.9) | 56.3 (56.3 - 56.3) | 0.2 (0.2 - 0.2) | 0.2 (0.2 - 0.2) | 0.3 (0.3 - 0.3) |
| Arabidopsideae trib. nov. | 2 | 42.0 (13.7 - 70.3) | 39.6 (11.4 - 67.8) | 47.2 (18.6 - 75.9) | 0.3 (0.0 - 0.5) | 0.3 (0.0 - 0.5) | 0.4 (0.1 - 0.7) |
| Kernereae | 3 | 41.3 (22.2 - 61.2) | 40.0 (21.1 - 61.1) | 49.2 (40.5 - 61.5) | 0.1 (0.0 - 0.2) | 0.1 (0.0 - 0.2) | 0.2 (0.1 - 0.2) |
| Chamireae | 1 | 38.9 (38.9 - 38.9) | 38.4 (38.4 - 38.4) | 39.9 (39.9 - 39.9) | 1.8 (1.8 - 1.8) | 1.9 (1.9 - 1.9) | 1.3 (1.3 - 1.3) |
| Crucihimalayeae | 2 | 27.8 (25.7 - 29.9) | 23.2 (21.7 - 24.7) | 45.1 (42.5 - 47.7) | 0.1 (0.1 - 0.2) | 0.1 (0.1 - 0.2) | 0.3 (0.1 - 0.5) |
| Fourraeeae | 2 | 24.2 (22.4 - 25.9) | 23.1 (21.3 - 24.9) | 43.2 (42.5 - 43.9) | 0.1 (0.0 - 0.1) | 0.1 (0.0 - 0.1) | 0.1 (0.1 - 0.1) |
| Yinshanieae | 1 | 23.2 (23.2 - 23.2) | 22.2 (22.2 - 22.2) | 42.1 (42.1 - 42.1) | 0.0 (0.0 - 0.0) | 0.0 (0.0 - 0.0) | 0.1 (0.1 - 0.1) |
| Schrenkielleae trib. nov. | 1 | 19.9 (19.9 - 19.9) | 12.4 (12.4 - 12.4) | 36.6 (36.6 - 36.6) | 0.0 (0.0 - 0.0) | 0.0 (0.0 - 0.0) | 0.1 (0.1 - 0.1) |
| Conringieae | 1 | 19.9 (19.9 - 19.9) | 18.3 (18.3 - 18.3) | 51.4 (51.4 - 51.4) | 0.0 (0.0 - 0.0) | 0.0 (0.0 - 0.0) | 0.1 (0.1 - 0.1) |
| Turritideae | 1 | 9.7 (9.7 - 9.7) | 12.4 (12.4 - 12.4) | 4.0 ( 4.0 - 4.0) | 0.0 (0.0 - 0.0) | 0.0 (0.0 - 0.0) | 0.0 (0.0 - 0.0) |

Table S7. Details of samples taken from Walden et al. (2020) and various other studies for this study's plastome phylogeny. Raw data were reanalysed using this study's bioinformatic pipeline.

| Sample (mostly = GenBank accession) | Tribe | Species | Main lineage |
|---|---|---|---|
| AP000423 | Arabidopsideae trib. nov. | *Arabidopsis thaliana* | I |
| JQ806762 | Microlepidieae | *Pachycladon cheesemanii* | I |
| JX205495 | Microlepidieae | *Pachycladon enysii* | I |
| KJ716483 | Brassiceae | *Raphanus sativus* | II |
| KM637653 | Aphragmeae | *Aphragmus involucratus* | II |
| KT962846 | Eutremeae | *Eutrema halophilum* | II |
| KT962847 | Eutremeae | *Eutrema botschantzevii* | II |
| KU170142.1 | Microlepidieae | *Pachycladon fastigiatum* | I |
| KY126841.1 | Arabideae | *Arabis stelleri* | IV |
| KY912021.1 | Anastaticeae | *Anastatica hierochuntica* | |
| KY912024.1 | Euclidieae | *Euclidium syriacum* | III |
| KY912025.1 | Anastaticeae | *Farsetia stylosa* | |
| KY912030.1 | Anchonieae | *Matthiola incana* | III |
| LN866844 | Cochlearieae | *Cochlearia borzaeana* | |
| LN866845 | Cochlearieae | *Cochlearia islandica* | |
| LN866846 | Cochlearieae | *Cochlearia pyrenaica* | |
| LN866847 | Cochlearieae | *Cochlearia tridactylites* | |
| LN866848 | Cochlearieae | *Ionopsidium acaule* | |
| LN877374 | Hesperideae | *Hesperis matronalis* | III |
| LN877375 | Dontostemoneae | *Clausia aprica* | III |
| LN877376 | Buniadeae | *Bunias orientalis* | III |
| LN877377 | Buniadeae | *Bunias erucago* | III |
| LN877378 | Buniadeae | *Bunias orientalis* | III |
| LN877380 | Arabidopsideae trib. nov. | *Arabidopsis carpatica* | I |
| LN877381 | Arabidopsideae trib. nov. | *Arabidopsis cebennensis* | I |
| LN877382 | Arabidopsideae trib. nov. | *Arabidopsis halleri* | I |
| LN877383 | Arabidopsideae trib. nov. | *Arabidopsis lyrata* | I |
| LN877384 | Arabidopsideae trib. nov. | *Arabidopsis pedemontana* | I |
| LN877385 | Camelineae | *Capsella rubella* | I |
| LN877386 | Camelineae I | *Camelina sativa* | I |
| LT629930 | Cochlearieae | *Ionopsidium megalospermum* | |
| MF169880.1 | Alysseae | *Alyssum gmelinii* | IV |
| MK637648 | Alysseae | *Alyssum alyssoides* | IV |
| MK637649 | Asperuginoideae trib. nov. | *Asperuginoides axillaris* | IV |
| MK637650 | Alysseae | *Aurinia corymbosa* | IV |
| MK637651 | Alysseae | *Alyssum dasycarpum* | IV |
| MK637652 | Boechereae | *Anelsonia eurycarpa* | I |
| MK637654 | Fourraeeae | *Hurkaea josiae* | IV |
| MK637655 | Alysseae | *Alyssum linifolium* | IV |
| MK637657 | Alyssopsideae | *Alyssopsis mollis* | I |
| MK637658 | Aphragmeae | *Aphragmus nepalensis* | II |
| MK637659 | Arabideae | *Aubrieta parvifolia* | IV |
| MK637660 | Thlaspideae | *Alliaria petiolata* | II |
| MK637661 | Aethionemeae | *Aethionema saxatile* | basal |
| MK637662 | Asteae | *Asta schaffneri* | V |
| MK637663 | Aethionemeae | *Aethionema spinosum* | basal |
| MK637664 | Microlepidieae | *Arabidella trisecta* | I |
| MK637665 | Arabideae | *Arabis verna* | IV |
| MK637666 | Eudemeae | *Brayopsis alpaminae* | V |
| MK637667 | Boechereae | *Boechera breweri* | I |
| MK637668 | Boechereae | *Boechera davidsonii* | I |
| MK637669 | Biscutelleae | *Biscutella laevigata* | |
| MK637670 | Boechereae | *Boechera laevigata* | I |
| MK637671 | Boechereae | *Boechera puberula* | I |
| MK637672 | Boechereae | *Boechera pulchra* | I |
| MK637673 | Boechereae | *Boechera stricta* | I |
| MK637674 | Boechereae | *Boechera suffrutescens* | I |
| MK637675 | Biscutelleae | *Biscutella vincentina* | |
| MK637676 | Dontostemoneae | *Clausia aprica* | III |
| MK637677 | Cardamineae | *Cardamine bulbifera* | I |
| MK637678 | Chamireae | *Chamira circaeoides* | V |
| MK637679 | Cardamineae | *Cardamine enneaphyllos* | I |
| MK637680 | Cardamineae | *Cardamine glanduligera* | I |
| MK637681 | Cardamineae | *Cardamine hirsuta* | I |
| MK637682 | Calepineae | *Calepina irregularis* | II |
| MK637683 | Alysseae | *Clypeola jonthlaspi* | IV |
| MK637684 | Cardamineae | *Cardamine kitaibelii* | I |
| MK637685 | Brassiceae | *Crambe kralikii* | II |
| MK637686 | Crucihimalayeae | *Crucihimalaya lasiocarpa* | I |
| MK637688 | Brassiceae | *Crambe maritima* | II |
| MK637689 | Conringieae | *Conringia orientalis* | II |
| MK637691 | Cardamineae | *Cardamine pentaphyllos* | I |
| MK637692 | Cremolobeae | *Cremolobus peruvianus* | V |
| MK637693 | Conringieae | *Iljinskaea planisiliqua* | II |
| MK637694 | Conringieae | *Iljinskaea planisiliqua* | II |
| MK637695 | Cardamineae | *Cardamine quinquefolia* | I |
| MK637697 | Chorisporeae | *Chorispora tenella* | III |
| MK637698 | Dontostemoneae | *Clausia trichosepala* | III |
| MK637699 | Arabideae | *Draba aizoides* | IV |
| MK637700 | Physarieae | *Dithyrea californica* | I |
| MK637701 | Descurainieae | *Descurainia erodiifolia* | I |
| MK637702 | Hemilophieae trib. nov. | *Dipoma iberideum* | I |
| MK637703 | Arabideae | *Draba incana* | IV |
| MK637704 | Dontostemoneae | *Dontostemon micranthus* | III |
| MK637705 | Arabideae | *Draba oligosperma* | IV |
| MK637706 | Lepidieae | *Delpinophytum patagonicum* | I |
| MK637707 | Descurainieae | *Descurainia sophia* | I |
| MK637708 | Chorisporeae | *Diptychocarpus strictus* | III |
| MK637709 | Arabideae | *Draba verna* | IV |
| MK637710 | Physarieae | *Dimorphocarpa wislizenii* | I |
| MK637711 | Erysimeae | *Erysimum bicolor* | I |
| MK637712 | Eutremeae | *Eutrema integrifolium* | II |
| MK637713 | Eudemeae | *Eudema nubigena* | V |
| MK637714 | Erysimeae | *Erysimum odoratum* | I |
| MK637715 | Eutremeae | *Eutrema salsugineum* | II |
| MK637716 | Halimolobeae | *Exhalimolobos weddellii* | I |
| MK637717 | Fourraeeae | *Fourraea alpina* | II |
| MK637718 | Brassiceae | *Guiraoa arvensis* | II |
| MK637719 | Calepineae | *Goldbachia laevigata* | II |
| MK637720 | Heliophileae | *Heliophila amplexicaulis* | V |
| MK637721 | Biscutelleae | *Heldreichia bupleurifolia* | |
| MK637722 | Brassiceae | *Henophyton deserti* | II |
| MK637723 | Brassiceae | *Horwoodia dicksoniae* | II |
| MK637724 | Heliophileae | *Heliophila juncea* | V |
| MK637725 | Heliophileae | *Heliophila longifolia* | V |
| MK637726 | Hillielleae | *Hilliella paradoxa* | |
| MK637727 | Hesperideae | *Hesperis persica* | III |
| MK637728 | Descurainieae | *Hornungia petraea* | I |
| MK637729 | Heliophileae | *Heliophila pusilla* | V |
| MK637730 | Hemilophieae trib. nov. | *Hemilophia sessilifolia* | I |
| MK637731 | Hesperideae | *Hesperis tristis* | III |
| MK637732 | Anchonieae | *Iskandera alaica* | III |
| MK637733 | Iberideae I | *Iberis amara* | |
| MK637734 | Cardamineae | *Iodanthus pinnatifidus* | I |
| MK637735 | Subularieae | *Idahoa scapigera* | V |
| MK637736 | Isatideae | *Isatis tinctoria* | II |
| MK637737 | Kernereae | *Kernera saxatilis* | II |
| MK637738 | Biscutelleae | *Lunaria annua* | |
| MK637739 | Lepidieae | *Lepidium echinatum* | I |
| MK637740 | Euclidieae | *Leiospora exscapa* | III |
| MK637741 | Crucihimalayeae | *Ladakiella klimesii* | I |
| MK637742 | Lepidieae | *Lepidium perfoliatum* | I |
| MK637743 | Lepidieae | *Lepidium sativum* | I |
| MK637744 | Chorisporeae | *Litwinowia tenuissima* | III |
| MK637745 | Anchonieae | *Matthiola alyssifolia* | III |
| MK637746 | Schizopetaleae | *Mathewsia auriculata* | V |
| MK637747 | Oreophytoneae | *Murbeckiella boryi* | I |
| MK637748 | Halimolobeae | *Mancoa hispida* | I |
| MK637749 | Cremolobeae | *Menonvillea linearis* | V |
| MK637750 | Anchonieae | *Matthiola longipetala* | III |
| MK637751 | Malcolmieae | *Malcolmia maritima* | I |
| MK637752 | Stevenieae | *Macropodium nivale* | IV |
| MK637753 | Thelypodieae | *Mostacillastrum orbignyanum* | II |
| MK637754 | Anchonieae | *Matthiola ovatifolia* | III |
| MK637755 | Coluteocarpeae | *Microthlaspi perfoliatum* | II |
| MK637756 | Isatideae | *Myagrum perfoliatum* | II |
| MK637757 | Oreophytoneae | *Murbeckiella pinnatifida* | I |
| MK637758 | Megacarpaeeae | *Megacarpaea polyandra* | |

### Table S7. Continued.

| Sample (mostly = GenBank accession) | Tribe | Species | Main lineage |
|---|---|---|---|
| MK637759 | Biscutelleae | *Megadenia pygmaea* | |
| MK637760 | Cremolobeae | *Menonvillea spathulata* | V |
| MK637761 | Notothlaspideae | *Notothlaspi austreale* | V |
| MK637762 | Anastaticeae | *Notoceras bicorne* | |
| MK637763 | Coluteocarpeae | *Noccaea caerulescens* | II |
| MK637764 | Euclidieae | *Neotorularia torulosa* | III |
| MK637765 | Coluteocarpeae | *Noccaea vesicaria* | II |
| MK637766 | Sisymbrieae | *Sisymbrium aegyptiacum* | II |
| MK637767 | Oreophytoneae | *Oreophyton falcatum* | I |
| MK637768 | Cremolobeae | *Menonvillea linearis* | V |
| MK637769 | Brassiceae | *Orychophragmus violaceus* | II |
| MK637770 | Boechereae | *Phoenicaulis cheiranthoides* | I |
| MK637771 | Thelypodieae | *Parodiodoxa chionophila* | II |
| MK637772 | Boechereae | *Polyctenium fremontii* | I |
| MK637773 | Thlaspideae | *Pseudocamelina glaucophylla* | II |
| MK637774 | Physarieae | *Paysonia grandiflora* | I |
| MK637775 | Halimolobeae | *Pennellia longifolia* | I |
| MK637776 | Anastaticeae | *Parolinia ornata* | |
| MK637777 | Chorisporeae | *Parrya pinnatifida* | III |
| MK637778 | Physarieae | *Physaria pinetorum* | I |
| MK637779 | Megacarpaeeae | *Pugionium pterocarpum* | |
| MK637780 | Kernereae | *Petrocallis pyrenaica* | II |
| MK637781 | Camelineae | *Pseudoarabidopsis toxophylla* | I |
| MK637782 | Arabideae | *Pseudoturritis turrita* | IV |
| MK637783 | Kernereae | *Rhizobotrya alpina* | II |
| MK637784 | Cardamineae | *Rorippa islandica* | I |
| MK637785 | Biscutelleae | *Ricotia lunaria* | |
| MK637786 | Descurainieae | *Robeschia schimperi* | I |
| MK637787 | Sisymbrieae | *Sisymbrium aculeolatum* | II |
| MK637788 | Euclidieae | *Strigosella africana* | III |
| MK637789 | Smelowskieae | *Smelowskia altaica* | I |
| MK637790 | Sisymbrieae | *Sisymbrium altissimum* | II |
| MK637791 | Smelowskieae | *Smelowskia annua* | I |
| MK637792 | Subularieae | *Subularia aquatica* | V |
| MK637793 | Isatideae | *Schimpera arabica* | II |
| MK637794 | Smelowskieae | *Smelowskia calycina* | I |
| MK637795 | Stevenieae | *Stevenia cheiranthoides* | IV |
| MK637796 | Thelypodieae | *Streptanthus cooperi* | II |
| MK637797 | Halimolobeae | *Sphaerocardamum divaricatum* | I |
| MK637798 | Smelowskieae | *Smelowskia flavissima* | I |
| MK637799 | Thelypodieae | *Sarcodraba karraikensis* | II |
| MK637800 | Microlepidieae | *Stenopetalum lineare* | I |
| MK637801 | Asteae | *Scoliaxon mexicanus* | V |
| MK637802 | Sisymbrieae | *Sisymbrium officinale* | II |
| MK637803 | Brassiceae | *Savignya parviflora* | II |
| MK637804 | Boechereae | *Sandbergia perplexa* | I |
| MK637805 | Thelypodieae | *Stanleya pinnata* | II |
| MK637806 | Brassiceae | *Schouwia purpurea* | II |
| MK637807 | Arabideae | *Scapiarabis saxicola* | IV |
| MK637808 | Anchonieae | *Sterigmostemum violaceum* | III |
| MK637809 | Schizopetaleae | *Schizopetalon walkeri* | V |
| MK637810 | Thlaspideae | *Thlaspi arvense* | II |
| MK637811 | Turritideae | *Turritis glabra* | I |
| MK637812 | Descurainieae | *Tropidocarpum gracile* | I |
| MK637813 | Thelypodieae | *Thelypodium laciniatum* | II |
| MK637815 | Iberideae II | *Teesdalia nudicaulis* | |
| MK637816 | Brassiceae | *Vella lucentina* | II |
| MK637817 | Yinshanieae | *Yinshania acutangula* | I |
| MK637818 | Yinshanieae | *Yinshania furcatopilosa* | I |
| MK637819 | Yinshanieae | *Yinshania henryi* | I |
| MK637820 | Plagiolobeae | *Plagioloba crenulata* | II |
| MK637821 | Schizopetaleae | *Atacama nivea* | V |
| MK637822 | Conringieae | *Conringia clavata* | II |
| MK637823 | Anastaticeae | *Farsetia occidentalis* | |
| MK637824 | Halimolobeae | *Halimolobos jaegeri* | I |
| MK637825 | Malcolmieae | *Malcolmia bicolor* | I |
| MK637826 | Thelypodieae | *Neuontobotrys berningeri* | II |
| MK637827 | Physarieae | *Nerisyrenia comporum* | I |
| MK637828 | Sisymbrieae | *Sisymbrium leucocladum* | II |

| Sample (mostly = GenBank accession) | Tribe | Species | Main lineage |
|---|---|---|---|
| MK637829 | Shehbazieae | *Shehbazia tibetica* | III |
| MK637830 | Boechereae | *Yosemitea repanda* | I |
| NC_009265 | Aethionemeae | *Aethionema saxatile* | basal |
| NC_009266 | Aethionemeae | *Aethionema grandiflorum* | basal |
| NC_009267 | Alyssopsideae | *Olimarabidopsis pumila* | I |
| NC_009268 | Arabideae | *Arabis hirsuta* | IV |
| NC_009269 | Cardamineae | *Rorippa islandica* | I |
| NC_009270 | Camelineae | *Capsella bursanpastoris* | I |
| NC_009271 | Crucihimalayeae | *Crucihimalaya wallichii* | I |
| NC_009272 | Arabideae | *Draba nemorosa* | IV |
| NC_009273 | Lepidieae | *Lepidium virginicum* | I |
| NC_009274 | Anastaticeae | *Lobularia maritima* | |
| NC_009275 | Cardamineae | *Nasturtium officinale* | I |
| NC_015139.1 | Brassiceae | *Brassica rapa* | II |
| NC_016734 | Brassiceae | *Brassica napus* | II |
| NC_026445 | Cardamineae | *Cardamine impatiens* | I |
| NC_026446 | Cardamineae | *Cardamine resedifolia* | I |
| NC_028272.1 | Brassiceae | *Brassica juncea* | II |
| NC_028517.1 | Camelineae | *Capsella grandiflora* | I |
| NC_028726.1 | Schrenkielleae trib. nov. | *Schrenkiella parvula* | II |
| NC_028727 | Eutremeae | *Eutrema yunnanense* | II |
| NC_028728 | Eutremeae | *Eutrema heterophyllum* | II |
| NC_030450.1 | Brassiceae | *Brassica nigra* | II |
| NC_030515.1 | Megacarpaeeae | *Pugionium dolabratum* | |
| NC_030516.1 | Megacarpaeeae | *Pugionium cornutum* | |
| NC_030775.1 | Brassiceae | *Cakile arabica* | II |
| NC_033498.1 | Brassiceae | *Orychophragmus diffusus* | II |
| NC_033499.1 | Brassiceae | *Orychophragmus taibaiensis* | II |
| NC_033500.1 | Brassiceae | *Orychophragmus hupehensis* | II |
| NC_034286.1 | Brassiceae | *Sinalliaria limprichtiana* | II |
| NC_034287.1 | Brassiceae | *Sinalliaria limprichtiana* | II |
| NC_034299.1 | Alysseae | *Alyssum desertorum* | IV |
| NC_034359.1 | Euclidieae | *Solms-laubachia eurycarpa* | III |
| NC_034360.1 | Megacarpaeeae | *Megacarpaea delavayi* | |
| NC_034361.1 | Euclidieae | *Rudolf-kamelinia korolkowii* | III |
| NC_034363.1 | Lepidieae | *Lepidium meyenii* | I |
| NC_034367 | Aethionemeae | *Aethionema arabicum* | basal |
| NC_035303.1 | Brassiceae | *Sinapis arvensis* | II |
| NC_035512.1 | Hesperideae | *Hesperis sylvestris* | III |
| NC_035513.1 | Anastaticeae | *Lobularia libyca* | |
| NC_035514.1 | Anastaticeae | *Morettia canescens* | |
| NC_035515.1 | Euclidieae | *Braya humilis* | III |

Sample details of outgroup species.

| Sample (mostly = GenBank accession) | Family | Species | Main lineage |
|---|---|---|---|
| MK637687 | Cleomaceae | *Cleome lutea* | |
| EU431223 | Caricaceae | *Carica papaya* | |
| HQ901196 | Malvaceae | *Gossypium hirsutum* | |
| MK637814 | Tropaeolaceae | *Tropaeolum majus* | |
| doi:10.5061/dry-ad.kc5b8_BM | Bataceae | *Batis maritima* | |
| doi:10.5061/dry-ad.kc5b8_RO | Resedaceae | *Reseda odorata* | |
| doi:10.5061/dry-ad.kc5b8_Ssp | Salvadoraceae | *Salvadora sp* | |

Table S8. Overlapping genes among the two bait sets used, the Brassicaceae-specific set targeting 1,827 exons from 764 genes (Nikolov et al. 2019; B764 in the table), and the Angiosperms353 v1 universal bait set (Johnson et al. 2019; A353 in the table). As a result of both bait sets being developed independently, a total of 36 genes showed overlap. In order to avoid studying the same genetic marker twice, we discarded sequencing data from these 36 genes from the B764 set in our analyses, as these were generally shorter target reads than for the A353 set.

| **A353 gene name** | **B764 gene name** |
|---|---|
| 4932 | AT2G31955 |
| 5018 | AT1G08490 |
| 5138 | AT5G61770 |
| 5257 | AT1G43860 |
| 5264 | AT5G67530 |
| 5333 | AT5G13680 |
| 5398 | AT3G20780 |
| 5406 | AT1G03190 |
| 5426 | AT1G49540 |
| 5449 | AT1G55880 |
| 5528 | AT1G17760 |
| 5596 | AT4G29380 |
| 5599 | AT3G51050 |
| 5664 | AT1G07010 |
| 5670 | AT2G39090 |
| 5699 | AT5G17290 |
| 5859 | AT4G29830 |
| 5921 | AT5G48600 |
| 5922 | AT3G12080 |
| 6016 | AT3G08650 |
| 6041 | AT3G53760 |
| 6051 | AT3G02660 |
| 6098 | AT1G31780 |
| 6139 | AT1G75330 |
| 6175 | AT5G06830 |
| 6176 | AT5G64050 |
| 6221 | AT3G06950 |
| 6295 | AT4G33030 |
| 6373 | AT5G48470 |
| 6383 | AT3G48500 |
| 6450 | AT5G10920 |
| 6487 | AT5G14760 |
| 6533 | AT5G46180 |
| 6572 | AT2G21280 |
| 6717 | AT5G18570 |
| 6961 | AT4G02790 |

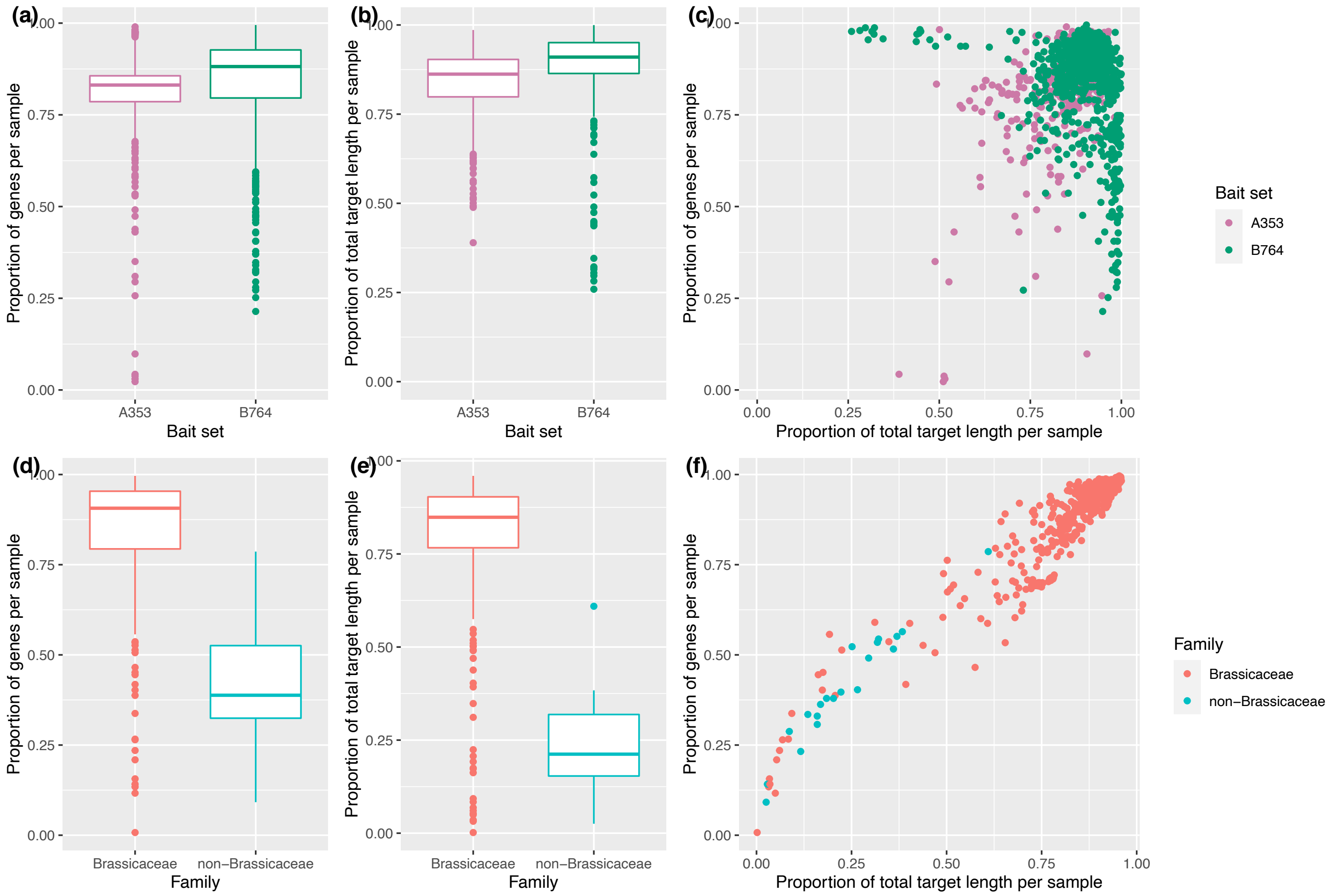


Fig. S1. Nuclear data recovery (in terms of proportions of genes and total target length), split by family (Brassicaceae vs. non-Brassicaceae; lower panels) and bait set (A353 vs. B764; upper panels). In general, our approach was more successful for samples from the Brassicaceae family than for other families, which is likely the result of the inclusion of the Brassicaceae-specific bait set B764. Whereas recovery was generally very good for both bait sets, both proportion of genes and proportion of total target length were slightly higher in the B764 bait set, compared to the A353 bait set.

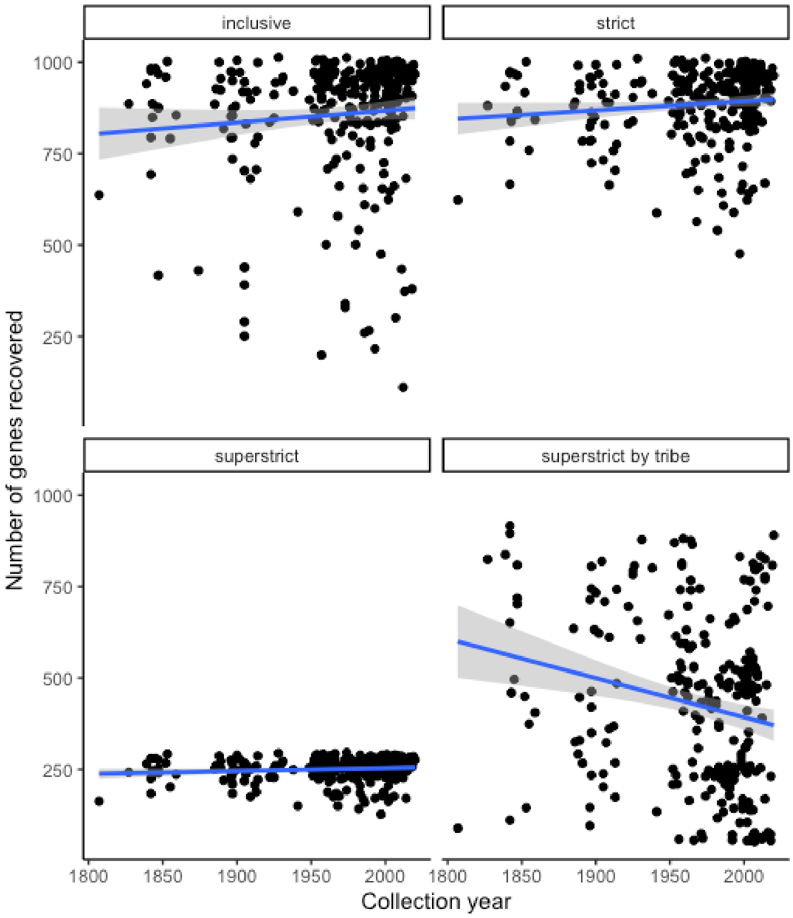


Fig. S2. Number of genes recovered (and passing selection using our more strict routines; see Materials & Methods in the main text) vs. sample collection year with linear regression (n = 277). Many pre-1900 samples were still good enough to recover hundreds of nuclear genes from. Note that gene number drop with sampling routine for the more strict routines, but that this is a combined effect of removing the more variable genes (high SNP proportions) and removing samples with relatively low recovery in terms of gene number and total target length (see table 1 in main text).

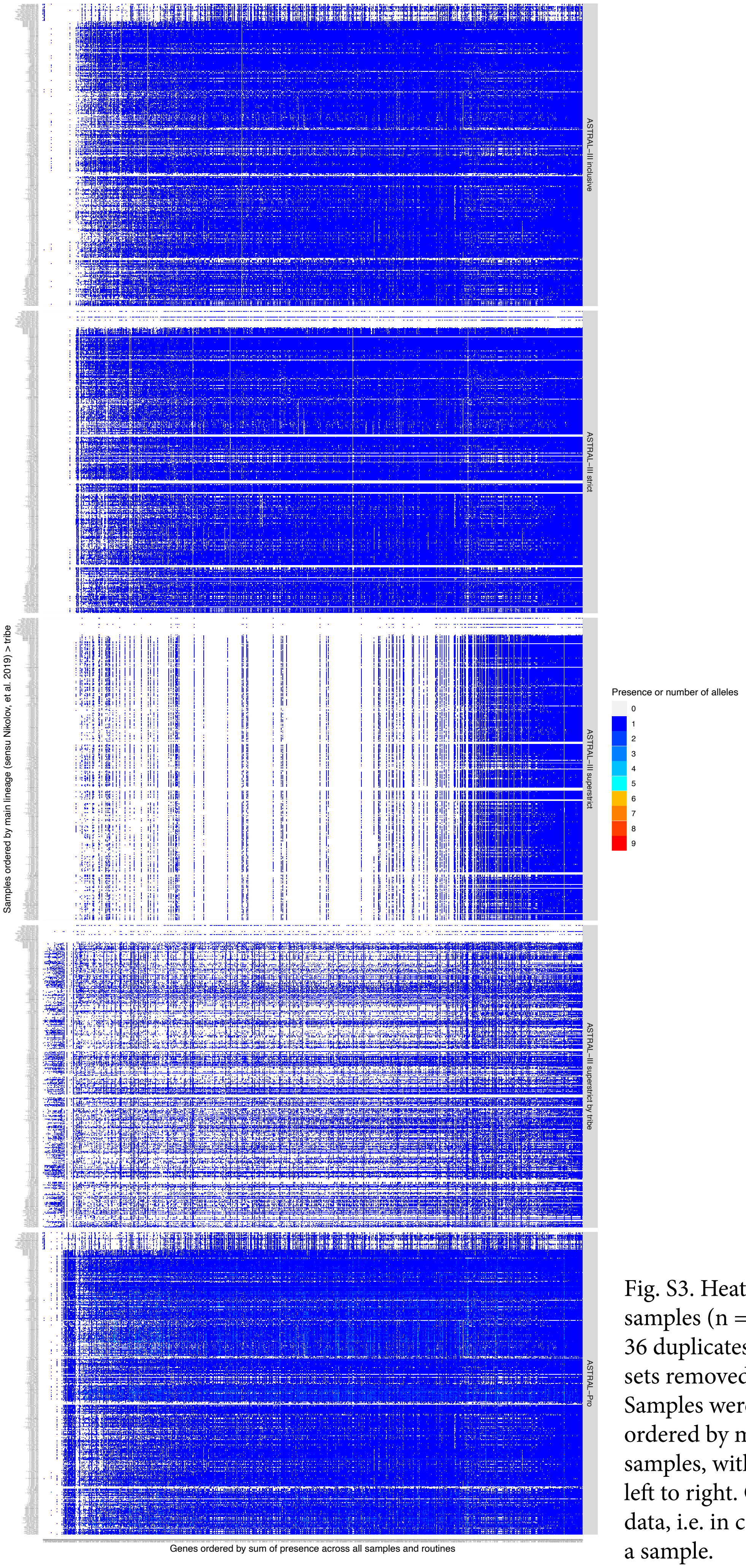


Fig. S3. Heatmap of SNP proportions for all samples (n = 397) and all genes (n = 1,081; 36 duplicates between A353 and B764 bait sets removed) included in our sampling. Samples were clustered by tribe, genes were ordered by mean SNP proportion over all samples, with mean values increasing from left to right. Grey boxes indicate missing data, i.e. in case a gene was not recovered for a sample.

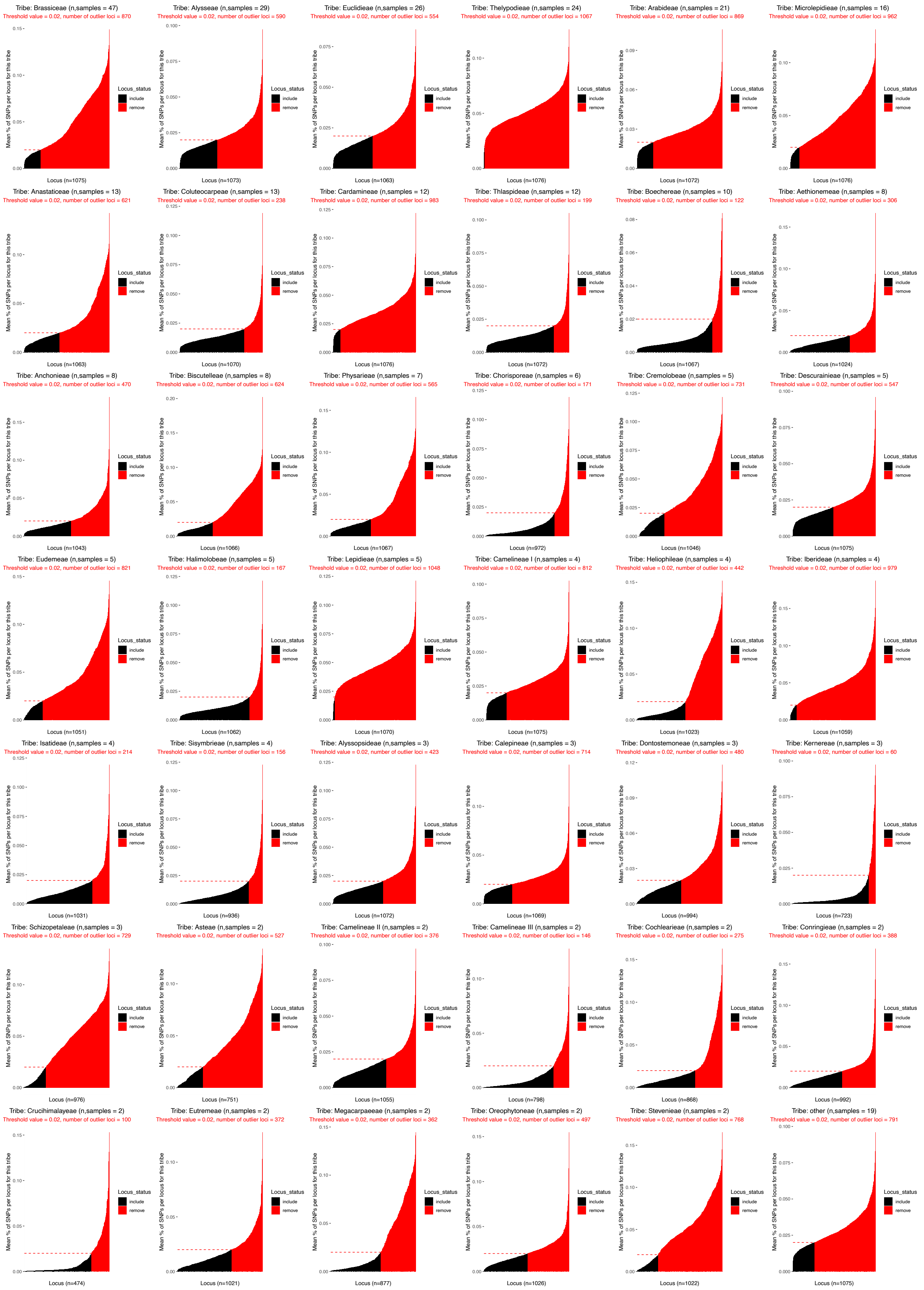


Fig. S4. HybPhaser paralog assessment for the routine 'superstrict by tribe'. Genes to be removed for phylogenetic analyses were removed based on a SNP proportion of >0.02 (shown in red). Because the assessment was based on a mean SNP proportion considering a collection of samples (in this case by tribe), it was possible that different genes were removed from different tribes. Tribes for which only a single sample was included were binned with samples that so far have not been confidently placed in a tribe (i.e. 'unplaced').

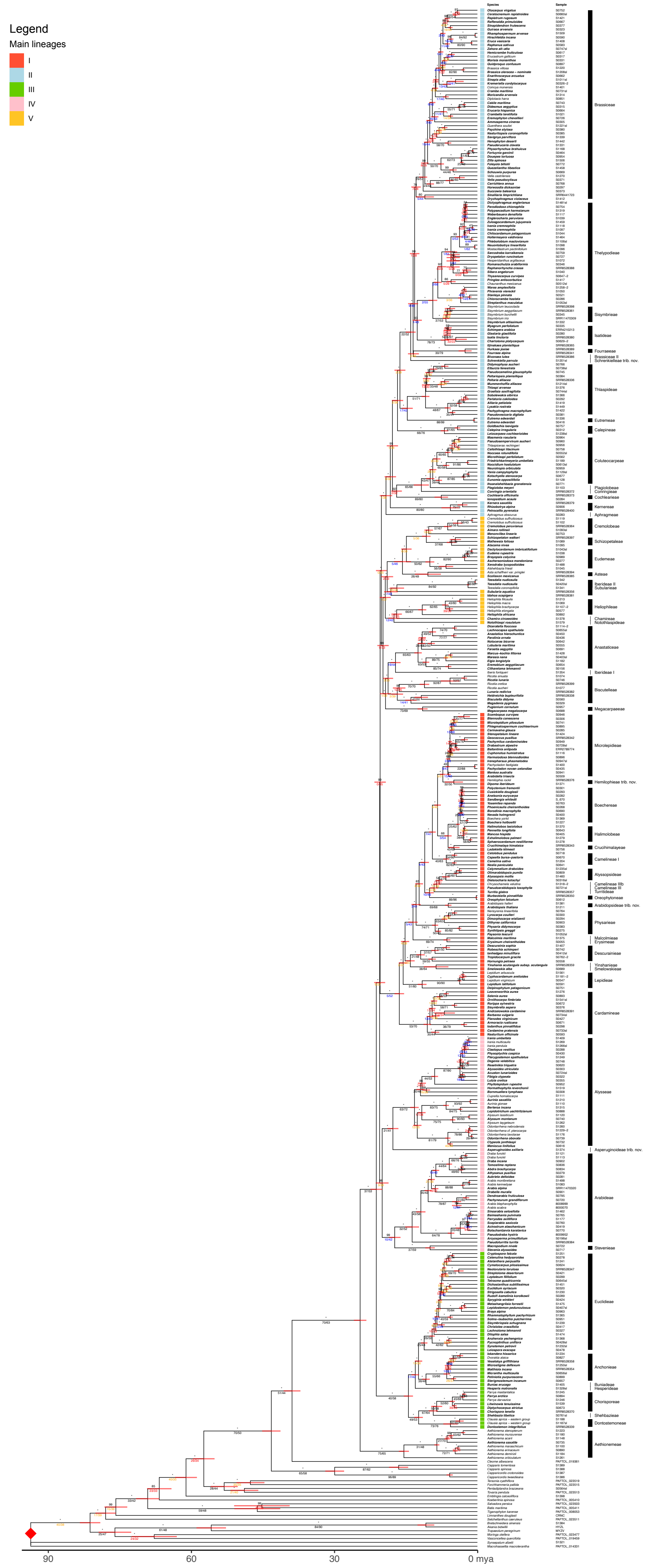


Fig. S5. Mustard family genus-level phylogeny from a maximum likelihood analysis on a 297 nuclear genes supermatrix ('superstrict routine') with node calibration of 90 mya at the stem of order Brassicales. Genus type species are highlighted in bold. This is the same figure as fig. 1 in the main text, but with full annotations. Support values: bootstrap values above branches (* for 100% support); gene/site concordance factors below branches. Red bars show 95% HPD of node ages.

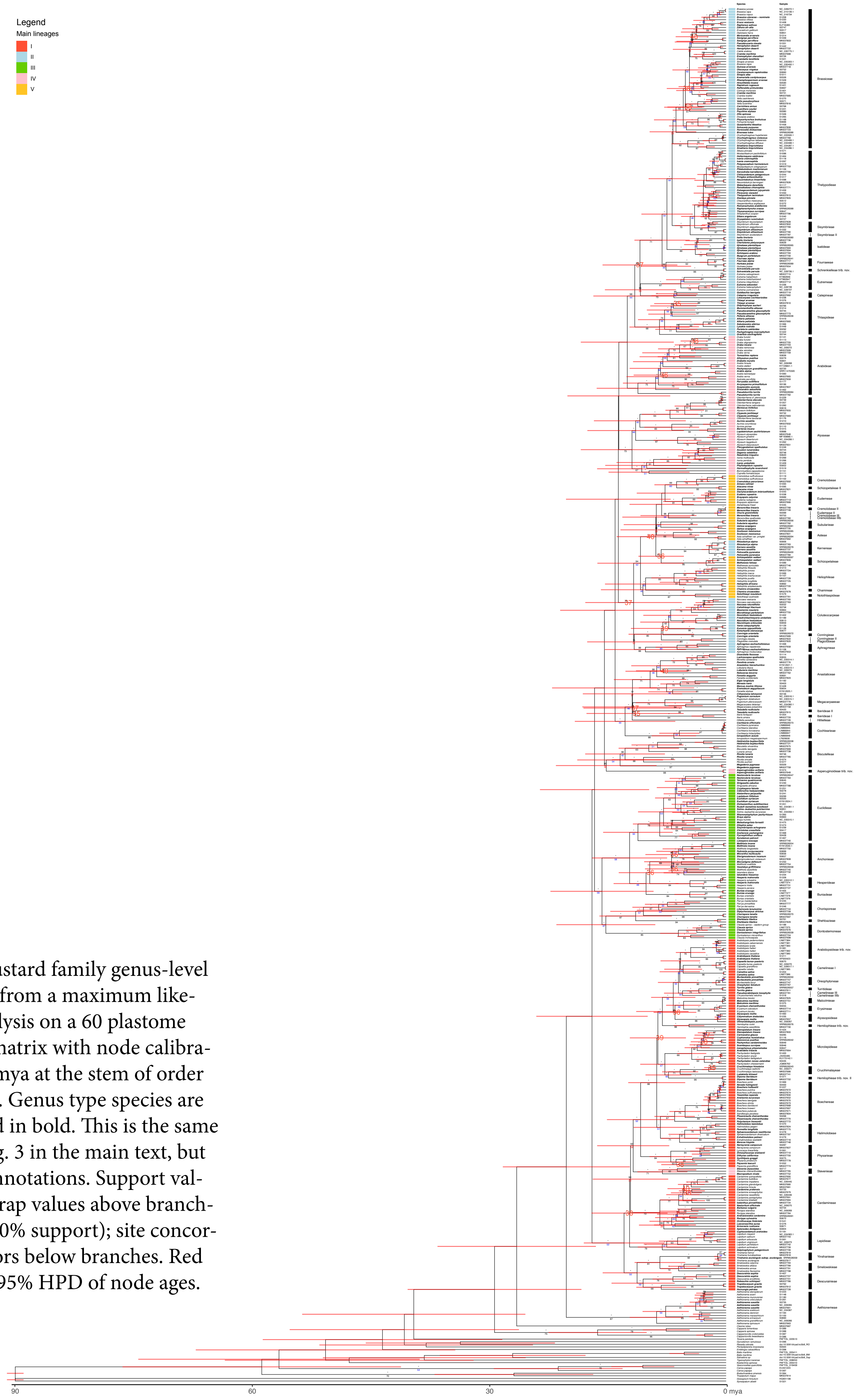


Fig. S6. Mustard family genus-level phylogeny from a maximum likelihood analysis on a 60 plastome loci supermatrix with node calibration of 90 mya at the stem of order Brassicales. Genus type species are highlighted in bold. This is the same figure as fig. 3 in the main text, but with full annotations. Support values: bootstrap values above branches (* for 100% support); site concordance factors below branches. Red bars show 95% HPD of node ages.

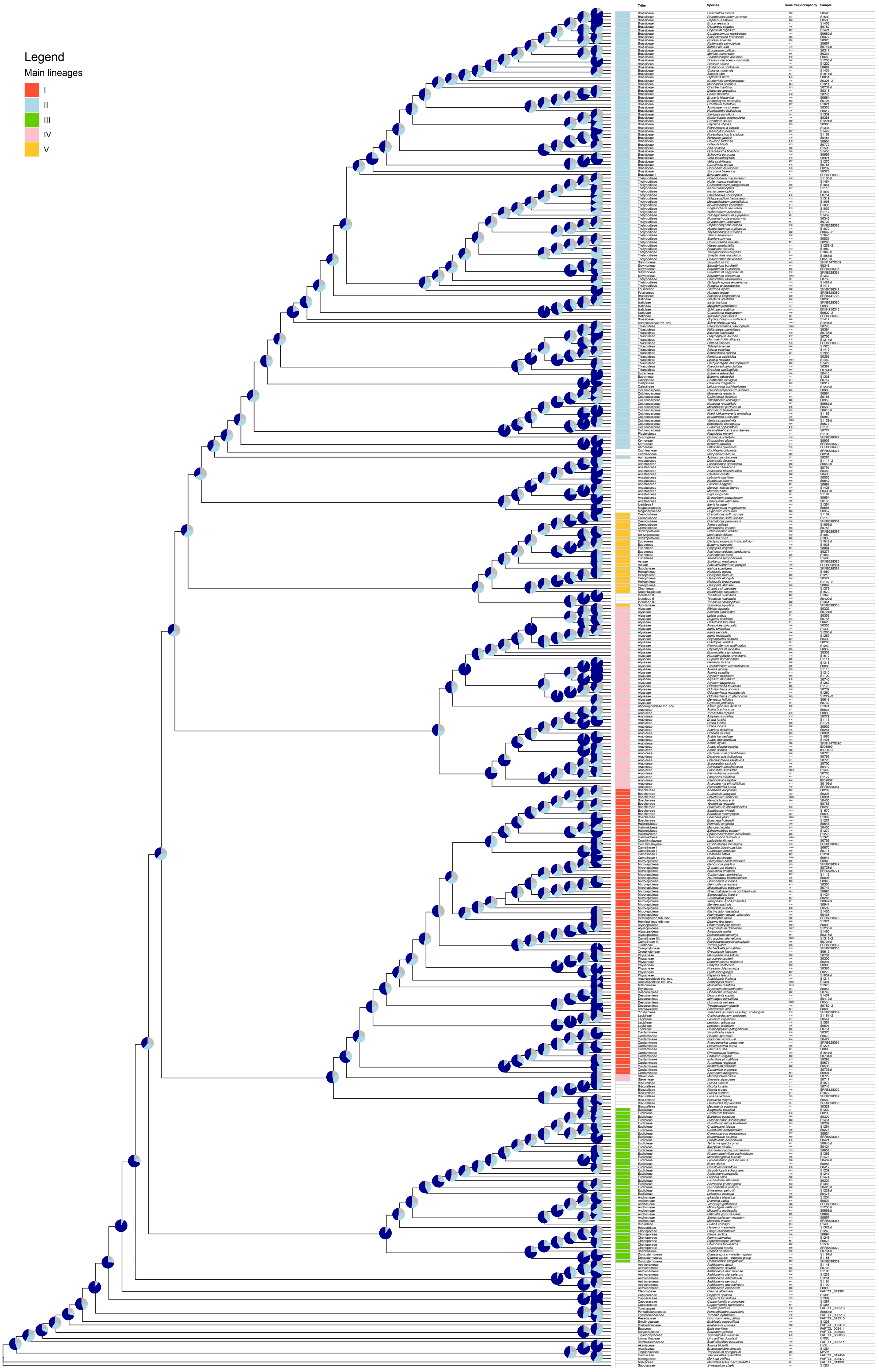


Fig. S7. Brassicaceae genus-level family phylogeny from coalescent approach with ASTRAL-III, ‘inclusive’ routine. Pie charts represent quartet scores: dark blue for the first quartet, light blue for the first alternative quartet, and grey for the second alternative quartet.

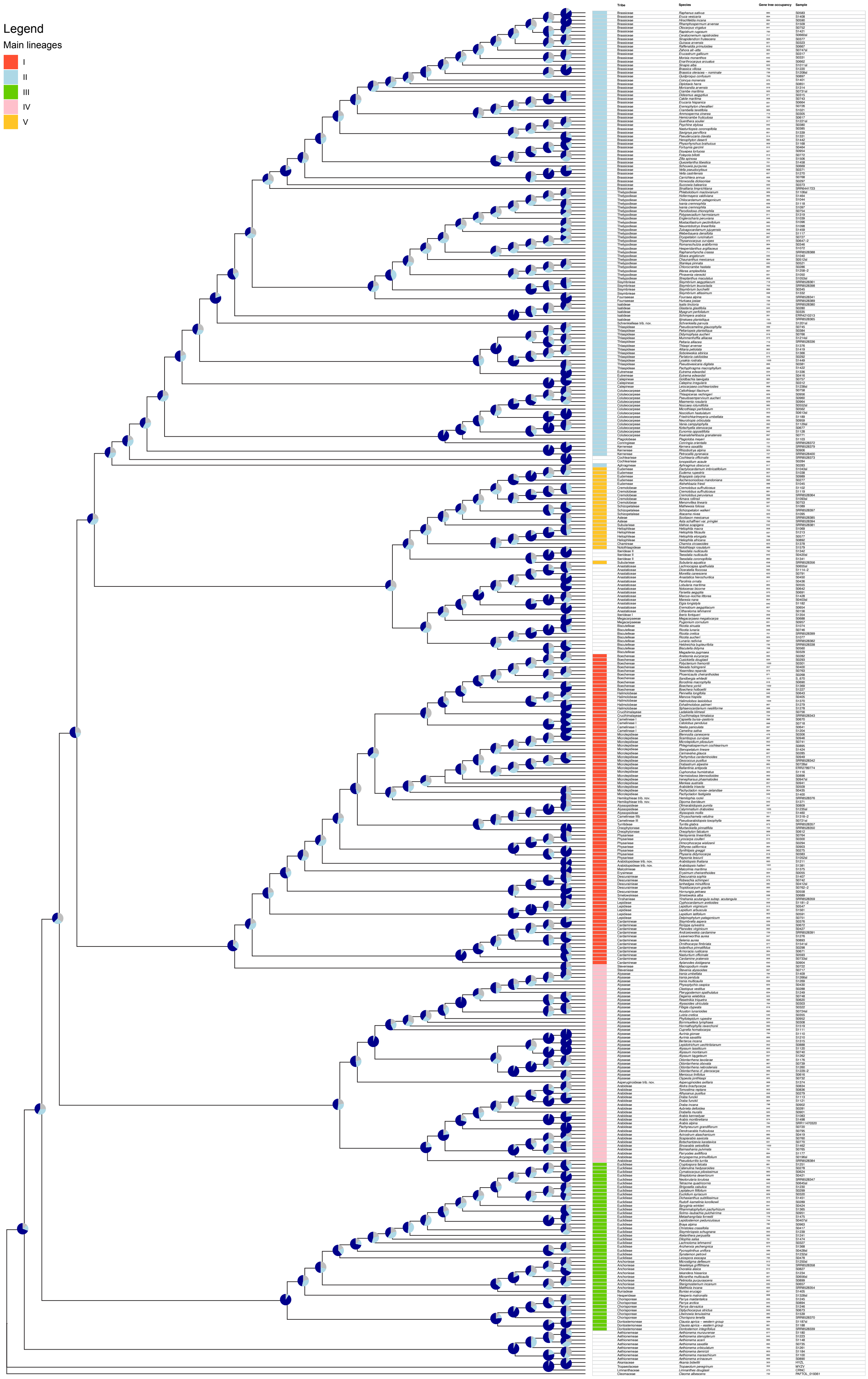


Fig. S8. Brassicaceae genus-level family phylogeny from coalescent approach with ASTRAL-III, ‘strict’ routine. Pie charts represent quartet scores: dark blue for the first quartet, light blue for the first alternative quartet, and grey for the second alternative quartet.

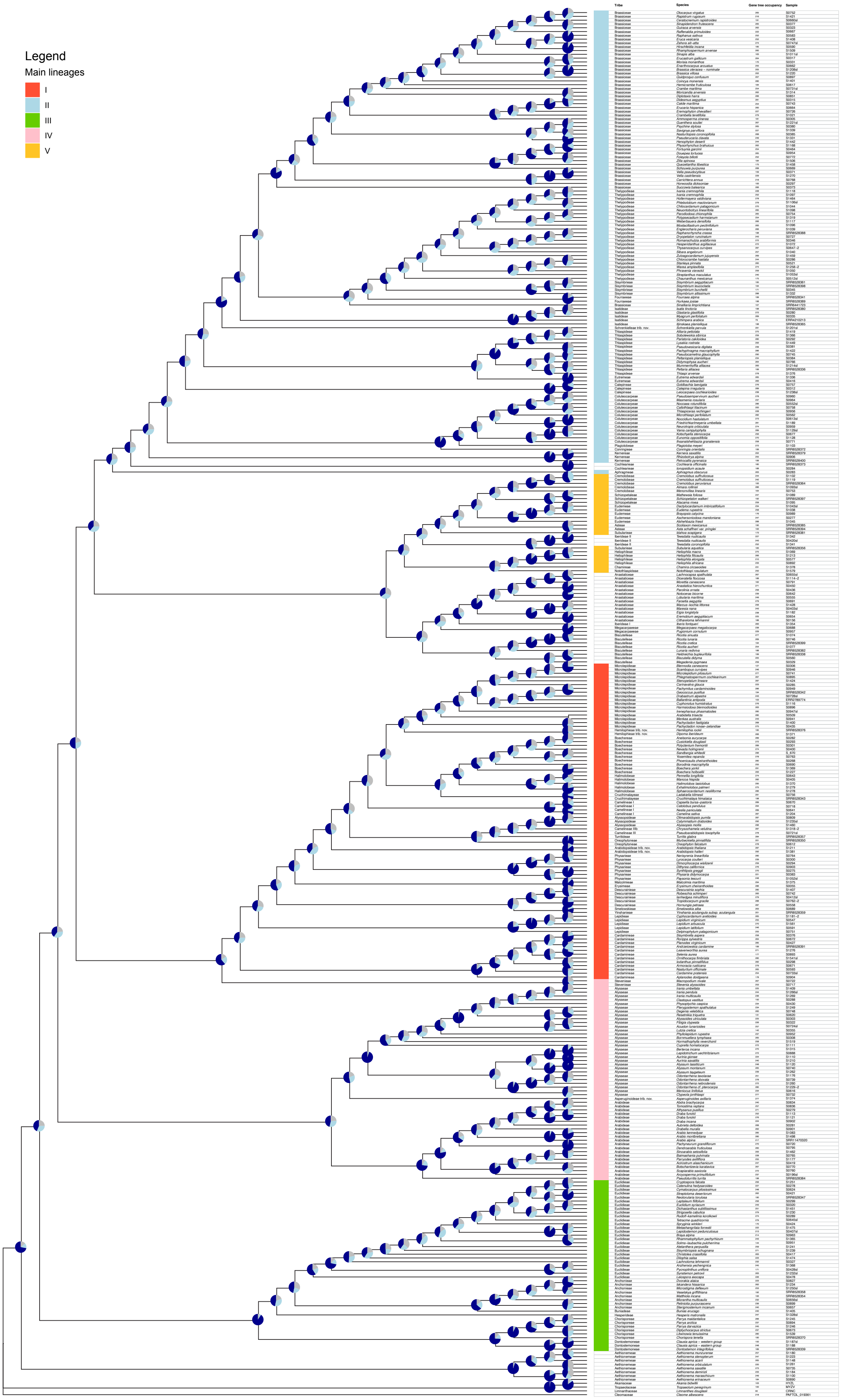


Fig. S9. Brassicaceae genus-level family phylogeny from coalescent approach with ASTRAL-III, 'superstrict' routine. Pie charts represent quartet scores: dark blue for the first quartet, light blue for the first alternative quartet, and grey for the second alternative quartet.

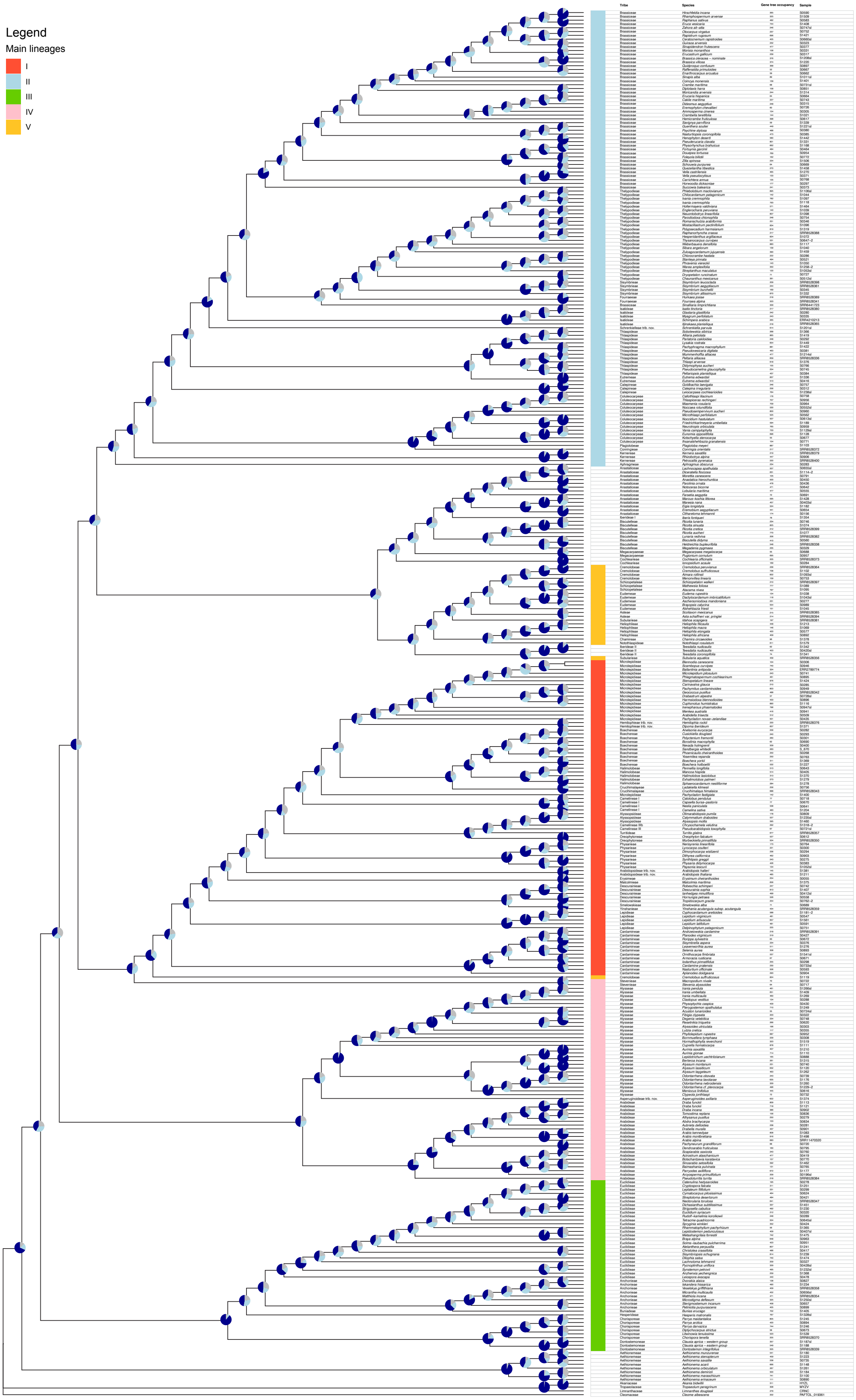


Fig. S10. Brassicaceae genus-level family phylogeny from coalescent approach with ASTRAL-III, 'super-strict-by-tribe' routine. Pie charts represent quartet scores: dark blue for the first quartet, light blue for the first alternative quartet, and grey for the second alternative quartet.

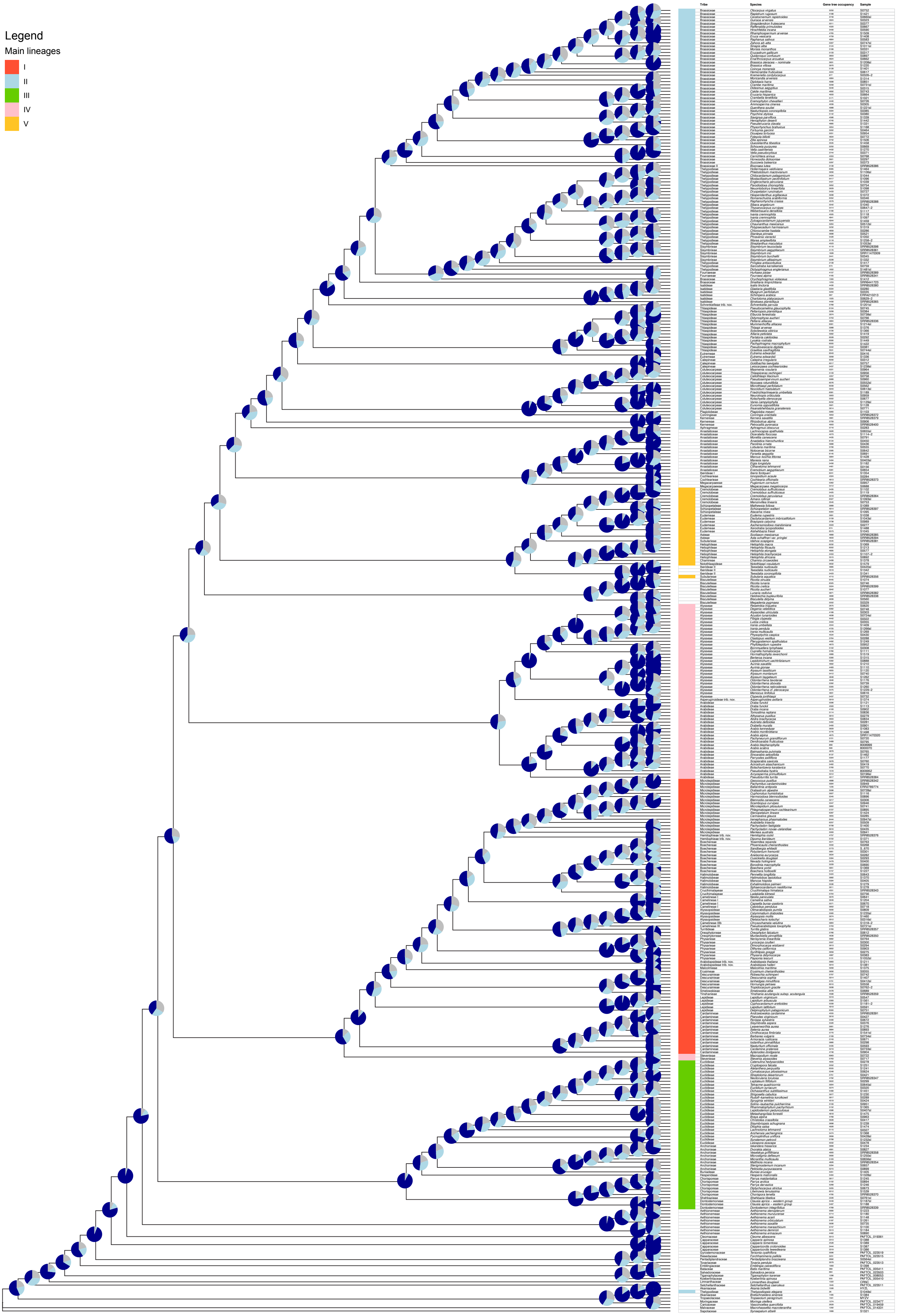


Fig. S11. Brassicaceae genus-level family phylogeny from coalescent approach with ASTRAL-Pro. Pie charts represent quartet scores: dark blue for the first quartet, light blue for the first alternative quartet, and grey for the second alternative quartet. Note that in this ASTRAL-Pro approach, gene tree occupancy can be higher than the total number of genes targeted, because for many loci multiple alleles were recovered.

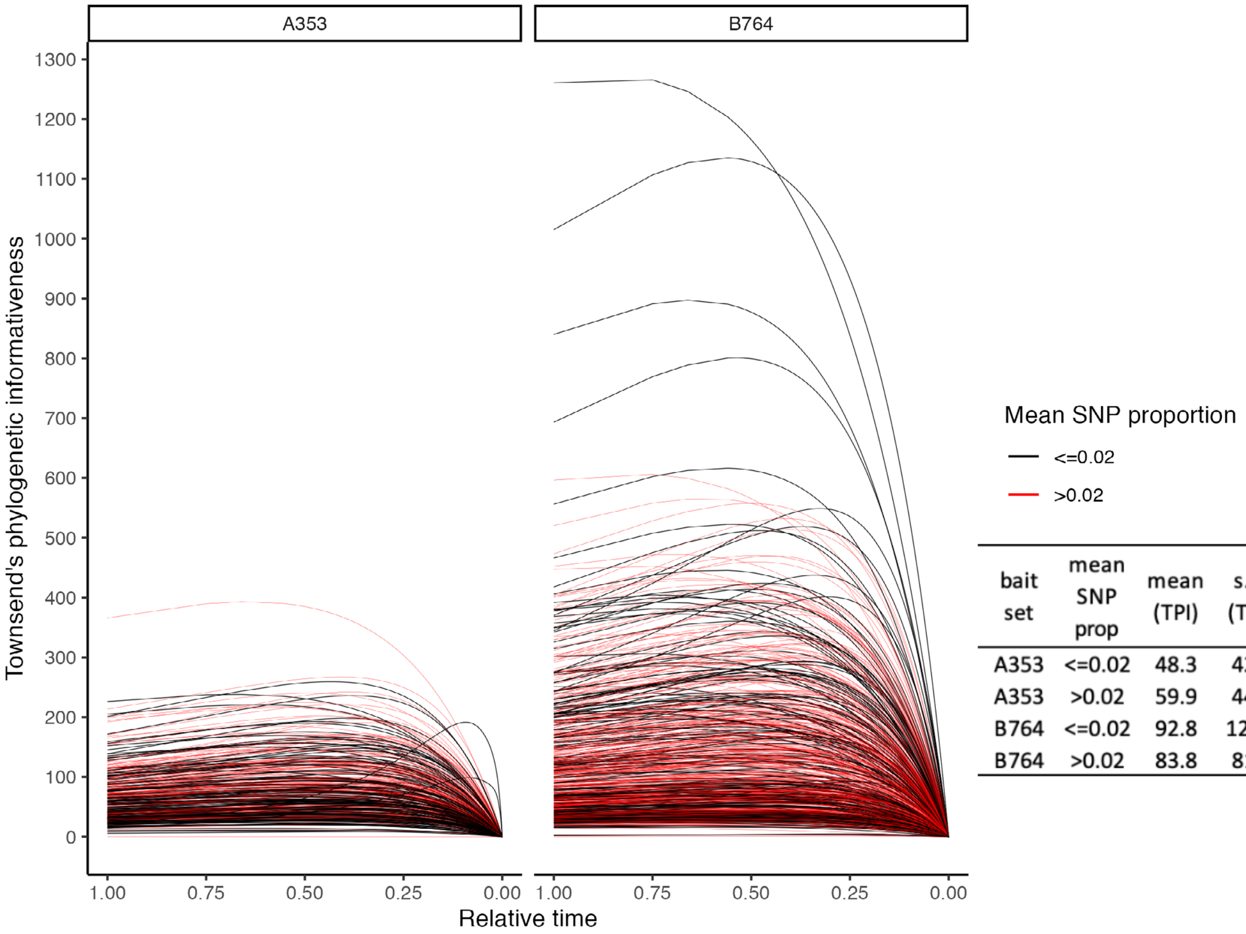


| bait set | mean SNP prop | mean (TPI) | s.d. (TPI) | n |
|---|---|---|---|---|
| A353 | <=0.02 | 48.3 | 42.4 | 125 |
| A353 | >0.02 | 59.9 | 44.8 | 191 |
| B764 | <=0.02 | 92.8 | 120.2 | 185 |
| B764 | >0.02 | 83.8 | 81.4 | 438 |

Fig. S12. Townsend's phylogenetic informativeness (TPI) for all genes, grouped by bait set (B764 vs. A353) and colored by mean SNP proportion (black for mean SNP proportion <=0.02, red for >0.02).

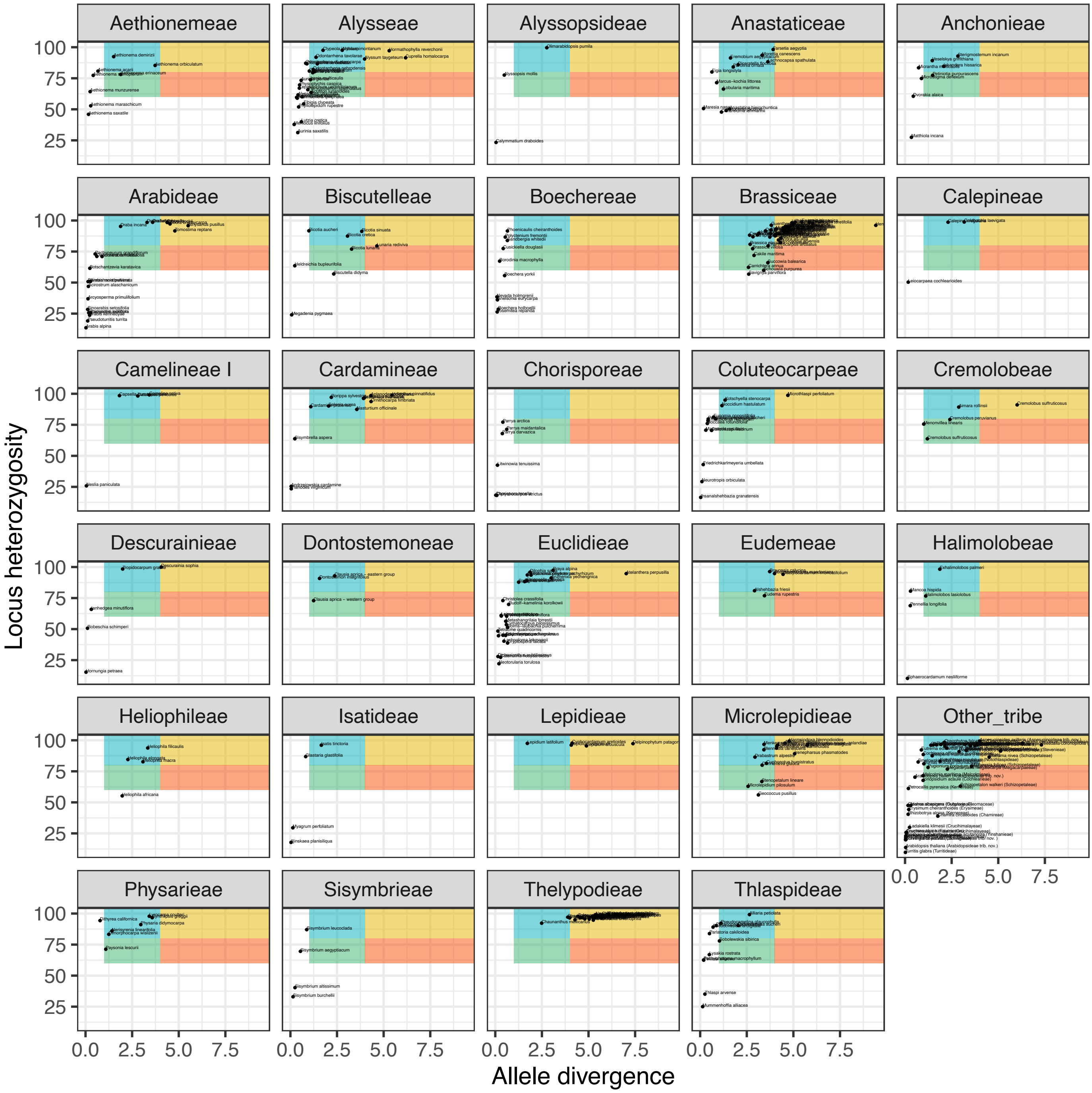


Fig. S13. HybPhaser locus heterozygosity (LH) versus allele divergence (AD). Values are means per sample over all genes available for a sample. Assessment is for the 'strict' routine. Several tribes show consistently high values, such as the Brassiceae, Lepidieae, Microlepidieae, and Thelypodieae, and are likely all from species plagued by (many) polyploidization events. To avoid LH and AD values calculated from too little data, this analysis used the 'strict' routine, meaning that samples with poor recovery (proportion of genes recovered <0.20 and/or proportion of total target length recovered <0.40) were first removed. Although it is not possible to identify polyploids based on AD and LH values alone, with numbers being much related to the family studied, tentative regions of suspected polyploids have been highlighted as follows. Green: likely old polyploids; blue: possible hybrids; yellow: highly polyploid; red: likely very old polyploids.

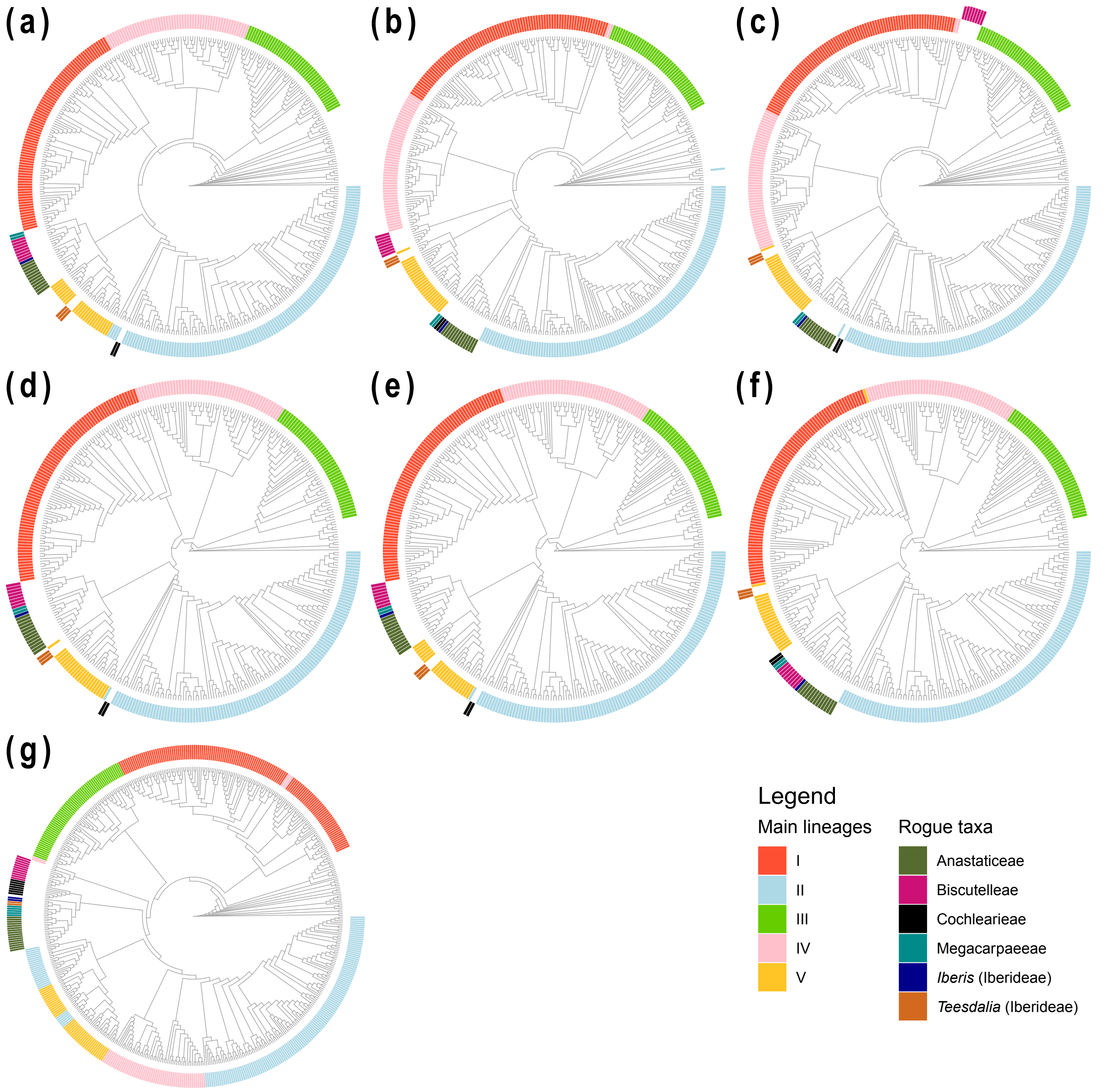


Fig. S14. Comparison of the recovery of rogue taxa in the different mustard family genus-level phylogenies. Phylogenies (a–f) based on various subsets of nuclear data, and reconstructed using different approaches; phylogeny (g) based on plastome data from (mostly) the same samples, but with additional data from Walden et al. (2020) to further complete the sampling. (a) Nuclear supermatrix maximum likelihood approach with genes from 'superstrict routine' with IQ-TREE. (b) Multiple alleles per nuclear locus coalescent approach using all genes with ASTRAL-Pro. (c) Nuclear coalescent approach with ASTRAL-III, 'inclusive' routine. (d) Nuclear coalescent approach with ASTRAL-III, 'strict' routine. (e) Nuclear coalescent approach with ASTRAL-III, 'superstrict' routine. (f) Nuclear coalescent approach with ASTRAL-III, 'superstrict by tribe' routine. (g) Plastome supermatrix maximum likelihood approach with IQ-TREE.
